# Energetic gradients emerge in developing motor-microtubule structures

**DOI:** 10.64898/2026.05.18.725774

**Authors:** Ana I. Duarte, Gabriel L. Salmon, Heun Jin Lee, Bibi Najma, Minakshi Ashok, Soichi Hirokawa, Henk W. Ch. Postma, Rachel Banks, Matt Thomson, Rob Phillips

## Abstract

Living matter produces a variety of beautiful spatiotemporal structures and patterns that are not enduringly present in their nonliving counterparts. These ordered, non-equilibrium steady states are often sustained through the consumption of energy. Here, we investigate the energetic cost of assembling an ordered aster from an initially disordered, uniform mixture of cytoskeletal microtubules and kinesin motors. Using a calibrated fluorescent ATP reporter, we measure reproducible radial ATP gradients on scales of tens of microns that establish within, and persist over, tens of minutes, alongside coupled spatial gradients in motor density. These appreciable gradients are predicted by a reaction-diffusion model that acknowledges the localization of ATP consumption to regions where both molecular motors and microtubules are sufficiently abundant to encourage consumption, as confirmed by finite element modeling. With our results, we compare the power per volume required by our cytoskeletal networks with the known power per volume expenditure in cells. Comparison of our measured results with estimates of the dissipative processes available to motor-microtubule mixtures leads to the hypothesis that maintaining spatial motor gradients dominates the energetic demand in this system. Our direct quantification of energetic fluxes across space unlocks future explorations of what steady states are accessible to cells, and how the cytoskeleton drives broad spatial organization.

**Significance:** How much energy do organisms pay to form and maintain their organizing biochemical patterns? Existing measurements of cellular metabolism and energy expenditures largely resolve net or supply-side biochemical fluxes, without spatial information, impeding the study of this basic question. Here, we develop an experimental approach to directly measure the distributions of biochemical energy that respond to power expenditures of cytoskeletal motor-microtubule networks as they form aster structures, reminiscent of those found in the mitotic spindle. As these structures self-assemble, calibrated readouts in real molecular units register large, reproducible, and long-lived gradients of ATP. We interpret these measurements by developing theory to account for the functional destinies of energy expenditure. These advances clarify outstanding questions of energy in living matter.

---

**A** key driver of the rich patterns in living matter is a steady investment of energy. For example, the establishment of morphogen gradients in developmental patterning arises from its synthesis, spatial redistribution, and degradation (1–4). Similarly, cytoskeletal motor systems hydrolyze adenosine triphosphate (ATP) and other triphosphates to achieve processes ranging from intracellular transport, to cell motility, to chromosome segregation. Motivated by these processes, we were inspired to develop a physical understanding of the connection between energy fluxes and the emergence of biological order in space and time in the context of the particular example of microtubule-motor assemblies.

The energetic basis of the processes within living cells are based upon a few fundamental energy currencies, which can be thought of as biological batteries. This metaphor is useful because it reminds us that batteries are indifferent to the particulars of what they are wired up to—they can drive anything from the light in a flashlight to motorized toys. Biological processes are powered by several key biological batteries including membrane potentials, redox reactions and trinucleotide hydrolysis. Indeed, for the molecular motor driven reactions that power the structures of the cytoskeleton, ATP and GTP hydrolysis are central. Thus, we were curious about how ATP consumption in space and time drives the dynamics of structure formation.

Recently, it has become possible to measure the total energy consumption of both living organisms and the molecular machinery that drives them (5–9), joining theoretical methods to infer energy expenditures from observables (10). Despite the foundational value of these investigations, to date almost all these measurements report such metabolic expenditures as spatial and temporal averages. These empirical limitations mean that putatively profound heterogeneities in expenditures across regions and processes of the cell remain unknown and unmeasured.

How significantly does dissipation localize in space and time among biomolecular assemblies? How big and sustainable are bulk energetic and dissipative gradients sculpting living matter? To address these open questions and complement earlier foundational studies, we set out to make precise, real time measurements of energy abundance and consumption resolved on the micron scale. In particular, here we report the visualization of spatial ATP concentration gradients across cytoskeletal networks, giving insight into how structure, composition and morphology drive energy dissipation.

We use motor-microtubule assemblies as a highly controllable and tunable system due to their minimal components and self-organizing properties *in vitro* as well as their biological ubiquity (11–16). An abundance of work has established that connected dimeric motor domains can cross-link microtubules, creating ordered networks (11, 12, 17, 18). To control the position, size and start of microtubule cross-linking, we optogenetically link motor proteins together as shown in Figure 1, as previously developed in our labs (18). Motors harness energy to drag microtubules into ordered structures by hydrolyzing an ATP molecule for each step they take along the microtubule. We measure the energy consumed by the motors throughout space and time using a fluorescent, QUEEN-based ATP probe (19). We illustrate the ratiometric probe mechanism shown in Figure 1(B).

**Fig. 1.**
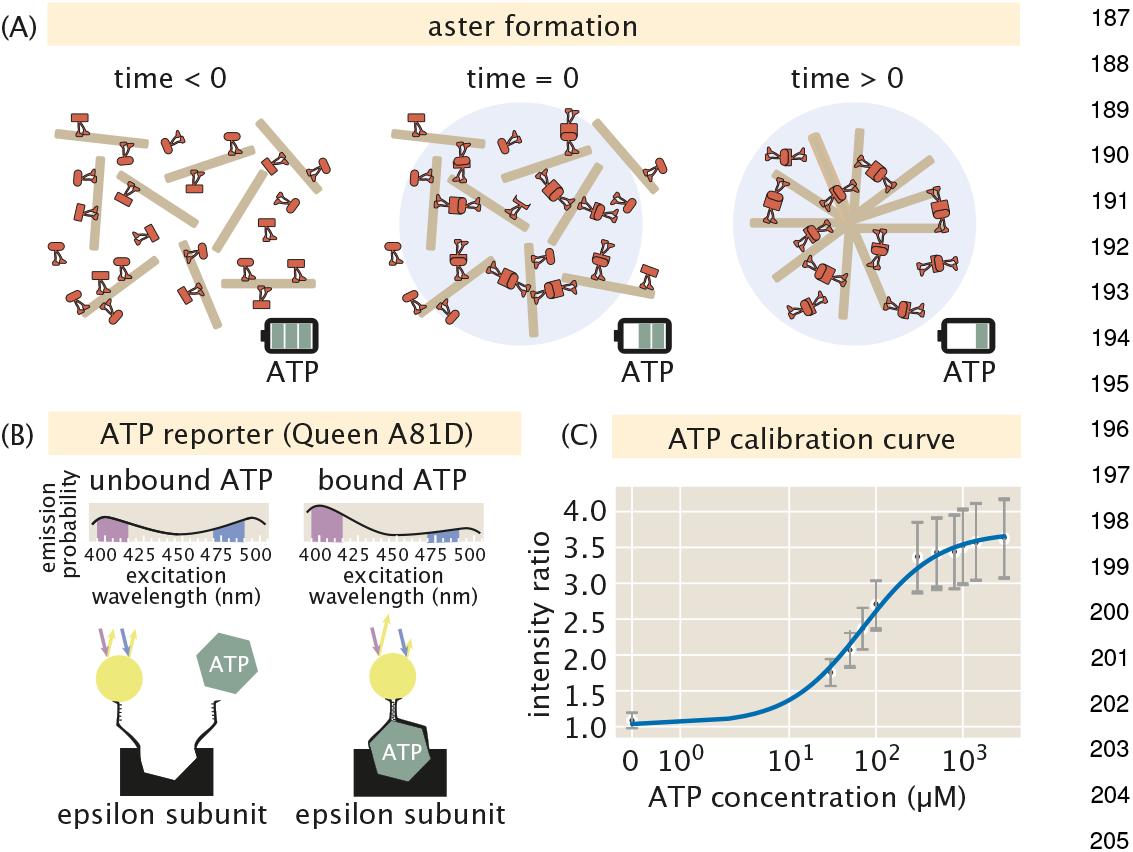
Schematic of the experimental system used to measure spatiotemporal evolution of ATP. (A) The formation of an aster using light activated motor dimerization. Before light activation, motors independently walk on microtubules, uniformly hydrolyzing ATP. At *t* = 0 a circular light pattern is projected onto the sample. Motor proteins inside the illuminated region dimerize, crosslinking microtubules. As time elapses, microtubules are dragged into an aster resulting in spatially heterogeneous depletion of ATP. (B) The binding mechanism for ATP to the ATP probe. Binding ATP to the probe causes the number of emission counts due to an excitation of 405 nm light to increase, while emission counts from 480 nm light excitation decreases. By comparing the ratio of the emission counts at 405 nm and 480 nm light excitations, the concentration of ATP can be inferred. (C) A calibration curve mapping known ATP concentrations to fluorescent light intensity ratios. Each black circle represents the mean ratio value for a given image and gray error bars report the standard deviation of the image.

In addition, to complement the measurements and to provide a framework for understanding them, we examine a reaction-diffusion model that describes the emergent ATP spatiotemporal gradients and explore the implications of that model with finite element calculations. Combining our experimental and theoretical results, we determine that the measured network formation power is indeed many orders of magnitude greater than the theoretical power of equilibrium processes.

## Results

### Direct measurement and visualization of emergent ATP gradients

The key elements of our experimental design are shown in Figure 1. As noted above, using spatially and temporally controlled illumination, we can generate patterns such as the radially symmetric aster shown in the schematic. Our principal experimental goal is to measure the rate of consumption of ATP as a function of position and time, a goal that is realized by using the fluorescent, ratiometric ATP reporter (19) depicted in Figure 1(B). The probe mechanism creates a change in the protonation state (20) of the fluorophore when ATP binds (21), triggering a shift in the fluorophore’s absorption spectrum (22). By using known standards, as shown in Figure 1(C), we construct a calibration curve that permits us to measure the ATP concentration in a given spatial region. Given that the characteristic scale of ATP concentrations in our experiments are of order a few hundred *µ*M, our calibration curve, in Figure 1(C), shows that the ATP reporter is sensitive in precisely the concentration range needed to spatially resolve ATP consumption in the system. The Supplemental Information, provides a detailed description of how we handled uneven illumination (section S2.B)and photobleaching (section S2.C), a prerequisite to generate such a calibration curve. There we also describe how we used two-dimensional images to infer the properties of our three dimensional system (section S7).

This measurement scheme equips us to simultaneously resolve the quantity of both ATP (the “fuel”) and molecular motors (the “consumer”) over space and time while asters form, as shown for representative time courses in Figure 2 (Top) and (Bottom). Local motor concentrations in space are inferred from fluorescence footage using a separate calibration curve, as discussed further in the Supplementary Information (section S2.D). As motors step along and exert torques on microtubules, they accumulate in the centers of assembling asters, reaching a peak concentration of *≈*3 *µ*M, creating self-organized polar order and material flow, as witnessed in the time course of Figure 2(Top). Concurrently, our measurements report how an initially uniform concentration of ATP is steadily reduced in time, depleting from an initial concentration of 500 *µ*M to around 350 *µ*M in the first four minutes, to *≈*100 *µ*M in about twenty minutes, as shown by Figure 2(Bottom). Importantly, the depletion of ATP is manifestly nonuniform over space, forming a steepening gradient of ATP, with less ATP in the aster center than at its periphery. These concentration fields are displayed as radial profiles in Figure 3(A) and Figure 3(B), registering rich time and space dependencies. Specifically, conservatively summarizing ATP gradients by their change in concentration across a one hundred micron radial region near the edge of an aster (between radial positions *r* = 200 *µ*m near the edge and *r* = 100 *µ*m) registers gradients of 0.5 *µ*M*/µ*m to 1 *µ*M*/µ*m across replicate asters (see Supplementary Material section S3).

**Fig. 2.**
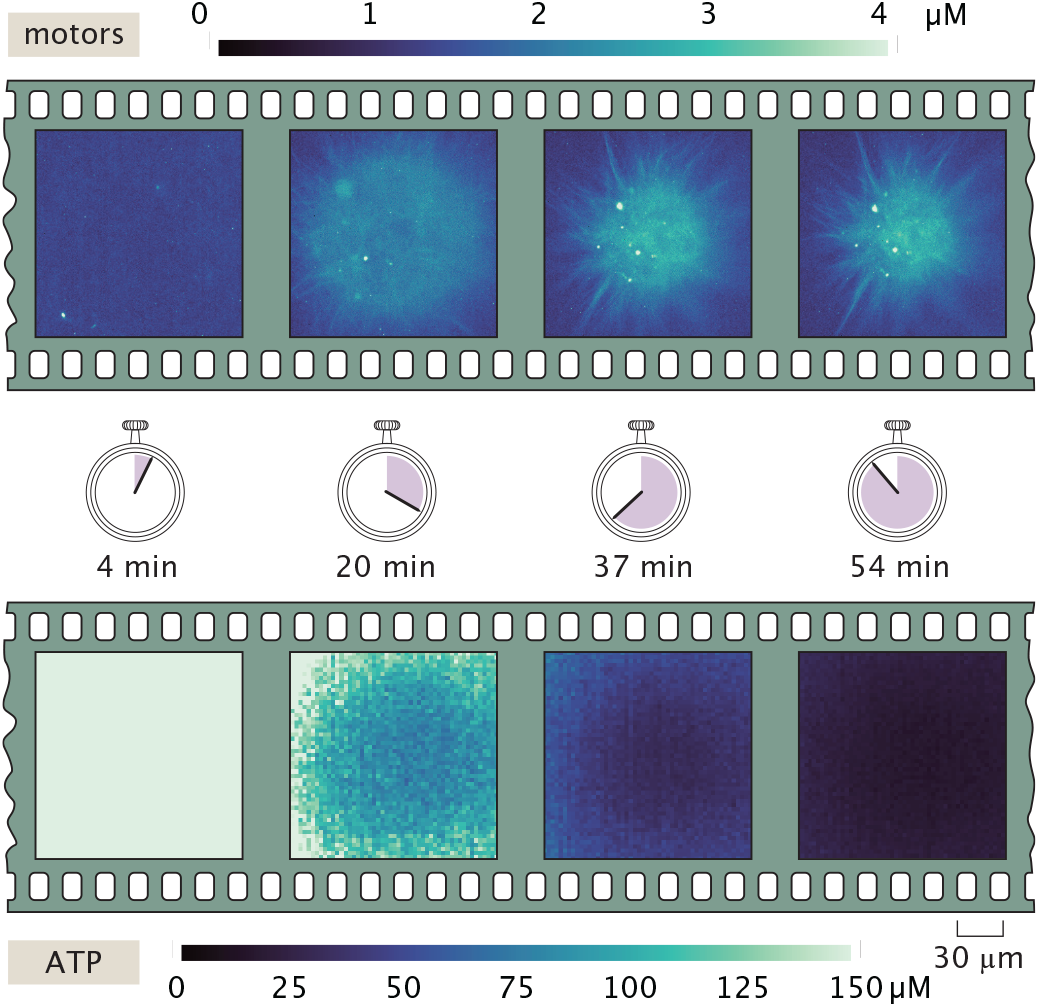
Experimental measurements resolve coupled gradients of motors and ATP across space and time. (Top) Experimentally measured spatial distributions of molecular motors and (Bottom) ATP over four time points during the self-organization of an aster. As time evolves, motor proteins concentrate near the aster center; a coupled ATP gradient develops, with greatest depletion in the aster’s center where motors are most abundant.

**Fig. 3.**
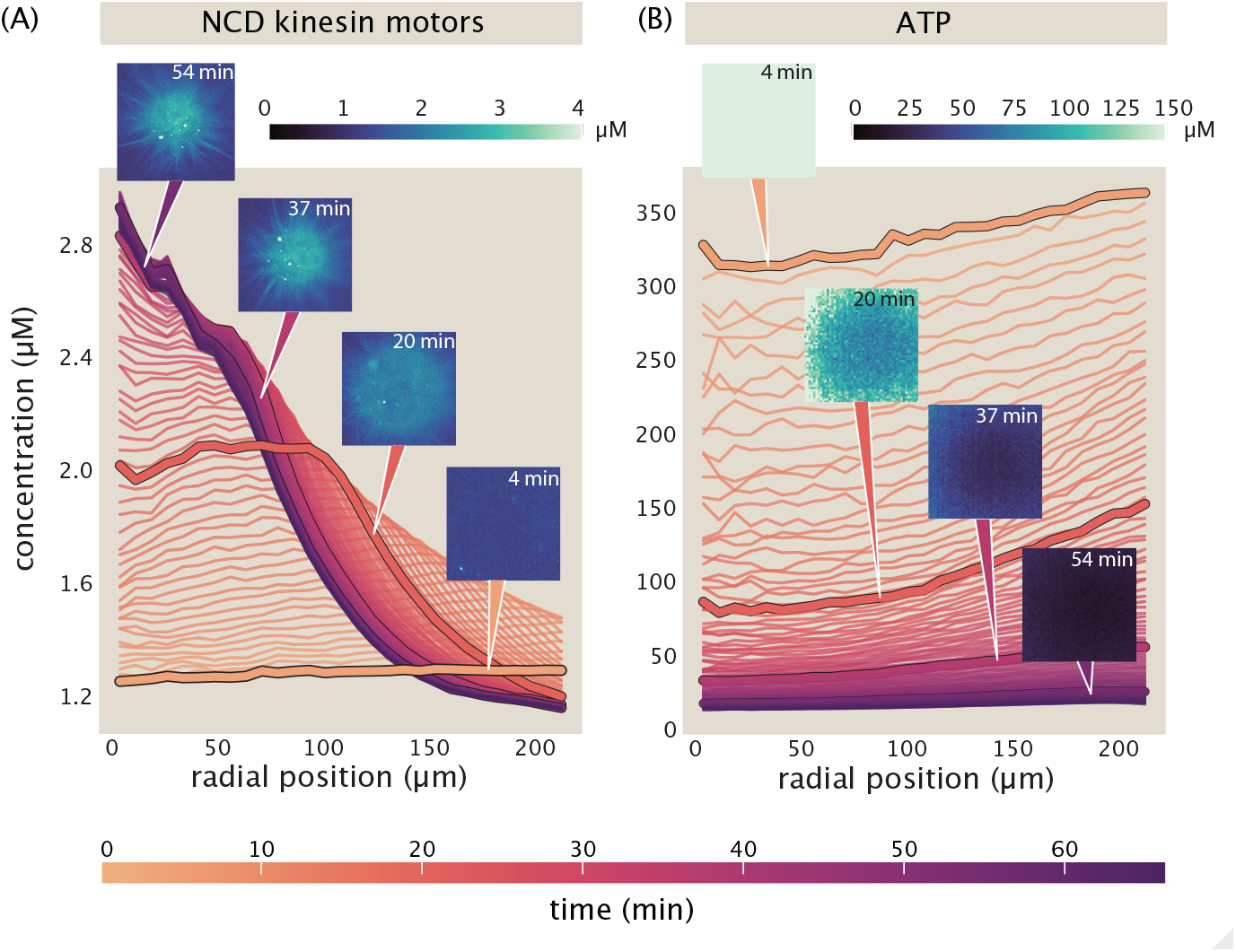
Measurements of developing ATP and motor gradients with respect to radial position in asters. (A) Azimuthally-averaging the data shown in Figure 2(Top) gives radial concentration profiles of motors. (B) Similarly, azimuthal averaging of the data shown in Figure 2(Bottom) yields the radial concentration profiles of ATP. These gradients reveal clear, rich nonuniformities over time and space. Further replicate aster experiments show largely coherent behaviors, with scales of variation across replicate experiments visible in the Supplementary Information section S3B.1

In general, the uncertainties in such interesting physical scales (such as gradients and dissipations) reported in this paper are most earnestly captured by their aggregate variations across many separate aster formation experiments, as detailed in the Supplementary Information, section S3, Figures S25-S28.

### Appreciating the biological magnitude of gradients

Are the spatial gradients in ATP reported by our calibrated measurements in Figure 2 and Figure 3 with a characteristic scale of *≈µM/µm* biologically meaningful? To develop intuition for that question, we examine how the measured ATP concentrations vary relative to the *K*_*M*_ of the motors. At intermediate times, as highlighted by Figure 4(A), ATP drops from a high value of *≈*100 *µ*M ATP that is several times larger than the characteristic concentration scales governing motor stepping activity (namely the enzymatic Michaelis-Menten constant *K*_*M*_ *≈* 23 *µ*M ATP, as reported by steady-state ATPase assays (23, 24)). Further, at sufficiently late times, ATP depletes sufficiently so as to be above this characteristic *K*_*M*_ at the edge of the aster but below this *K*_*M*_ in the center of the aster (as illustrated by the 37 min pink-fuchsia inset of Figure 3(B)).

**Fig. 4.**
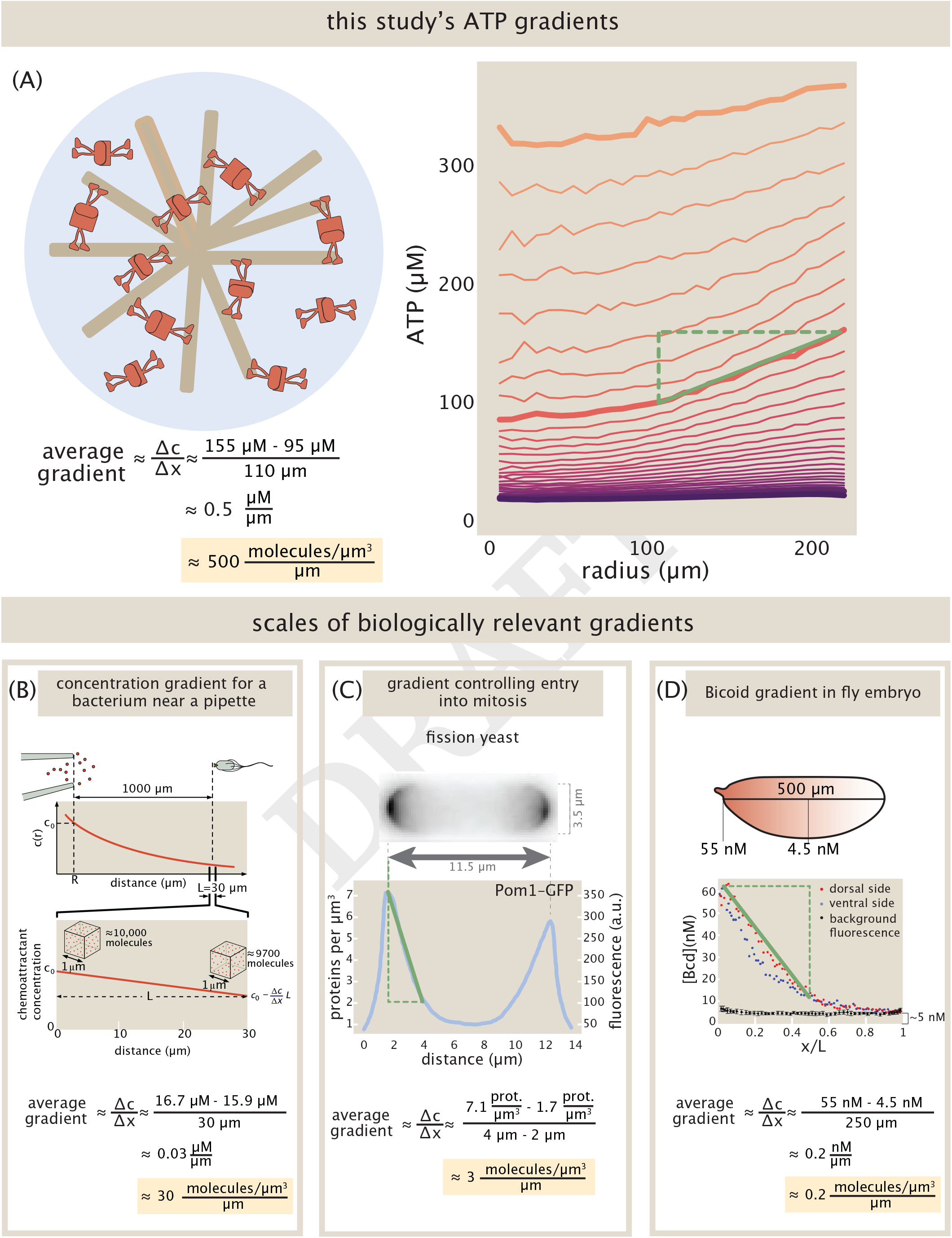
Appreciating the relative scales of biologically developed gradients. (A) Our measured ATP gradients experience approximately a factor of two depletion across the scale of the aster length. (B) Bacterial cells respond to gradients of chemoattractants (Plot based on V. Sourjik et al. (25)). (C) In fission yeast, the protein kinase Pom1 localizes at the cell poles creating steep gradients on each side of the cell. (Plot adapted from J. Moseley et al. (26).) (D) The *Bicoid* gradient organizing early fly development manifests the majority of its decrease on the scale of half its body length. (Plot adapted from T. Gregor et al., (29).)

Thus, the spatial variation of ATP inside a single aster lies squarely in the functional range that controls whether or not motors have enough ATP to move close to their maximum stepping rates.

Another way to size up these gradients is to compare them to other important gradients as we detail in the Supplemental Information section S10. Figures 4(B), (C), and (D) provide several candidate reference points. In Figure 4(B), we illustrate a classic experimental study (25) on bacterial chemotaxis in which bacteria navigate gradients of approximately 30 (molecules/*µ*m3)/*µ*m (25). As seen in Figure 4(C), the protein kinase Pom1 in fission yeast is part of the signaling pathway that controls entry to mitosis, Pom1 localizes in the poles of the yeast cell (26). We estimate the gradient in these yeast cells is approximately 3 (proteins/*µ*m3)/*µ*m. As another example, we consider the transcription factor *Bicoid*, which famously contributes to delivering the positional information that organizes the fruit fly *Drosophila*’s body plan, as shown in Figure 4(D) (2, 27–29). This molecular gradient develops with high precision across embryos, and robustly adopts a characteristic profile with a gradient of *≈*0.2 (molecules/*µ*m^3^)/*µ*m. Thus we see that the maximal ATP gradients we measure are ten to a few thousand times steeper than the chemoattractant, Pom1 and *Bicoid* gradients used to drive bacterial chemotaxis, cellular division and sculpt embryos, respectively. (Further numerical discussion, including complementary notions of steepness, is found in the Supplementary Information section S10.) Accordingly, these measurements attest that such gradients could have biologically salient impacts.

### Mapping power consumption in space and time

Another way of visualizing the results of our measurements is to take the ATP data from successive instants and convert it to a power. As shown in Figure 5, the power can be evaluated directly in units of ATP/s, revealing power of order few *×*10^8^ ATP*/*s. This magnitude of power expenditures is reproducibly observed across replicate asters as described in detail in the Supplementary Information, section 3B.2, Figures S25 and S29; peak dissipation rates usually occur at early times, and vary between *≈*3 *×* 10^8^ ATP*/*s and *≈*8 *×* 10^8^ ATP*/*s across separate self-organizations of distinct asters. We interpret this few-fold variation in the dissipations across replicate aster experiences as representative of aggregate uncertainty in these scales (see SI).

**Fig. 5.**
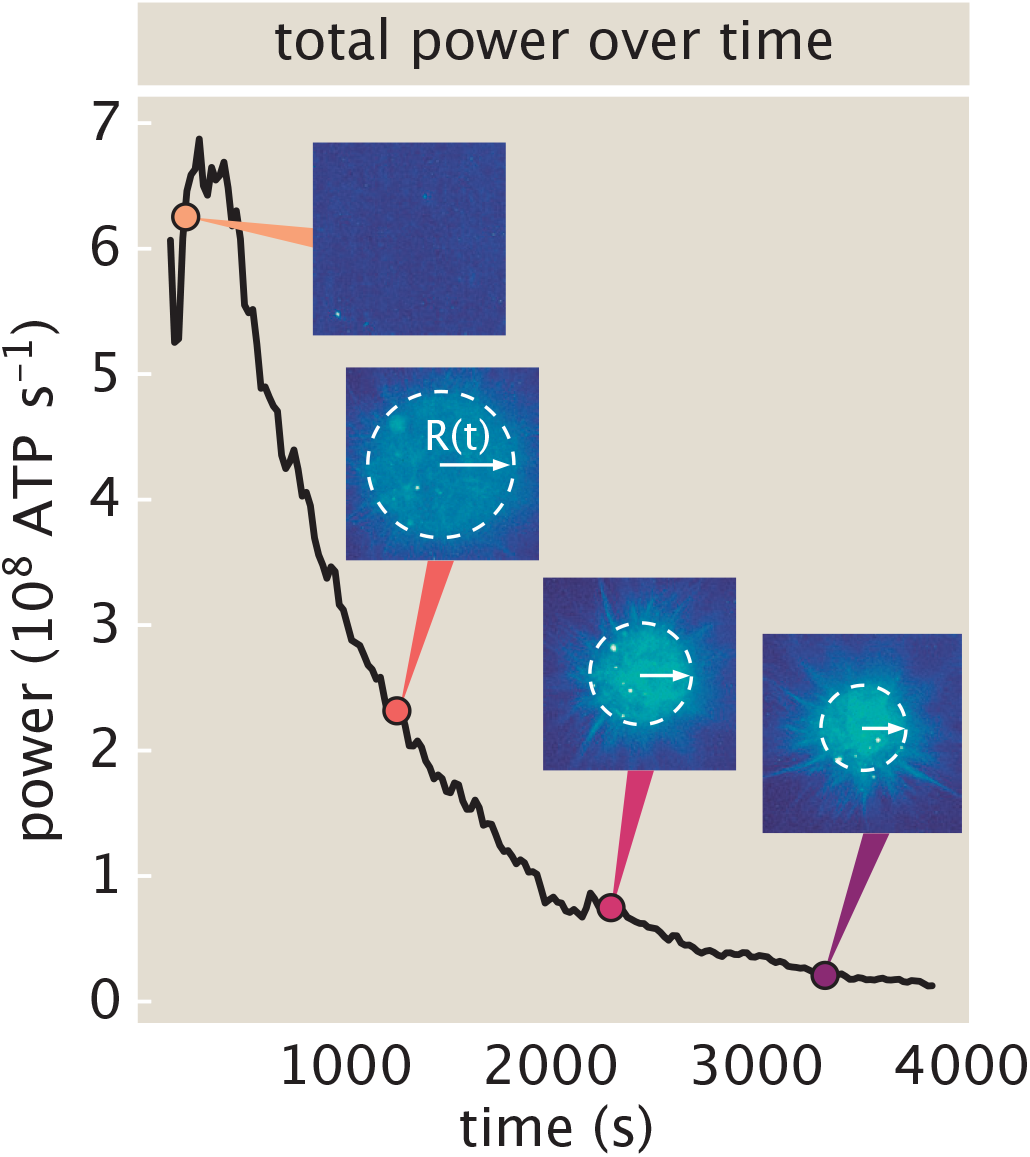
Experimental readouts of total power during aster formation. Time derivatives of ATP reflect local and global power consumption, acknowledging the imaging geometry. The number of ATP molecules consumed per time over the whole aster volume significantly changes over time. Under these conditions (1.2 *µ*M motor proteins and 500 *µ*M initial ATP), the magnitude of power tends to follow the size of the aster. Initially, while the aster is most rapidly contracting, ATP is consumed most rapidly, then approaches a baseline power level once the aster is no longer dramatically changing in size. This trend is visible in the majority of aster replicates which can be seen in the Supplementary Information Figure S29.

Note that these changes in the ATP present in an aster over time technically reflect both a consumption power inside the aster’s volume, and any (not explicitly measured) replenishment flux of ATP into the aster due to diffusion or fluid flow. In the present setting, such replenishment fluxes should counter consumption: since ATP increases with radial position (including at the aster’s boundary), average diffusive flux directs inwards (by Fick’s law), and net fluid flow is also inwards in this contractile setting (see Supplementary Information section S2J). Accordingly, the net ATP consumption rates reported by measurements here may be regarded as reasonably close lower bounds to full underlying consumption rates.

These real thermodynamic units of our calibrated measurements allow us to compare our total measured power with estimates of what the energy from hydrolysis events is used to pay for. In addition, we can directly contrast these measurements with power expenditures reported in far different biological contexts. We return to reckon with what sets the scale of these measured power values in the final *Results* section. Next, we ask how one might develop quantitative intuition for the distribution of the ATP in the aster in both space and time.

### Understanding measured gradients with reaction-diffusion modeling

The profiles revealed in Figure 6 characterize the radial and temporal dependence of both the motors and the ATP. We present a dynamical equation in Figure 6(A) which acts as a minimal model to describe the resultant ATP profiles through space and time. The rate of change of ATP in a small material volume element can be attributed both to ATP molecules entering and leaving that small region (diffusion) and to the hydrolysis of those ATP molecules by molecular motors that are in the material volume element of interest (reaction). The reaction term is defined as the product of the hydrolysis rate of a motor protein, the motor profile (which gives the density of motors in that material volume element), the bound fraction of motors to microtubules, and the probability an ATP is bound to a motor. The bound fraction of motors to microtubules is described by a kinetic model of binding as derived in the Supplementary Information section S8A. The probability an ATP is bound to a motor protein is determined by Michaelis-Menten-like dynamics for the rate of ATP consumption by motors. However, the denominator of that term also includes terms that reflect competitive inhibition of the reaction due to ADP and P_*i*_. Note that there is another more pernicious dynamic taking place during our experiments, namely, photobleaching. Section S6 in the Supplemental Information describes how we measure and account for photobleaching, including through finite element simulations of a diffusion-photobleaching equation in section S6F.

**Fig. 6.**
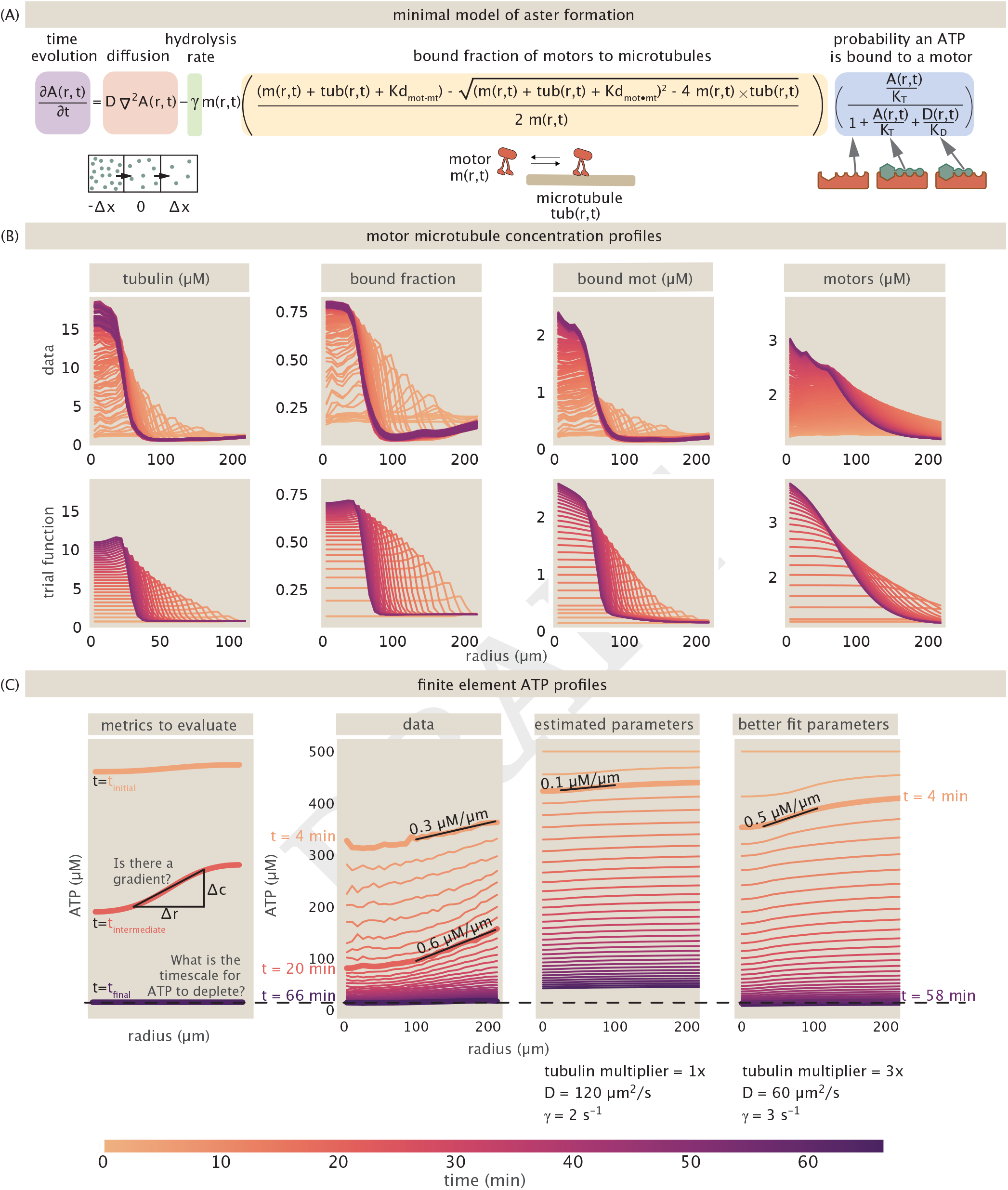
Minimal mathematical model and finite element simulations of ATP concentration in space and time. (A) The reaction-diffusion equation used to simulate ATP concentration is written with illustrations of the diffusive term, the binding states of motors to microtubules, and the binding states of ATP (*A*) and ADP (*D*) to the motor protein. (B) The distribution of microtubules and motors as a function of distance from the center of the aster as a function of time show the development of a gradient over time. The bottom row shows the empirical and approximate fits used as input into the finite element calculations. The key point of the motor and microtubule temporal distributions is that they are used as input into the consumption-diffusion model for the ATP. (C) We compare the gradients produced by finite element simulations on two metrics: how quickly the ATP depletes and the slope of the largest gradient. Next to the data, we plot the simulation using our best estimated parameters, namely a diffusion constant of 120 *µ*m^2^*/*s and a hydrolysis rate of 2 s^*−*1^. Additionally, the inputted tubulin profile is the trial function profile in panel (B). The right most plot shows parameters that better satisfy our evaluation metrics using a diffusion constant of 60 *µ*m^2^*/*s, a hydrolysis rate of 3 s^*−*1^, and multiplying the maximum value of the tubulin profile by a factor of 3.

One powerful way to analyze the solutions to equations such as that shown in Figure 6(A) in diverse geometries is by appealing to numerical methods. In our case, we used the finite element method to compute the space-time history of ATP as shown in Figure 6(C). To generate these plots, we input trial functions for the time-dependent motor and tubulin profiles, which are based on measured data. The form of these profiles are discussed in detail in the Supplementary Information section S4A.4 and are depicted in Figure 6(B), along with the resultant bound fraction of motors and bound motor concentration across time and space. To assess the quality of our simulation, we consider two semi-quantitative checks: does the global concentration of ATP deplete on a similar time scale to that measured in the experiment and is the size of the maximum gradient comparable to the maximum gradient in the experimental profiles? In Figure 6(C), the left plot illustrates these two dynamical metrics. The second graph to the left plots the experimental ATP profiles, which we note show a maximal slope of *≈*0.6 *µ*M*/µ*m and globally depletes in *≈*66 minutes. The two plots on the right show the results of our simulation. The second plot from the right uses our best estimates for the parameters that we believe apply to our experiment, which we expound on in the Supplementary Information section S4A. For this particular choice of parameters, the simulated profiles have a much shallower gradient and globally deplete ATP slower. This inspired us to systematically sweep the parameter values to find those that more faithfully reproduce the measured profiles. We find in the right most plot that lowering the diffusion constant by half, raising the motor hydrolysis rate by 50% and increasing the maximum tubulin concentration by a factor of 3 results in an ATP depletion profile that better matches our data. Here we see a maximum gradient of 0.5 *µ*M*/µ*m and global ATP depletion in 58 minutes. Note that these alternative parameter values are still within the uncertainty of knowledge of their values.

Note that our goal with these simulations is not to fit data curves, but rather to examine plausible behaviors for realistic parameter choices. With reasonable parameter choices, within a factor of a few of our assumed parameter values, we find gradients in ATP of the right scale appear in our simulation. While we attempted to characterize the most significant processes impacting the rate of ATP consumption, we leave the open questions of what other mechanisms may impact the shape of gradients. These may include fluid flow, motor cooperativity, motor jamming, motor detachment, and motor force-stalling, to name just a few.

More generally, the adequate accord of such a reaction-diffusion model for how gradients develop in ATP with measurements attests to general principles for how molecular gradients can form in space. This mechanism differs in fundamental conceptual ways from the simple and classical mechanism that develops the *Bicoid* gradient in *Drosophila*. There, in *Drosophila*, some reaction (degradation) rate *k* uniform in space conspires with a nonuniform boundary condition in space (specifying a uniform production in time at one end of the embryo), and competes with diffusion at coefficient *D* to set a steady-state gradient with a characteristic length scale 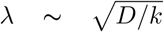. However, applying identical logic to the present motor-microtubule system would be hasty (and would anticipate an incorrectly and negligibly-long analogous gradient length scale).

Instead, we contend that gradients in the present active matter system ensue very differently. Here, gradients in ATP develop when its hydrolysis reaction grows sufficiently nonuniform in space (thanks to the underlying self-organization of motors and microtubules), against uniform boundary conditions in space. Specifically, gradients ensue from sufficiently dramatic localization of hydrolysis rates, whose effects accumulate for long enough times compared to the characteristic diffusive smoothing times that appreciable gradients sustain over pattern formation. Indeed, this connection between a field’s spatial gradient and that accumulated over time in a consumption rate field can be expressed as a precise mathematical statement: the steepest gradient in concentration is at most the maximal gradient in hydrolysis rates accumulated over earlier times (see Supplementary Information, section S13). Such concentration gradients thus testify to reaction rate gradients. In sum, then, the precise competition of characteristic physics which govern whether a gradient appears, and its quantitative extent, differs from other celebrated classical gradients organizing biology. That an emergent patterning of reaction rates itself controls the appearance of sustained gradients may be a principle that operates beyond the specific motor-microtubule setting explored in this work.

### Comparing and appraising underlying physical origins for measured power

Provoked by our measurements and these analyses in the preceding sections, a fundamental and urgent question arises. What does that ATP hydrolysis “pay for?” Of course, mechanistically, we know that ATP is being consumed by motors as they carry out their walk along microtubules. But here we mean it differently and functionally. That energy is dissipated through elementary processes such as ordering and contraction of the microtubule network. How much energy do these processes cost?

Microscopically, a huge variety of dissipative processes are taking place during the microtubule-motor rearrangements attending aster formation. As shown in Figure 5 the power varies considerably at different stages of the aster formation process. To that end, we explore the power associated with a variety of processes that we imagine are taking place concurrently and would cost different amounts at different stages of the aster formation process.

Given the measured power, we were intrigued to compare it to the power associated with a variety of elementary dissipative processes that take place during aster formation as shown in Figure 7. For example, as is evident from the radius as a function of time in Figure 5, the volume of the microtubule aster is decreasing over time. As shown in Figure 7(A) and described in detail in the Supplementary Information section S14C, we can perform a simple estimate of the power associated with this contraction as the pressure-volume work done divided by the elapsed time. We find that the pressure-volume power is five orders of magnitude smaller than the measured power. As shown in Figure 7(B), a second dissipative process is the frictional sliding of the various microtubules during the contraction process. A naïve estimate is obtained by replacing a given microtubule by a corresponding sphere of the same dimensions and to work out the Stokes drag (see Supplementary Information section S14E). As in the case of the pressure-volume power, this results in a power that is five orders of magnitude smaller than the measured power of Figure 5. As discussed in the Supplementary Information section S14E.1, a better estimate can be made in which the microtubule is treated as a rod rather than a sphere and in this case the computed power is even smaller. It is possible that crowding effects could amend these estimates. Another approach to estimating the power is offered by field theories of nematic ordering in which the state of the system is characterized by the spatially varying tensor *Q*_*ij*_(**r**, *t*). This estimate is trickier to make since the parameters in such a field theory of motor-microtubule systems are not well known. Nevertheless, as seen in Figure 7(C) (and described in more detail in the Supplementary Information section S14F), the power we estimate associated with such ordering is many orders of magnitude smaller than the measured power. The final dissipative process highlighted in Figure 7(D) is that of building and maintaining a nonequilibrium gradient of motors radially outward from the center of the aster. One way to think about such a gradient is that if there were not some active transport carrying motors towards the aster center, then diffusion would smooth out that gradient. As we show in Supplementary Information section S14G, there is a well-defined prescription for estimating the power to maintain such a gradient using statistical physics; we find that this estimated expenditure at late times is roughly an order of magnitude lower than our measured powers at late times. This interesting result suggests the hypothesis that a significant fraction of the ATP hydrolysis consumed by the motors is “spent” to build and then maintain this gradient.

**Fig. 7.**
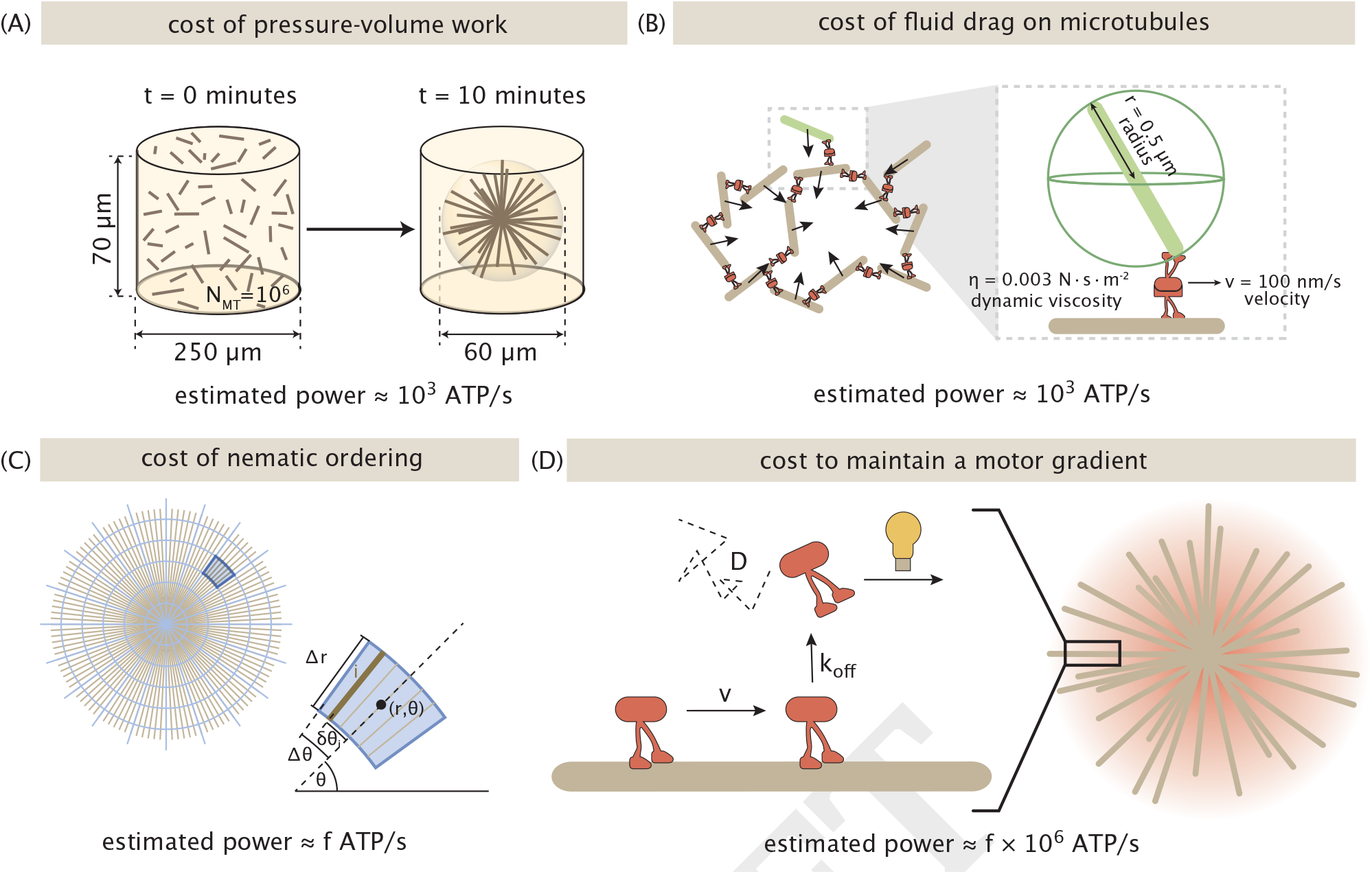
Mechanistic processes taking place during aster formation and their estimated power. Each schematic considers a different dissipative processes that occurs during the formation of an aster. (A) Schematic of the power of compressing an “ideal gas” of microtubules. (B) The power of dragging microtubules through a viscous medium. (C) Power estimate for inducing nematic ordering in a random array of microtubules. (D) The power to maintain a concentration gradient of motors in the aster.

## Discussion

It is practically a cliché to note that living organisms are “out of equilibrium.” And yet, because of separation of time scales, equilibrium ideas are often useful in a variety of nonequilibrium settings. To render our approach to these problems more precise, it is useful to measure the rate at which energy is being consumed to maintain systems in these nonequilibrium states. One useful way to think about such dissipative processes in living organisms is to recast them not as a cost, but rather as a mechanism.

Recent work has made exciting strides in characterizing energy expenditure in a variety of systems via techniques that include calorimetry (5, 6), oxygen consumption (30–33), and fluorescent metabolite measurements (30, 34, 35). Some of these studies capture the total energy expenditures of some biological system of interest and reveal intriguing mismatches between the measured dissipation and contribution such as the mechanical power. Accompanying theoretical efforts have reached similar conclusions (36, 37). Models have also even started to probe how biochemical energy sources might vary in space (38). Yet scarce experimental measurements impede their calibrated application to cells (35, 36). Further, these methods do not always have the spatial resolution that can isolate individual mechanisms, their contributions to the total dissipation, nor related gradients on the cellular or sub-cellular scale.

Having the ability to measure gradients is crucial since gradients are ubiquitous and functionally critical for processes such as body symmetry and division axes (29), division time, and cell size (26). These gradients resist thermodynamic equilibrium, requiring continuous energy input for their formation and maintenance. Motivated by the need to develop a physical understanding of how energy fluxes give rise to biological order, we developed an *in vitro* system of motor-microtubule assemblies to ask how ATP consumption across space and time drives the dynamics of structure formation. Within this framework, ATP gradients emerge as a consequence of motor protein activity. These gradients serve as a measurable signature of the energetic mechanisms associated with the rearrangement of microtubules during aster formation.

In our *in vitro* system, such gradients emerge from complex interactions between motors, microtubules, flows, diffusion, and ATP/ADP competitive binding and are not readily predicted from individual molecular properties. As shown by our finite element modeling, this cooperation of interactions can be quantitatively addressed by rigorous modeling and real calibrated and thermodynamically-meaningful measurements. These systems display gradients of order *≈*few *×*10^*−*1^ *µ*M*/µ*m over tens of microns— hundreds of times steeper than celebrated developmental gradients discussed in Figure 4.

The differences between the magnitudes of the gradients we measured and those observed in specific cellular and organismal contexts (Figure 4), possibly come from a host of additional complexities in those systems, which we need to account for in future studies with living systems. For example, in whole cells, there are metabolic energy sources (including mitochondria) and sinks of energy other that motors (filaments, ATP driven enzymes). It is a fascinating, abiding question for our field, to ask whether or how the conspiracy of both sources and sinks manifests appreciable gradients (or not) of biochemical sources of energy, as started in a beautiful recent work (38). We can begin addressing these complexities by finite element modeling, built upon our initial steps described here. In terms of future studies, our analysis cautions that the dissipation rate estimated by mean-field quantities (such as average, not local, concentrations) can generically overestimate the true macroscopic dissipation rate, as we dissect further in the Supplementary Information section S12 using an application of Jensen’s inequality. This highlights the need for spatially resolved measurements. To understand the nonequilibrium nature of whole cells and their assemblies into tissues and organisms, we acknowledge that an approach with at least a similar level of rigor and scope as the one we have demonstrated needs to be undertaken in these further contexts.

Our experiments, calibrated in real molecular units, resolve substantial dissipation rates, up to a few *×*10^8^ ATPs per second (*>* 10^*−*17^ W*/µ*m^3^), which for comparison, exceeds the average optical power densities of our activation light of 10^*−*17^ W*/µ*m^3^. As illustrated in Figure 8, these power values measured in contracting asters enjoy informative comparisons with typical metabolic power expenditures of bacteria (*≈*10^3^ ATP/s/cell in slow-metabolizing anaerobic *P. aeruginosa* (39); *≈*10^6^ ATP/s/cell (40, 41) in basal metabolism to *≈*10^7^ ATP/s/cell in exponential growth (42, 43) in organisms like *E. coli*); mouse oocytes (approximately 10^5^ ATP/*µ*m^3^/s totaling 2 *×* 10^10^ ATP/s/oocyte as inferred from NADH turnover) (30); *Xenopus* embryos (on average, approximately *≈* 1.5 *×* 10^4^ ATP/*µ*m^3^/s in the two-cell-stage to *≈* 2.1 *×* 10^4^ ATP/*µ*m^3^/s at the cleavage stage as inferred by calorimetry, with cell cycle oscillations about one-hundred-fold smaller (6)); or other, extensile, active matter microtubule systems (few *×*10^3^ ATP/s/*µ*m^3^) (5); see Supplementary Information section S11 for further numerical discussion. Our experiments further report ATP gradients develop in space (with steepnesses of order at least a few tenths of micromolar per micron, sustained over tens of microns).

**Fig. 8.**
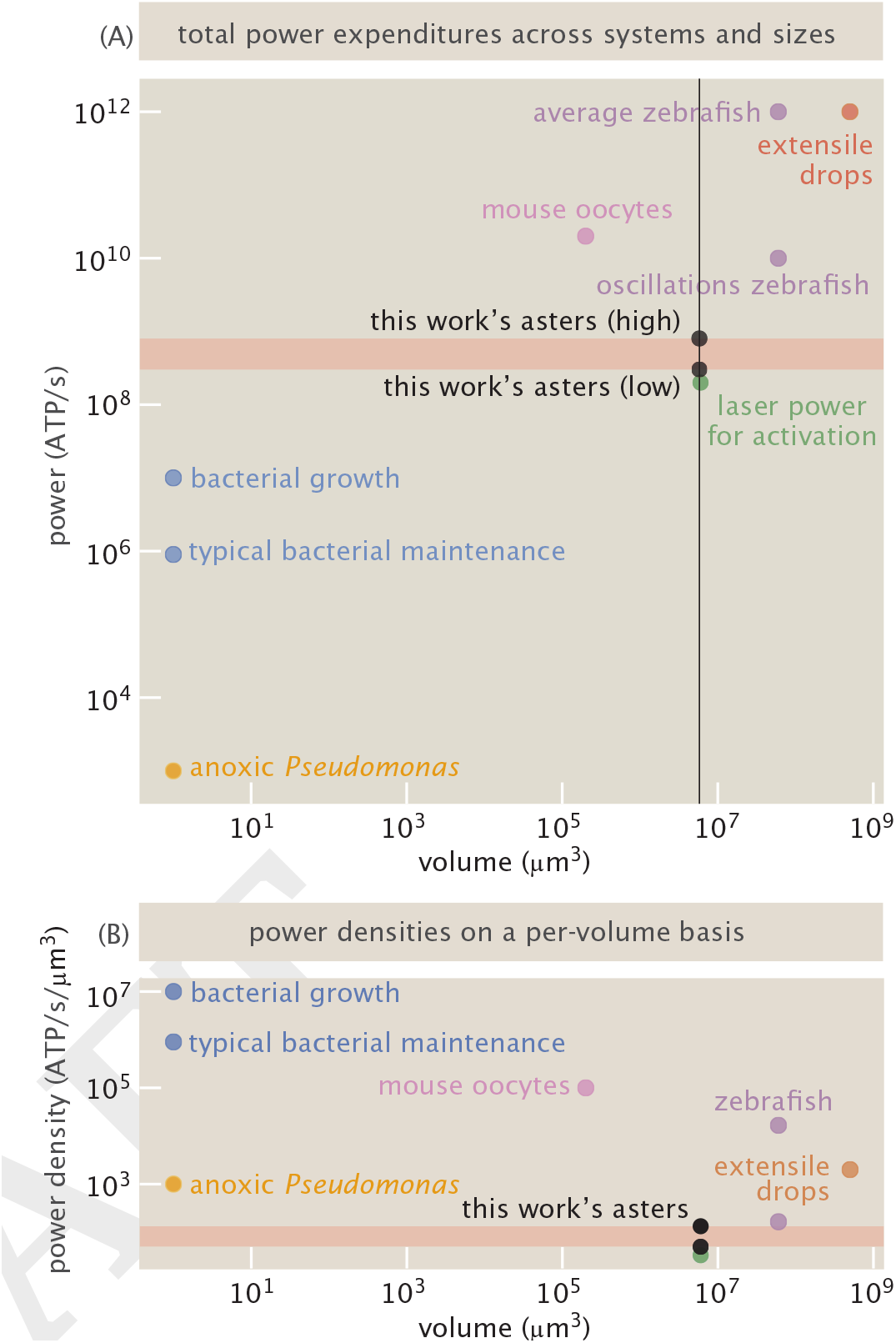
Comparisons of this work’s measured thermodynamic dissipations with those of relevant organismal and active matter contexts. (A) Absolute power versus total volume for embryos, microbial cells, and distinct experimental contexts. (B) The power expenditures of these contexts measured on a per-volume basis. See SI *§*11 for further details.

Beyond the aim of developing an energetic consumption census, the formation of ATP gradients motivates questions that ask: might these energetic gradients hold significance to the cell? As we have highlighted in the results, the energetic cost of maintaining a gradient is comparable to energy expenditure we observe. Widely celebrated cellular gradients, including the morphogen gradient in Drosophila (29), which assigns the embryo’s anterior-posterior body axis, the protein kinase gradient in fission yeast (26), which regulates the timing of cellular division, ras-related nuclear protein gradients (44), which control spindle assembly, and others, are so beloved due to how they drive cellular function. Perhaps, ATP gradients have similar implications, localizing energy availability to incite cellular activity in specified areas. Only with measurements and mathematical models yielding space-time understandings of energetic dissipation can one truly know how cells exists away from equilibrium.

## Materials and Methods

Experimental, computational, and theoretical methods and results are described extensively in the supplementary information.

## Data, Materials, and Software Availability

Data and analysis code used to generate this study’s analyses are available open source; see https://github.com/RPGroup-PBoC/am_atp.

## ACKNOWLEDGMENTS

We gratefully thank Frank Jü licher, Justin Bois, Xingbo Yang, Erwin Frey, Sara Mahdavi, Avi Flamholz, Peter Foster, Catherine Ji, Victor Gomez, Vahe Galstyan, Mohammadamin Tajik, and members of the Phillips and Thomson groups for insightful discussions. We would also like to thank the David Van Valen and Rebecca Voorhees labs for providing resources for performing protein expression and purification. We thank the NIH for support through award numbers DP1OD000217 (Director’s Pioneer Award) and NIH MIRA 1R35 GM118043-01 to RP. MT thanks The Moore-Simons Project on the Origin of the Eukaryotic Cell. AID and GLS were supported by the NSF Graduate Research Fellowship DGE-1745301. GLS thanks the NSF-Simons National Institute for Theory and Mathematics in Biology (NITMB) Fellowship supported via grants from the NSF (DMS-2235451) and Simons Foundation (MP-TMPS-00005320).

## References

1. T Gregor, W Bialek, RR de Ruyter van Steveninck, DW Tank, EF Wieschaus, Diffusion and scaling during early embryonic pattern formation. Proc. Natl. Acad. Sci. 102, 18403–18407 (2005).

2. T Gregor, AP McGregor, EF Wieschaus, Shape and function of the bicoid morphogen gradient in dipteran species with different sized embryos. Dev. Biol. 316, 350–358 (2008).

3. KW Rogers, AF Schier, Morphogen gradients: From generation to interpretation. Annu. Rev. Cell Dev. Biol. 27, 377–407 (2011).

4. JA Drocco, O Grimm, DW Tank, E Wieschaus, Measurement and perturbation of morphogen lifetime: Effects on gradient shape. Biophys. J. 101, 1807–1815 (2011).

5. PJ Foster, et al., Dissipation and energy propagation across scales in an active cytoskeletal material. Proc. Natl. Acad. Sci. 120, e2207662120 (2023).

6. J Rodenfels, KM Neugebauer, J Howard, Heat oscillations driven by the embryonic cell cycle reveal the energetic costs of signaling. Dev. Cell 48, 646–658.e6 (2019).

7. Q Yu, D Zhang, Y Tu, Inverse power law scaling of energy dissipation rate in nonequilibrium reaction networks. Phys. Rev. Lett. 126, 080601 (2021).

8. Y Cao, H Wang, Q Ouyang, Y Tu, The free-energy cost of accurate biochemical oscillations. Nat. Phys. 11, 772–778 (2015).

9. Y Deng, D Beahm, I S R Srpeshkar, Measuring and modeling energy and power consumption in living microbial cells with a synthetic atp reporter. BMC biology 19, 101–121 (2021).

10. J Li, JM Horowitz, TR Gingrich, N Fakhri, Quantifying dissipation using fluctuating currents. Nat. Commun. 10, 1666 (2019).

11. F Nédélec, T Surrey, AC Maggs, S Leibler, Self-organization of microtubules and motors. Nature 389, 305–308 (1997).

12. T Surrey, F Nédélec, S Leibler, E Karsenti, Physical properties determining self-organization of motors and microtubules. Science 292, 1167–1171 (2001).

13. T Sanchez, DTN Chen, SJ DeCamp, M Heymann, Z Dogic, Spontaneous motion in hierarchically assembled active matter. Nature 491, 431 (2012).

14. M Dogterom, T Surrey, Microtubule organization in vitro. Curr. Opin. Cell Biol. 25, 23–29 (2013).

15. PJ Foster, S Fürthauer, MJ Shelley, DJ Needleman, Active contraction of microtubule networks. eLife 4, e10837 (2015).

16. D Needleman, Z Dogic, Active matter at the interface between materials science and cell biology. Nat. Rev. Mater. 2, 17048 (2017).

17. ME Janson, et al., Crosslinkers and Motors Organize Dynamic Microtubules to Form Stable Bipolar Arrays in Fission Yeast. Cell 128, 357–368 (2007).

18. TD Ross, et al., Controlling organization and forces in active matter through optically defined boundaries. Nature 572, 224–229 (2019).

19. H Yaginuma, Y Okada, Live cell imaging of metabolic heterogeneity by quantitative fluorescent ATP indicator protein, QUEEN-37c. bioRxiv (2021).

20. T Nagai, A Sawano, ES Park, A Miyawaki, Circularly permuted green fluorescent proteins engineered to sense Ca2+. Proc. Natl. Acad. Sci. 98, 3197–3202 (2001).

21. H Yaginuma, et al., Diversity in ATP concentrations in a single bacterial cell population revealed by quantitative single-cell imaging. Sci. Reports 4, 6522 (2014).

22. RY Tsien, The green fluorescent protein. Annu. Rev. Biochem. 67, 509–544 (1998).

23. KA Foster, JJ Correia, SP Gilbert, Equilibrium binding studies of non-claret disjunctional protein (Ncd) reveal cooperative interactions between the motor domains. J. Biol. Chem. 273, 35307–35318 (1998).

24. KA Foster, SP Gilbert, Kinetic studies of dimeric Ncd: evidence that Ncd is not processive. Biochemistry 39, 1784–1791 (2000).

25. V Sourjik, HC Berg, Receptor sensitivity in bacterial chemotaxis. Proc. Natl. Acad. Sci. 99, 123–127 (2001).

26. JB Moseley, A Mayeux, A Paoletti, P Nurse, A spatial gradient coordinates cell size and mitotic entry in fission yeast. Nature 459, 857–860 (2009).

27. W Driever, C Nüsslein-Volhard, A Gradient of Bicoid Protein in *Drosophila* Embryos. Cell 54, 83–93 (1988).

28. W Driever, C Nüsslein-Volhard, The Bicoid Protein Determines Position in the *Drosophila* Embryo in a Concentration-Dependent Manner. Cell 54, 95–104 (1988).

29. T Gregor, DW Tank, EF Wieschaus, W Bialek, Probing the limits to positional information. Cell 130, 153–164 (2007).

30. X Yang, G Ha, DJ Needleman, A coarse-grained NADH redox model enables inference of subcellular metabolic fluxes from fluorescence lifetime imaging. Elife 10, e73808 (2021).

31. SA Mookerjee, AA Gerencser, DG Nicholls, MD Brand, Quantifying intracellular rates of glycolytic and oxidative ATP production and consumption using extracellular flux measurements. J. Biol. Chem. 292, 7189–7207 (2017).

32. ME Handel, MD Brand, SA Mookerjee, The whys and hows of calculating total cellular ATP production rate. Trends Endocrinol. & Metab. 30, 412–416 (2019).

33. CA Schmidt, KH Fisher-Wellman, PD Neufer, From OCR and ECAR to energy: Perspectives on the design and interpretation of bioenergetics studies. J. Biol. Chem. 297, 101140 (2021).

34. R Sakamoto, MP Murrell, F-actin architecture determines the conversion of chemical energy into mechanical work. Nat. Commun. 15, 3444 (2024).

35. NK Bennett, et al., Defining the ATPome reveals cross-optimization of metabolic pathways. Nat. Commun. 11, 4319 (2020).

36. E Arunachalam, W Ireland, X Yang, D Needleman, Dissecting flux balances to measure energetic costs in cell biology: techniques and challenges. Annu. Rev. Condens. Matter Phys. 14, 211–235 (2023).

37. M Murrell, When a bath is not enough: State-dependent energy injection and nonmonotonic dissipation in cytoskeletal matter. PRX Life 4, 027001 (2026).

38. R Kumar, IG Johnston, Estimating physical conditions supporting gradients of ATP concentration in the eukaryotic cell. Biophys. J. (2025).

39. JA Ciemniecki, CL Ho, RD Horak, A Okamoto, DK Newman, Mechanistic study of a low-power bacterial maintenance state using high-throughput electrochemistry. Cell 187, 6882–6895 (2024).

40. SJ Schink, E Biselli, C Ammar, U Gerland, Death rate of e. coli during starvation is set by maintenance cost and biomass recycling. Cell Syst. 9, 64–73 (2019).

41. M Lynch, GK Marinov, The bioenergetic costs of a gene. Proc. Natl. Acad. Sci. United States Am. 112, 15690–15695 (2015).

42. AH Stouthamer, Theoretical study on amount of ATP required for synthesis of microbial cell material. Antonie Van Leeuwenhoek J. Microbiol. 39, 545–565 (1973).

43. M Mori, C Cheng, BR Taylor, H Okano, T Hwa, Functional decomposition of metabolism allows a system-level quantification of fluxes and protein allocation towards specific metabolic functions. Nat. Commun. 14, 4161 (2023).

44. D Oh, CH Yu, DJ Needleman, Spatial organization of the Ran pathway by microtubules in mitosis. Proc. Natl. Acad. Sci. 113, 8729–8734 (2016).

