## Supplementary Information for "Energetic gradients emerge in developing motor-microtubule structures"

##### This PDF file includes:

Supporting text
Figs. S1 to S87
Tables S1 to S5
SI References

**12 Jensen's inequality shows that hydrolysis rates can be underestimated by spatially-averaged values** **118**

**13 Spatial gradients in ATP are upper bounded by those in underlying reaction rates (accumulated over time)** **120**

**14 Power Estimates** **122**

### Supporting Information Text

#### 1. Experimental preparation of contractile active matter system

**A. Flow cell preparation.** To prevent non-specific binding of proteins to glass surfaces, microscope slides and coverslips were cleaned and coated with a polyacrylamide solution based off of a protocol written by the Dogic Lab (1) with slight modifications described in detail in previous work from our labs (2). In brief, the protocol involves cleaning via three sonication steps, in the detergent Hellmanex, ethanol and then KOH, with rinses of double-distilled water and/or ethanol in between each step. There is then an incubation step in HCl to strip away any trace metals. The glass is then coated in silane, which acts as a bridge between polyacrylamide and glass, washed and finally coated and stored in a polyacrylamide solution.

Using these treated slides and coverslips, we construct the flow cells. We rinse off the glass with double-distilled water and dry with $N_2$ gas, ensuring there is no contact with the side of the glass that will face the inside of the chamber. We place channels cut out of parafilm flat on the slide. Rectangular channels are cut with dimensions 3 mm by 18 mm using a Silhouette Studio machine. We then place a coverslip on top of the parafilm, ensuring the parafilm channel extends past both sides of the coverslip. We place the slide on a hot plate at 65 degrees Celcius and gently press down on the coverslip to melt the parafilm and allow it to anneal to the slide and coverslip. The final depth of the chamber ranges from $60 - 100\,\mu$m depending on the amount of pressure used when pressing the coverslip. This can be quantified by using brightfield microscopy to measure the distance between the top and bottom focal planes of the parafilm.

Chambers are loaded by pipetting liquid on one side, placing the pipette up against the space between the coverslip and slide. Capillary action helps the fluid fill the chamber. Each side is sealed with Picodent Twinsil Speed, a fast-setting, silicone polymer.

**B. Sample Preparation and Reaction Mixture.** The reagents for our imaging assays are comprised of a common buffer with varying amounts of ATP, motor proteins, and microtubules, which we specify for each experiment type in the following paragraphs. The common buffer is comprised of 66.7 mM PIPES at pH 6.8, 4.7 mM $MgCl_2$, 0.83 mM EGTA, a crowding agent (20-22% glycerol, Sigma, G5516), a surface passivating agent (0.50 mg/mL Pluronic F-127, Sigma, P2443), and an oxygen-scavenging system to prevent photobleaching (0.37 mg/mL pyranose oxidase, Sigma, P4234; 7.2 mg/mL glucose, Thermo Fisher Scientific, USA; 9 $\mu$g/mL catalase, Sigma, C40; 5.4 mM DTT, Thermo Fisher Scientific, USA; 2.0 mM Trolox, Sigma, 238813).

For aster experiments, the final reaction mixture consisted of 500 $\mu$M of MgATP (Sigma A9187), 2.8 $\mu$M Queen A81D (3), 0.6 $\mu$M Ncd-mCherry-micro, 0.6 $\mu$M Ncd-mCherry-iLID, and GMPCPP-stabilized microtubules (1.3 $\mu$M tubulin).

For aster ATP calibration experiments, we varied the concentration of MgATP (ranging from 50 $\mu$M to 3000 $\mu$M) and included 2.8 $\mu$M Queen A81D (3), and GMPCPP-stabilized microtubules (1.5 $\mu$M tubulin). Motor proteins were excluded to ensure no hydrolysis or ATP interferes with the calibration measurement.

Sample preparation and handling were performed in a dark room with red light to minimize early light activation of the optogenetic proteins. We prepared the reaction mixture right before loading it into the flow cell and sealed it using Picodent Twinsil Speed. Experiments occured at room temperature (approximately 25°C).

**C. Activation and Imaging Protocol.** In experiments focusing on aster formation, we selected one position within the flow channel, which is illuminated by an excitation region with a diameter of 400 $\mu$m. In the case of experiments related to ATP hydrolysis, the entire field of view is illuminated. Typically, one experiment is conducted per flow channel.

The fluorescent motors (mCherry labeled) and the Queen A81D ATP probe (3) were imaged simultaneously every 20 seconds using a $\times 10$ objective. The exposure time values for 405nm and 480nm excitations were 33-150 ms and 50-160 ms, respectively, and 100-300 ms for 587 nm mCherry excitation.

**D. ATP Calibration.** In order to convert fluorescence counts emitted by the ATP probe to ATP concentration units, we perform a calibration procedure where we image the ATP probe in buffers with titrated amounts of known ATP standards. We prepare the common buffer as described above in section B. We include the ATP probe at 2.8 $\mu$M with GMPCPP-stabilized microtubules (1.5 $\mu$M tubulin) and no motor proteins. Known ATP standards are prepared by using powder magnesium salt ATP to make solutions of varying concentrations (0 $\mu$M to 3000 $\mu$M ATP), which we add to our reaction mix.

As illustrated in Figure 2(B) of the Main Text, the ATP probe is a ratiometric based sensor that emits photons for excitations at 405 and 480 nm. The binding state of ATP to the probe causes the ratio of photons emitted from 405 nm vs 480 nm excitation to change. Thus, we perform epifluorescence imaging with excitation filters (405 and 480 nm) and a $\times 10$ objective. The exposure time values for 405nm and 480nm excitation are 150ms and 160ms, respectively. Typically, five positions within the same flow channel are imaged as technical replicates. The experiments are performed at room temperature (approximately 25°C).

After using our image analysis procedure below to correct for background signal and uneven illumination (see sections A and B) on images collected from both 405 nm and 480 nm emissions, we find the ratio of fluorescent emission counts at each pixel location of the image by dividing each image element-wise,

$$\text{ratio}_{ij} = \frac{\text{image}(405\text{ nm})_{ij}}{\text{image}(480\text{ nm})_{ij}} \quad [1]$$

We then compute the mean value and standard deviation of pixel ratios in a given image. We plot the mean ratio value against ATP concentration in Figure 2(C) of the Main Text, where the dots are the mean values and the error bars are the standard deviation. To establish a calibration, we fit the data to a Hill function following the equation,

$$R = (R_{\max} - R_{\min})\frac{\frac{[\text{ATP}]}{K_{\text{M}}}}{1 + \frac{[\text{ATP}]}{K_{\text{M}}}} + R_{\min}, \quad [2]$$

where $R$ is the intensity ratio of the bound and unbound channels, $R_{\max}$ is the ratio value as the ATP concentration goes to infinity, $R_{\min}$ is the ATP value at zero ATP concentration, [ATP] is the concentration of ATP, and $K_{\text{M}}$ is the Michaelis-Menten constant.

***D.1. The ATP probe is selective to ATP rather than ADP.*** When aiming to accurately report ATP concentrations with a fluorescent based ATP reporter, it is imperative that the reporter, QUEEN, does not falsely fluoresce upon binding ADP rather than ATP. The designers of the probe test for this sensitivity in two mutants of the QUEEN protein by titrating the concentrations of ATP and ADP and measuring the resultant fluorescent readouts. They find that upon adding increasing amounts of ADP with the probe, there is no change in the fluorescent readout, meaning there is not false fluorescence due to ADP, see the blue curves

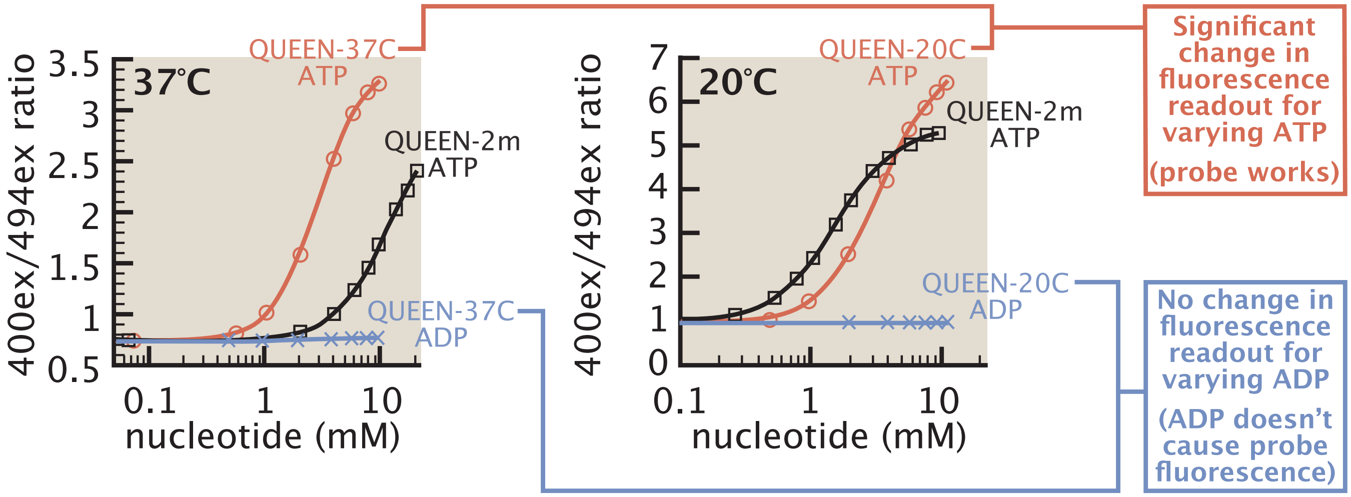

**Fig. S1. The QUEEN ATP probes are selective for ATP and against ADP.** This figure is adapted from supplementary figure 1 (B) and (D) in (3). These plots show the fluorescent probe response to titrating concentrations of ATP and ADP for QUEEN-37C, on the left, and QUEEN-20C, on the right. Both plots also show the response of QUEEN-2m to ATP at each temperature. Comparing the red and blue curves, the probe only exhibits a dynamic response to ATP with no response to varying concentrations of ADP.

in Figure S1. Meanwhile, when they add increasing concentrations of ATP, they do find a change in the fluorescent readout, indicative that the probe is indeed functional, see the red curves in Figure S1. However, we note the authors did not directly test the sensitivity of the QUEEN mutant we use in our study, QUEEN-7µ A81D. Figure S2 outlines a family tree of the QUEEN proteins mentioned here to give a sense of the how similar the protein sequences are. The nearest common ancestor to all the mutant proteins discussed here is QUEEN-7µ. This probe sandwiches a circularly permuted enhanced green florescent protein (cpEGFP) in between the epsilon subunit of the *Bacillus* PS3 ATP synthase. The epsilon subunit is the region of the ATP synthase that binds ATP. The probe we use has a single amino acid mutation at the 81st position from an alanine (A) to an aspartic acid (D). This single change lowers the affinity of the probe to ATP having a $K_{\mathrm{d}} = 0.07$ mM (4) as opposed to a $K_{\mathrm{d}} = 0.006$ mM, making this mutant more suitable to the ATP concentrations we aim to measure in our study. Both of these probes are designed to function at 25°C. The authors performed further mutations to modulate the dynamic ranges of probes at varying temperatures. As intermediates, they made QUEEN-2m, which is designed for use at 20°C and QUEEN 84BP, which is designed for use at 37°C. To make these probes, they combined parts of the epsilon subunit from *Bacillus* PS3 and *Bacillus subtilis* in different proportions. Then, to fine tune their desired dynamic range, they made two amino acid mutations yielding QUEEN-37C and QUEEN- 20C each with a $K_{\mathrm{d}} \approx 3$ mM. These versions of QUEEN are the two they demonstrate to have no ADP sensitivity. In this study, we have taken as an assumption that given the robust changes in the sequences giving rise to both QUEEN-37C and QUEEN-20C, that the epsilon subunits of ATP synthases are highly selective to only ATP.

**E. Flow Chamber, Sample Preparation, and Imaging Protocol for Microtubule Gliding Assays.** The flow chambers were constructed as previously described (5). In brief, we built a chamber by assembling an amino-silanized coverslip and an acrylamide-coated microscope slide separated by parafilm spacers.

We treated the amino-silanized glass surface with glutaraldehyde to attach antibodies. For this purpose, the chamber was incubated with 10% (v/v) glutaraldehyde (Sigma, G7776) for 30 minutes. After removing unreacted glutaraldehyde by rinsing the chamber with MilliQ water, it was then incubated with a 0.02 mg/ml solution of anti-FLAG antibody (F3165, Sigma) to specifically bind motor proteins to the glass surface. The remaining exposed surface was blocked with a 0.2% (w/v) Pluronic F-127 (Sigma, P2443) and 2 mg/ml $\beta$-casein (Sigma, C6905) solution for 5 minutes. Motors were bound to the surface by incubating Ncd-YFP-iLID FLAG-tag motor proteins (at $\approx 1$ nM in 10 mg/ml bovine serum albumin (BSA, JT Bakers), 1 mM DTT, and 500 $\mu$M mgATP in M2B buffer (80 mM PIPES, 1 mM EGTA, 2 mM $MgCl_2$)) for 5 minutes. Unbound motors were washed out with M2B, and then AlexaFluor 647 labeled GMPCPP-stabilized microtubules in M2B with 1mM DTT were flowed in. After 5 minutes of incubation at room temperature, the flow cell was rinsed with M2B to remove unbound microtubules.

The imaging buffer consisted of M2B buffer with 0.5 mg/ml $\beta$-casein, 0.50 mg/mL Pluronic F-127, 3 mM $MgCl_2$, 2 mM Trolox, and an oxygen scavenging system (0.37 mg/ml pyranose oxidase, 7.2 mg/ml glucose, 9 $\mu$g/ml catalase, 5.4 mM DTT, 2.0 mM Trolox). In microtubule gliding experiments, we varied the concentrations of mgATP (ranging from 20 $\mu$M to 5000 $\mu$M), ADP (ranging from 0 $\mu$M to 5000 $\mu$M), and potassium phosphate (pH 7.0, ranging from 0 mM to 40 mM). Following the addition of the reaction mixture, the flow cell was sealed using Picodent Speed. The experiments were repeated at least three times at room temperature (approximately 25°C).

Image acquisition of AlexaFluor 647-labeled GMPCPP-stabilized microtubules was performed in a sealed chamber using Total Internal Reflection Fluorescence (TIRF) microscopy with an HCX PL Apo 100×/1.47 TIRF objective. Imaging was performed at one frame per second for 100 seconds. We also imaged Ncd motor proteins (YFP labeled) bound to the glass surface. Individual microtubules were tracked using custom-written Python code to determine their speed.

**F. Protein Purification.** NCD kinesin motors proteins were prepared as described in previous works from our labs (2). Briefly, plasmids were transfected into Sf9 cells and incubated at 27°C while shaking at 120 rpm for 3 days. Proteins are selected for via a FLAG affinity tag and stored at −20°C in 50% glycerol.

Microtubules purchased from PurSolutions LLC were reconstituted without the inclusion of any dye. They were stabilized with GMPCPP. The exact details of this preparation can be found in a previous work from our labs (2).

We transfected BL21(DE3) Competent Cells cells with a pBiEx-1 vector containing pRSET-QUEEN 7mu A81D (Addgene # 129351, gift from Yasushi Okada) (3). Cells were left to grow in a shaker rotating at 250 rpm and incubated at a temperature 37°C. To facilitate growth, we periodically added 20% glucose to the culture. Once the culture reached an optical density of around 0.5, we induced with IPTG and continued to shake overnight, reducing the temperature to 25°C. We then centrifuged cells at 4000 rpm for 30 minutes at 4°C and removed the supernatant. Cells were resuspended in lysis buffer (1 mg/ml lysozyme, 1 cOmplete™ EDTA free protease inhibitor tablet/50 mL, 50 mM imidazole, 0.05 mM MgATP, 5 mM BME, 50 mM sodium phosphate, 4 mM MgCl2, 250 mM NaCl) and incubated for an hour at 4°C while stirring. The lysate is then homogenized with a cell disrupter and then clarified by centrifuging at 145,000 rpm for an hour. The clarified lysate is combined with nickle-NTA beads and stirred for an hour to allow protein binding. Beads and proteins are isolated via three washes (50 mM imidazole, 0.1 mM MgATP, 5 mM BME, 50 mM sodium phosphate, 4 mM MgCl2, 250 mM NaCl) in a filtration column. We add elution buffer (500 mM imidazole, 0.01 mM MgATP, 5 mM BME, 50 mM sodium phosphate, 4 mM MgCl2, 250 mM NaCl) and collect the pass through into a 50 kDa protein concentrator. The eluted proteins are centrifuged at 4000 G until reaching the desired volume. His-tags are severed by adding a TEV protease at a 1/25 ratio to proteins. Proteins are dialyzed overnight in 1 L of dialysis buffer (same as the wash buffer - 50 mM imidazole, 0.1 mM MgATP, 5 mM BME, 50 mM sodium phosphate, 4 mM MgCl2, 250 mM NaCl). Then, the proteins are washed in 1X M2B buffer in a 50 kDa conical and stored at −20°C in 50% glycerol and 1 mM DTT.

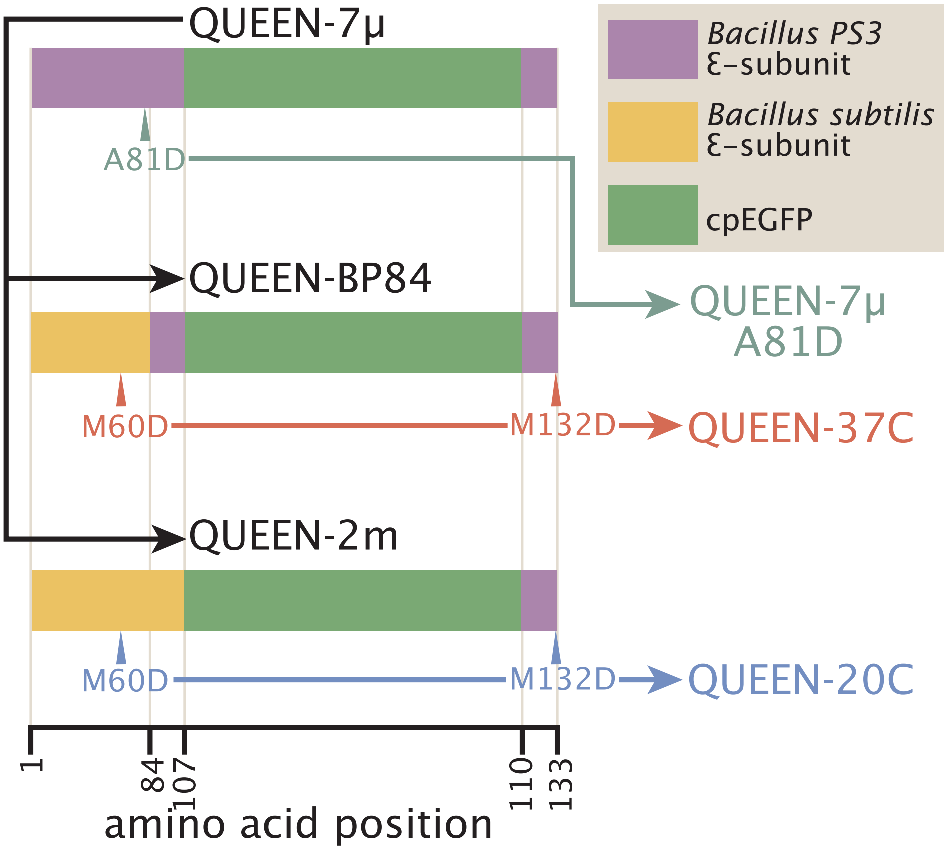

**Fig. S2. Family tree of QUEEN ATP reporters.** This schematic, based on supplementary figure 1(A) of (3), shows the mutations differentiating the versions of probes created by the Yasushi Okada laboratory. Single amino acid mutations are shown with a triangle with a label where the first letter is the original amino acid, the number is the position, and the second letter is the new amino acid. Three base sequences are created starting with QUEEN-7μ which uses the epsilon subunit of *Bacillus* PS3, represented by the purple block. QUEEN BP84 places part of *Bacillus subtilis'* epsilon subunit at the beginning of the protein sequence, represented by the yellow block. QUEEN-2m replaces all of the sequence before the cpEGFP with *Bacillus subtilis'* epsilon subunit.

#### 2. Data analysis protocol

Here we detail our image processing method. We aim to be very careful in checking how our operations modify the data. The predominant goal of the following text is to identify where variation sources occur in our data and how to remove such variation.

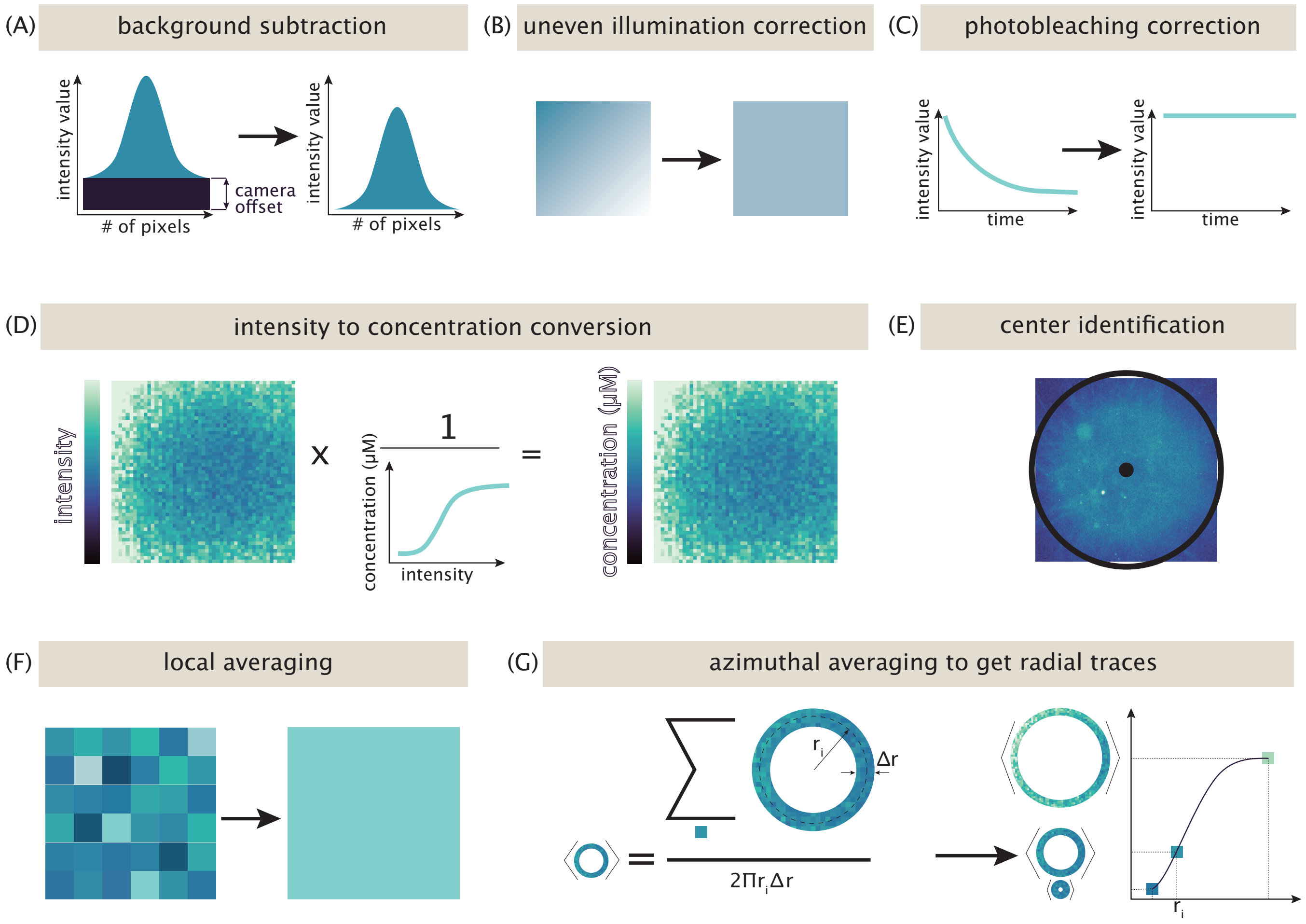

**Fig. S3. Data processing pipeline.** A road map of the analysis performed to turn images into graphs. Each step in the pipeline signifies an action performed on the images collected during an aster formation experiment in which we wanted to quantify the amount of ATP as a function of time.

**A. Background Subtraction.** Cameras artificially add an offset to the intensity values of an image to ensure that no pixels are recorded as having less than zero signal (6). We identify this offset by taking a "dark image", where we close the camera shutter, preventing light from reaching the camera. We use the averaged image of multiple dark images, as shown in Figure S4, as the background image subtracted from all fluorescent images.

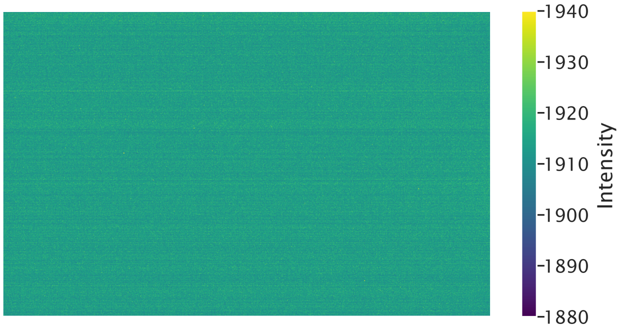

**Fig. S4. Camera offset is found by taking a "dark image".** The average of six images are taken with the camera shutter closed. The average offset value is 1913.8 intensity counts. This image is subtracted from all fluorescent images.

**B. Uneven Illumination Correction.** Microscopy images of uniform samples can often have artificial gradients of intensity across space. Commonly, this artifact arises from vignetting cast by magnification tubes, non-uniform light sources, and off axis light (7). Here, we work to distinguish the fraction of an image's variance that is due to a spatial gradient, implying uneven illumination, rather than local fluctuations.

To search for the origin of variation, we will divide our image into a grid. As we prove in Section B.1, the total variance, $\sigma^2_{\text{tot}}$, of the image will equal the average variance within a grid block, $\left\langle \sigma^2_{\text{in}} \right\rangle$ plus the variance of the average grid block value, $\sigma^2_{\text{btwn}}$,

$$\sigma^2_{\text{tot}} = \left\langle \sigma^2_{\text{in}} \right\rangle + \sigma^2_{\text{btwn}}. \quad [3]$$

If the variance within grid blocks dominates the total variance term, then most of the variation in an image arises from the noise of nearby pixel values, see the second row of Figure S5. However, if the variance between grid blocks (the variance of grid block average values) dominate, the image variance is attributable to intensity differences across regions of the image. In this section, we explore homogeneous ATP images, so we expect all the variance to be within grid blocks. If instead most of the variance is between blocks, this is a sign of uneven illumination, which manifests as a gradient of intensity from one side of the image to the other, see the first row of Figure S5. Thus, we can use this grid block method as a metric for the amount of uneven illumination present in an image.

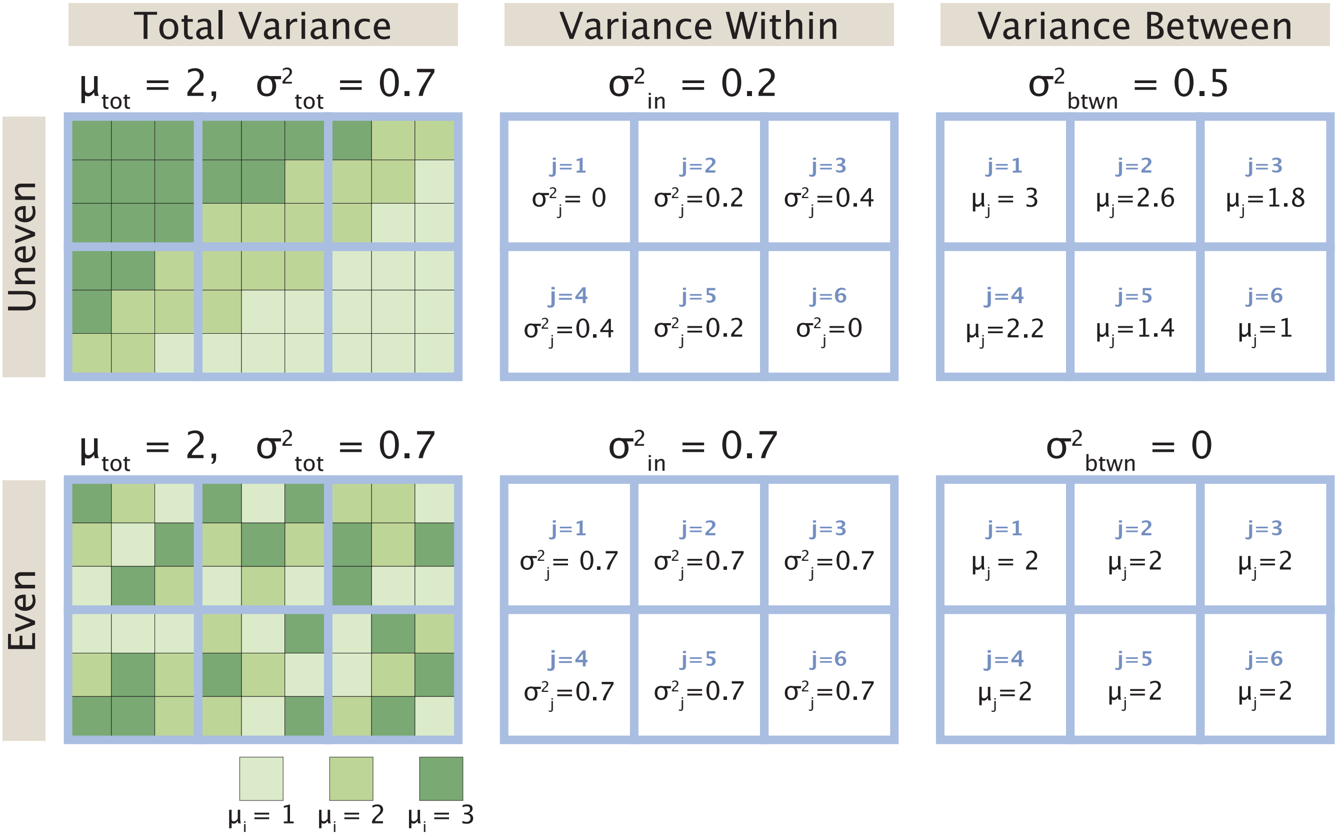

**Fig. S5. Partitioning of variance for unevenly versus evenly illuminated images.** Here, we go through the exercise of finding the variance within and between blocks for two synthetic images. We assign each pixel value in the first column to have a value between one and three. By eye, we would consider the image in the first row to be unevenly illuminated, since there is an intensity gradient across the image. We would consider the image in the second row to be evenly illuminated, since there is no apparent intensity pattern. Splitting the image into six grid blocks of nine pixels each, we quantify our assumptions. The second column (Variance Within) reports the variance of pixels within each grid block. Taking the average of these variances, we report $\sigma^2_{\text{in}}$, our metric for the variance within grid blocks (as defined by Equation 11). In the third column (Variance Between), the mean value of each grid block is reported. Taking the variance of these means gives $\sigma^2_{\text{btwn}}$, our metric for the variance between grid blocks (as defined by Equation 12). While the total image mean and variance for both images are the same, we find that more variance is between blocks in the first row and all the variance is within blocks in the second row. Thus when an image is unevenly illuminated, the variance between blocks dominates.

When dividing our image into into $B$ blocks each containing $N_B$ pixels, we must take note of the variance partitioning in the limits $N_B = 1$ and $N_B = N$, where $N$ is the total number of pixels in the image. In the limiting case that each block only contains one pixel, $N_B = 1$, the variance within a block will be zero and the variance between blocks will equal the total variance of the image. In the opposite limit, where there is one large block containing all pixels, $N_B = N$, the variance within the block is the total variance of the image and there is no variance between blocks. Thus, we must pick an intermediate $N_B$ block size. In Figure S6, we plot the fraction of variance within grid blocks versus the number of pixels per block for all measured ATP conditions. We find that the data fits well to a cubic function of the logarithm of the number of pixels per block. Since we are looking for a middle ground block size that does not favor variance partitioning in either limit, we look near the inflection points of the curves. The light gray boxes contain the inflection points of all curves in both bound and unbound ATP channels. In this region, the slopes of all curves are low, implying that regardless of the choice of box size, the fraction of variance within blocks is about the same. Thus, we arbitrarily select the midpoint of the gray box (on a linear scale), represented by the gray line, as the block size we will use to compare the changes of variance partitioning when correcting images.

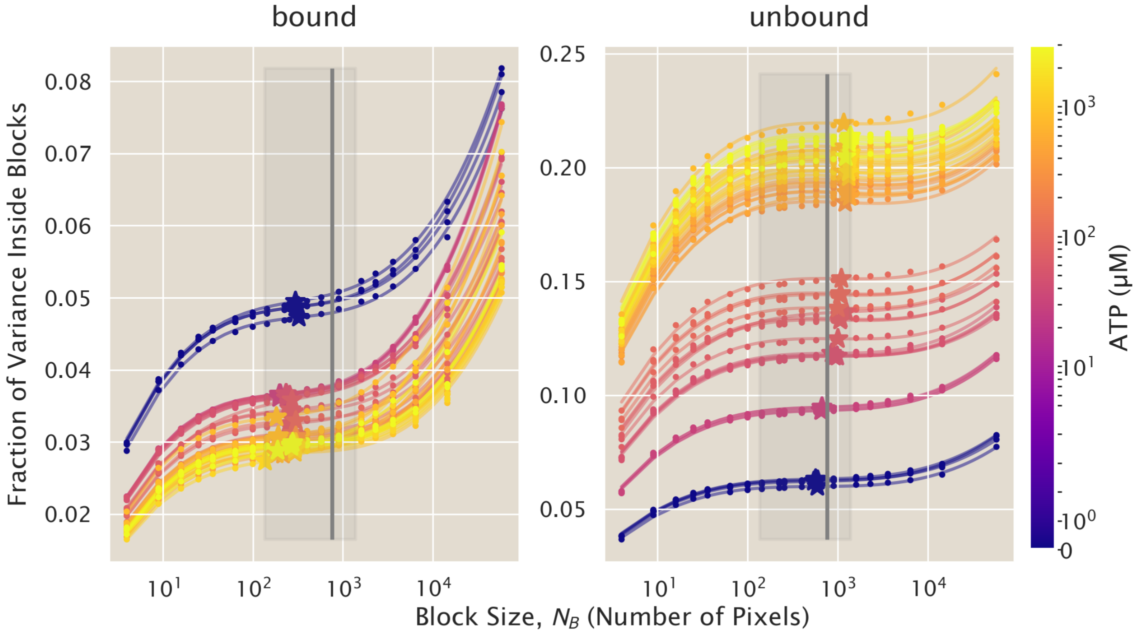

**Fig. S6. The variance within grid blocks varies with the size of the grid block.** We measure how the variation within a block changes with the number of pixels per grid block (represented by circular markers). The trend fits well to a cubic polynomial of the form $\sigma^2_{\text{in}} = ax^3 + bx^2 + cx + d$, where $x = \ln(N_B)$ and $N_B$ is the number of pixels in a grid block. Each solid line fits the data for a single image and is color coded by the concentration of ATP in the sample. We plot the inflection point for each fit (represented by star markers). The gray box is the region containing all the inflection points from both the bound and unbound channel's images. The gray line plots the midpoint (on a linear scale) of the gray box region. The value of the curves at the gray line as compared to the value at line's inflection point is nearly the same. So, we take this value as an arbitrary size by which we can compare the variance partitioning within versus between blocks as we perform uneven illumination corrections on our images.

From Figure S6, we see a very small amount of the variance is located within grid blocks (the y-axis values are low), meaning the vast majority of variance is between grid blocks, implying uneven illumination. We now correct the uneven illumination by fitting our image with a 2D quadratic polynomial of the form,

$$\text{2Dquad}(x, y) = ax^2 + by^2 + cxy + dx + ey + f, \qquad [4]$$

where $x$ and $y$ are spatial coordinates, as described by (8). Next, we create a normalization matrix from our 2D quadratic polynomial fit,

$$\alpha = \frac{\langle \text{2Dquad}(x, y) \rangle}{\text{2Dquad}(x, y)}, \qquad [5]$$

where we divide the average polynomial fit value by the value of each filter pixel value. We then multiply the original image by this filter to find an evened image,

$$\text{im}_{\text{ev}} = \alpha \cdot \text{im}. \qquad [6]$$

By eye, the post-filter image, "Evened Image", in Figure S7, does not appear to retain any of the light gradient present in the "Raw Image". We can confirm numerically that after correction, nearly all the image variance is within blocks. In Figure S8, the fraction of variance within blocks is plotted for all images, with most images containing over 98% of their variance within grid blocks.

Most of the variance before correction occurred along the $y = -x$ diagonal, as seen in Figure S7. Visually, we see the result of the 2D polynomial filter in Figure S9, which plots the intensity value of the image versus the distance along the $y = -x$ diagonal, represented by the multiplication of the x and y pixel coordinate. Indeed, we see a negative slope before correction and an approximately zero slope after correction.

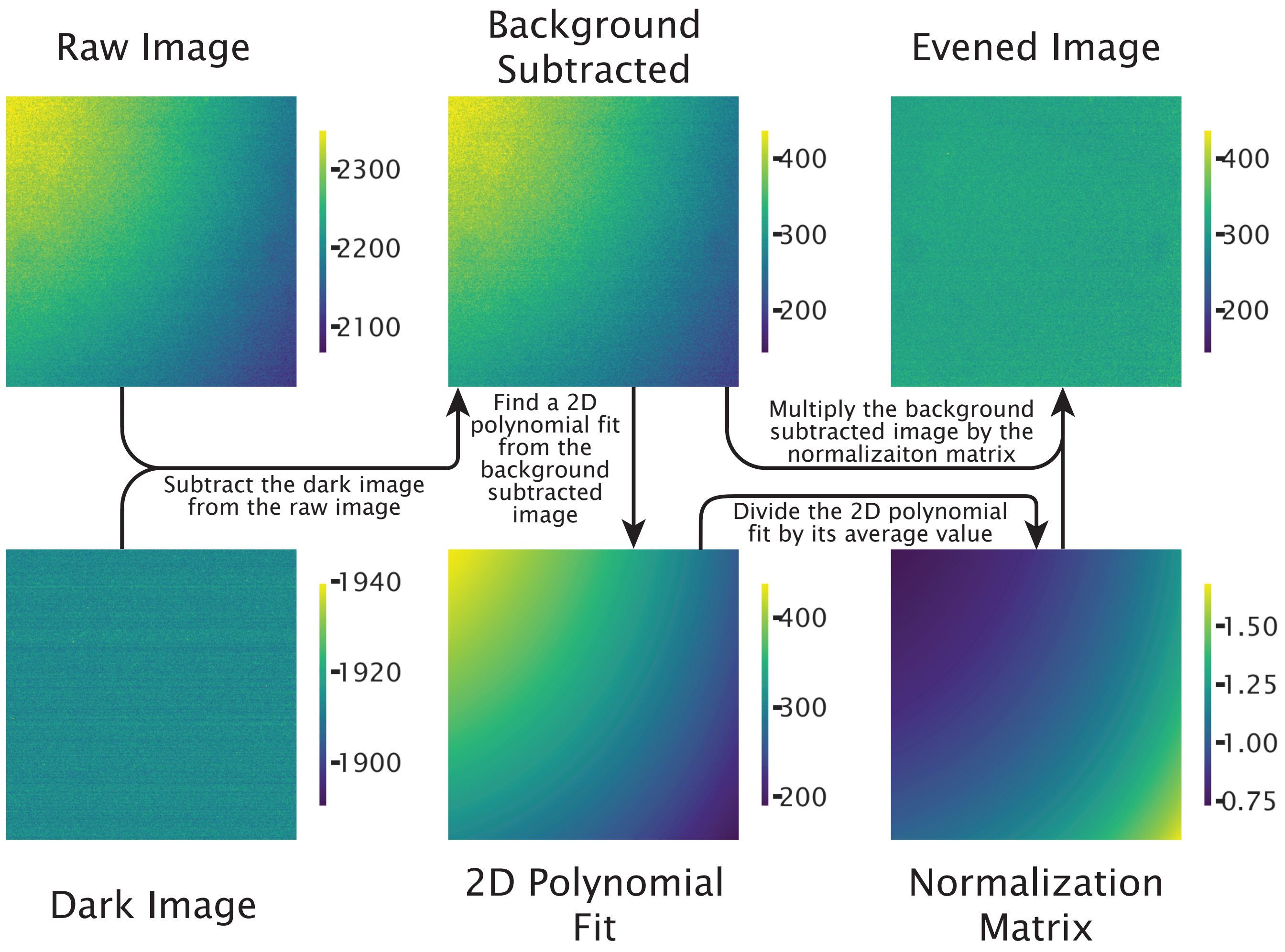

**Fig. S7. Uneven illumination correction process.** We detail the pipeline to correct uneven illumination in images of homogeneous ATP samples. A camera shutter closed, "dark", image is subtracted from the "Raw Image" to create the "Background Subtracted" image. Then, a 2D polynomial is fit to the "Background Subtracted" image using Equation 4. A normalization matrix is created from the 2D polynomial fit using Equation 5. Finally, the "Background Subtracted" image is multiplied by the normalization matrix to create an "Evened Image".

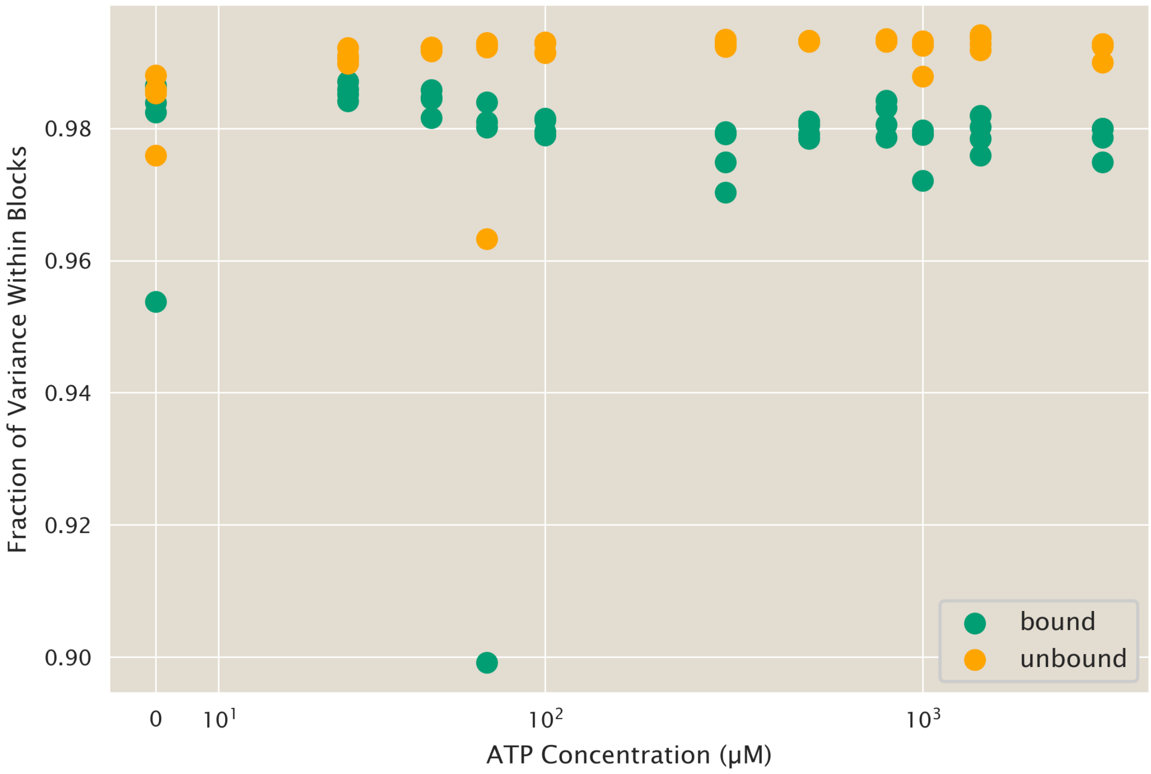

**Fig. S8. Nearly all of the image variance is located within grid blocks after correction.** After applying a 2D polynomial filter to the images, over $85\%$ of variance is located within grid blocks, and over $98\%$ for most images. There is no apparent correlation between variance fraction and ATP concentration. Green dots mark data from images taken in the bound ATP channel, while orange dot mark data from the unbound ATP channel.

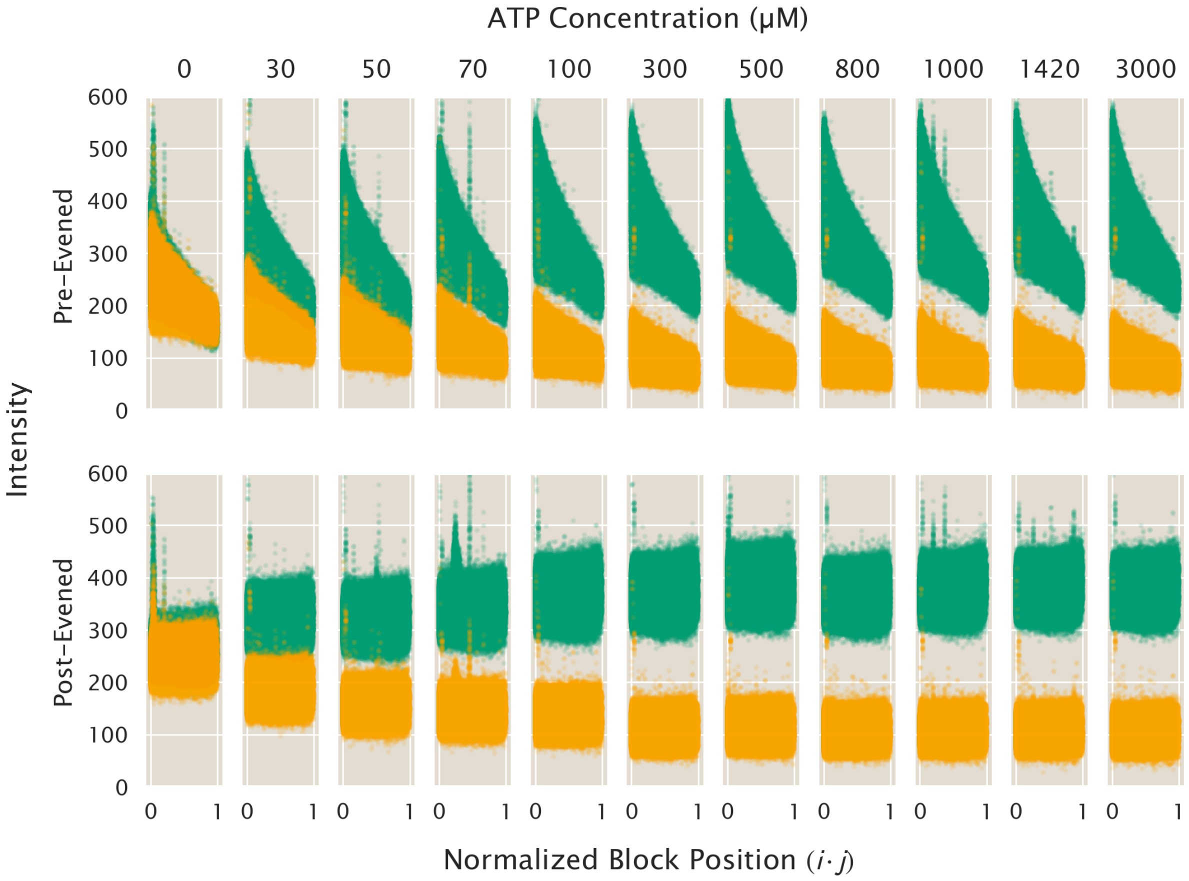

**Fig. S9. Intensity trend with pixel location.** The intensity value of a pixel versus the location of the pixel is plotted along the upper left to lower right diagonal of the image, as assigned by the multiplication of the pixel coordinates, $i \cdot j$. The green dots represent data from the bound ATP channel (excitation 405 nm) and the orange dots present data from the unbound channel (excitation 480 nm). In the top row of plots, images taken before uneven illumination correction, regardless of ATP concentration or imaging channel, show a negative slope. In the second row, images after uneven illumination correction have approximately no slopes, indicating that the filter correctly removed any illumination bias along the diagonal.

Regardless of the gradient direction, for an image of a homogeneous ATP sample to be even, we would expect grid block averages should be the same across the image. This would imply that there is not a correlation between the intensity of the block and it's location. In Figure S10(A) and Figure S10(B), we mark the average value for every grid block of every image. The average values are color coded by their grid block's location, which is mapped out in Figure S10(C). For each ATP concentration, there is no clear trend in the intensity based on block location (with the exception of the corner blocks - the deepest purple and red dots - which are subject to edge effects).

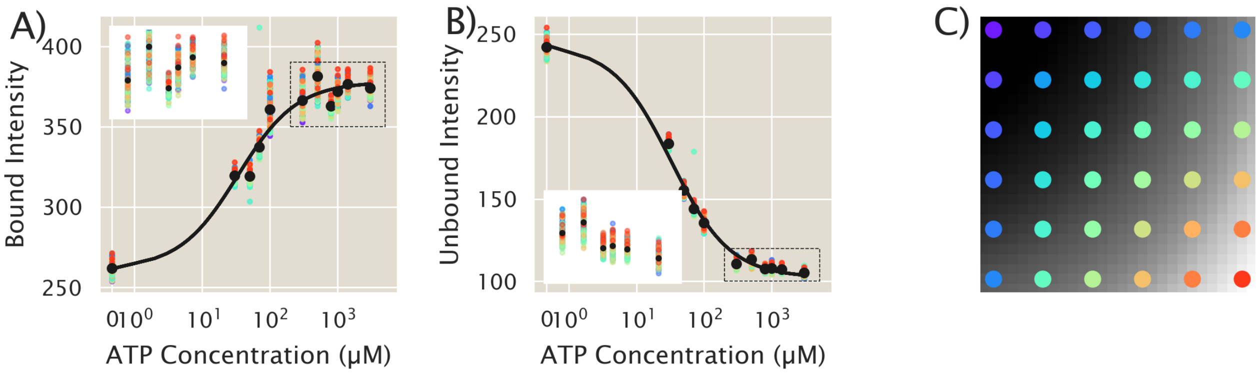

**Fig. S10. After correction, grid block values of homogeneous ATP images do not depend on block location.** The average grid block intensity values for all images are plotted with respect to the ATP concentration of the sample, where A) reports data from the bound ATP channel and B) reports data from the unbound ATP channel. The data points are color coded by the location of their grid block. Black dots are the average value for each ATP concentration. A subset of data is plotted in the white inset of both figure A) and B) for a zoomed in view of the data spread. The data in the inset is denoted by the dotted black rectangle. A map of the grid block location by color is shown in C) overlayed on an unevened ATP image.

We additionally fit intensity versus ATP concentration for each grid block using a Michaelis-Menten curve of the form,

$$I = (I([\mathrm{ATP}] = \infty) - I([\mathrm{ATP}] = 0)) \frac{[\mathrm{ATP}]}{K_{\mathrm{M}} + [\mathrm{ATP}]} + I([\mathrm{ATP}] = 0), \qquad [7]$$

where $I$ is the intensity of the block, and $K_{\mathrm{M}}$ is the Menten constant. We plot the fit parameters versus the block position in Figure S11. Again, there is no significant trend based on the region of the image and the fit parameters are roughly centered about the mean fit parameter.

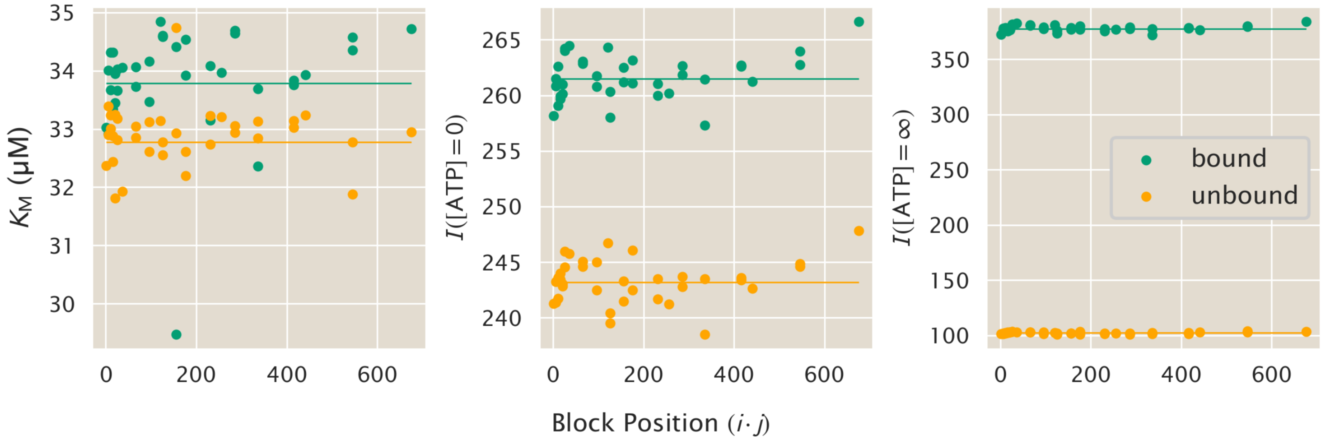

**Fig. S11. Michaelis-Menten fit parameters of the evened image do not have a trend.** We fit the intensity versus ATP concentration to a Michaelis-Menten curve of the form of Equation 7. Plotting the fit parameters versus grid block position, as denoted by multiplying the $i, j$ coordinates of a grid block, we find no trend based on position. The bound ATP channel is plotted in green, while the unbound channel is orange. The fit to the averaged evened images are represented by a horizontal line.

The method we describe works to eliminate gradients created by the optical setup and can thus be applied to any uniform image. If the sample has a real, physical gradient, applying this method would smooth out any existing spatial heterogeneity on top of the gradients imposed by the optical setup. In this study, we form asters that create ATP gradients and we want to ensure our analysis quantifies these gradients without capturing any uneven illumination. Thus, for movies of our experiments, we compute a normalization matrix on the first frame of the movie and use this matrix to correct all subsequent images. The first frame has not been subjected to light that activates structure formation, so this image is homogeneous. To ensure it is valid to correct subsequent images with the normalization matrix of the first frame, we show that the normalization matrices of homogeneous ATP samples are the same across multiple images and across ATP concentrations. In Figure S12, we plot the mean value of the normalization factor for 36 grid blocks distributed through the image, where each data point is a unique image. The standard deviation is very low, a fraction of a percent, compared to the mean value, indicating the same normalization matrix could be used to describe images taken at a variety of ATP concentrations. However, we do note that each light channel should have its own filter correction as the mean normalization values for images taken at the 405 nm and the 480 nm channel emissions differ.

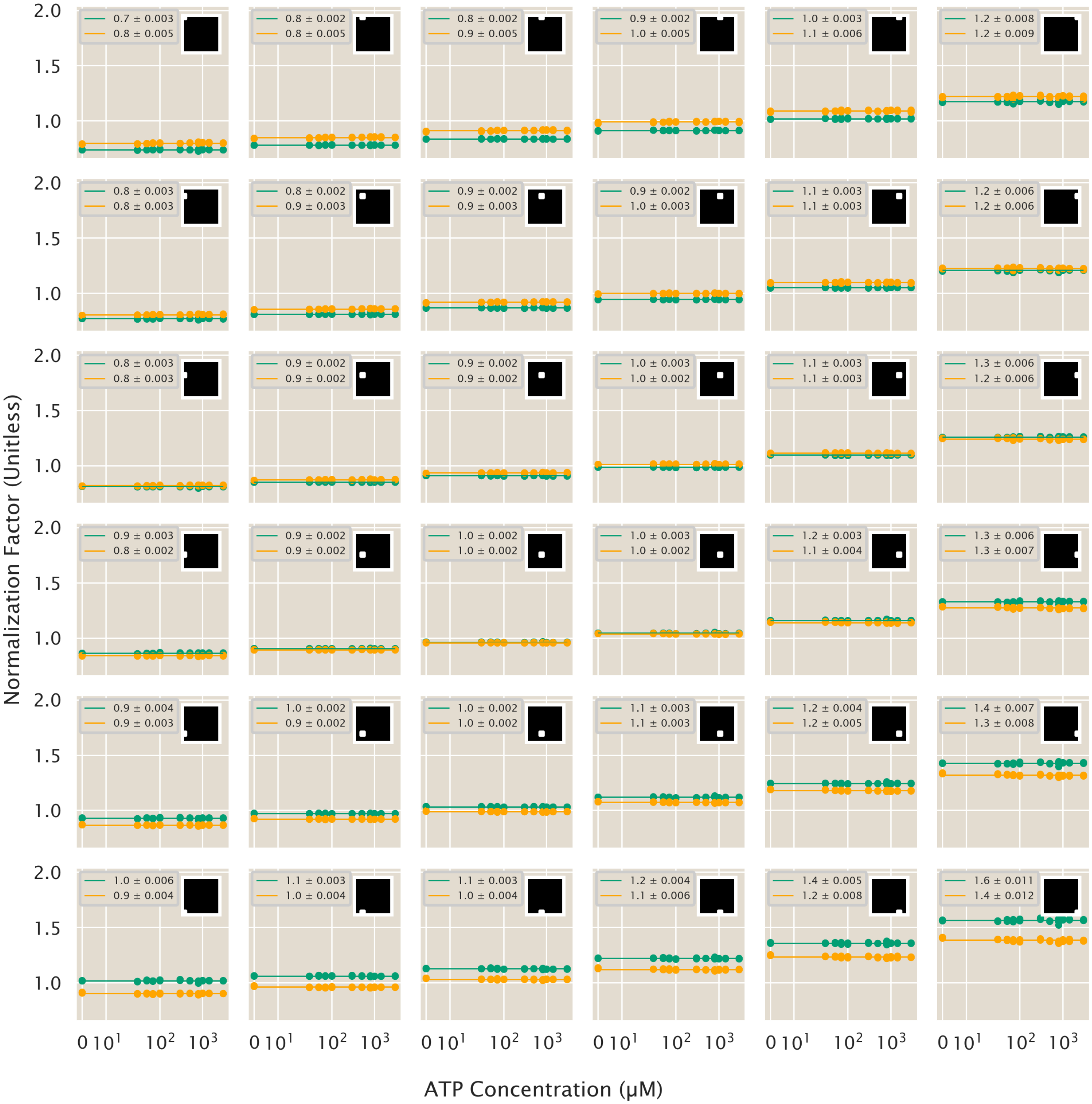

**Fig. S12. The normalization matrices across ATP concentrations report the same value at a given grid block.** Each subplot charts the average intensity value for a given grid block where the inset shows the location of the grid block within the image. The horizontal lines plot the evened image average and the standard deviation is low, a fraction of a percent compared to the mean, indicating the average matrix well represents each image's matrix.

In this data set, we measure the intensity values reported by the ATP probe for a range of known ATP concentrations. Thus, from these images, we can establish a calibration of the intensity values for given ATP concentrations. Taking the ratio of the bound to unbound image intensity values, we fit the data to a Michaelis-Menten function,

$$R = (R_{\max} - R_{\min})\frac{\frac{[\mathrm{ATP}]}{K_{\mathrm{M}}}}{1 + \frac{[\mathrm{ATP}]}{K_{\mathrm{M}}}} + R_{\min}, \quad [8]$$

where $R$ is the intensity ratio of the bound and unbound channels, [ATP] is the concentration of ATP, and $K_{\mathrm{M}}$ is the Menten constant, as shown in Figure S13.

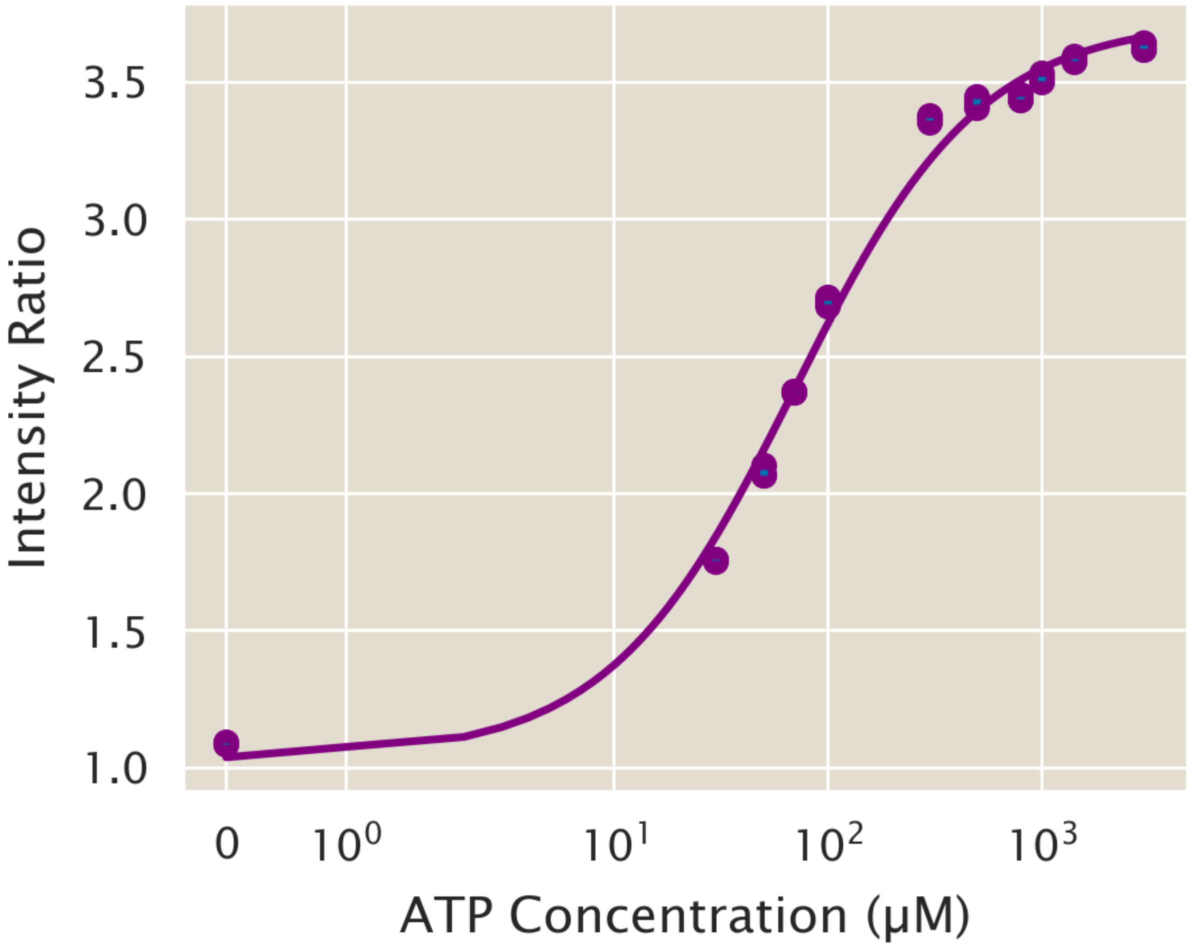

**Fig. S13. ATP Calibration Curve.** Intensity versus ATP concentration values are plotted where each ATP concentration has four replicates. The data is fit to a Michaelis-Menten curve of the form of Equation 8. Here, $K_\mathrm{M} = 70\,\mu\mathrm{M}$, $R_\mathrm{max} = 3.7$, and $R_\mathrm{min} = 1$

We are left to ask what are the error bars on our calibration? In answering this question, we additionally ask how the standard deviation of intensity values scale with increasing ATP concentration. We plot the histograms of intensity values for evened images of varying ATP concentrations in Figure S14. We find that as the ATP concentration increases, so does the mean, while the standard deviation remains nearly constant. We plot a Gaussian distribution given our histogram's mean and standard deviation and find the intensity, $I$, is well predicted by a Gaussian of the form,

$$\frac{1}{\sqrt{2\pi\sigma}} \exp\left(\frac{-(I-\mu)^2}{2\sigma^2}\right). \qquad [9]$$

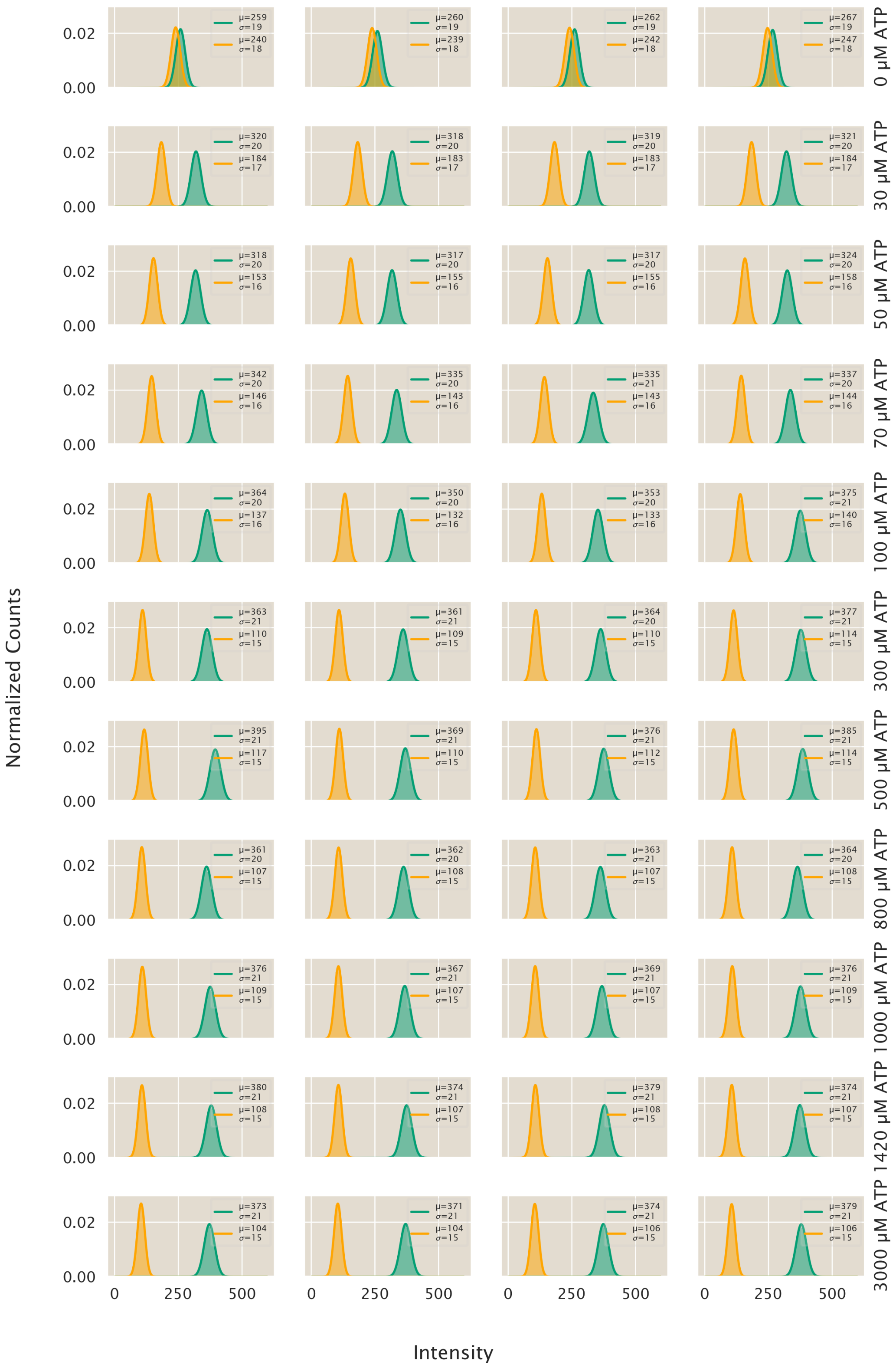

**Fig. S14. Intensity histograms of evened images are Gaussian distributed.** Each row contains intensity distributions for four replicate images at a given ATP concentration. Plugging the mean and standard deviations of the histograms into Equation 9, a Gaussian curve describing the distribution is plotted by the solid line. With increasing ATP concentration, there is little change in the standard deviation relative to the mean. Green curves represent the bound excitation channel, while orange curves represent the unbound excitation channel.

While dividing two Gaussian functions does not result in a Gaussian, we find the ratio of bound to unbound intensities can still be extremely well approximated by a Gaussian, as shown in Figure S15. The histograms of the bound to unbound ratios appreciably widen with increasing ATP concentration, reflecting that as the ATP concentration increases, so too does the mean and standard deviation of empirical ratio values.

Understanding how the standard deviation scales with the mean of the intensity ratio distribution can help inform us how to compute the error of our calibration at various ATP concentrations. In Figure S16, the standard deviation is shown to trend linearly with the mean of the histograms. For the intensity values measured by the bound and unbound channels, there is a very small slope, 0.02, such that the standard deviation is relatively unaffected by the mean. However, for the ratio of these two channels, the slope is, 0.18, which results in notable changes in the standard deviation for our range of ATP concentrations. Thus, our next step will be to use these results to model the error on our ATP calibration fit.

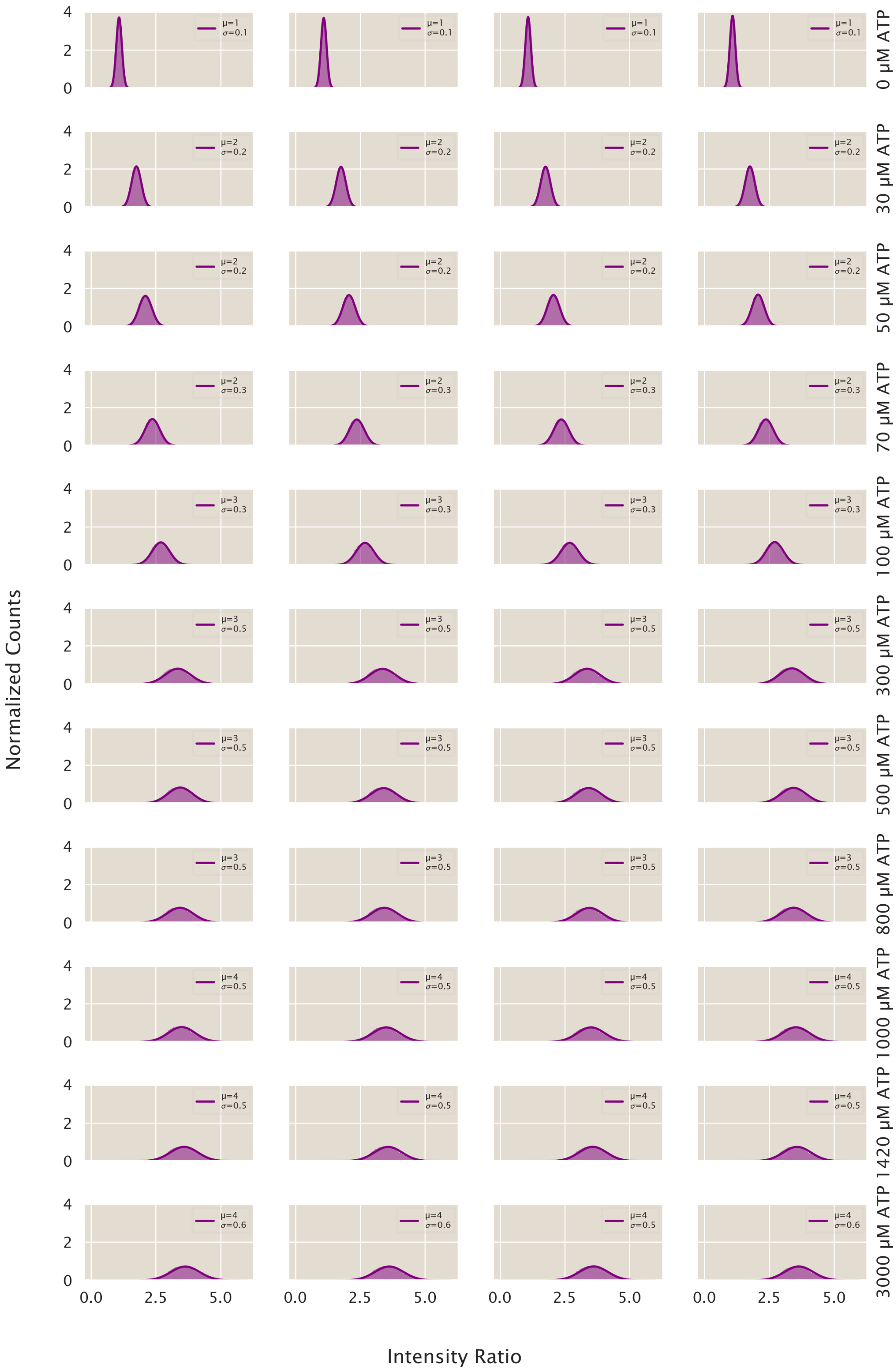

**Fig. S15. Intensity ratios are Gaussian distributed.** Each row contains intensity ratio distributions for four replicate images at a given ATP concentration. Plugging the mean and standard deviations of the histograms into Equation 9, a Gaussian curve describing the distribution is plotted by the solid line. As ATP concentration increases, there is significant broadening of the ratios histogram, such that the standard deviation increases.

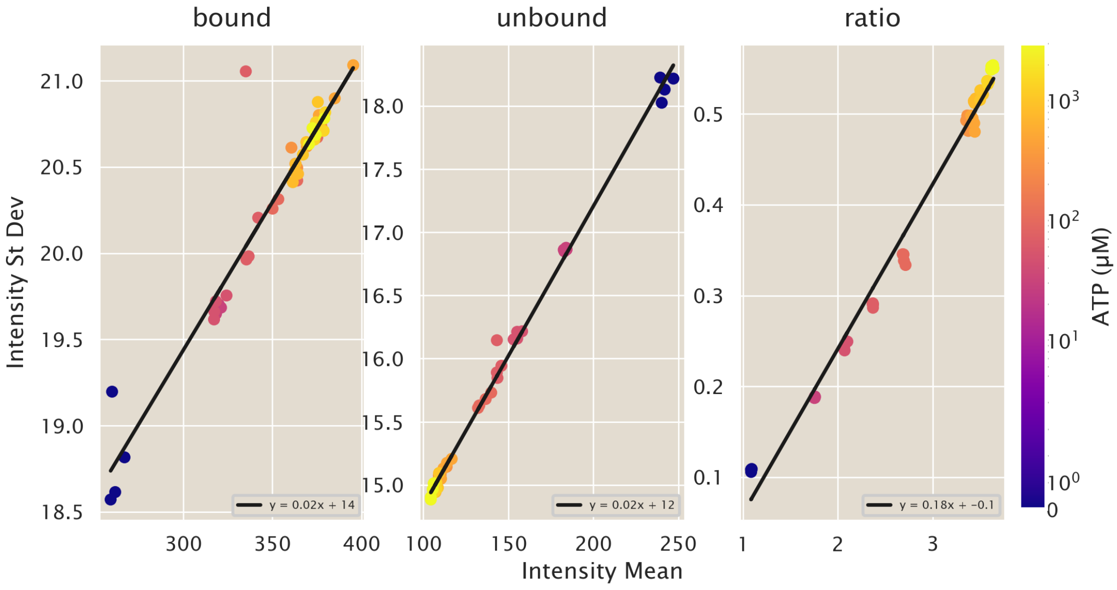

**Fig. S16. The mean and standard deviation of intensity histograms are linearly related.** Here we plot the standard deviation versus the mean across ATP concentrations and linearly fit the data (black line). For both the bound and unbound intensity channels, there is a weak positive slope of 0.02 between the mean and standard deviation of the intensity distributions. However, there is a larger slope of 0.18 for the distribution of the ratio of intensities. Thus, the linear relation describes the broadening of the histograms seen in Figure S15 .

***B.1. Proof the Total Variance is the Sum of the Variances Within and Between Grid Blocks.*** The total variance of the image, can be written as

$$\sigma^2_{\text{tot}} = \frac{1}{N}\sum_{i}^{N}(I_i - \langle I\rangle)^2 \qquad [10]$$

where $i$ sums over each pixel, $I_i$ is the intensity value of a given pixel, and $\langle I\rangle$ is the average intensity value of all pixels in the image. To understand if most of the variance in the images occur within given regions of the image or between regions of the image, we divide the image into a grid with $B$ blocks each containing $N_B$ pixels. Inside of a given grid block, indexed $j$, the variance is

$$\sigma^2_{\text{in}} = \sigma^2_{\text{j}} = \frac{1}{N_B}\underbrace{\sum_{i}^{N_B}}_{\text{in } j}\left(I_i - \langle I\rangle_j\right)^2, \qquad [11]$$

which sums over all pixels, $i$, that are within block $j$, and where $\langle I\rangle_j$ is the average of pixel intensity values within block $j$. Finally, we define the variance between blocks as,

$$\sigma^2_{\text{btwn}} = \frac{1}{B}\sum_{j}^{B}\left(\langle I\rangle_j - \langle I\rangle\right)^2. \qquad [12]$$

Let us rewrite the form of the total variance to see how it relates to the variance inside and between blocks,

$$\sigma^2_{\text{tot}} = \frac{1}{B}\sum_{j}^{B}\frac{1}{N_B}\underbrace{\sum_{i}^{N_B}}_{\text{in } j}(I_i - \langle I\rangle)^2. \qquad [13]$$

Expanding the square yields,

$$\sigma^2_{\text{tot}} = \frac{1}{B}\sum_{j}^{B}\frac{1}{N_B}\underbrace{\sum_{i}^{N_B}}_{\text{in } j}\left(I_i^2 - 2I_i\langle I\rangle + \langle I\rangle^2\right). \qquad [14]$$

We add and subtract $-2I_i \langle I \rangle_j + \langle I \rangle_j^2$ to simplify terms to the form of inside block variance. Highlighting terms in red that become the inside block variance,

$$
\begin{aligned}
\sigma_{\text{tot}}^2 &= \frac{1}{B}\sum_j^B \frac{1}{N_B}\underbrace{\sum_i^{N_B}}_{\text{in } j}\left(I_i^2 - 2I_i\langle I\rangle + \langle I\rangle^2 - 2I_i\langle I\rangle_j + \langle I\rangle_j^2 + 2I_i\langle I\rangle_j - \langle I\rangle_j^2\right) \\
&= \frac{1}{B}\sum_j^B \frac{1}{N_B}\underbrace{\sum_i^{N_B}}_{\text{in } j}\left(\left(I_i - \langle I\rangle_j\right)^2 - 2I_i\langle I\rangle + \langle I\rangle^2 + 2I_i\langle I\rangle_j - \langle I\rangle_j^2\right) \\
&= \frac{1}{B}\sum_j^B\left(\sigma_j^2 + \frac{1}{N_B}\underbrace{\sum_i^{N_B}}_{\text{in } j}\left(2I_i\left(\langle I\rangle_j - \langle I\rangle\right) - \langle I\rangle_j^2 + \langle I\rangle^2\right)\right)
\end{aligned}
\tag{15}
$$

Bringing all terms that do not depend on $i$ out of the first sum,

$$
\sigma_{\text{tot}}^2 = \frac{1}{B}\sum_j^B\left(\sigma_j^2 - \langle I\rangle_j^2 + \langle I\rangle^2 + \frac{2}{N_B}\left(\langle I\rangle_j - \langle I\rangle\right)\underbrace{\sum_i^{N_B}}_{\text{in } j} I_i\right). \tag{16}
$$

Note that $\frac{1}{N_B}\underbrace{\sum_i^{N_B}}_{\text{in } j} I_i$ is simply $\langle I\rangle_j$, so we can write

$$
\sigma_{\text{tot}}^2 = \frac{1}{B}\sum_j^B\left(\sigma_j^2 - \langle I\rangle_j^2 + \langle I\rangle^2 + 2\left(\langle I\rangle_j - \langle I\rangle\right)\langle I\rangle_j\right). \tag{17}
$$

With some algebra, we can simplify our expression,

$$
\begin{aligned}
\sigma_{\text{tot}}^2 &= \frac{1}{B}\sum_j^B\left(\sigma_j^2 - \langle I\rangle_j^2 + \langle I\rangle^2 + 2\langle I\rangle_j^2 - 2\langle I\rangle_j\langle I\rangle\right) \\
&= \frac{1}{B}\sum_j^B\left(\sigma_j^2 + \langle I\rangle_j^2 - 2\langle I\rangle_j\langle I\rangle + \langle I\rangle^2\right) \\
&= \frac{1}{B}\sum_j^B\left(\sigma_j^2 + \left(\langle I\rangle_j - \langle I\rangle\right)^2\right) \\
&= \frac{1}{B}\sum_j^B \sigma_j^2 + \frac{1}{B}\sum_j^B\left(\langle I\rangle_j - \langle I\rangle\right)^2.
\end{aligned}
\tag{18}
$$

Interestingly, the second term on the right hand side is exactly Equation 12, the variance between blocks,

$$
\sigma_{\text{tot}}^2 = \frac{1}{B}\sum_j^B \sigma_j^2 + \sigma_{\text{btwn}}^2. \tag{19}
$$

We can interpret the first term as the average of the variances within blocks,

$$
\sigma_{\text{tot}}^2 = \left\langle\sigma_{\text{in}}^2\right\rangle + \sigma_{\text{btwn}}^2 \tag{20}
$$

Thus, the total variance in the image is average variance inside blocks plus the variance between blocks. Now, we have a direct way to determine how much variance is between blocks, implying uneven illumination.

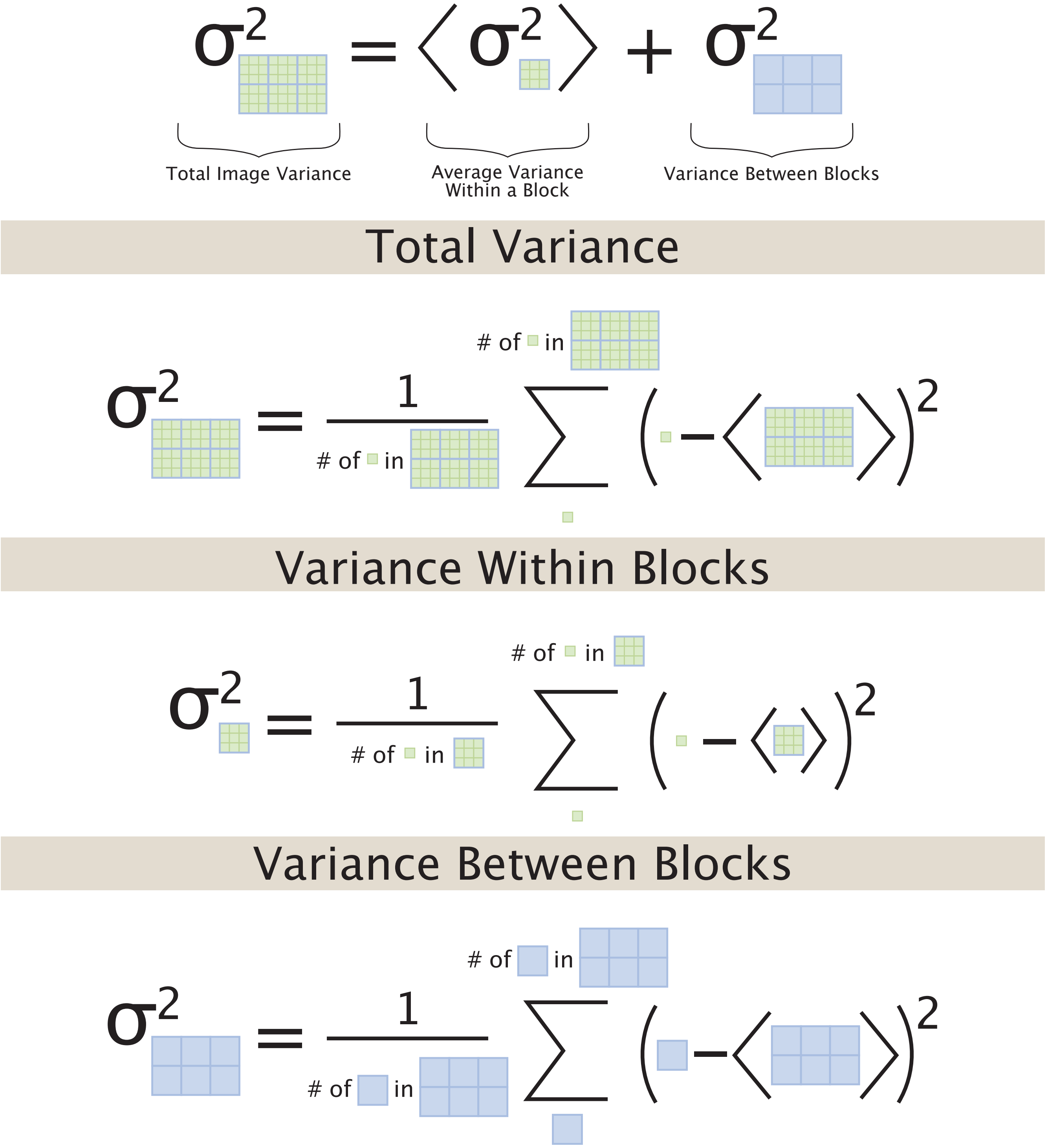

**Fig. S17. Graphical representation of variance between and within blocks.** The top panel is the equation for the total variance in terms of the variance within a block and between blocks. The subsequent three panels graphically depict the definition of each type of variance, where a single green block represents one pixel, $i$, and a blue block represents one grid block, $j$, containing $N_B$ pixels.

**C. Correcting time courses for photobleaching.** Imaging time courses of ATP calibration images, as experimentally prepared as described in section D, we find the form of ATP probe photobleaching, which we describe in detail in section 6. In brief, we model the form of photobleaching as a single exponential plus a constant in time, as written in equation 43. Fitting ATP calibration experiments, described in section D, with many measured time points to equation 43, we establish a correction factor that depends on the time of image acquisition and the interval of time between acquisitions, see Figure S45. We divide all pixel intensity values of a given image by the scalar correction value given by equation 43 at the time point of the image, as described by equation 48. The exponential and constant parameters used are determined by the fit values at the 20 second acquisition interval shown in Figure S45.

**D. Converting intensity units to concentration units.** As described in section D, ATP data is collected via imaging in both the 405 nm and 480 nm excitation channels. In order to describe ATP in concentration units, we first divide the pixel intensity values element-wise between the two images, as written in equation 1. Then, we can invert the equation for the calibration curve, equation 2, to convert ratio units to ATP concentration units, giving the conversion,

$$[ATP] = K_d \frac{R - R_{\text{min}}}{R_{\text{max}} - R}. \quad [21]$$

We note that this conversion will yield infinity or "not-a-number" values for ratios greater than $R_{\text{max}}$ or give negative ATP concentrations for ratios less than $R_{\text{min}}$. Thus, we bound the ratio values setting any ratio greater than $R_{\text{max}}$ equal to $R_{\text{max}}$ and any values less than $R_{\text{min}}$ to $R_{\text{min}}$. While most values fall within the physical ratio range, stochasticity in image acquisition gets magnified by dividing collected intensity values meaning some pixels yield ratios that are nonphysical.

We then apply the conversion in equation 21 to each pixel of the ratio image.

**E. Motor Image Analysis.** Given that our primary objective in this work is to accurately measure ATP in our experiment, we have spent considerable effort detailing how to quantitatively analyze fluorescent images. These results are, of course, better understood by simultaneous measurements of motor proteins, given that the motors are the ATP consuming element in the system. Thus, we need to apply similar analysis schemes to quantify the motor proteins in our collected fluorescent images. While the majority of our analysis is analogous to that done for the ATP images, there is one significant difference in how we analyze the motor channel.

When imaging ATP, our fluorescent probe is excited by two light excitations at wavelengths that overlap with the motor dimerization light wavelength. Due to this quirk of the experiment, we shine the light that excites the ATP probe and dimerizes the motors in a small circular subset of the camera detector's field of view. In contrast, the wavelength of light that excites the motor's fluorophore has no impact on motor dimerization, so we collect a full field image at the size of the camera detector. This means there is a large border region where no structure formation occurs since motors are undimerized, however, we still collect images of motor fluorescence, as diagrammed in Figure S18. These regions are useful to us in this analysis, since we can use these as homogeneous images from which to compute the dynamics of motor photobleaching and create a calibration curve to turn motor fluorescence intensity into concentration units. When we analyzed ATP images, we needed separate datasets from the aster datasets, since the overlapping fluorescence and motor dimerization channels meant we could not obtain homogeneous ATP samples in an aster dataset. Having separate datasets has its merits, predominantly, the experimentalist need not worry about gradients in the fluorophores due to advection or diffusion into the circular aster excitation region, we address this concern for the motor analysis in section E.2. On the other hand, it can be convenient to have bespoke calibrations for each unique aster, which we have the opportunity to measure here.

***E.1. Motor Photobleaching.*** Applying the same principles from Section A, we analyze how the fluorescence of motor proteins change over the course of the experiment. As done previously, we subtract the background signal from camera noise (Section A) and apply an uneven illumination correction (Section B). We then average the intensities of the images for each time point and plot the points over time. Once again, we find that fits to a single exponential plus a constant best represent our data given that the motors are in a reservoir, as discussed in Section D.1.

In Figure S19 we show the photobleaching fits for many asters spanning a range of motor concentrations. These fits demonstrate a time constant of approximately 1400 seconds or just over 20 minutes. This is a significantly longer time constant as compared to the ATP probe which boasted a very fast time constant of approximately 20 seconds. This implies the associated fluorophores with these proteins undergo different photophysics from the ATP probe. When examining the fit of the constant plateau value, we find the intensity only drops on average 9%. This is less of a drop than the ATP probe.

We compare the variation in fits across experiments and find in Figure S20(B) and (C), there is no trend in the fit parameters with the starting motor concentration. This is to be expected since the rate of bleaching should depend on intrinsic properties of the fluorophore rather than the number of fluorophores. In Figure S20(A), we show all the individual fits and the fit parametrized by the average of all fit values. By the end of all the experiments, where any variations are most prominent, the range of fitted photobleaching curves is about 6%. This is not a significant range, so we do not correct for motor photobleaching in our experiments.

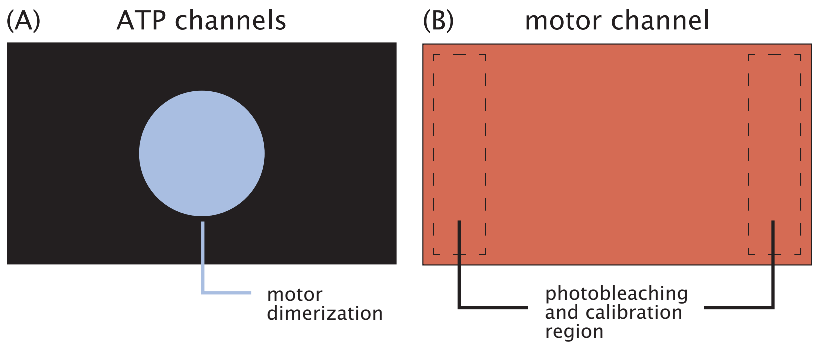

**Fig. S18. Geometry of light excitations.** We illustrate the patterns of light excitations for the different wavelengths of light. (A) We depict the blue wavelengths that both dimerize motors and excite the ATP probe. This light is only projected in a circle at the center of the image. (B) We show the red light used illuminates the entire image. We outline the region where we perform the motor photobleaching and calibration analysis in dashed lines.

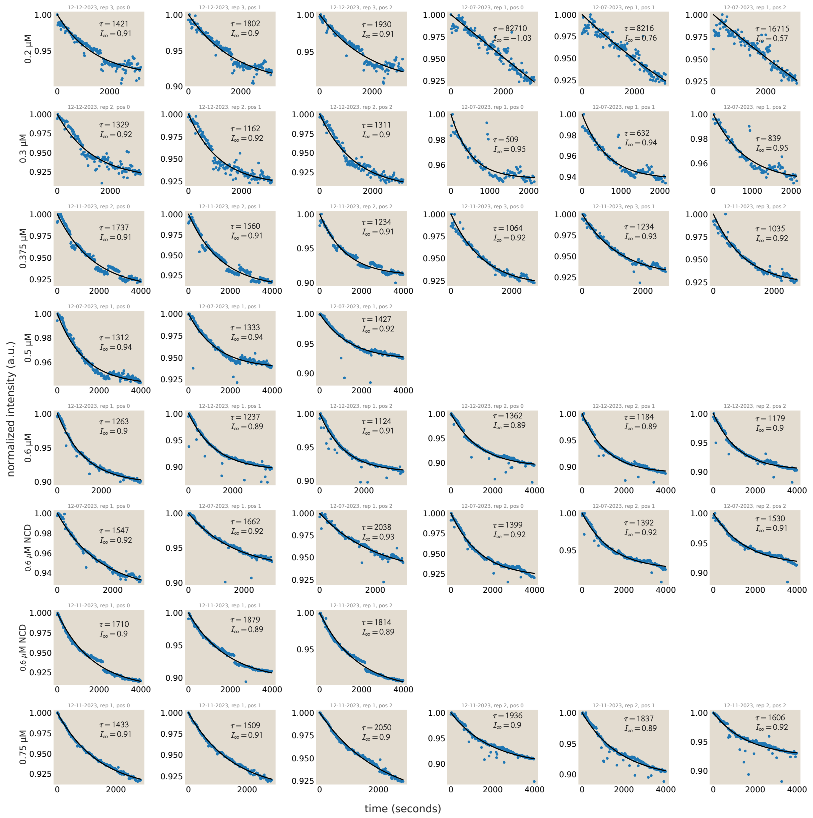

**Fig. S19. Photobleaching dynamics of motor proteins can be explained by a single exponential plus a constant.** Here, we compiled the photobleaching fits for asters that vary in starting motor concentrations. When fitting, the intensity scales are normalized by the maximum value in the plot. The dots represent the averaged value of images at each time point and the black line shows the single exponential plus a constant fit.

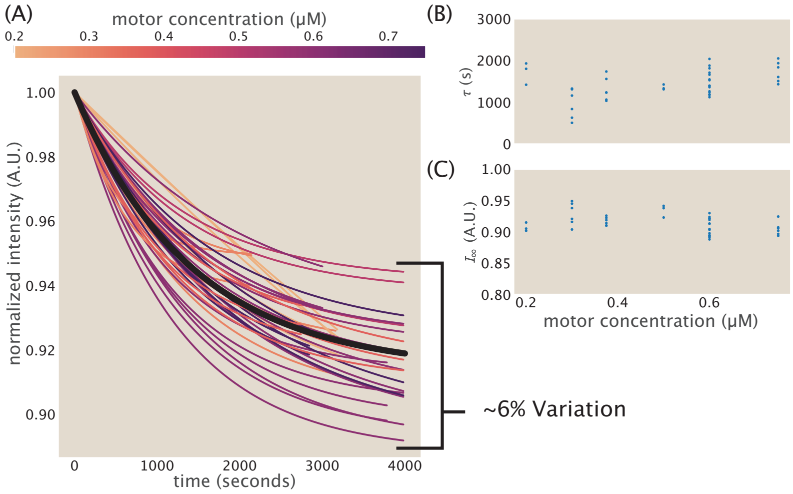

**Fig. S20. Variation in motor photobleaching fit curves.** (A) The fits for each aster are plotted colored by the starting motor concentration of each aster. The black line plots the fit as determined from averaging the values of the fit parameters. We see by the end of aster formation, there is about a 6% variation in the multiplicative value of the fit correction. (B) There is no trend in the decay constant fit parameter with motor concentration. (C) There is also no trend in the plateau constant value with motor concentration.

***E.2. Spatial inhomogeneities do not impact our analysis method.*** Since we perform our analyses upon the border region of aster movies, we may be concerned that spatial inhomogeneities created by advection or diffusion of motors into the aster region may confound our photobleaching measurements. We perform a simple test to assess how concerned we should be about spatial inhomogeneities.

First, noting that the regions along the border that we analyze are to the left and right of the circular region (see Figure S18(B)), we average the intensities of pixels in each column of our analyzed region. We then plot the average intensities as a function of the column's location. This gives us a proxy for how the intensity of motors changes moving from the boundary of the image toward the center of the aster as depicted in Figure S21(A). We examine this trace for both the first frame and the last frame. We can see that intensity increases moving towards the aster region and plateaus. Despite us doing uneven illumination corrections, this could be due to edge effects. But importantly, the shape remains the same between the first and last frame. If motor gradients were to arise over the course of aster formation, these profiles would differ along the position coordinate. We can look at this more quantitatively by taking the relative change in the profiles at each position. In Figure S21, we plot this relative change as

$$\frac{I(x,t_0) - I(x,t_f)}{I(x,t_f)}, \quad [22]$$

where $I$ is the pixel intensity, $x$ is the axis along which we measure, $t_0$ is the first time point and $t_f$ is the final time point. We find that across positions the relative change is close to uniform as shown in Figure S21. This is indicative of photobleaching, since the development of a gradient should non-uniformly change the motor intensity values corresponding to the regions where motors have been enriched. In Figures S22 and S23, we plot the percent change for each aster's left and right regions to get a sense of the variation between aster preps.

***E.3. Motor Calibration.*** Following the aforementioned procedures for image analysis, we subtract camera noise and perform uneven illumination corrections. Since we are examining the boundary regions of aster movies of varying motor concentrations, all that is left to do is to simply average the measured intensity values of the first time point of aster movies, while motors are nominally still homogeneous. We plot the average of each individual aster experiment as a function of the initial motor concentration in Figure S24. We find that the data is fit nicely by a linear function.

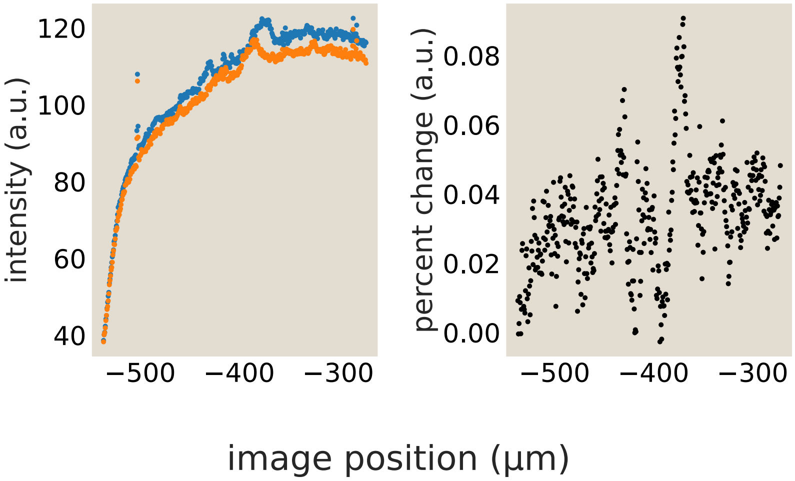

**Fig. S21. Determination of intensity percent change across space in the far-field image regions.** (A) The intensity across space in the far left region of the image. The x-axis is defined by setting the center of the image to zero. The blue curve reports the intensity of the first image while the orange curve reports the intensity in the last image acquired. (B) The percent change as defined by Equation 22 is plotted based on the intensities plotted in (A).

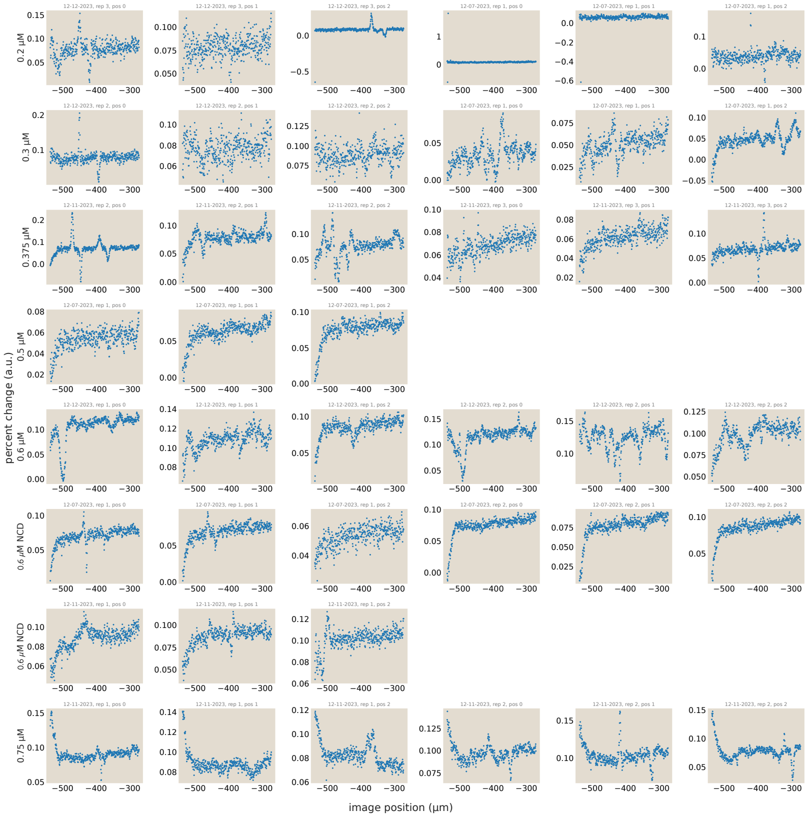

**Fig. S22. Variation in left, far-field intensity percent changes.** The percent change based on the first and last image, as diagrammed in Figure S21 is computed for the left region of each aster.

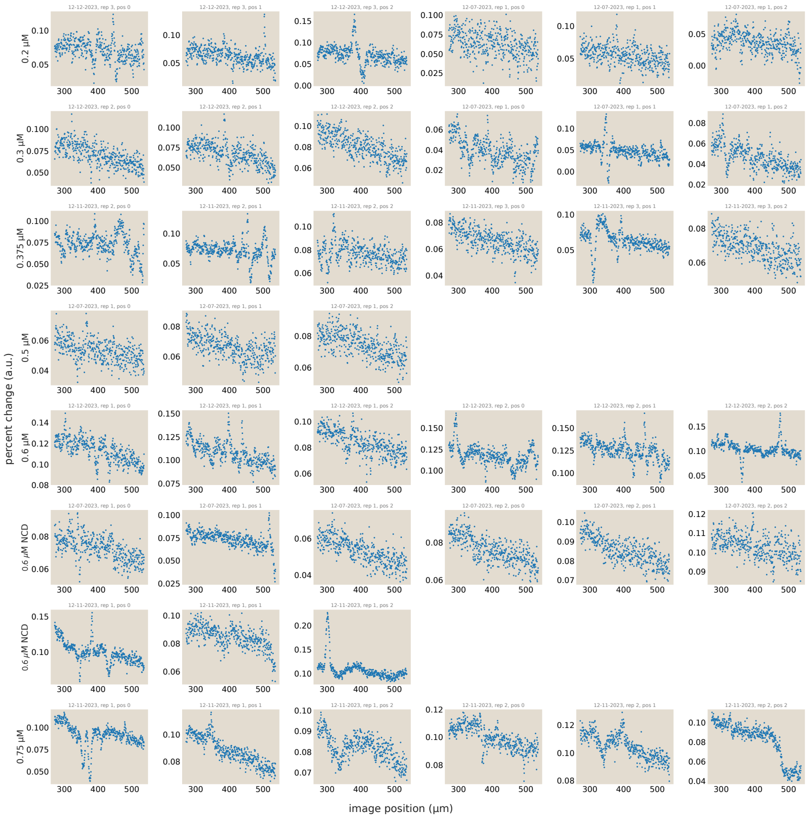

**Fig. S23. Variation in right, far-field intensity percent changes.** The percent change based on the first and last image, as diagrammed in Figure S21 is computed for the right region of each aster..

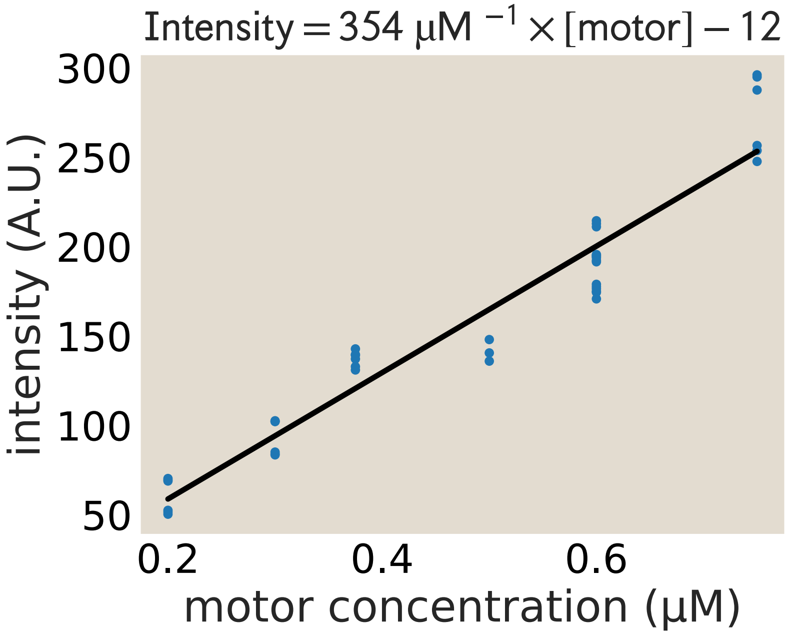

**Fig. S24. Motor intensity to concentration calibration.** Using far-field regions of motor images, we average the intensity values of the first frame for asters at varying starting motor concentrations. We assume all motors are homogeneous in the first image. The intensity scales linearly with concentration.

**F. Center Tracking.** To accurately determine how concentrations change with the aster radius, it is important to accurately identify the center of an aster. Images of fluorescently labeled motor proteins in asters generally show a bright core region with fainter arms. Especially while developing, asters show diffuse, ovular boundaries. These boundaries of asters are not always geometrically contiguous or universally high in contrast with the background over the aster region. These features fundamental to real asters deeply complicate the use of automated segmentation and tracking of asters using even sophisticated image analysis pipelines and thresholding procedures. Specifically, for instance, we built and assessed a battery of semisupervised thresholding and tracking pipelines, but all methods failed to achieve satisfactory identification of asters and positions across the experimental conditions and replicates we measured. Accordingly, to ensure that this segmentation and tracking is performed robustly for every dataset, we performed manual segmentation and tracking of all asters as they formed across all data. Specifically, to accomplish this, we drew elliptical boundaries of asters over a dense number of key frames in each aster movie (using an instance of the open source Computer Vision Annotation Tool (CVAT)), and where relevant performed simple temporal interpolation of these aster boundaries between explicitly tracked key frames. This produced excellent and internally consistent tracking results.

**G. Local averaging of ATP images.**

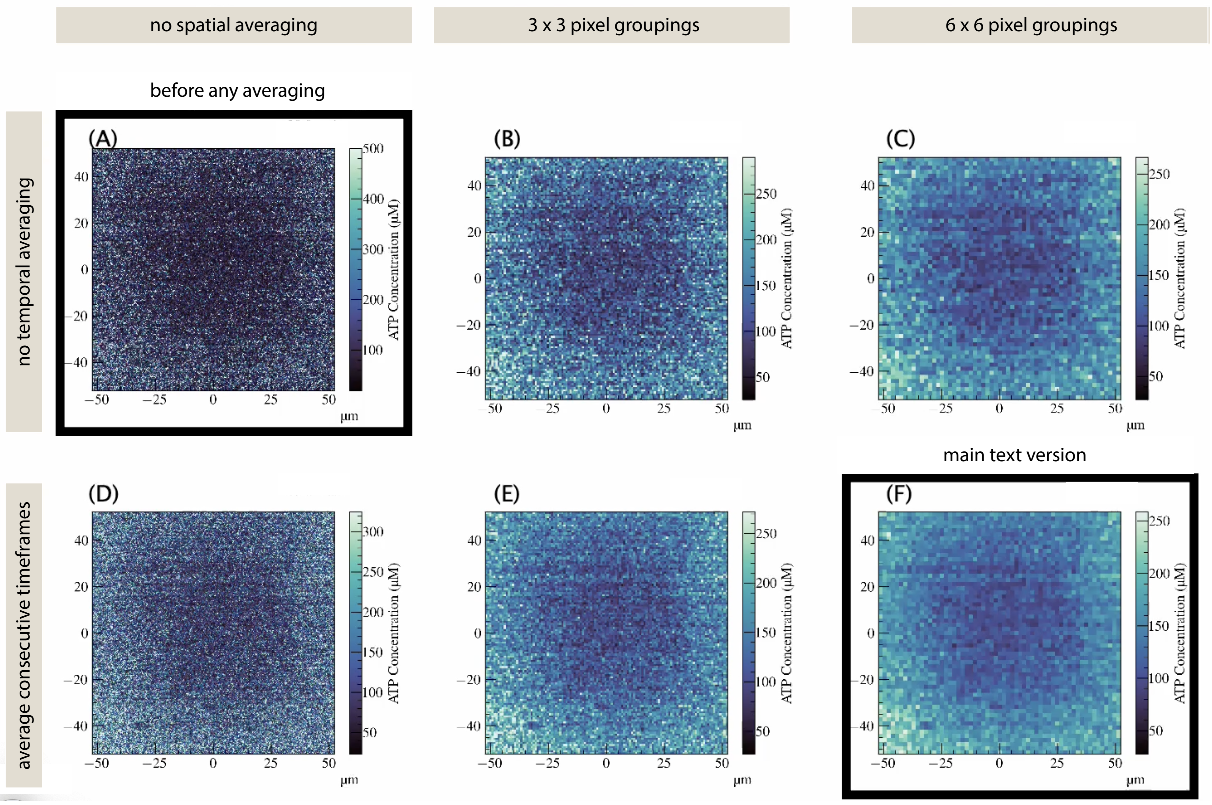

**Fig. S25. Visualization of averaging images in space and time.** Images (A), (B), and (C) are not averaged in time while images (D), (E), and (F) are the results of averaging two consecutive images in time. The columns demonstrate how spatial averaging impacts the images. (A) and (D) have no spatial averaging. Each pixel in (B) and (E) are the results of averaging a 3x3 group of pixels from the raw image. Each pixel in (C) and (F) are the results of averaging a 6x6 group of pixels from the raw image. (A) is an image before any averaging occurs and (F) is the same image after the space time averaging procedure we use for data in the Main Text is applied.

Acquiring microscopy images is subject to detector noise and other stochasticity. We find that the concentration images of the ATP probe are especially susceptible to pixel to pixel variation. To both better visualize concentration gradients and to assess quantitative trends more cleanly, we locally average our images to reduce noisy fluctuations. We first average in space by replacing the concentrations of individual pixels in a group with the average concentration of all the pixels in the group. If we take an $N \times N$ group of pixels as the spatial averaging window, then an $m \times n$ image will have dimensions $m/N \times n/N$. If an image was not already a multiple of our group size, we cropped the image evenly on both sides. We tried several grouping sizes and selected to use a group size of 6 px × 6 px. See Figure S25 for an example of various spatial averaging group sizes.

We also modestly average ATP images over time. For this, we do a running average, replacing each pixel value of images of adjacent timepoints with the average value of that pixel location. Here and in plots we average over just two consecutive images that were taken 20 seconds apart. Thus for an original set of images over time of size, $l$, we now have $l - 1$ timepoint images.

#### H. Computing azimuthal averages for radial concentration profiles.

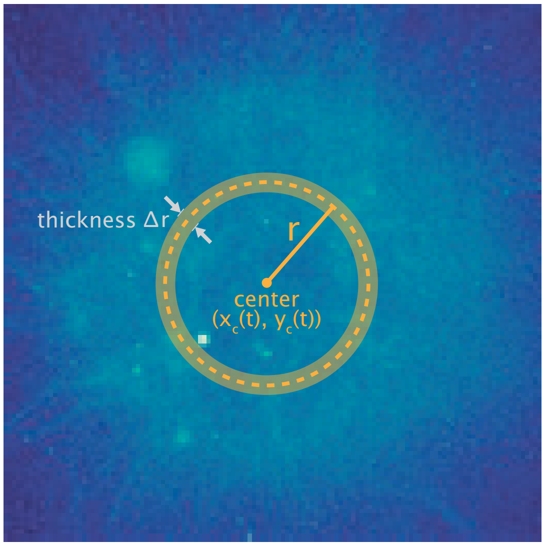

**Fig. S26.** Schematic of how average radial concentrations are calculated from aster images. The ATP or motor concentration at a radius $r$ away from the aster's current center reflects the angularly-averaged value of concentrations in pixels found within an annulus of finite width $\Delta r$ centered at radius $r$.

Given inferred local ATP and motor concentrations at each pixel position $(x, y)$ in an image plane, these profiles and their emerging gradients become more illuminating when expressed with respect to the natural coordinates of the aster. Specifically, since tracking annotations locate the center $(x_c(t), y_c(t))$ of each aster as it contracts and moves over time $t$, it is useful to summarize the average ATP concentration $\langle\text{ATP}\rangle(r, t)$ and motor concentration $\langle m\rangle(r, t)$ shown by points at a radial separation $r \equiv \sqrt{(x - x_c(t))^2 + (y - y_c(t))^2}$ away from the center. (Naturally, given the circular light pattern that excites their activity, asters are largely angularly symmetrical around their centers; our tracking annotations can resolve modest and transient elliptical eccentricity, but these departures from circularity are very modest and justify a polar coordinate system.) Specifically, let $r_{\max}$ be largest radial separation of interest for an aster (which we take as the largest (namely, initial) radius the aster adopts over all times up to a final time). We average concentrations inside each annulus in a fixed set of $m$ radial positions spanning the center to this farthest edge $r_{\max}$ of the aster. That is, the $i$th annulus is centered at a radius $r_i = i\Delta r$, with a width $\Delta r$, where these radial bin widths are defined as $\Delta r \equiv \left\lfloor \frac{r_{\max}}{m} \right\rfloor$). This geometry, commonly used to generate the radial profiles reported throughout the main text and this supplement, is illustrated in Fig. S26.

**I. Computing total power consumption in asters.** The rate at which ATP decreases inside the volume of an aster reflects the total dissipation of free energy powering its active dynamics at a given time. (Specifically, see §J for a discussion about the connection of this raw observable to total underlying power consumption rates.) To estimate the ATP consumption rate over a whole aster at a given time, we use a simple first-order finite difference. Specifically, our spatial maps of ATP concentration can be multiplied by the experimental observation volume $v$ (in the flow cell and microscope) to give the total number of ATP molecules at a given two-dimensional pixel position; integrating all pixels inside the boundary of an aster at a given time yields the aster's present total ATP; and subtracting this total ATP at two consecutive timepoints estimates the power. With a uniform imaging height $h$ and a pixel area $dA$, the observation volume is $v = dA\ h$, giving,

$$\text{power}(t) \approx h \frac{\int_{\text{aster}} dA\ ([\text{ATP}](\mathbf{r}, t + \Delta t) - [\text{ATP}](\mathbf{r}, t))}{\Delta t}. \qquad [23]$$

An exactly equivalent scheme to calculate such a power, convenient once armed with radial profiles as described by $S$H, is to radially integrate the ATP concentration over all radii spanning the aster's volume, namely

$$\text{power}(t) \approx h \frac{\int_{\text{aster}} (2\pi r\ dr)\ ([\text{ATP}](r, t + \Delta t) - [\text{ATP}](r, t))}{\Delta t}. \qquad [24]$$

The geometry of this simple calculation is illustrated in Fig. S27.

We remark that in general, more precisely estimating derivatives of noisy data can be fraught with substantive technical subtlety (9). We refrain from sophisticated filtering or inference schemes in favor of this simple finite difference scheme for transparency (and reducing obfuscating effects from parameter-dependent filtering schemes), and because this scheme suffices to localize both the order of magnitude and the broad temporal trends of these experimental consumption rates. As an additional guide to the eye, a short moving average in time is also provided as black lines in figures such as Fig. S28 along with raw values. We expect that for other investigators or scientific questions, more sophisticated signal processing operations could be justified if greater precision is required to resolve some of these dynamics.

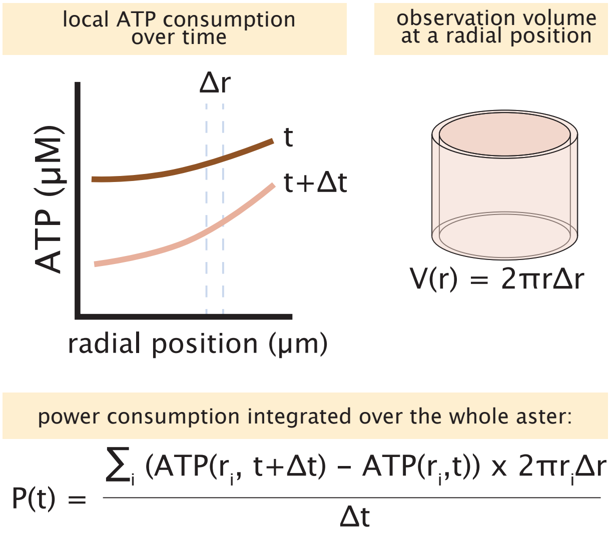

**Fig. S27.** Illustration of simple scheme to estimate the ATP consumption rate over a whole aster's volume.

**J. Measured ATP consumption rates are lower bounds on aster power consumption rates.** Here we further motivate why net changes in ATP in an aster's volume may be understood as lower bounds on true underlying consumption rates, as discussed in the main text's *Results* section.

Let the total number of ATP molecules (in a specified aster's volume) be $\mathrm{ATP}(t)$. Let the motors inside this volume consume ATP at some true hydrolysis rate $h(t)$, which actually would be proportional to the free energy dissipation rate of greatest interest accomplishing such functions as mechanical work, maintenance of gradient, etc. However, an aster is not perfectly separated from the rest of the biochemical reservoir: at its boundary, we expect replenishment from diffusive influx countering the small gradients in ATP concentration manifested at these edges, plus any replenishment from inward fluid flow. For such an aster with a positive radial concentration gradient and inward contractile fluid flow, both of these transport processes should have positive sign countering the negative sign of the consumption rate: let their total impact be $r(t)$. Then the total net rate of change in ATP, namely what we measure, is simply,

$$\frac{d\mathrm{ATP}}{dt} = h(t) - r(t). \qquad [25]$$

So, the true consumption rate is $h(t) = r(t) - \frac{dATP}{dt}$. In other words, only when net replenishments are negligibly small, $r(t) \to 0$, would our literal measured rate-of-change in ATP $-\frac{d\mathrm{ATP}}{dt}$ give exactly the consumption rate $h(t)$. Otherwise, when small net replenishment $r(t) > 0$ occurs, the consumption rate $h(t)$ is lower bounded by the directly observed rates of change $-\frac{d\mathrm{ATP}}{dt}$. We note that the big question of what functions energy expenditures pay for, given that a number of candidate physical answers have much smaller apparent dissipative costs than measured rates-of-change $-\frac{d\mathrm{ATP}}{dt}$ as explored by the main text, is only made more serious by these measurements being a lower bound on the true underlying dissipation rate.

Of course, this reasoning depends on some fortuitous properties of these dynamics and geometry. In general, diffusive contributions could have either replenishing or reductive effects (depending on the accumulated net signs of gradients over a volume's boundaries that set the net diffusive current), and more complex fluid flows than manifested here in contracting asters may similarly accomplish either effect.

#### 3. Empirical variability and uncertainty in aster behavior and dissipation

**A. Notions of uncertainty.** Detecting biomolecules like ATP using measurements from optical fluorescent probes, as we do here in a much wider tradition of modern biology, is ultimately an act of inference with many sources of stochasticity and uncertainty. We use a foundation of layered control and calibration experiments to accomplish this inference accurately and control governing parameters as tightly as feasible, as described in wide detail throughout this Supplementary Information. Ultimately, however, while some sources of persistent uncertainty manifest conspicuously in tractable forms (e.g. in the form of estimated (co)variances in calibration curve fit parameters), others (such as variations in optical stability, biochemical preparation and function beyond experimental control, spurious noise from numerical derivatives, etc.) are more insidious, though likely even more impactful. Given these ambient sources of uncertainty, and our focused set of coarse observables we care most about identifying accurately in this work (namely, the orders of magnitudes of ATP gradients in space and time), a more earnest representation of the accumulated operational uncertainty in our measurements is given by the dispersion of these phenomenologies across macroscopic replicate aster formation experiments. We interpret the dispersion in these measurements, e.g. the variation manifested across replicate applications of the whole inference procedure, to more sincerely reflect uncertainty on the core numbers of interest reported in this work than a superficially-formal technical propagation of only the most tractable errors. (In particular, the manifold challenges intrinsic to estimating numerical derivatives of noisy data forewarn that too superficially precise an error accounting before numerical derivatives are taken is likely quixotic. Aggregated variation across experiments of such numbers more sincerely reflect their variation.)

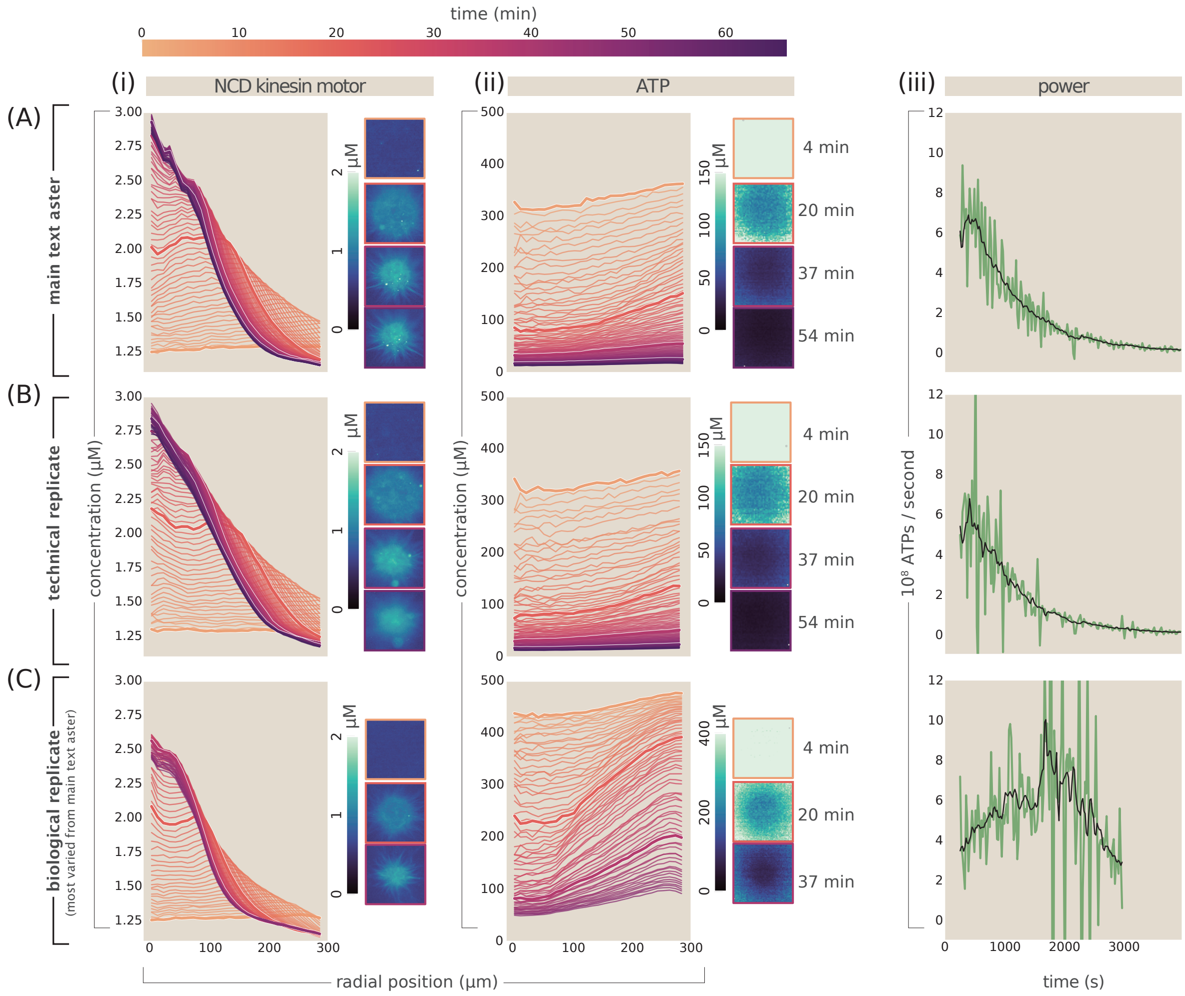

**Fig. S28.** Characteristic variation in aster formation. (A) The motor profile (i), ATP profile (ii), and power trajectory (iii) for the characteristic aster trajectory discussed most extensively in the main text. (B) A separate aster with representative motor, ATP, and dissipation behaviors; this technical replicate of the aster discussed in the main text was prepared in a separate position in the experimental flow cell in a separate optical excitation of motors to initiate self assembly, but using the same biochemical reaction mixture. These trajectories reflect a scale of how coherently typical asters behave. (C) Trajectories from a third aster, chosen intentionally to highlight the largest type of variation in behavior from the typical aster referred to in the main text. This third aster was formed using a completely separate biochemical preparation on a separate date (and so form what can be considered a "biological replicate"). These three asters join a fuller ensemble of replicates displayed in Figs.S29-S32.

**B. Measured variability across aster formations.** The active matter systems we investigate are intrinsically non-ergodic, stochastic systems sensitive to initial and ambient conditions, both controlled (temperature, motor concentrations, etc.) and uncontrolled (thermodynamic noise, laser stability, etc.). These factors mean there is fundamental variability in the trajectories of aster geometry accompanying gradients and measured dissipation over time. Thus, even when experimental reactions occur under the same biochemical conditions (concentrations of motors, microtubules, and ATP), this real variability persists. Here, we give a sense of the type and extent of variability in replicate aster formation trajectories.

As a focused glimpse of the scales of variability, Figure S28 compares three asters. (These asters are highlighted here to reflect both consensus in major features and variability in numerical features of interest.) The first row (A) illustrates the aster data referred to most centrally in the main text; the second row (B) showing a technical replicate of a separate aster (with optical excitation and imaging in the same flow channel in the same reaction mix but at a different position in the experimental flow cell); and the third row showing a more divergent "biological" replicate (another aster with a separately made reaction mix occurring on a different day).

Many replicate asters tend to show a fairly coherent behavior with each other. For instance, the aster from the main text (shown in Figure S28(row A)) shows commonalities with many other aster trajectories (like another aster represented in

Figure S28(row B)). This coherence includes shared features like the scale of developing gradients, the timescale of energy consumption, and the monotonically decreasing power of asters over time. Yet, variability does manifest in some other asters, like the especially variable aster example given in Figure S28(row C). Notably, some of these asters appear to show nonmonotonic power consumption over time. Comparisons of aster motor profiles and ATP profiles across a much wider selection of asters are given in Figures S29 and S30, along with their manifested concentration gradients and power consumptions over time in Figures S31 and S32, respectively.

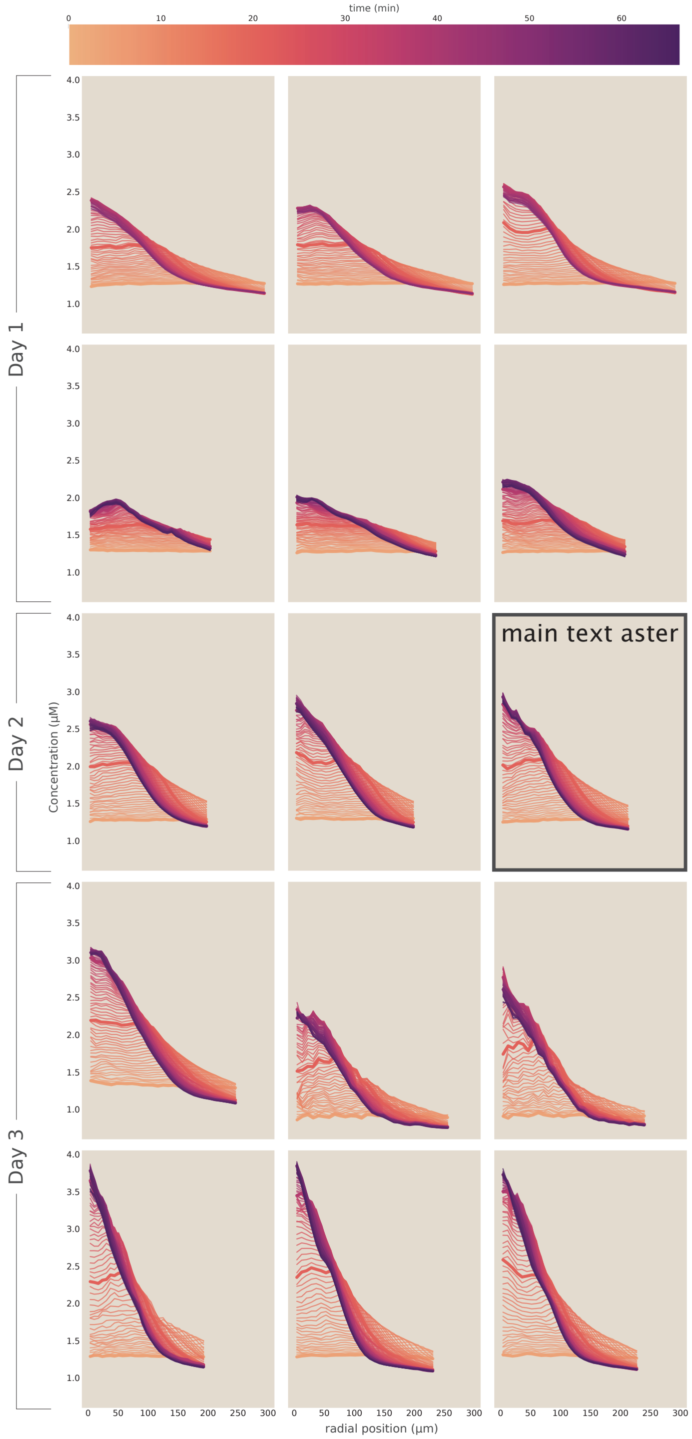

**Fig. S29.** Variation of motor concentration profiles across experiments, performed on common or distinct days in common or distinct experimental flow cells. Each inset plot is a separate aster experiment; the ATP, concentration gradient, and power trajectories for each experiment are shown in analogous inset plot locations in the grids of Fig. S30, Fig. S31, and Fig. S32.

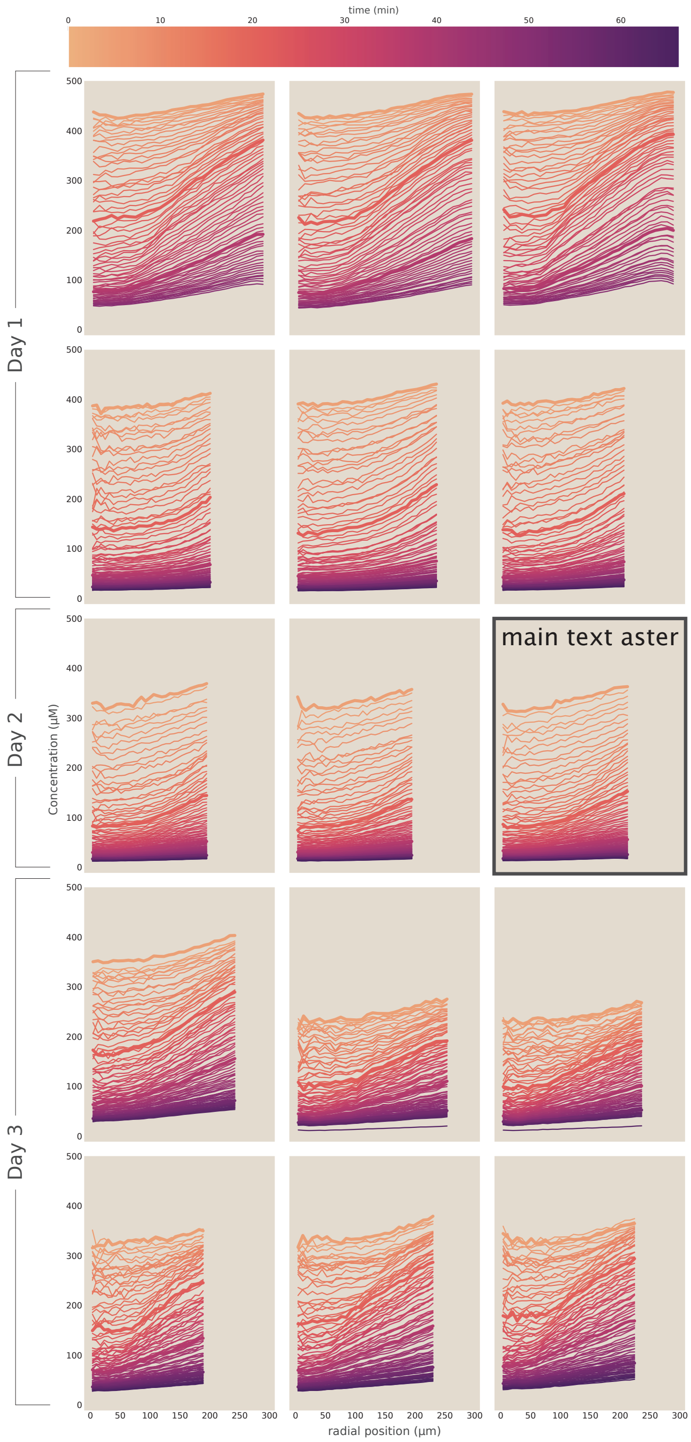

**Fig. S30.** Variation of ATP profiles across aster experiments, performed on common or distinct days in common or distinct experimental flow cells. Each inset plot is a separate aster experiment; the motor, ATP concentration gradient, and power trajectories for each experiment are shown in analogous inset plot locations in the grids of Fig. S29, Fig. S31, and Fig. S32.

545 ### *B.1. Empirical scale of variability in spatial gradients.*

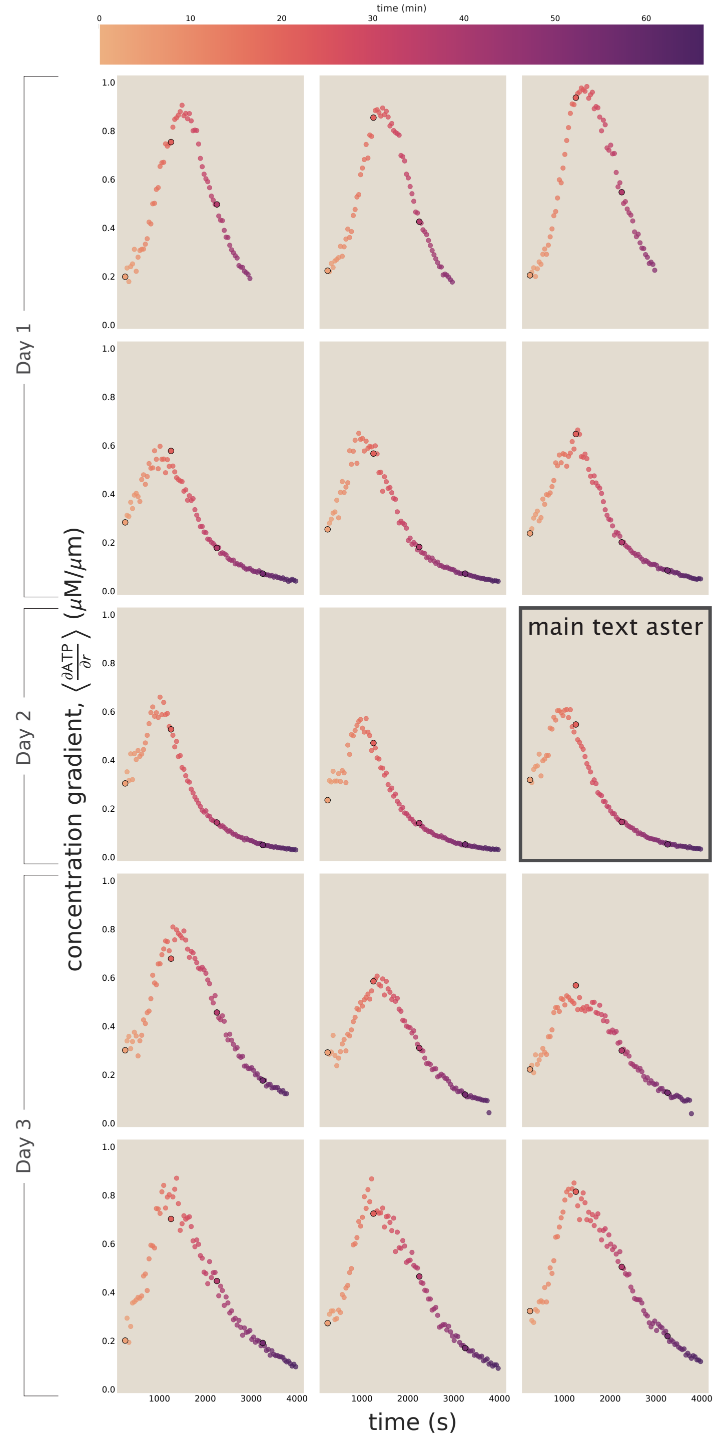

**Fig. S31.** Variation of emergent concentration gradients in ATP across aster experiments, performed on common or distinct days in common or distinct experimental flow cells. Specifically, the slopes visible in the concentration profiles of Fig. S30 are quantitatively approximated as the ATP gradient over the $r = 100\ \mu$m to $200\ \mu$m region, approximately spanning the edge to the middle third of most asters, calculated as the (unsmoothed, raw) first order finite difference $\left\langle \frac{\partial \mathrm{ATP}}{\partial r} \right\rangle \approx \frac{\mathrm{ATP}(r=100\mu\mathrm{m},t) - \mathrm{ATP}(r=200\mu\mathrm{m},t)}{100\mu\mathrm{m}}$. (Datapoint circles with black edges correspond to the same timepoints highlighted in bold lines in this larger suite of Figs. S28 - S32.) Each inset plot is a separate aster experiment; the motor, ATP, and power trajectories for each experiment are shown in analogous inset plot locations in the grids of Fig. S29, Fig. S30, Fig. S32.

Figure S31 illustrates how variably aster experiments develop concentration gradients in ATP. Specifically, we quantitatively summarize ÅTP gradients by estimating their magnitudes in an outer radial region near aster edges, as $\left\langle \frac{\partial \mathrm{ATP}}{\partial r} \right\rangle \approx \frac{\mathrm{ATP}(r=100\mu\mathrm{m},t) - \mathrm{ATP}(r=200\mu\mathrm{m},t)}{100\mu\mathrm{m}}$, and plot these over time for every aster. Every aster's steepest gradient develops at an intermediate time (surprisingly coherently, within a few hundred seconds or less of a typical apotheosis time $t_* \sim 1.5 \times 10^3$ seconds). As shown in the Fig. S31, these steepest gradients vary between $\approx 0.5\ \mu$M/$\mu$m (the scale conservatively quoted in the main text) and up to $\approx 1\ \mu$M/$\mu$m across asters.

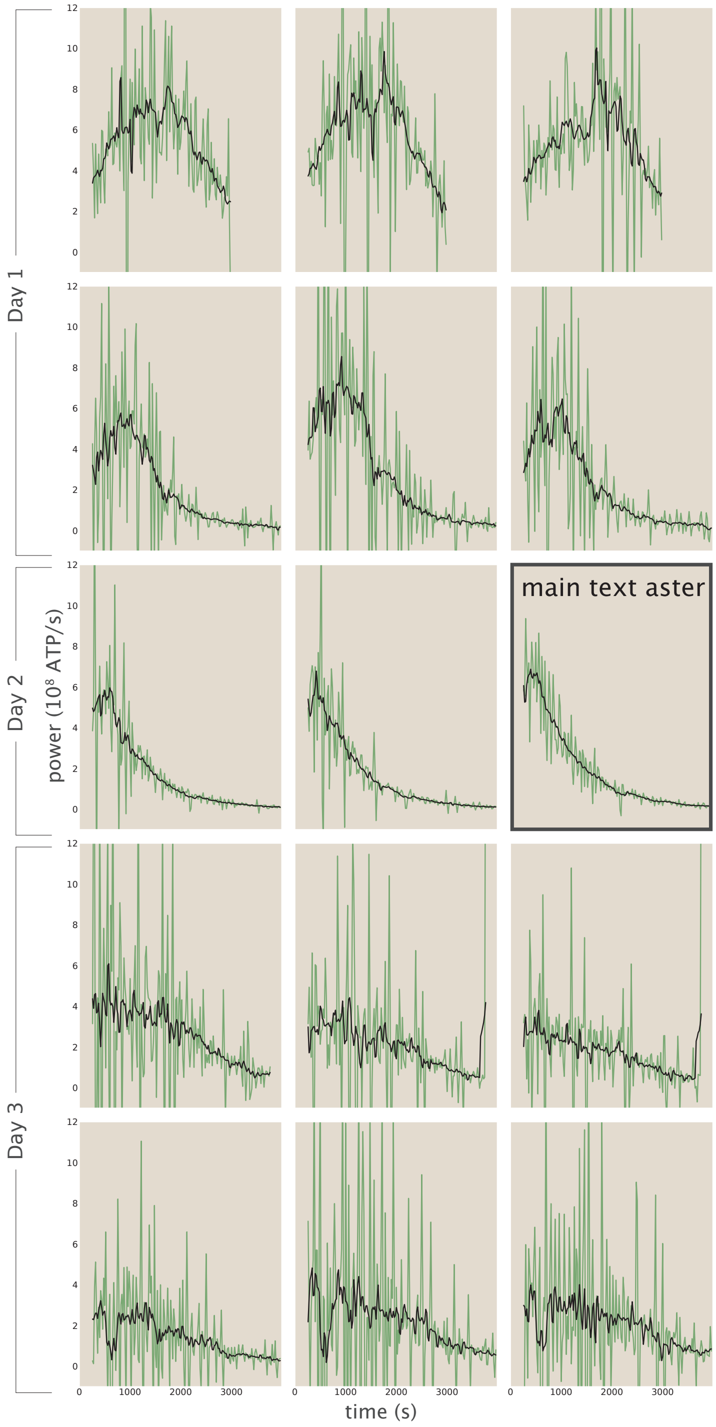

**Fig. S32.** Variation of power dissipation trajectories across aster experiments, performed on common or distinct days in common or distinct experimental flow cells. Green lines show raw first-order finite differences between the ATP extant at consecutive instants in time. Since such numerical derivatives are notoriously noisy, as a guide to the eye, the black lines display a short moving average of these trajectories. Each inset plot is a separate aster experiment; the motor, ATP, and concentration gradient trajectories for each experiment are shown in analogous inset plot locations in the grids of Fig. S29, Fig. S31, and Fig. S30.

***B.2. Empirical scale of variability in power dissipation.*** Being based on numerical derivatives of noisy temporal data, power estimates made in this study are perhaps the components most susceptible to intrinsic and extrinsic stochasticity in our analysis. We refrained from performing sophisticated or bespoke signal processing to estimate temporal gradients, in favor of taking finite differences between adjacent time intervals for simplicity and transparency. What is clearly resolvable despite this straightforward approach, is the typical scale of power consumption rates and their typical tendencies over time for each aster. Specifically, as reported by Figure S32, initial power consumptions vary between $\approx 2\times 10^8$ ATP/s and $\approx 6\times 10^8$ ATP/s across individual asters, with peak dissipations varying from $\approx 3\times 10^8$ ATP/s to $\approx 8\times 10^8$ ATP/s. (So, these data unanimously affirm a typical scale of oder few $\times\, 10^8$ ATP/s.) Twelve of the fifteen asters decrease their power consumptions apparently monotonically in time, whereas three of the fifteen aster formations display power that first appears to increase, reach a peak, then decrease.

#### 4. Continuum field equations for ATP hydrolysis

To reckon quantitatively with the experimentally measured phenomenology of ATP dissipation reported by our measurements, we investigated a slew of different theoretical pictures for how ATP changes in time. Examining the ATP concentration in a small region of the system, several mechanisms can cause such changes in ATP concentration. Diffusion, with coefficient $D$, and fluid flow, $v(r,t)$, drive molecular transport into or out of the region. ATP can also be consumed by motor proteins, $m(r,t)$, as a function, $h(\mathrm{ATP}(r,t), m(r,t), \mathrm{tub}(r,t); r, t)$, which depends on the concentrations of ATP, motor proteins, and microtubules, $\mathrm{tub}(r,t)$. ATP could be created by ATP synthase in a cell or by enzymatic regeneration systems *in vitro*; however, we do not include any method for regeneration in our system, so we do not consider creation here. All together, the change of ATP over time can be described by a reaction-diffusion-advection equation,

$$\frac{\partial \mathrm{ATP}}{\partial t} = D\nabla^2 \mathrm{ATP} - h\left(\mathrm{ATP}(r,t), m(r,t), \mathrm{tub}(r,t); r, t\right) - \mathbf{v}(r,t)\cdot\nabla \mathrm{ATP}, \qquad [26]$$

which assumes isotropic homogeneous diffusion and incompressible fluid flow.

To build understanding of the time scale of ATP depletion and the steepness of developing gradients, our analysis deeply explored a myriad of possible candidate dependencies of the hydrolysis rate $h\left(\mathrm{ATP}(r,t), m(r,t), \mathrm{tub}(r,t); r, t\right)$ on the underlying governing fields. (For the narrow scope of this work investigating gradients, this analysis largely neglected uniformly contractile fluid flow profiles (effectively taking $\mathbf{v}(r,t)\to 0$), though the character of this manifest contraction is clearly biophysically interesting, as inquired by work in our laboratories and others (2).) Specifically, using finite element modeling reproducing our three dimensional experimental geometry and boundary conditions, we thoroughly investigated how successfully models at increasing levels of sophistication could (or could not) recapitulate the important phenomenological features of our experiments.

Our models acknowledge the stoichiometric inhibition of the motors' hydrolysis of ATP by accumulating ADP. As demonstrated in Figure S33, we find that including ADP dependence in the hydrolysis term slows the global depletion time of ATP throughout the system. Specifically, we found that the hydrolysis rate is better modeled by including the proportionality term,

$$p_{\text{mot-ATP}}(\mathrm{ATP}(r,t)) \equiv \frac{\frac{\mathrm{ATP}}{K_T}}{1+\frac{\mathrm{ATP}}{K_T}+\frac{\mathrm{ADP}}{K_D}}, \qquad [27]$$

where $K_\mathrm{T}$ and $K_\mathrm{D}$ are the Michaelis-Menten constants of ATP and ADP binding to the motor protein respectively, rather than a more naive, simpler form like

$$p_{\text{mot-ATP}}(\mathrm{ATP}(r,t)) \equiv \frac{\frac{\mathrm{ATP}}{K_T}}{1+\frac{\mathrm{ATP}}{K_T}}. \qquad [28]$$

This picture is consistent with the biochemical knowledge established elsewhere by foundational literature (10). We did a similar test for the accumulation of phosphate and found that based on biochemical literature values of $K_\mathrm{P}$, the Michaelis-Menten term of phosphate to the motor protein, this term was negligible.

Acknowledging this dependence on ADP, we investigated a model where the local hydrolysis rate is defined as the product of $\gamma$, the average per-motor ATP hydrolysis rate, $m$, the local motor concentration, and $p_{\text{mot-ATP}}$, the probability an ATP binds to a motor as described above,

$$h(r,t,\mathrm{ATP}) \equiv \gamma\; m(r,t)\; p_{\text{mot-ATP}}(\mathrm{ATP}(r,t), \mathrm{ADP}(r,t)). \qquad [29]$$

In Figure S34 we explore how well this model reflects the phenomenology of our experiment as we tune the diffusion and hydrolysis rate parameters. Simulations of this model could recapitulate the transient appearance of meaningful gradients in ATP, but *only* when parameterized by unphysically small effective values of the diffusion coefficient $D$. The timescale to deplete ATP in experiments most closely matches the simulations when the average motor hydrolysis rate is $0.5\,\mathrm{s}^{-1}$.

Aiming to visualize gradients for more physical diffusion constants, we attempted a subsequent minimal model, whose form we discuss in the main text. This further model acknowledges that the rate at which motors hydrolyze should depend not only on local motor concentration, but the proportion of these motors actually bound and stepping along microtubules. That is, the hydrolysis rate adopts the form,

$$h(r,t,\mathrm{ATP}) \equiv \gamma\; m(r,t)\; p_{\text{mot-ATP}}(\mathrm{ATP}(r,t))\; p_{\text{mot-tub}}\left(\mathrm{tub}(r,t), m(r,t)\right) \qquad [30]$$

where

$$p_{\text{mot-tub}} \equiv \frac{(m(r,t)+\mathrm{tub}(r,t)+K_{d_{\text{mot}\cdot\text{mt}}}) - \sqrt{(m(r,t)+\mathrm{tub}(r,t)+K_{d_{\text{mot}\cdot\text{mt}}})^2 - 4\,m(r,t)\,\mathrm{tub}(r,t)}}{2\,m(r,t)}, \qquad [31]$$

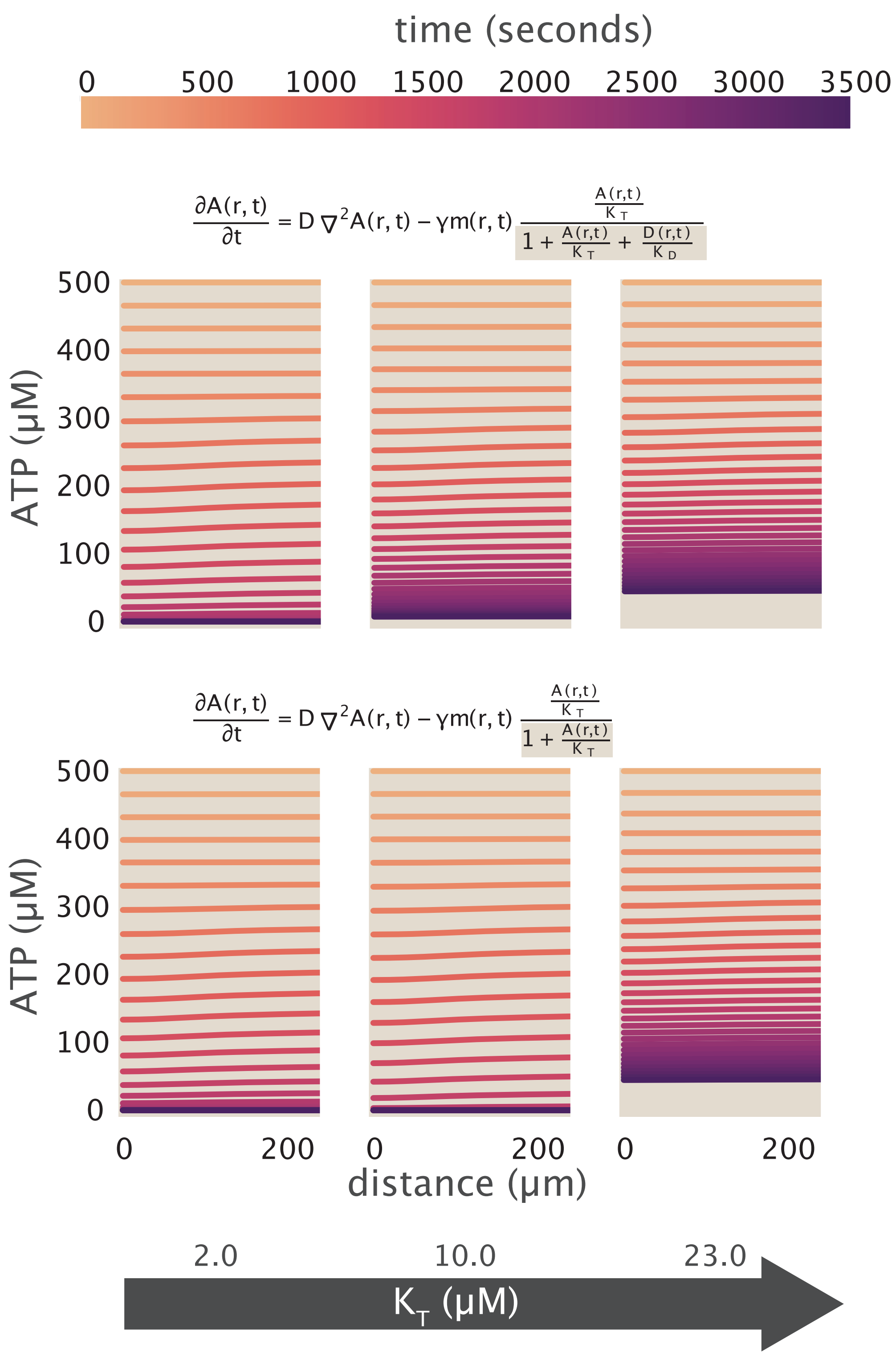

**Fig. S33. Including ADP dependence in our model of ATP binding to motor proteins slows the time it takes for the system to run out of ATP.** In the top row, we include competitive inhibition of ADP binding to the motor protein rather than ATP, while in the second row we ignore any effects of ADP in solution. ATP depletes slower in the top row, where we account for ADP, than the second row. We also show the effect of increasing the Michaelis-Menten constant for ATP binding to motor proteins, $K_T$, across the columns.

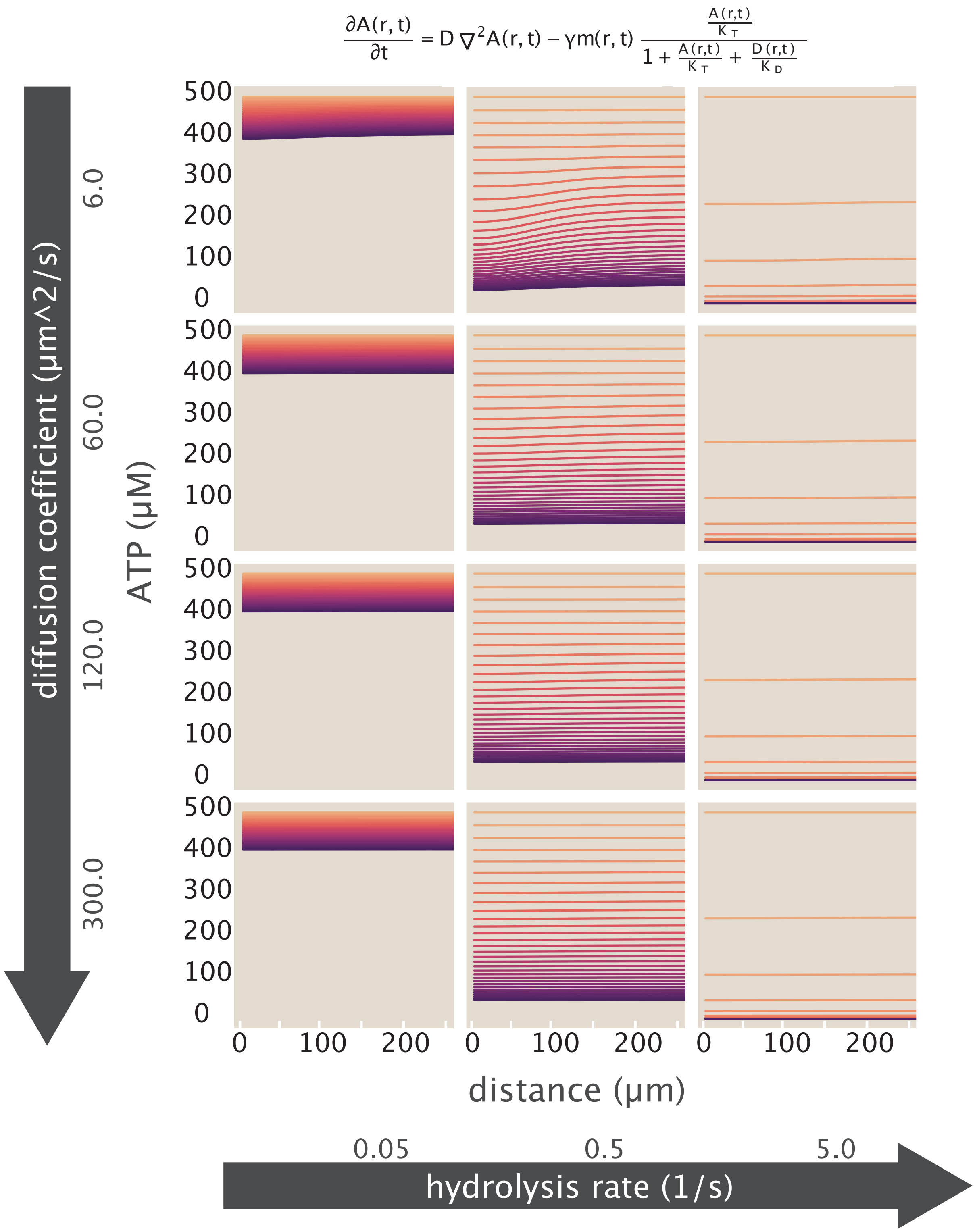

**Fig. S34. Tuning the diffusion constant very low can produce a gradient in ATP, while tuning the hydrolysis rate modulates how quickly ATP is depleted.** We show the results of sweeping the diffusion constant in the rows. We only see noticeable gradients when $D = 6\,\mu\mathrm{m}^2/\mathrm{s}$. The results of sweeping the hydrolysis rate is showed in the columns. As the hydrolysis rate increases, the time to deplete ATP becomes shorter.

is the probability that a motor is bound to a microtubule (as derived in section 5). Simulations of this model can recapitulate gradients in ATP, using parameter values that are largely physically plausible (at least, falling within factors of a few of putative (and very poorly constrained) values of diffusion coefficients $D$, $K_M$, $K_T$, $\gamma$, local tubulin concentrations, etc.) We provide a systematic sensitivity analysis of how this feasible model behaves in Figure S35. Relative to the earlier model, this additional dependence on microtubule concentration achieves the important effect of sharpening the spatial variation in the hydrolysis rate $h$ itself.

We stress that even such an elaborate model is certainly not to be regarded as the final theoretical word on these rich and intricate dynamics, which surely invite much further theoretical thinking. Instead they are intended as tools for exploring hypotheses and for conceptual imagination. They give two main takeaways. First, gradients in ATP can develop under our experimental conditions using acceptably physically plausible reaction-diffusion dynamics and parameter values. Second, such models succeed at producing gradients in ATP concentrations because these dynamics sufficiently strongly localize reactions in regions of space within typical times that diffusion would otherwise smooth ATP. (This fact induces to a broader conceptual principle, as we articulate in a simple analytical bound, see section 13.)

In the subsections that follow, we describe and document the operational procedures we used to perform our extensive simulations of the above models and reach these conclusions.

#### A. Finite Element Simulation Parameter Choices.

***A.1. ATP diffusion constant.*** We can compute an expected diffusion constant for ATP in our system using the Stokes-Einstein equation,

$$D = \frac{k_{\mathrm{B}}T}{6\pi\mu r}, \tag{32}$$

where $\mu$ is the viscosity of the media and $r$ is the radius of the molecule, given that our system is at low Reynold's number. Using the viscosity of our system ($\mu = 2.4 \times 10^{-3}\,\mathrm{N\cdot s/m^2}$ (11)), the diffusion constant of a species only depends on its radius,

$$D = \frac{k_{\mathrm{B}}T}{6\pi\mu r} = \frac{4\,\mathrm{pN\cdot nm}}{6\pi r(2.4\times 10^{-3}\,\mathrm{pN\cdot s}/\mu\mathrm{m}^2)} \approx \frac{90\ \mathrm{nm}}{r}\,\frac{\mu\mathrm{m}^2}{\mathrm{s}}, \tag{33}$$

given that $k_{\mathrm{B}}T \approx 4\,\mathrm{pN\cdot nm}$. We can now compute the relevant diffusion constants for each molecule in the experiment: ATP has a radius of 0.7 nm, (BNID 106798), which leads to a diffusion coefficient of

$$D_{\mathrm{ATP}} \approx 130\,\mu\mathrm{m}^2/\mathrm{s}. \tag{34}$$

This well matches literature values. ATP in cytosol has been measured to have a diffusion constant of $250 - 400\,\mu\mathrm{m}^2/\mathrm{s}$. Since our system is more viscous due to a significant fraction of glycerol (by a factor of 2.4 if we take the viscosity of cytosol to that of water), appropriately scaling the diffusion reduces the diffusion constant to $\approx 100 - 170\,\mu\mathrm{m}^2/\mathrm{s}$.

***A.2. Hydrolysis rate.*** Literature hydrolysis rates for single molecule experiments of NCD motor proteins range from 2 ATP/s to 25 ATP/s (12–18). We performed gliding assays using our motor proteins and measured the speed of microtubule gliding across various ATP concentrations as shown in Figure S36. We fit the measured motor speeds to the equation

$$S = d\gamma\frac{[\mathrm{ATP}]/K_{\mathrm{T}}}{1+\mathrm{ATP}]/K_{\mathrm{T}}}, \tag{35}$$

where $d$ is the length of a motor step (8 nm), $\gamma$ is the per motor ATP hydrolysis rate, and $K_{\mathrm{T}}$ is the Michaelis-Menten constant for ATP binding to the motor protein. The black line in Figure S36 is characterized by the fit parameters $\gamma = 6\ \mathrm{s}^{-1}$ (within the range of literature values) and $K_{\mathrm{T}} = 31\,\mu\mathrm{M}$.

We additionally have performed standardized bulk ATPase plate reader measurements. Using the Enzyme Linked Inorganic Phosphate Assay from Cytoskeleton, we found the bulk hydrolysis rate (averaging all motor proteins in the assay irrespective of their binding state to microtubules) was 0.5 ATP/s.. Thus, considering this measurement and our gliding measurements, when running FEM simulations, we take the hydrolysis rate of ATP by motor proteins to be of order a few.

***A.3. Michaelis-Menten and dissociation constants.*** We choose the Michaelis-Menten and dissociation constants in our simulation based on reported literature values. For $K_{\mathrm{T}}$, the Michaelis-Menten constant of ATP binding to motor proteins, we take $23\,\mu\mathrm{M}$ as reported by (19). For $K_{\mathrm{D}}$, the Michaelis-Menten constant for ADP binding to motor proteins, we use $34\,\mu\mathrm{M}$, as given by (10), which studies conventional kinesin, but we take this value as an estimate of the correct order of magnitude. We use $K_{\mathrm{d_{mot\cdot mt}}} = 60\,\mu\mathrm{M}$ for the dissociation constant of motors binding to microtubules, as previously reported by our labs (16).

***A.4. Motor and tubulin concentration profiles.*** When inputting concentration profiles into our simulations, our goal is to represent the characteristic features of the species' dynamics, not to perfectly fit our data. With this in mind, we build trial functions based on simple expressions that appropriately capture the magnitude and location of greatest concentration, the approximate length scale of gradients, and the infinite boundary concentration conditions.

For the motor concentration profile function, $m(r,t)$, we use a sigmoidal based trial function,

$$m(r,t) = \frac{m_{\max}(t)}{1+\exp\left(\frac{r-r^*(t)}{\lambda(t)}\right)} + m_{\min}(t), \tag{36}$$

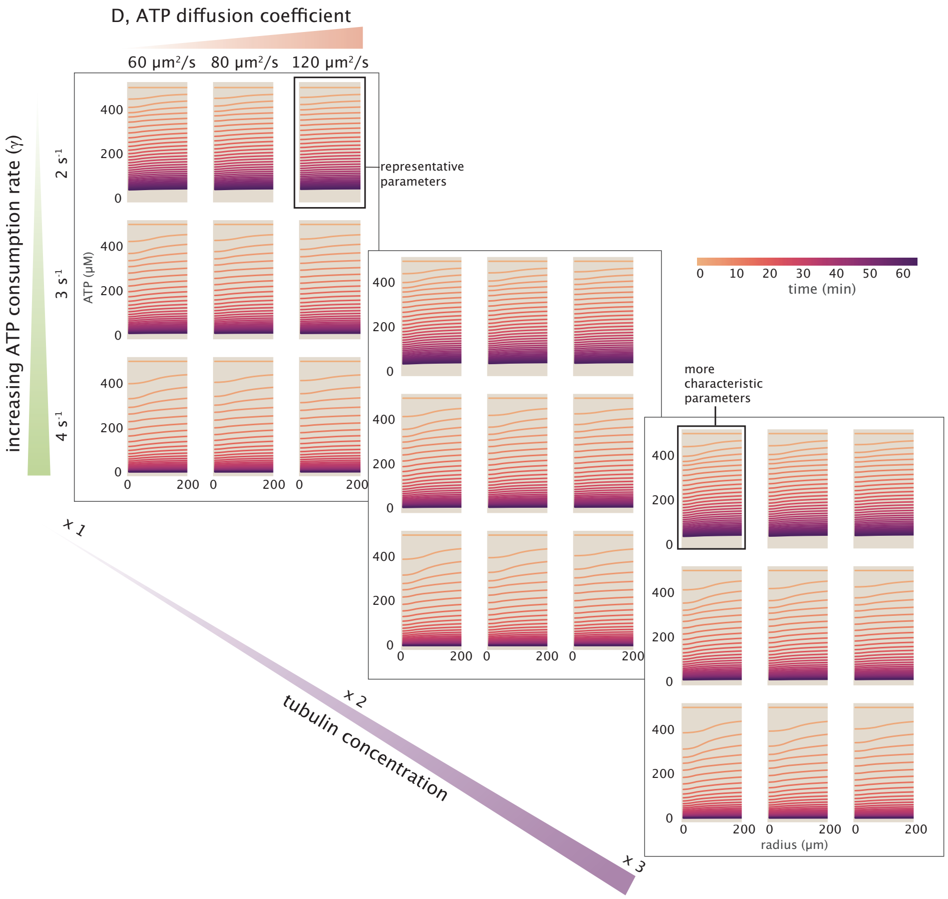

**Fig. S35. Including the binding of motor proteins to microtubules, we show how modulating diffusion, motor hydrolysis rate, and microtubule concentration impacts the phenomenology of ATP depletion.** We find that decreasing the diffusion constant and increasing the microtubule concentration creates larger gradients in ATP. Increasing the hydrolysis rate causes faster global depletion of ATP.

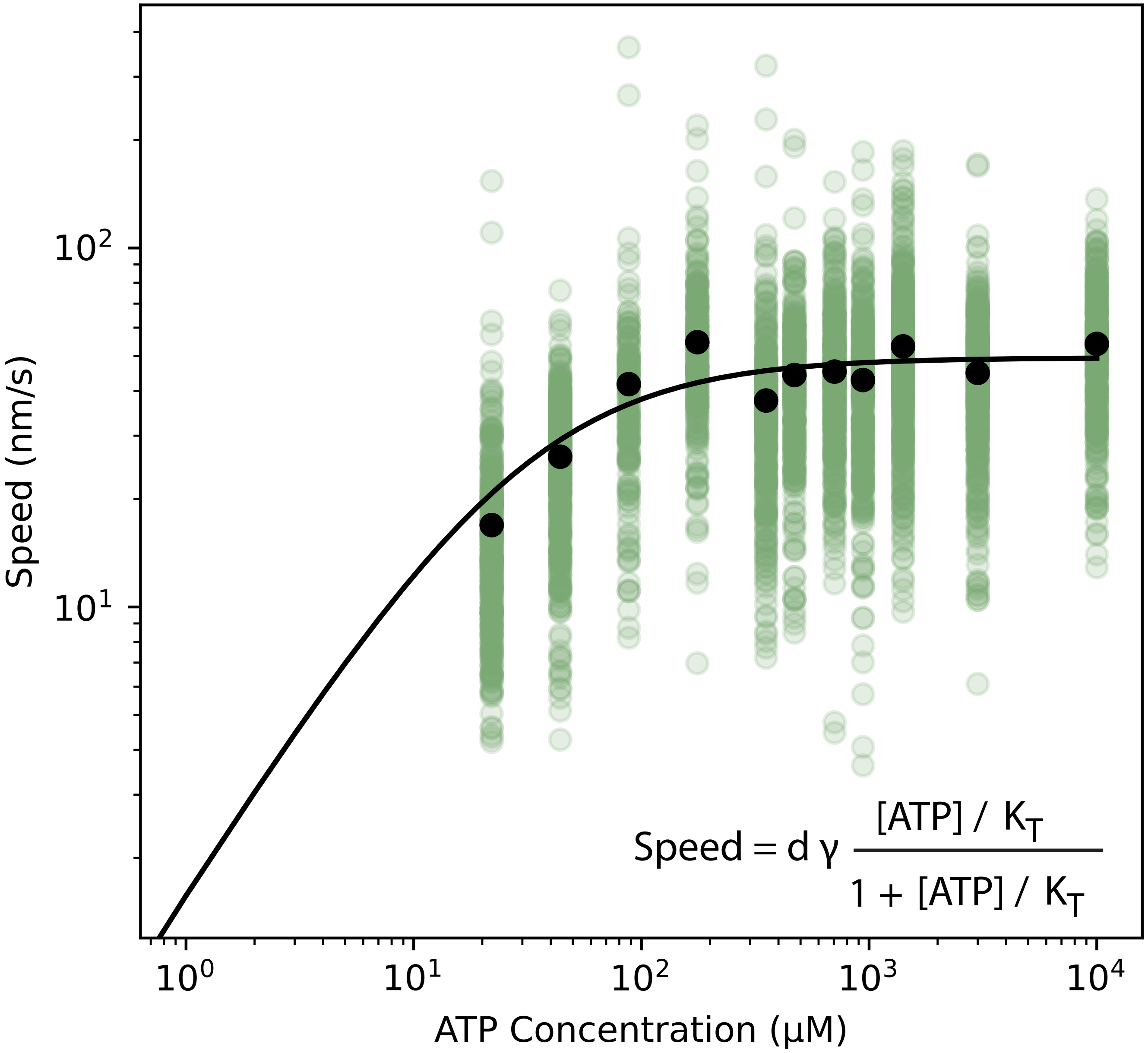

**Fig. S36. Speeds of NCD motor proteins at various ATP concentrations as measured by gliding assays.** Green dots indicate speeds of individualy tracked microtubules in a gliding assay. The black line fits the data to Equation 35, where the hydrolysis rate $\gamma = 6$ s$^{-1}$ and the Michaelis-Menten constant $K_{\mathrm{M}} = 31\,\mu\mathrm{T}$.

where $m_{\max}(t)$ is the time-dependent value of the maximum motor concentration, $r^*(t)$ is the time-dependent radial position of the maximum motor concentration, $\lambda(t)$ is the time-dependent exponential decay constant, and $m_{\min}(t)$ is the time-dependent minimum motor concentration, which is the concentration in the limit the radius goes to infinity. In Figure S37, we graphically depict equation 36 indicating empirically how the parameters depend on time. We fit data for each time-dependent parameter to simple functions written in each subplot and represented by the orange line in Figure S37.

Microtubules are made of tubulin. To reconstitute microtubules, we polymerize a known concentration of tubulin and determine microtubule concentration with the knowledge that there are 1600 tubulin per $1\,\mu$m microtubules (20). Our labs found our microtubules are approximately $1\,\mu$m in length as described in previous works (2). However, when accounting for the binding of motors to tubulin in our simulations, we write an equation to represent the filaments in terms of tubulin rather than microtubules. Additionally, the true units of this trial function is concentration of tubulin binding sites where we scale experimental tubulin concentrations by a factor of 13, the number of binding sites per tubulin.

For the tubulin profile, we stitch together two gaussian functions that meet at the maximum tubulin peak. To make an analytic trial function, we multiply gaussian functions by a sigmoid with a very small length scale (we find $0.01\,\mu$m is sufficient) to approximate a step function centered about the tubulin peak location. Thus, we represent the tubulin profile as,

$$\begin{aligned}\text{tub}(r,t) = {} & \underbrace{\frac{1}{1+\exp\left(\frac{r-r^*(t)}{0.01\,\mu\text{m}}\right)}}_{\text{left step function}} \underbrace{\left((\text{tub}_{\max}(t)-\text{tub}_0(t))\exp\left(-\frac{(r-r^*(t))^2}{\sigma_{\text{left}}}\right)+\text{tub}_0(t)\right)}_{\text{left sigmoid function}} \\ & + \underbrace{\frac{1}{1+\exp\left(-\frac{r-r^*(t)}{0.01\,\mu\text{m}}\right)}}_{\text{right step function}} \underbrace{\left((\text{tub}_{\max}(t)-\text{tub}_\infty(t))\exp\left(-\frac{(r-r^*(t))^2}{\sigma_{\text{right}}(t)}\right)+\text{tub}_\infty(t)\right)}_{\text{right sigmoid function}},\end{aligned} \qquad [37]$$

where $r^*(t)$ is the time-dependent radial position of the maximum tubulin concentration, $\text{tub}_\text{m}\text{ax}(t)$ is the time-dependent value of the maximum tubulin concentration, $\text{tub}_0(t)$ is the time-dependent value of the tubulin concentration at $r=0$, $\sigma_{\text{left}}(t)$ is the time-dependent gaussian length scale for the left gaussian, $\text{tub}_\infty(t)$ is the time-dependent value of the tubulin concentration at $r=\infty$, and $\sigma_{\text{right}}(t)$ is the time-dependent gaussian length scale for the right gaussian. We fit each of these expressions to our empirical data as represented by the orange curves in Figure S38.

Both the motor and tubulin trial functions can be compared to the experimentally measured concentration profiles in Figure S40.

**B. Finite Element Simulations of Continuum Models.** Our work is predicated on an interplay between experimental measurements and a theoretical description of our results. As seen in the main body of the paper, we have considered an elementary reaction-diffusion model that describes the spatiotemporal evolution of ATP as a result of the consumption of ATP by the motors as well as the diffusion of ATP.

Although the equations are simply stated, for the case in which the motor distribution is varying in space and time, the solution of these equations is prohibitive (i.e. to the best of our knowledge, analytically out of reach) except in the most highly idealized circumstances. As a result, we were interested in having a robust numerical approach to solving the continuum equations describing our experiments, including the effects of photobleaching, in the context of the complex geometry of our microscope coverslips and optically activated aster regions.

To that end, in several different contexts we used the finite element method to carry out numerical solutions of our reaction-diffusion-photobleach equations. Though the equations of mathematical physics can often be solved in highly-idealized geometries using special functions, once the symmetry of those geometries is gone, analytical solutions are often no longer available or helpful. The finite element method provides a particularly simple alternative. The geometry of interest is discretized into a set of points known as the nodes. The field quantities of interest are defined on each of those nodes, for example, $c_n$ refers to the concentration of the field of interest at the $n^{th}$ node. To find the value of the field variables at positions other than the nodes, the finite element interpolates using "shape functions." For example, if we are to solve the one-dimensional diffusion equation, then we would write the concentration in the form

$$c(x) = \sum_{n=1}^{N} c_n N_n(x), \qquad [38]$$

where $N_n(x)$ is the shape function which in the simplest version of the approach is nothing more than linear interpolation! The key point is that once we adopt a finite-element discretizations, instead of solving for an entire unknown function $c(x)$, we solve for the finite set of values $(c_1, c_2, c_3, \cdots c_N)$.

Part of the power of the finite element method is that it allows us the flexibility to consider arbitrary geometries and to quickly move between different effects by tuning the relevant partial differential equations. In particular, we have used the commercial finite element software known as COMSOL. As seen in Figure S39, to carry out a simulation of interest, we need to set up the calculations by describing the geometry, choosing which partial differential equations to solve, what the boundary and initial conditions are and how to mesh our region of interest. For those interested, we will provide the exact COMSOL simulation setup files corresonding to our various numerical studies.

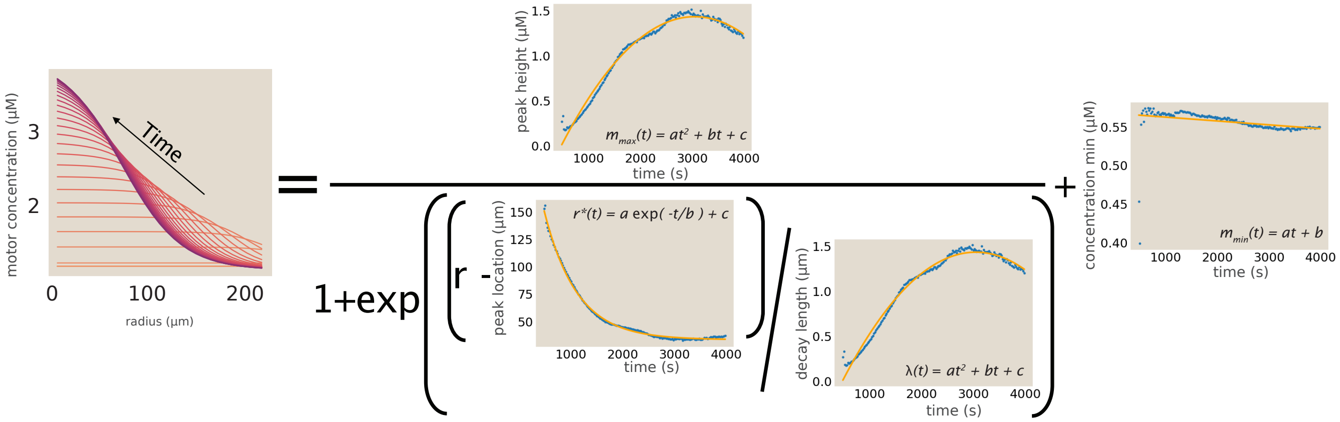

**Fig. S37. Graphical depiction of the motor trial function.** Empirical data for the characteristic parameters of the motor trial function are indicated with blue dots. The orange lines depict the expression we fit to the data. Each subplot indicates the expression we use and the coefficients $a,\ b,$ and $c$ are the fit parameters.

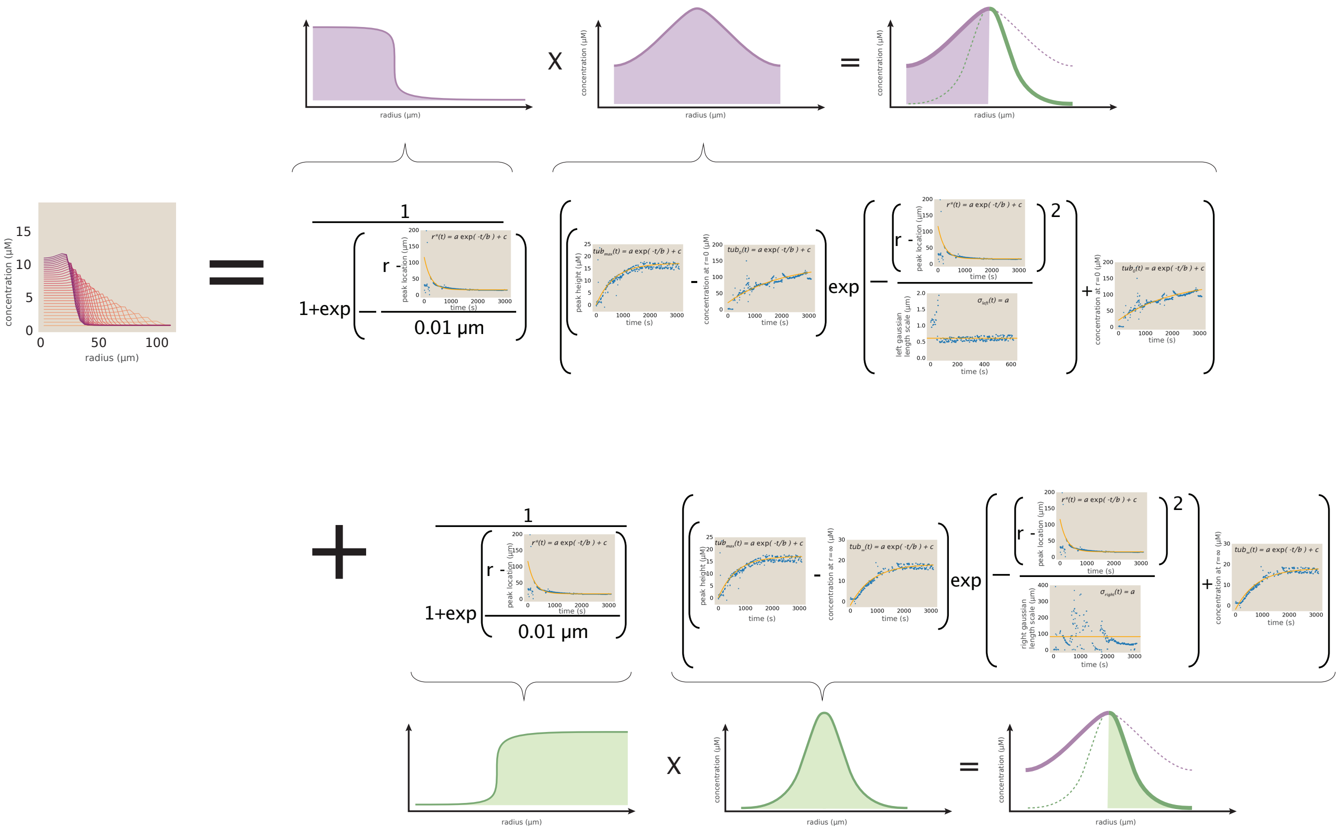

**Fig. S38. Graphical depiction of the tubulin trial function.** Empirical data for the characteristic parameters of the tubulin trial function are indicated with blue dots. The orange lines depict the expression we fit to the data. Each subplot indicates the expression we use and the coefficients $a,\ b,$ and $c$ are the fit parameters.

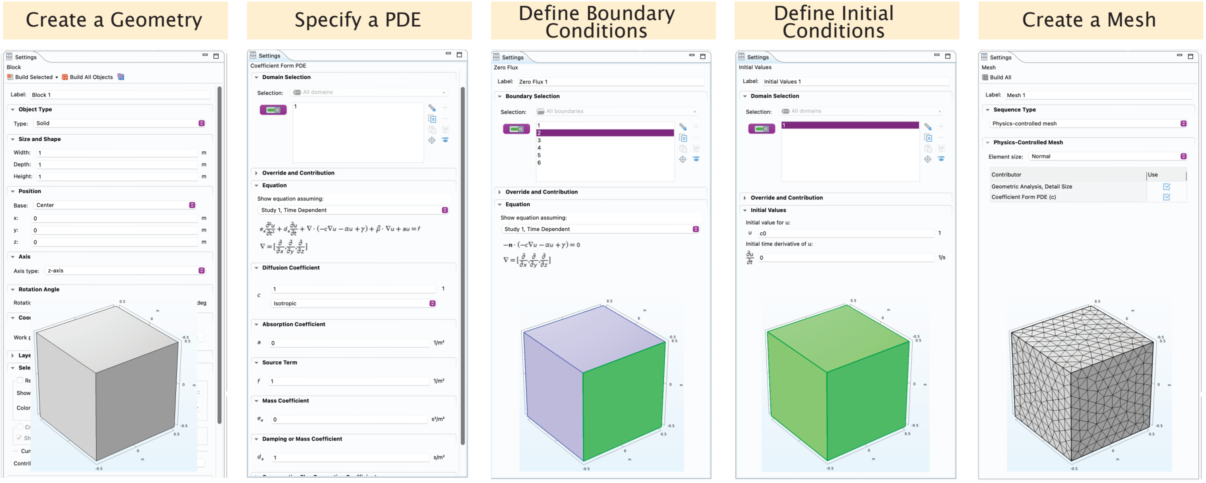

**Fig. S39. Setting up a finite element calculation in COMSOL.** The use of COMSOL is dependent upon a series of steps that in total make it possible to solve some partial differential equation of interest. The five columns show how to create a geometry, specify which partial differential equation we are solving, how to define boundary conditions and initial conditions and how to set up the finite element mesh.

In the main body of the paper, we showed how the finite element method was used to compute the depletion over time of ATP as a result of a gradient in the distribution of motors. Later in the SI, we will discuss how we used the finite element method to make careful simulations of how photobleaching might have influenced our measurement of the ATP concentration.

#### 5. Binding dynamics of motor proteins to microtubules

One of the most important theoretical questions broached in our paper is whether the competing dynamics of ATP diffusion and consumption could give rise to the kinds of observed gradients we measure. To that end, we undertook a series of numerical studies using the finite element method to explore the distribution of ATP in space and time during the development of an aster. As seen in Figure S40, we can plot the distribution of microtubules (left) and motors (right) as a function of time. Further, as indicated in the figure, we can use our understanding of motor-microtubule binding to infer the fraction of bound motors as a function of position within the aster at various times. The reason such a calculation is important is that it reflects an underlying difference in the rate of ATP consumption at various points within the aster.

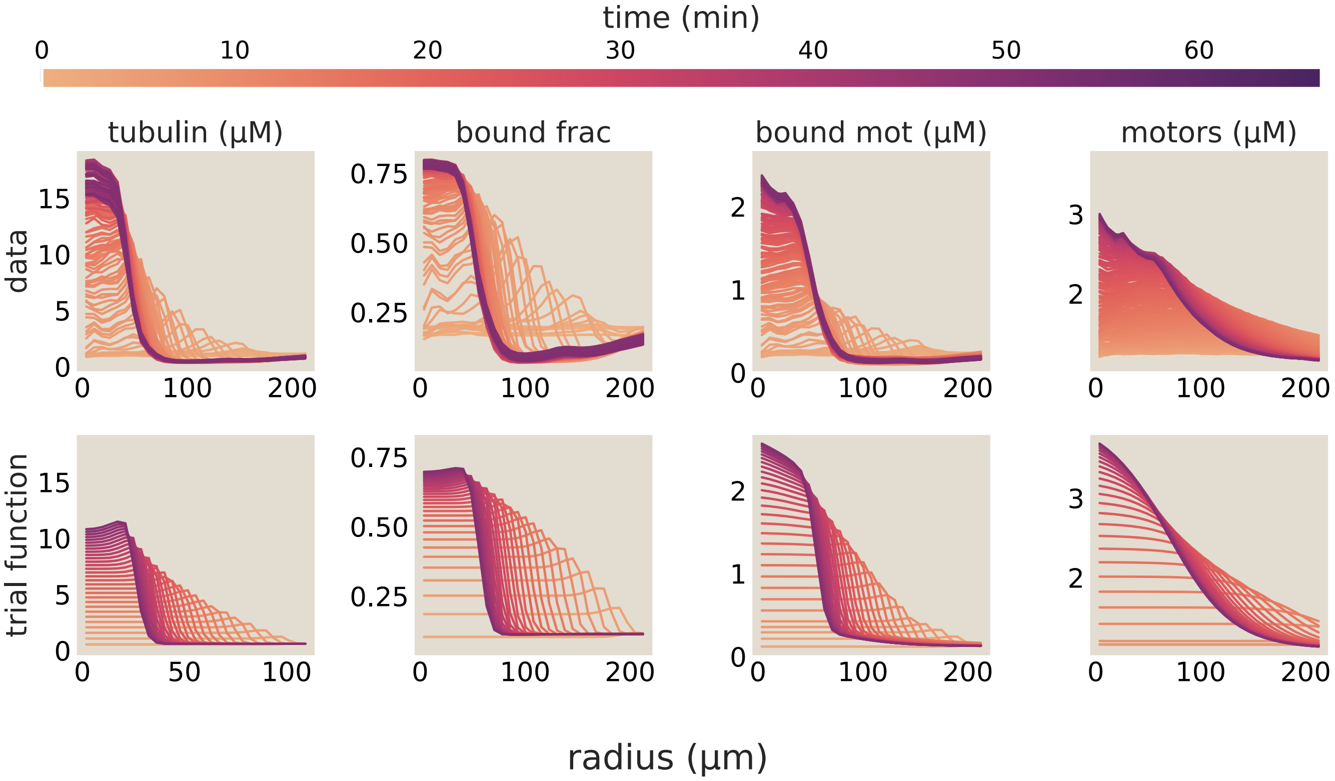

**Fig. S40. Concentration profiles for motors and microtubules.** The distribution of microtubules and motors as a function of distance from the center of the aster as a function of time show the development of a gradient over time. The bottom row shows the empirical and approximate fits used as input into the finite element calculations. The key point of the motor and microtubule temporal distributions is that they are used as input into the consumption-diffusion model for the ATP.

To make progress on the FEM analysis of the ATP consumption in space and time, our reaction-diffusion framework needs as input the current distribution of microtubules and motors. In particular, the spatial distribution of ATP consumption depends upon the fraction of bound motors. Where the concentration of both motors and microtubules are both "low" there will be relatively few motors stepping per unit time and hence a low allied ATP consumption rate. On the other hand, as seen in the figure, near the aster center where both motors and microtubules are at "high" concentration, the ATP consumption rate will be high.

To make these notions of "low" and "high" concentrations precise, we need to solve the statistical mechanical binding problem. We model motor binding to microtubules as a ligand-receptor binding reaction, where $[L]_{\text{tot}}$ is the total concentration of motor proteins, $[R]_{\text{tot}}$ is the total concentration of available microtubule binding sites, and $[C]$ is the concentration of bound motor-microtubule complexes. The corresponding free concentrations are $[L]_{\text{free}} = [L]_{\text{tot}} - [C]$ and $[R]_{\text{free}} = [R]_{\text{tot}} - [C]$.

In the usual limit where the ligand concentration is much larger than the receptor concentration, the free ligand concentration is essentially unchanged by binding. In that case, the fraction of receptor sites that are occupied is

$$p_R = \frac{[L]_{\text{free}}/K_{\text{D}}}{1 + [L]_{\text{free}}/K_{\text{D}}}. \qquad [39]$$

Here $K_{\text{D}}$ is the dissociation constant for motor-microtubule binding. For the case of interest here, however, the finite reservoirs of both motors and microtubule binding sites must be retained. The binding equilibrium is

$$K_{\mathrm{D}} = \frac{([L]_{\mathrm{tot}} - [C])([R]_{\mathrm{tot}} - [C])}{[C]}. \quad [40]$$

Solving this quadratic equation for the concentration $[C]$ of bound motor-microtubule complexes gives

$$[C] = \frac{[L]_{\mathrm{tot}} + [R]_{\mathrm{tot}} + K_{\mathrm{D}} - \sqrt{([L]_{\mathrm{tot}} + [R]_{\mathrm{tot}} + K_{\mathrm{D}})^2 - 4[L]_{\mathrm{tot}}[R]_{\mathrm{tot}}}}{2}. \quad [41]$$

The physically meaningful solution uses the minus sign, since the plus sign would give a bound-complex concentration larger than one of the available reservoirs. The fraction of motors that are bound to microtubules is therefore

$$p_b = \frac{[C]}{[L]_{\mathrm{tot}}} = \frac{[L]_{\mathrm{tot}} + [R]_{\mathrm{tot}} + K_{\mathrm{D}} - \sqrt{([L]_{\mathrm{tot}} + [R]_{\mathrm{tot}} + K_{\mathrm{D}})^2 - 4[L]_{\mathrm{tot}}[R]_{\mathrm{tot}}}}{2[L]_{\mathrm{tot}}}. \quad [42]$$

This equation is also used in section A.

This result is the basis of the plots of bound motor fraction shown in Figure S40 where we see that at different positions within the aster, there are quite distinct motor bound fractions, and hence, ATP consumption rates. Of course, to make these calculations also demands a variety of parameters that describe the kinetics of binding and hydrolysis, which we discuss in section A

#### 6. Investigation of photobleaching

From an experimental perspective, one of the main objectives of the present work is to determine the concentration of ATP at different positions and times within our microtubule-motor system. However, since our method of making that measurement uses fluorescence, it is incumbent upon us to quantify the unwanted effects of photobleaching, which leads to a reduction in fluorescent signal, as opposed to the physical reduction we are really interested in due to ATP consumption.

To understand and account for the character and extent of photobleaching as it affects our measurements, we consider these dynamics from several angles. We perform control experiments quantifying the intensity reduction in a uniform mixture of fluorescent ATP probes absent any motor activity. We understand the quantitative phenomenology reported by these experiments by building models accounting for the role of a reservoir outside the excitation region, both via simple kinetic and finite element modeling.

In the following sections, we review experiments and models to understand how our probe photobleaches. The ultimate conclusion leads us to an understanding, from many angles, that despite variability in photobleaching dynamics at early times, little to no variation occurs between replicates at intermediate to late times. Thus, to be rigorous and conservative, all of the data reported in the main text are reported only after a stabilization cutoff time, and are corrected for photobleaching once there is a consensus between experimental controls. Next, we discuss this procedure in detail.

**A. Phenomenology of Photobleaching from Control Experiments.** We prepare an ATP calibration experiment, as described in section D, such that we pipette a fixed, known quantity of ATP into the same reaction mixture as an aster experiment; however, we do not include motor proteins, thereby preventing any ATP hydrolysis from occurring. We then image the sample using the same time dynamics as aster experiments, namely matching laser intensity currents, 1000 mA, exposure times, $150 - 160$ ms, and the interval between image acquisition, 20 s. Upon collecting image data, we process it using the pipeline described in section 2 to do background subtraction (section A) and uneven illumination corrections (section B). We then average the intensity value of each image and plot the intensity values over time. This produces scatter plots such as those shown in Figure S41. We see empirically from the lines in Figure S41 that photobleaching in our system is very well described by fitting a single exponential plus a constant of the form

$$f_p(t) = \frac{I(t)}{I_0} = (1 - I_\infty)e^{-t/\tau} + I_\infty, \quad [43]$$

where $I(t)$ is the intensity at time $t$, $I_0$ is the intensity at the start of the experiment, $I_\infty$ is the intensity as time approaches infinity, and $\tau$ is the characteristic decay constant in time.

To fit this decay due to photobleaching, we first divide all data points by the initial value of the intensity, $I_0$, which, for a strictly monotonic function, is the maximum value of the data. Thus, we pin the time zero intensity to 1. Using scipy.optimize.minimize, we find the parameter set, $\underline{p} = (p_0, p_1, \ldots, p_n)$, for a fitting function, $f_p(t)$, that minimizes the sum of the square of the residuals,

$$\mathrm{SSR} = \sum_t (I(t) - f_p(t))^2, \quad [44]$$

where $I(t)$ is the value of the intensity at a time t. From here on, we will define the variables of the parameter set such that $p_0 = \tau$, $p_1 = I_\infty$.

Why does photobleaching adopt this form of an initial exponential decay to a plateau of about 80% of the initial intensity? Ordinarily, photobleaching in a uniformly-illuminated, confined chamber could instead be modeled by a single exponential decay,

$$f_p(t) = e^{-t/\tau}, \quad [45]$$

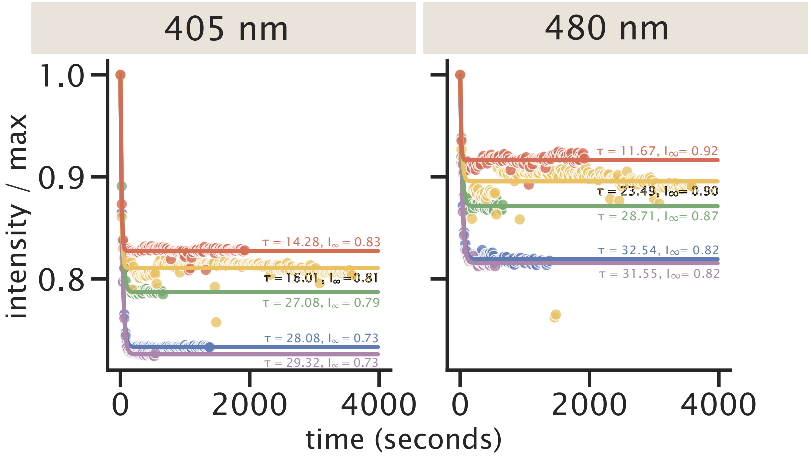

**Fig. S41. Examples of representative, appreciably-variable, raw photobleaching trajectories across (putatively-identical) experiments, and accompanying fits.** Fluorescence intensities $I(t)$ of 405 nm (left column) and 480 nm (right column) excitation channels were each fit adequately to an exponential decay, Equation 43, with a single timescale $\tau$ and a long-time offset $I_\infty$ (once normalized by maximum intensity values at time zero, $I_0$). Lines show best-fit curves and dots are raw data. Each color represents an experimental repeat for data taken at $500\mu M$ ATP and at 20 s intervals.

since the only thing that could change the state of the probe is bleaching according to the dynamical equation,

$$\frac{dI}{dt} = -\frac{1}{\tau} I. \quad [46]$$

However, our situation is more nuanced because the excitation region is small compared to the size of the chamber, meaning non bleached probes can diffuse in, slowing the time scale of intensity decay. We show that a single exponential indeed fits poorly to our data in Figure S42(A). But, as we commented before in Eq. 43, a single exponential plus a constant seems to do the trick. One may be concerned about whether this model is physically reasonable since at sufficiently long timescales, the underlying physical picture would still expect that all the probes to have been bleached. To reckon with this question, in section D.1, we show that when diffusive replenishment is invoked in a first order kinetic model where probes adopt two (bleached or unbleached) states, the decay of visible (unbleached) probe is modeled by a double exponential,

$$f_p(t) = (1 - I_\infty)e^{-t/\tau} + I_\infty e^{-t/\tau_2}, \quad [47]$$

where $\tau_2$ is a second emergent timescale in the physical problem. If this second time constant $\tau_2$ is very long, the behavior describing the photobleaching dynamics is well approximated by a single exponential plus a constant. In Figure S42(C), a double exponential fit seems to fit just as well as a single exponential plus a constant as shown in Figure S42(B). However, we note that the the solver kept the initial guess for $\tau_2$, the long decay constant. In adjusting initial guesses and solver tolerances, we conclude that experimental data do not offer sufficiently long time courses for the solver to get a good fit for this parameter. However, we do know it is much much longer than the timescale of our experiment. Indeed, in Figure S43, we show that for $\tau_2 > 3 \times 10^5$, the squared sum of residuals for most replicates have reached a plateau and there is no further refinement of this parameter to achieve. If our experiment contained data at much longer times, we could more clearly witness the effect of the second decay constant.

**B. Sensitivity of photobleaching to experimental conditions.** When correcting for photobleaching in aster data, one of the things we need to check is if the photobleaching decay fits vary with ATP concentration. In Figure S44, we plot the fit parameters as a function of ATP concentration. There is no clear trend with ATP, indicating that the probe photobleaching process is not ATP-concentration dependent. Thus, regardless of an aster's ATP concentration, we can process images using appropriate globally-inferred parameters to the photobleaching decay curves.

We also examine how the fit parameters scale with modulating the time interval between light excitations. We expect that for shorter times between pulses, the bleaching of probes will be more continuous leading to shorter decay constants, while for longer intervals between light pulses the time constant should take much longer to see reduction in light intensity, which we have simulated in section F. In Figure S45, we find the decay constant, $\tau$, scales about linearly with the interval between light excitations. In contrast, there is only small variation in the $I_\infty$ parameter with respect to the pulsing interval.

Amid the variation in these parameters discussed above, key quantitative regularities emerge. Across experiments, all the photobleaching decay time constants are shorter than $\tau \lesssim 35$ s. In addition, all plateau intensity values $I_\infty$ vary by less than 20%. Together, these two facts have the consequence that the late time behaviors of photobleaching trajectories agree within a few tens of percent of each other. This result can be understood by referring to equation 43, $I(t)/I_0 = (1 - I_\infty)e^{-t/\tau} + I_\infty$. After waiting a few decay constants—reached early in the experiment even for the slowest photobleaching trajectory—the first term $(1 - I_\infty)e^{-t/\tau}$ vanishes. Since the remaining plateau constant, $I_\infty$, varies only mildly (by less than 20%), the overall trajectories ultimately agree within a few tens of percent of each other after long enough times.

**C. Photobleaching correction procedure.** With the form of the time-dependent intensity reduction in hand, we can now account for the amount of depletion due to photobleaching over time to recover the "true" intensity. In particular, we interpret the measured intensity as the product of a weighting function and the true intensity if fluorophores were not bleached. Namely, when $\tilde{I}$ is the true intensity and $I$ is the measured intensity,

$$\tilde{I}(t) = \frac{I(t)}{f_p(t)} = \frac{I(t)}{(1 - I_\infty)e^{-t/\tau} + I_\infty}. \quad [48]$$

Since our probe calibrations are ratios of image intensities collected at two wavelengths, we can independently correct images from each wavelength and then take the ratio. This means the effective correction to the overall ratio is $\frac{f_{480}(t)}{f_{405}(t)}$, yielding,

$$\frac{\tilde{I}_{405}(t)}{\tilde{I}_{480}(t)} = \frac{I_{405}(t) f_{480}(t)}{I_{480}(t) f_{405}(t)}. \quad [49]$$

In Figure S46 we plot the correction values over time for five characteristic experimental replicates. In addition, we plot the consensus of the fitted parameters, $\tau$ and $I_\infty$, as determined by the linear fit evaluated at the acquisition interval, 20 seconds, in Figure S45. The kinetic variability in fit parameters across experiments propagates to implied corrections. The differences are most apparent at early times before the fit functions plateau. We can see how these corrections modify the number of ATP versus time for a sample aster experiment in Figure S47.

As expected, the corrected ATP readouts are greater than the uncorrected measurement, yet still deplete over time as the aster consumes ATP. However, control photobleaching trajectories vary considerably across replicates and conditions at early time points. In Figure S47, these discrepancies manifest in corrected ATP trajectories until roughly 250 seconds ($\sim 4$ min).

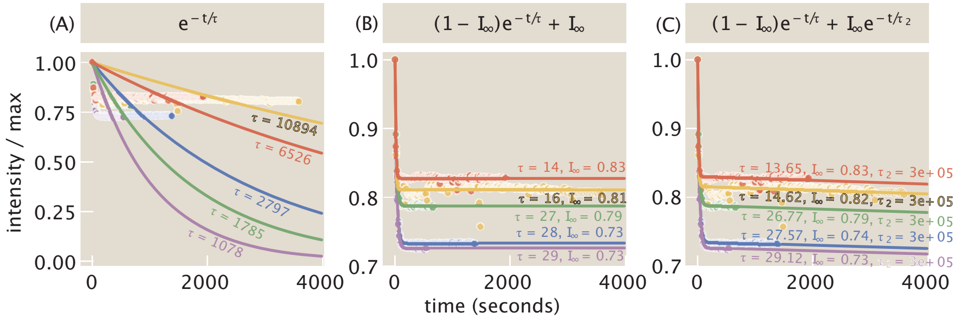

**Fig. S42. Photobleaching data fit to several functional forms.** (A) Single exponential fits do not do an acceptable job of capturing the intensity depletion due to photobleaching and lead to large values of the decay constant. (B) Fitting to a single exponential plus a constant provides a satisfactory fit to the form of measured intensity reduction. (C) A double exponential plus a constant also fits the measured intensity decline, though the solver does not capture the second, longer decay constant very well. All data in these plots are taken at $500\ \mu\mathrm{M}$ ATP and at $20$ s intervals. Each color represents the data from the same experimental repeats as depicted in Figure S41. Values for $\tau$ and $\tau_2$ have units of seconds and values for $I_\infty$ are unitless, since the intensities are normalized by the maximum intensity $I_0$ at time zero.

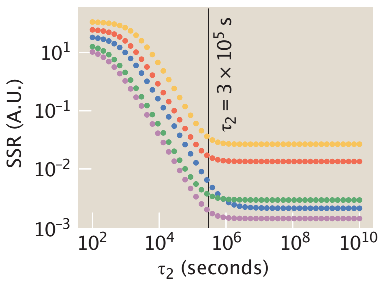

**Fig. S43. Evaluation of fit for a second decay constant.** Using the fit parameters in Figure S41, we plot how the minimization of the sum of the square of the residuals (SSR) to the data depends on the value of a second decay constant, $\tau_2$. For values larger than approximately $3 \times 10^5$ seconds, the SSR reaches a minimization limit beyond which there is no further improvement to the fit . Each color represents the data from the same experimental repeats as depicted in Figure S41.

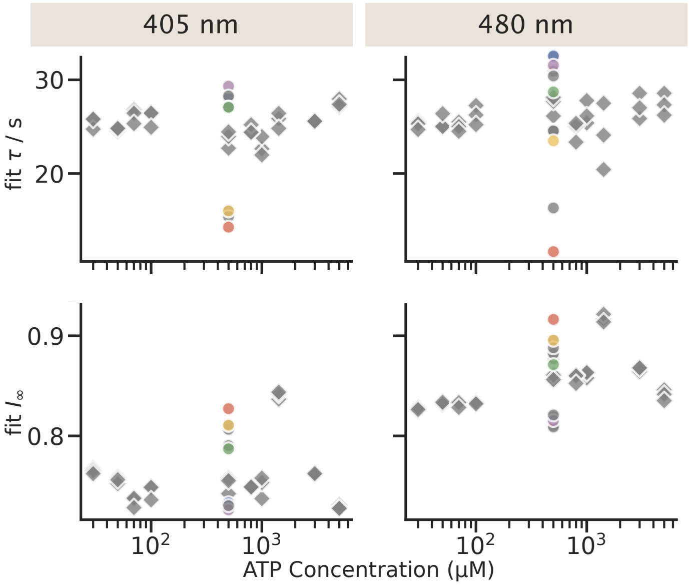

**Fig. S44. ATP dependence of the fit parameters for photobleaching.** The fit constants, $\tau$ and $I_\infty$, of the function, $I(t)/I_0 = (1 - I_\infty)e^{-t/\tau} + I_\infty$, are plotted as a function of ATP concentration. None of the fit parameters seem to have a clear trend as a function of ATP concentration. Circular points and diamond-shaped points display data from experiments conducted on separate dates. All data points are taken with the same imaging parameters. Each color represents the data from the same experimental repeats as depicted in Figure S41.

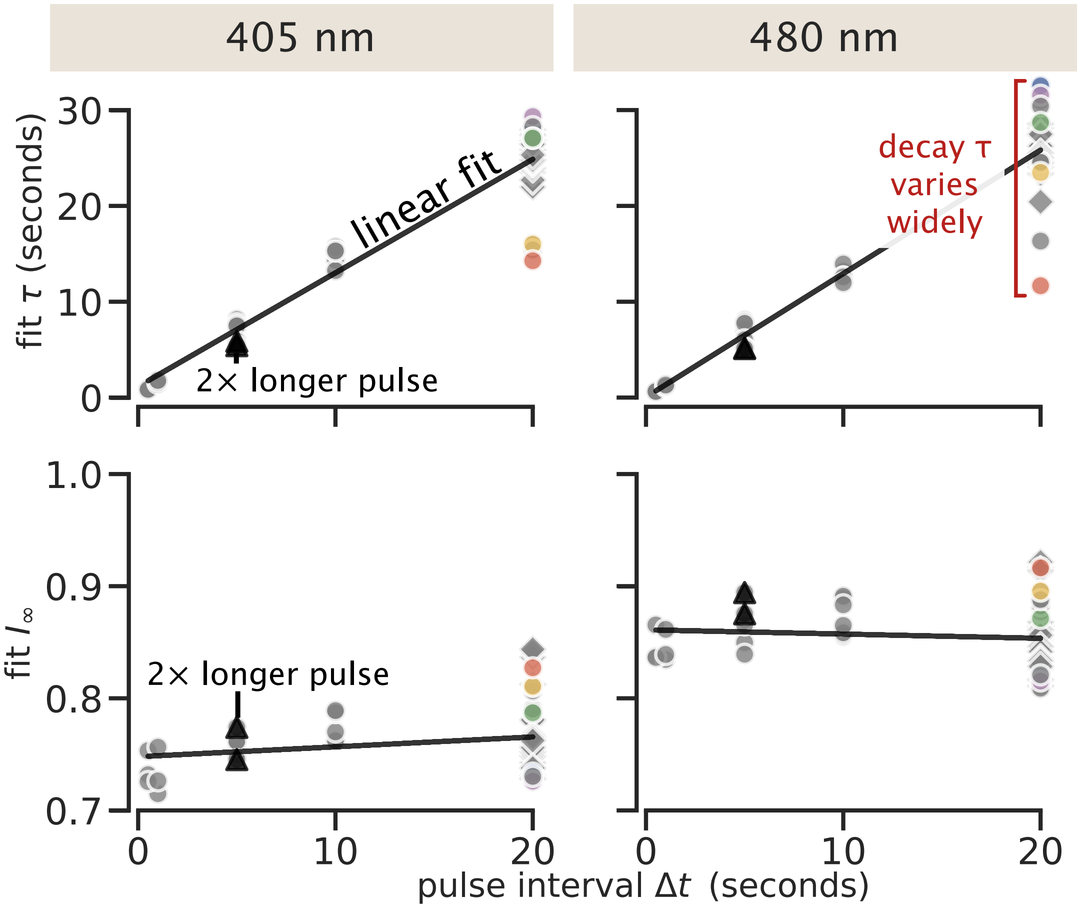

**Fig. S45. Intrinsic and systematic variability of photobleaching kinetic parameters across days and excitation pulse intervals $\Delta t$.** On average, the fit decay time $\tau$ and offset $I_\infty$ vary linearly with the interval $\Delta t$ between excitation pulses in the measurement, as reported by acceptable linear fits between these parameters to the pulse interval $\Delta t$ (shown as black lines). Five distinctly colored examples, the same experimental repeats as described by Figure S41, are taken at the pulse interval of $\Delta t = 20$ s, the interval typically used between frames in aster experiments, are highlighted, evidencing wide and ultimately meaningful variability in photobleaching kinetics across experiments. For additional context, two additional measurements with a 2×-longer excitation pulse (300 ms instead of the typical 150 ms) are displayed as triangles (▲), at a pulse interval of $\Delta t = 5$ s. Circular points and diamond-shaped points display data from experiments conducted on separate dates.

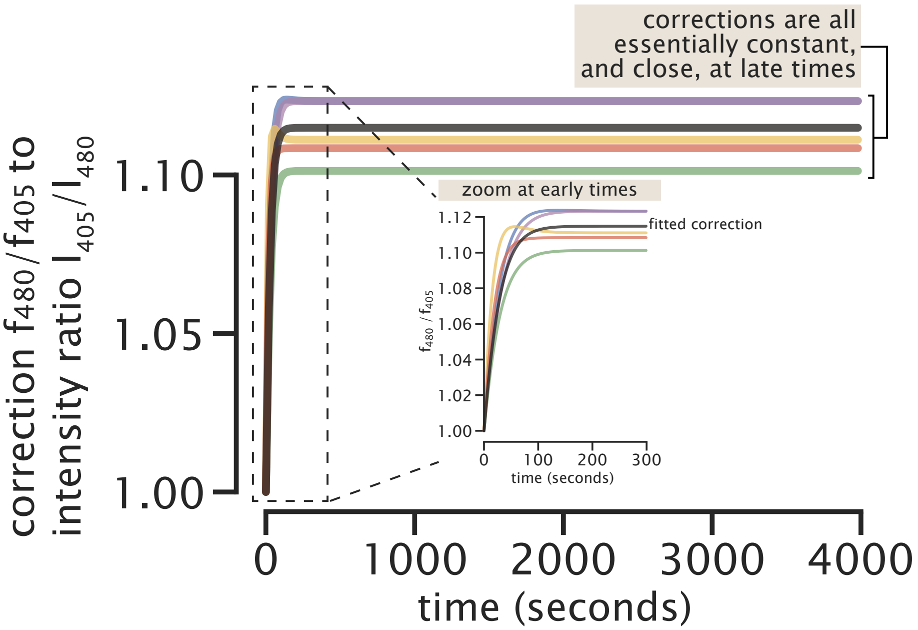

**Fig. S46. Variation in the ratiometric photobleaching correction implied by variability in photobleaching experimental fits.** Specifically, each fluorescence channel offers a photobleaching fit function $f(t) = I(t)/I_{\text{max}} = (1 - I_\infty)\exp\left[-t/\tau\right] + I_\infty$. The measured fluorescence intensity ratio $r(t) = I_{405}(t)/I_{480}(t)$ would be corrected by the ratiometric correction $f_{480}(t)/f_{405}(t)$—and then is ultimately transformed to ATP concentration by a separate nonlinear concentration calibration function. Experimental variability in the raw kinetic parameters $\tau$ and $I_\infty$ of each fluorescence channel thus propagate in rich ways to the shape of the ratiometric correction function. Notice even the appearance of non-monotonicity in time in the yellow and blue experimental fits' correction curves, which is facilitated by mismatches in the timescales of bleaching in the 405 nm and 480 nm channels. Each line is colored with the same scheme as the repeats described by Figure S41 and an additional line, the dark gray, plots the correction function when $\tau$ and $I_\infty$ are set by the value of the linear fit at 20 seconds in Figure S45.

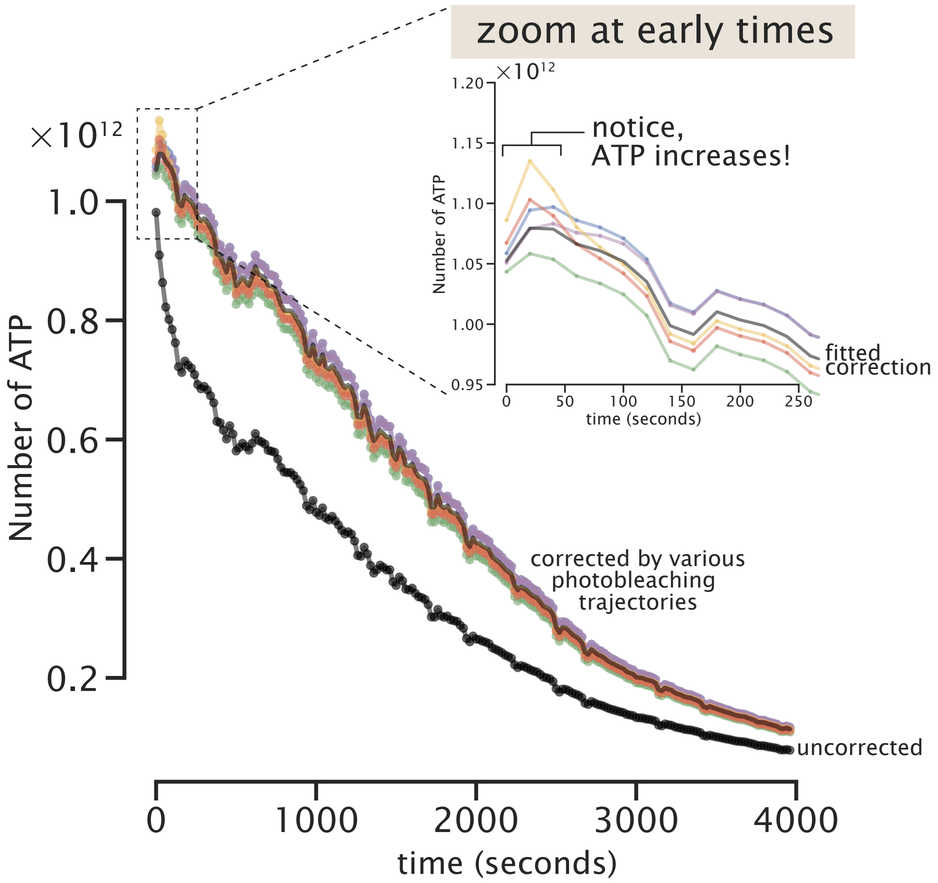

**Fig. S47. Implications of variability across experimental photobleaching trajectories for inferred ATP abundance over time.** The black trajectory shows the number of ATP molecules over time of a representative aster if photobleaching were left uncorrected. The colored curves above are the same underlying aster data subjected to exemplary but diverse ratiometric photobleaching corrections (ensuing from those in Figure S46 and colored correspondingly). The dark gray curve, is the result of correcting using the $\tau$ and $I_\infty$ values of the linear fit at 20 seconds in Figure S45. Notice that all corrected trajectories show an apparent *increase* in ATP over at least the first pair of data points, with e.g. the blue and purple trajectories sustaining this apparent increase in ATP over at least two time points. Given that motors are continually consuming ATP, we expect the appearance of such an increase in ATP is pathological/nonphysical, emphasizing how profoundly photobleaching trajectories confound the resolution of ATP dynamics at early time points.

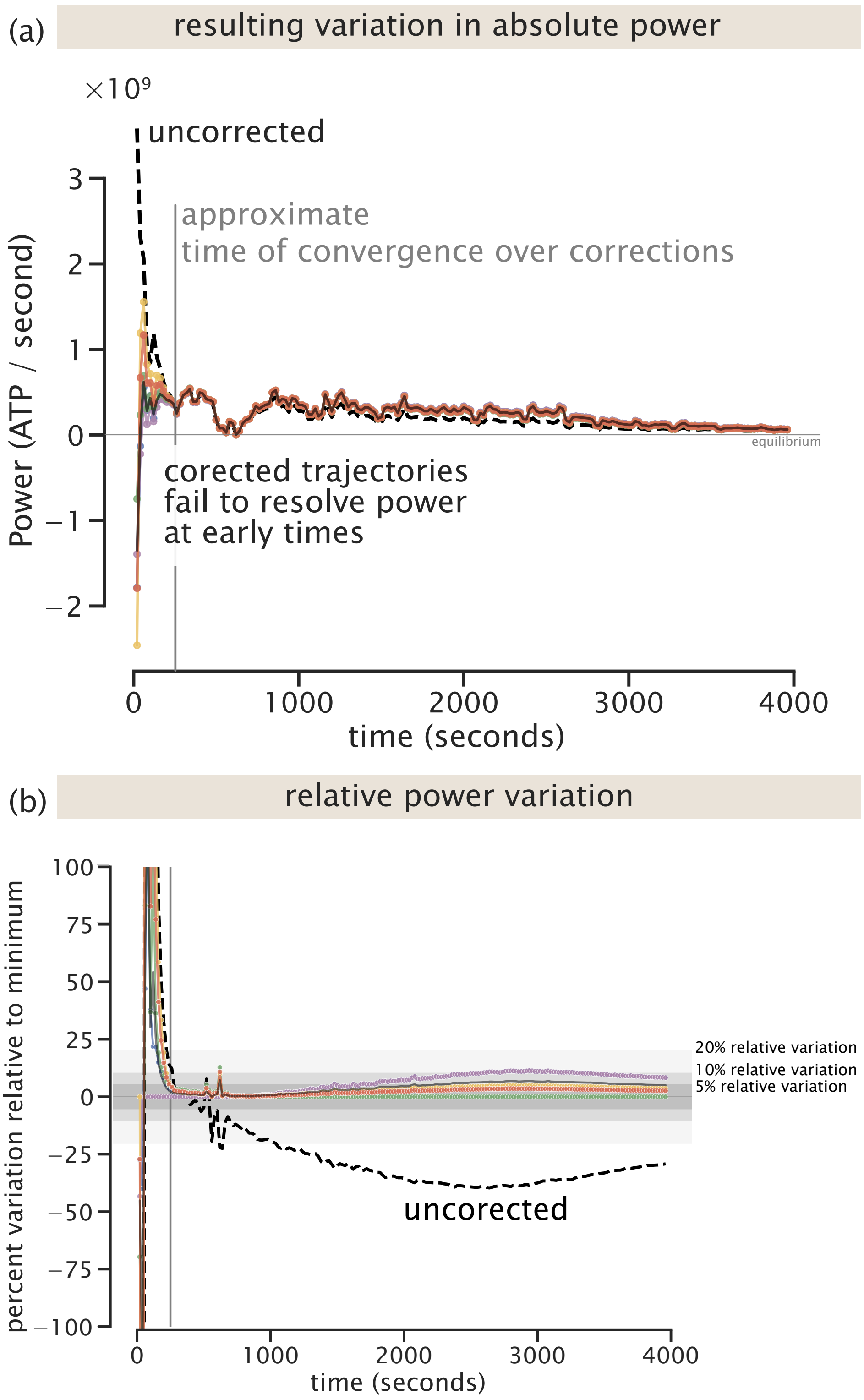

**Fig. S48. Implications of variability across experimental photobleaching trajectories for inferred power consumption (ATP/second) over time.** (a) The black trajectory shows the consumption rate of ATP molecules over time of a representative aster if photobleaching were left uncorrected. The colored curves are the same underlying aster data subjected to exemplary but diverse ratiometric photobleaching corrections (ensuing from those in Figure S46and colored correspondingly). The dark gray curve, is the result of correcting using the $\tau$ and $I_\infty$ values of the linear fit at 20 seconds in Figure S45. Notice the dramatic ambiguity in even the *sign* of the ATP consumption rate, atop the quantitative disagreement across corrections, at early times. However, after a few characteristic photobleaching time constants (approximately $t \gtrsim 250\ s$, indicated by the gray line), the diversely-corrected curves closely agree. (b) In determining a cutoff time for where the varied photobleach corrected ATP curves agree, we plot the percent variation at each time for each power curve as compared to the curve with the minimum power. The variation is computed as $\frac{\text{power}_i(t) - \min(t)}{\min(t)} \times 100$, where $i$ indexes over each corrected curve and $\min(t)$ refers to the minimum value of power at a given time $t$. The variation curves are laid upon three shaded regions depicting 5, 10, and 20 percent variation.

Further, some candidate corrections could even spuriously show a mild increase in ATP over the earliest time points. (We regard such an apparent increase as nonphysical, given the continual consumption of ATP by motors.) This potential problem is even more apparent when examining power (namely the derivative of ATP over time), as shown in Figure S48. Crucially, however, after roughly 250 seconds (nearly ten times longer than the fit decay constant $\tau$ values), power trajectories across all candidate corrections align nearly identically. This concordance of the inferred ATP and power trajectories only after 250 seconds recommends that these trajectories are reliably resolvable after this short initial period.

In Figure S49, we see that the initial period is early in the meaningful development of the aster structures. Figure S49(C) depicts the the state of motor and ATP gradients at the proposed cutoff time. The motor image does not yet show any structure formation, reinforcing our recommendation to start analyzing data after the initial $t \gtrsim 250$ s period. Taking a closer look at amount of gradients developed in ATP, the grey lines in Figure S49(A) and (B) show the ATP profiles at all measured time points before the cutoff. After correcting for photobleaching, we find that although ATP does begin depleting, ATP throughout the system is still abundant. Additionally nearly all the gradient formation in ATP occurs after the cutoff. Given that nearly all development of spatial gradients, in both ATP and motors, occur after the cutoff and ATP is still at meaningful abundances, we proceed with our analysis examining data following this practical cutoff.

**D. Theory of the Experiment: Analysis of Photobleach-Diffusion Equations.** To develop intuition for the way in which photobleaching alters our fluorescence signal and hence could contaminate our measurement, we worked to develop a "theory of the experiment" to compute how the dynamics of the effect we are trying to measure (i.e. ATP consumption) couple with the unwanted effects of photobleaching. This effort is one of many such examples within our laboratory where we have repeatedly found it very useful to do our best job of writing down a quantitative theoretical description of the experiment and using that description to calibrate our experiments themselves (21, 22). This is not a substitute for the actual experimental correction described in the previous section C, but rather a complementary approach for developing intuition for the scale and dynamics of the photobleaching effect.

In this section, we show how to set up quantitative models of the photobleaching and diffusion of our fluorescent probes and how the model equations can be solved as the basis of proposed photobleaching corrections in our experiments. First, in the ensuing sections D.1 and D.2, we write a model for the fluorescent probe independent of space, where the probe adopts two states, bleached or unbleached. This model can be used to account for the character of photobleaching in our experiments.

Figure S50 shows the geometry of one of our experimental devices. As shown, the experimental system is a giant reservoir encompassing a small region where illumination destroys the visibility of some molecules through photobleaching. This geometry sets up conditions for photobleaching to compete with bulk replenishment from the reservoir, as we now explore.

***D.1. Photobleaching in a reservoir takes the form of a double exponential.*** When exciting a region of our sample, there is some rate, $k_{\text{bleach}}$, at which probes in this region are photobleached. These probes are then lost from the pool of probes we can account for through fluorescence. Within the excitation region, there is some concentration of fluorescent probes, $c_{\text{in}}$, and outside of this region, there is additional pool of fluorescent probes with concentration, $c_{\text{out}}$. Probes can diffuse between these regions with rates $k_{\text{in}}$ and $k_{\text{out}}$. We write a model for the kinetics of probe movement. The time dependence of non-bleached probes in the illumination region is

$$\frac{dc_{\text{in}}(t)}{dt} = -k_{\text{bleach}}c_{\text{in}}(t) - k_{\text{out}}c_{\text{in}}(t) + k_{\text{in}}c_{\text{out}}(t), \qquad [50]$$

while the time dependence of non-bleached probes outside the illumination region is,

$$\frac{dc_{\text{out}}(t)}{dt} = k_{\text{out}}c_{\text{in}}(t) - k_{\text{in}}c_{\text{out}}(t). \qquad [51]$$

Writing the system of equations as in matrix form results in

$$\frac{d}{dt}\begin{bmatrix} c_{\text{in}} \\ c_{\text{out}} \end{bmatrix} = \begin{bmatrix} -k_{\text{bleach}} - k_{\text{out}} & k_{\text{in}} \\ k_{\text{out}} & -k_{\text{in}} \end{bmatrix}\begin{bmatrix} c_{\text{in}} \\ c_{\text{out}} \end{bmatrix}, \qquad [52]$$

and applying the ansatz,

$$\begin{aligned} c_{\text{in}}(t) &= c_{\text{in}}^{(0)}e^{\sigma t} \\ c_{\text{out}}(t) &= c_{\text{out}}^{(0)}e^{\sigma t}, \end{aligned} \qquad [53]$$

we can solve the system of equations as an eigenvalue problem. Plugging in the ansatz into the matrix form,

$$\sigma e^{\sigma t}\begin{bmatrix} c_{\text{in}}^{(0)} \\ c_{\text{out}}^{(0)} \end{bmatrix} = e^{\sigma t}\begin{bmatrix} -k_{\text{bleach}} - k_{\text{out}} & k_{\text{in}} \\ k_{\text{out}} & -k_{\text{in}} \end{bmatrix}\begin{bmatrix} c_{\text{in}}^{(0)} \\ c_{\text{out}}^{(0)} \end{bmatrix}, \qquad [54]$$

which we rearrange to find

$$0 = \begin{bmatrix} -k_{\text{bleach}} - k_{\text{out}} - \sigma & k_{\text{in}} \\ k_{\text{out}} & -k_{\text{in}} - \sigma \end{bmatrix}\begin{bmatrix} c_{\text{in}}^{(0)} \\ c_{\text{out}}^{(0)} \end{bmatrix}. \qquad [55]$$

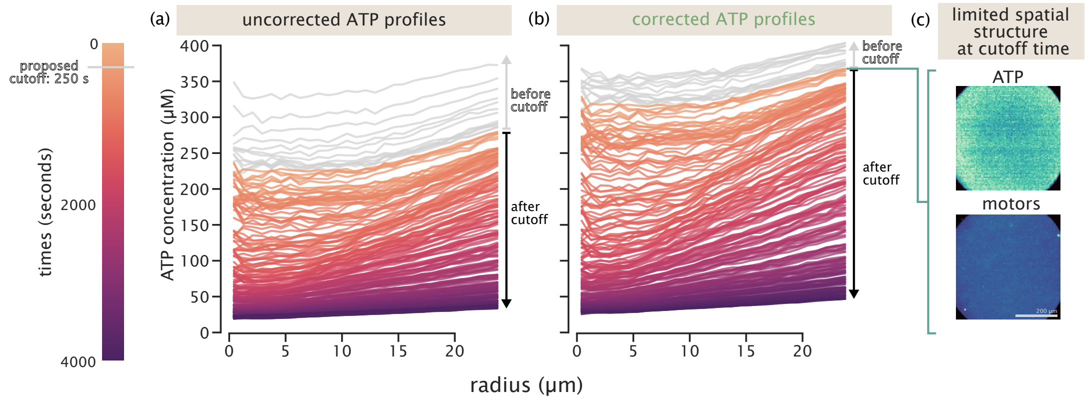

**Fig. S49. ATP traces over space and time, with and without an exemplary photobleaching correction, relative to a proposed cutoff time—before which we propose deeming ATP dynamics quantiatively-unresolvable.**(a) Spatiotemporal profiles of ATP if no photobleaching correction were applied. (b) Spatiotemporal profiles of ATP under an exemplary photobleaching correction (corresponding to the green example photobleaching trajectory fit in the preceding figures in this section. In panels (a) and (b), profiles before the proposed cutoff time of 250 s—the same cutoff indicated by in Figure S48 by the vertical gray line—are colored in grey (leaving the remaining profiles, which we propose retaining, colored by time). (c) For these representative aster data, the spatial profiles of both ATP and motors have developed only modest spatial gradients at the proposed cutoff time (250 seconds, 13 frames). This means the retained data contain the bulk of interesting spatiotemporal gradient formation attending the aster formation.

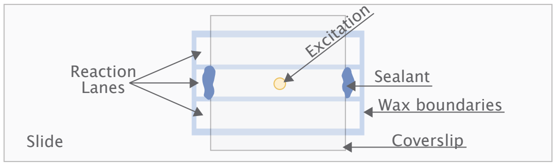

**Fig. S50. Anatomy of the experimental chamber.** To create our experimental chamber, we cut 3 mm × 18mm lanes out of parafilm wax. We sandwich the wax sheet between a coverslip and a slide leaving the ends of the lanes exposed. This allows us to flow a reaction mixture into each lane. We seal the reaction in the lanes using picodent twinsill speed, a polymeric seal.

Taking the determinant,

$$\begin{aligned} 0 &= (-k_{\text{bleach}} - k_{\text{out}} - \sigma)(-k_{\text{in}} - \sigma) - k_{\text{in}}k_{\text{out}} \\ &= \sigma^2 + (k_{\text{bleach}} + k_{\text{in}} + k_{\text{out}})\sigma + k_{\text{in}}(k_{\text{bleach}} + k_{\text{out}}) - k_{\text{in}}k_{\text{out}} \\ &= \sigma^2 + (k_{\text{bleach}} + k_{\text{in}} + k_{\text{out}})\sigma + k_{\text{in}}k_{\text{bleach}}. \end{aligned} \quad [56]$$

Solving for the roots of $\sigma$,

$$\sigma = -\frac{1}{2}(k_{\text{bleach}} + k_{\text{in}} + k_{\text{out}}) \pm \frac{1}{2}\sqrt{(k_{\text{bleach}} + k_{\text{in}} + k_{\text{out}})^2 - 4k_{\text{in}}k_{\text{bleach}}} \quad [57]$$

Plugging this partial differential equation into Mathematica and solving for $c_{\text{in}}(t)$, we find,

$$c_{\text{in}}(t) = \frac{c_0 \exp\left(-\frac{(\alpha+\gamma)t}{2}\right)}{2\gamma}\left(\beta - \beta\exp(\gamma t) + \gamma + \gamma\exp(\gamma t)\right), \quad [58]$$

where, for ease of reading, we define the constants

$$\begin{aligned} \alpha &\equiv k_{\text{bleach}} + k_{\text{in}} + k_{\text{out}}, \\ \beta &\equiv k_{\text{bleach}} - 3k_{\text{in}} + k_{\text{out}}, \\ \gamma &\equiv \sqrt{-4k_{\text{bleach}}k_{\text{in}} + (k_{\text{bleach}} + k_{\text{in}} + k_{\text{out}})^2} = \sqrt{-4k_{\text{bleach}}k_{\text{in}} + \alpha^2}. \end{aligned} \quad [59]$$

We simplify the form of $c_{\text{in}}(t)$ by distributing the exponential prefactor,

$$c_{\text{in}}(t) = \frac{c_0}{2\gamma}\left(\beta e^{-(\alpha+\gamma)t/2} - \beta e^{-(\alpha-\gamma)t/2} + \gamma e^{-(\alpha+\gamma)t/2} + \gamma e^{-(\alpha-\gamma)t/2}\right). \quad [60]$$

Combining the terms in the parenthesis by exponentials,

$$c_{\text{in}}(t) = \frac{c_0}{2\gamma}\left((\gamma + \beta)e^{-(\alpha+\gamma)t/2} + (\gamma - \beta)e^{-(\alpha-\gamma)t/2}\right), \quad [61]$$

This equation establishes that the visible abundance of probes in the monitored region, $c_{\text{in}}(t)$, can be described as a double exponential with two characteristic timescales $\tau_1$ and $\tau_2$, given by,

$$\begin{aligned} \tau_1^{-1} &= (\alpha + \gamma)/2 \\ &= \frac{1}{2}\left(k_{\text{bleach}} + k_{\text{in}} + k_{\text{out}} + \sqrt{-4k_{\text{bleach}}k_{\text{in}} + (k_{\text{bleach}} + k_{\text{in}} + k_{\text{out}})^2}\right) \end{aligned} \quad [62]$$

and

$$\begin{aligned} \tau_2^{-1} &= (\alpha - \gamma)/2 \\ &= \frac{1}{2}\left(k_{\text{bleach}} + k_{\text{in}} + k_{\text{out}} - \sqrt{-4k_{\text{bleach}}k_{\text{in}} + (k_{\text{bleach}} + k_{\text{in}} + k_{\text{out}})^2}\right). \end{aligned} \quad [63]$$

We remark that since $\gamma \geq 0$, we see that $\tau_1^{-1} \geq \tau_2^{-1}$ and so $\tau_1 \leq \tau_2$. That is, the more rapid decay process is captured by the first timescale $\tau_1$.

In Section A, we find that the predictions of this model do a reasonable job of describing the photobleaching in our experiments. Specifically, our data is best fit to a double exponential with a short and a long time constant, corresponding to $\tau_2 \gg \tau_1$ and yielding an effective plateau in fluorescence on experimental timescales.

To further grapple with other origins of photobleaching, we also explored what modifications to the model are introduced by probes sticking to the glass coverslip or slide. Next, we include our findings here for those practitioners interested.

***D.2. Photobleaching with surface bound probes in a reservoir takes the form of a triple exponential.*** As above, we write model for the bleaching dynamics inside and outside the excitation region, but now also take into account an additional effect where probes can be bound to the surface and do not diffuse. These probes can still bleach when inside the excitation region. We denote the concentration of probes inside the excitation region with variable $c_{\text{s-in}}$ and outside the excitation region as, $c_{\text{s-out}}$. We include two more rate constants, $k_{\text{on}}$ and $k_{\text{off}}$, for the rates of binding and unbinding to the surface. Thus, our system of equations is now,

$$\begin{aligned}\frac{dc_{\text{in}}(t)}{dt} &= -k_{\text{bleach}}c_{\text{in}}(t) - k_{\text{out}}c_{\text{in}}(t) + k_{\text{in}}c_{\text{out}}(t) - k_{\text{on}}c_{\text{in}}(t) + k_{\text{off}}c_{\text{s-in}}(t),\\ \frac{dc_{\text{out}}(t)}{dt} &= k_{\text{out}}c_{\text{in}}(t) - k_{\text{in}}c_{\text{out}}(t) - k_{\text{on}}c_{\text{out}}(t) + k_{\text{off}}c_{\text{s-out}}(t),\\ \frac{dc_{\text{s-in}}(t)}{dt} &= -k_{\text{bleach}}c_{\text{s-in}}(t) + k_{\text{on}}c_{\text{in}}(t) - k_{\text{off}}c_{\text{s-in}}(t),\\ \frac{dc_{\text{s}}(t)}{dt} &= k_{\text{on}}c_{\text{out}}(t) - k_{\text{off}}c_{\text{s-out}}(t)\end{aligned} \tag{64}$$

Writing the system of equations as in matrix form, we have

$$\frac{d}{dt}\begin{bmatrix} c_{\text{in}} \\ c_{\text{out}} \\ c_{\text{s-in}} \\ c_{\text{s-out}} \end{bmatrix} = \begin{bmatrix} -k_{\text{bleach}} - k_{\text{out}} - k_{\text{on}} & k_{\text{in}} & k_{\text{off}} & 0 \\ k_{\text{out}} & -k_{\text{in}} - k_{\text{on}} & 0 & k_{\text{off}} \\ k_{\text{on}} & 0 & -k_{\text{bleach}} - k_{\text{off}} & 0 \\ 0 & k_{\text{on}} & 0 & -k_{\text{off}} \end{bmatrix} \begin{bmatrix} c_{\text{in}} \\ c_{\text{out}} \\ c_{\text{s-in}} \\ c_{\text{s-out}} \end{bmatrix}. \tag{65}$$

As a first guess, we assume the binding kinetics to the surfaces are very slow, slower than the duration of the experiment. So we simplify this system of equations such that where surface bound probes are only subject to bleaching and do not communicate with the pool of other probes. Thus our system of equations is,

$$\begin{aligned}\frac{dc_{\text{in}}(t)}{dt} &= -k_{\text{bleach}}c_{\text{in}}(t) - k_{\text{out}}c_{\text{in}}(t) + k_{\text{in}}c_{\text{out}}(t),\\ \frac{dc_{\text{out}}(t)}{dt} &= k_{\text{out}}c_{\text{in}}(t) - k_{\text{in}}c_{\text{out}}(t),\\ \frac{dc_{\text{s-in}}(t)}{dt} &= -k_{\text{bleach}}c_{\text{s-in}}(t),\\ \frac{dc_{\text{s-out}}(t)}{dt} &= 0.\end{aligned} \tag{66}$$

Thus, the concentration of surface bound probes outside the excitation region is constant in time, and we can ignore the last equation. Now referring to the surface bound probes in the excitation region as $c_{\text{s}}$, we write the system of equations in matrix form,

$$\frac{d}{dt}\begin{bmatrix} c_{\text{in}} \\ c_{\text{out}} \\ c_{\text{s}} \end{bmatrix} = \begin{bmatrix} -k_{\text{bleach}} - k_{\text{out}} & k_{\text{in}} & 0 \\ k_{\text{out}} & -k_{\text{in}} & 0 \\ 0 & 0 & -k_{\text{bleach}} \end{bmatrix} \begin{bmatrix} c_{\text{in}} \\ c_{\text{out}} \\ c_{\text{s}} \end{bmatrix}. \tag{67}$$

As above, we prescribe the ansatz that the concentrations of each species change exponentially in time,

$$\begin{aligned} c_{\text{in}}(t) &= c_{\text{in}}^{(0)}e^{\sigma t}\\ c_{\text{out}}(t) &= c_{\text{out}}^{(0)}e^{\sigma t}\\ c_{\text{s}}(t) &= c_{\text{s}}^{(0)}e^{\sigma t}\end{aligned} \tag{68}$$

we can solve the system of equations as an eigenvalue problem. Plugging in the ansatz into the matrix form,

$$\sigma e^{\sigma t}\begin{bmatrix} c_{\text{in}}^{(0)} \\ c_{\text{out}}^{(0)} \\ c_{\text{s}}^{(0)} \end{bmatrix} = e^{\sigma t}\begin{bmatrix} -k_{\text{bleach}} - k_{\text{out}} & k_{\text{in}} & 0 \\ k_{\text{out}} & -k_{\text{in}} & 0 \\ 0 & 0 & -k_{\text{bleach}} \end{bmatrix} \begin{bmatrix} c_{\text{in}}^{(0)} \\ c_{\text{out}}^{(0)} \\ c_{\text{s}}^{(0)} \end{bmatrix}, \tag{69}$$

which we rearrange to find

$$0 = \begin{bmatrix} -k_{\text{bleach}} - k_{\text{out}} - \sigma & k_{\text{in}} & 0 \\ k_{\text{out}} & -k_{\text{in}} - \sigma & 0 \\ 0 & 0 & -k_{\text{bleach}} - \sigma \end{bmatrix} \begin{bmatrix} c_{\text{in}}^{(0)} \\ c_{\text{out}}^{(0)} \\ c_{\text{s}}^{(0)} \end{bmatrix}. \tag{70}$$

Taking the determinant,

$$\begin{aligned} 0 &= \Big((-k_{\text{bleach}} - k_{\text{out}} - \sigma)(-k_{\text{in}} - \sigma) - k_{\text{in}}k_{\text{out}}\Big)\Big(-k_{\text{bleach}} - \sigma\Big)\\ &= \Big(\sigma^2 + (k_{\text{bleach}} + k_{\text{in}} + k_{\text{out}})\sigma + k_{\text{in}}(k_{\text{bleach}} + k_{\text{out}}) - k_{\text{in}}k_{\text{out}}\Big)\Big(-k_{\text{bleach}} - \sigma\Big)\\ &= \Big(\sigma^2 + (k_{\text{bleach}} + k_{\text{in}} + k_{\text{out}})\sigma + k_{\text{in}}k_{\text{bleach}}\Big)\Big(-k_{\text{bleach}} - \sigma\Big).\end{aligned} \tag{71}$$

Solving for the roots of $\sigma$, the first set of big parenthesis is the same quadratic equation as before, giving the two roots,

$$\sigma = -\frac{1}{2}(k_{\text{bleach}} + k_{\text{in}} + k_{\text{out}}) \pm \frac{1}{2}\sqrt{(k_{\text{bleach}} + k_{\text{in}} + k_{\text{out}})^2 - 4k_{\text{in}}k_{\text{bleach}}}. \quad [72]$$

And the second big parenthesis yields a third root which is simply,

$$\sigma = -k_{\text{bleach}}. \quad [73]$$

Accordingly, very interestingly, bleaching with a population of probes stuck to a surface simply adds an exponential to the model, yielding a triple exponential function. To ask whether our control experiment data showed conspicuous signals of such surface sticking, we also subjected them to the triple exponential model, but concluded the fits were no better than fitting to a double exponential. Given that second time constant values already exceed the duration of the experiment, our data alone do not garner further information about a third timescale, if one should participate.

**E. Time-dependent bleaching rates from periodic excitations.** Our models in the preceding sections D.1 and D.2 investigated the case where the rate of photobleaching $k_b$ is continuous and constant in time. In a real experiment monitoring aster formation, however, the sample is illuminated only in pulses over time, yielding frames of movies. Accordingly, the rate at which probes bleach is time dependent, so that the decay of fluorescence intensity $u(t)$ is given by the ordinary differential equation,

$$\frac{\mathrm{d}u(t)}{\mathrm{d}t} = -k_b(t)u(t), \quad [74]$$

where $k_b(t)$ is the rate of photobleaching. Our experiments acquire images every 20 seconds, where light only excites the sample when an image is being taken presenting a fixed exposure time of 150 ms. To understand how such time-varying bleaching rates shape trajectories of photobleaching, we now explore properties of a simple model evoking the periodic excitations of experiments. Crucially, this understanding also delivers standard benchmark solutions that validate the numerical accuracy and stability of our most realistic, precise, and spatially-explicit simulations from finite element methods that follow next, in section F.

First, to capture pulsatile excitations, we prescribe an example time-dependent photobleaching rate function as,

$$k(t) \equiv k_b \frac{1 + \sin(2\pi t/\tau)}{2}, \quad [75]$$

where $\tau$ is the period of light pulses. This function is visualized in Figure S51(C). The rate of photobleaching is a periodic function between 0 and $k_b$. In this simple test, we do not account for diffusion of probes, only considering a degradation rate due to bleaching. The ODE for the concentration of bright probes is thus,

$$\frac{\mathrm{d}u(t)}{\mathrm{d}t} = -k(t)u(t) = -\frac{k_b u(t)}{2}\left(1 + \sin\left(\frac{2\pi t}{\tau}\right)\right). \quad [76]$$

We can solve for $u(t)$ by using the separation of variables technique, resulting in the expression

$$\int_{u(0)}^{u(t)} \frac{\mathrm{d}u(t')}{u(t')} = \int_0^t -\frac{k}{2}\mathrm{d}t' + \int_0^t -\frac{k}{2}\sin\left(\frac{2\pi t'}{\tau}\right)\mathrm{d}t'. \quad [77]$$

Integrating both sides, we find

$$\ln\left(\frac{u(t)}{u(0)}\right) = -\frac{k}{2}t + \frac{k\tau}{4\pi}\left(\cos\left(\frac{2\pi t'}{\tau}\right) - 1\right). \quad [78]$$

Finally, exponentiating both sides, we arrive at the time dependent concentration given by

$$u(t) = u(0)\exp\left(-\frac{k}{2}t + \frac{k\tau}{4\pi}\left(\cos\left(\frac{2\pi t'}{\tau}\right) - 1\right)\right). \quad [79]$$

We plot this result as the black lines in Figure S51.

We can compare the numerical solution to the ODE in Equation 76 using the forward Euler method,

$$u(t + \Delta t) = u(t) - \frac{k_b}{2}\Delta t\, u(t)\left(1 + \sin\left(\frac{2\pi t}{\tau}\right)\right). \quad [80]$$

Computing the numerical solution in python, we plot the result as the green circles in Figure S51(A). Both the numerical and analytic solutions are indistinguishable.

In general, we remark that this sinusoidally-varying bleaching rate is a special case of an integrating factor behavior, namely, Eq. 74 has the solution,

$$u(t) = u_0 \exp\left[-\int_0^{t'} dt' k(t')\right] \quad [81]$$

$$\equiv u_0 \exp\left[-\langle k(t)\rangle t\right], \quad [82]$$

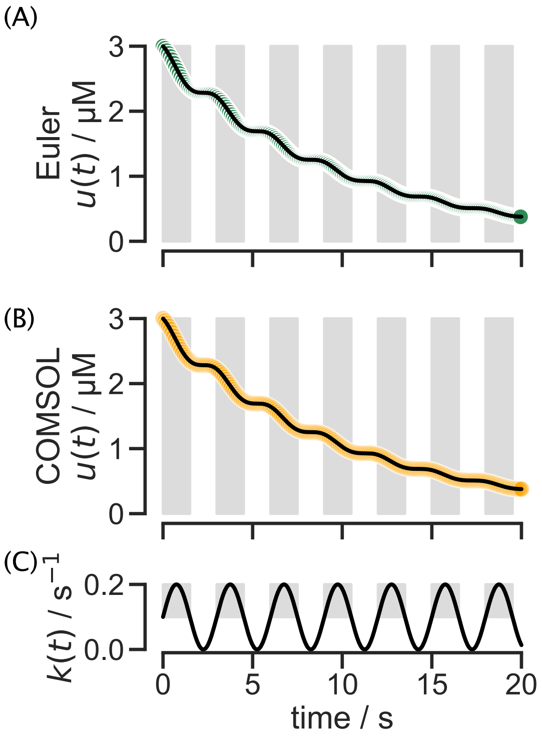

**Fig. S51. Numerically solving the intensity decay due to photobleaching when pulsing light matches the analytic solution for both the forward Euler method and FEM.** (A) The numerical solution, via the forward Euler method, for Equation 76 is plotted by the green circles under the analytical solution represented with a black line. (B) The numerical solution, via the finite element method, for Equation 76 is plotted by the orange circles under the analytical solution represented with a black line. (C) The driving pulse of the form of Equation 75 is plotted with the gray shading highlighting the half periods of greater bleaching contributions.

where $u_0$ is the value of the intensity at time zero, and in the second line we defined $\langle k(t)\rangle$ as a time averaged bleaching rate, $\langle k(t)\rangle \equiv \frac{1}{t}\int_0^t dt' k(t')$ up to time $t$. This behavior makes explicit that the observed intensity decay due to some time-varying photobleaching rate up to a time $t$ can be understood as effectively an exponential decay with a time-averaged bleaching rate. Thus, the overall effect of illumination by pulses is largely to trace out a softened decay curve set by an effective duty cycle of the illumination over the experimental timescale.

**F. Finite element simulations of our photobleaching models.** Having understood how a first-order kinetic description of bleaching balanced with reservoir replenishment (sections D.1 and D.2) and time-varying bleaching rates (section E ) manifest effectively plateauing intensity trajectories, we now graduate our modeling to a full description that explicitly accounts for the depletion and transport of fluorescent probes across space. We move from the ordinary differential equations of sections D.1 and D.2 to a partial differential equation treatment of the full reaction diffusion equations underlying our experimental conditions.

To develop both this full theory of our experiment and develop an understanding for the continuum mechanics of our system, we harness the power of the finite element method (FEM) already introduced in section B. To validate our simulations, in the next section F.1, we perform benchmarking checks with analytic solutions to the pulsatile dynamical context discussed in section E. Afterwards, in section F.2, we thoroughly explore the effects of modulating the diffusion constant and bleach rate on the time dynamics of intensity depletion.

***F.1. Validating finite element numerical conditioning with pulsing model benchmark.*** Before examining the full three-dimensional complexity of our problem, we first do numerical "controls" by comparing analytic and FEM solutions to the same simplified problem discussed in section E. Importantly, this verifies that these simulations are numerically well conditioned and behaved for experimentally relevant conditions.

In COMSOL, we defined a cylindrical geometry in which we applied Equation 76. We specify a uniform concentration initial condition and impose zero flux on the boundaries. Using the time dependent solver, we specify the start time, stop time and step size with the relative tolerance set to match the step size. We plot the results of the simulation, averaged over the cylinder, in Figure S51(B) as the orange circles. Again we find that the numerical solution using FEM is indistinguishable from the analytical solution.

Adding a layer of complexity, we now include diffusion in our COMSOL model and ensure that our simulations accurately describe the pulsing nature of our experiment. We now add a larger cylinder to the geometry which contains the reservoir of ATP probes but is never excited by light. Probes from this larger cylinder can diffuse into the smaller cylinder where light excitation can bleach the probes. We simulate two limits for which we already know the solutions, infinitely long and short times between pulses. When the interval between pulses is infinitely large, we would expect that as soon as the light

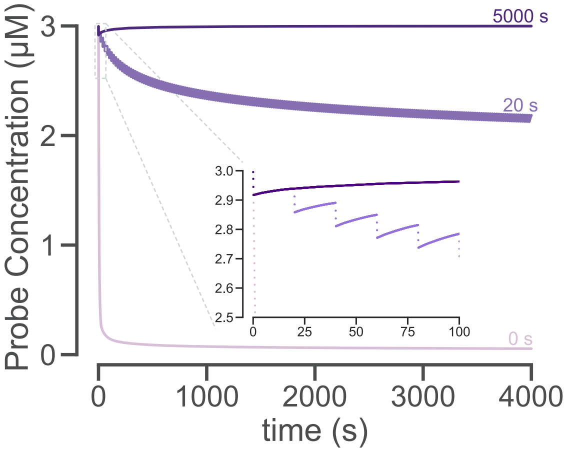

**Fig. S52. Examining the limits of interval times, we find simulations agree with expectations.** We simulate the depletion of bright probe concentration due to photobleaching with a diffusion constant of $40\,\mu\mathrm{m}^2/\mathrm{s}$ and a bleaching rate of $0.02\,\mathrm{s}^{-1}$ with three different intervals of time between pulses. In the limit of having an infinite time between pulses, or just a single pulse, we set the interval to be longer than the simulation time. The resultant curve (the darkest purple) initially undergoes decay with the other two curves, but after 20 seconds, recovers and plateaus back at the starting concentration. The middle trace plots the depletion when the light pulses every 20 seconds. The zoomed in inset of early times show a drop in concentration during each pulse and recovery for the interval between pulses. The second limit, having no time between light pulses (the lightest purple), shows monotonic depletion of bright probe concentration where a steady photobleaching smoothly depletes the bright probes regardless of their ability to diffuse.

turns off, we see recovery in the concentration of bright probes that remains constant for the rest of time. This is a FRAP experiment. The darkest line in Figure S52 shows just this. After one pulse which ends at 20 seconds, the concentration smoothly increases and then plateaus at about the starting concentration. In the opposite limit, when there is no interval between pulses, meaning the light is continuously on, the concentration of bright probes decays monotonically with no recovery, as seen by the lightest curve in Figure S52. Our experiment exists in an intermediate range, where light is pulsed every 20 seconds. The medium purple curve depicts this in Figure S52. We can see periodic intensity depletion and then recovery in the time between depletion. This is apparent in the zoomed-in inset.

***F.2. Finite element simulations of photobleaching and transport.*** We now use the finite element analysis to more precisely understand the dynamics of photobleaching in our experiments, acknowledging all relevant experimental effects. In particular, our experimental setup demands a "bleach-diffusion" equation treatment since the cover slip forms a "reservoir" of fluorescent molecules that can diffuse into the bleaching region. As a result, we have two competing time scales, namely, that of the photobleaching itself and the time scale over which diffusion of molecules from the reservoir can replenish the photobleached region. In the simulations that follow, we examine the dynamics that results from tuning these two time scales independently.

We set up a COMSOL simulation with the geometric constraints of our experiment. We model the excitation region as a small cylinder, with the depth of the chamber and diameter of the light pulse, inside a larger cylinder, with a diameter of the width of the chamber, see Figure S53. The use of the cylindrical geometry is simply a matter of convenience for the calculations since it makes it easier to set up and visualize when the problem has this symmetry.

This simulation explores the change of concentration $u(\mathbf{r}, t)$ of bright probes through space and time when there is no aster formation. We create a reaction containing microtubules, ATP, the ATP probe, but no motor proteins to prevent ATP hydrolysis and any structure formation. In this scenario, probes should be allowed to freely diffuse everywhere. Probes in the excitation region are subject to bleaching at a constant rate. Hence, in our simulations of this problem, we define two coefficient form partial differential equations, one to act in each region of the geometry. In the central region, there is both bleaching and diffusion, while in the exterior region, there is simply diffusion. The COMSOL form of "coefficient form partial differential equations" is

$$e_a \frac{\partial^2 u}{\partial t^2} + d_a \frac{\partial u}{\partial t} + \nabla \cdot (-c\nabla u - \alpha u + \gamma) + \beta \cdot \nabla u + au = f. \qquad [83]$$

Inside the excitation region, probes can diffuse and undergo bleaching. Thus, we set $d_a = 1$, $c = D$, $a = -k_{\mathrm{bleach}}$, and all other constants to zero such that

$$\frac{\partial u}{\partial t} - D\nabla^2 u + k_{\mathrm{bleach}} = 0. \qquad [84]$$

Outside the excitation region, probes cannot be bleached, so as a second coefficient form PDE, we set $d_a = 1$ and $c = D$, with all other constants set to zero, such that

$$\frac{\partial u}{\partial t} - D\nabla^2 u = 0. \qquad [85]$$

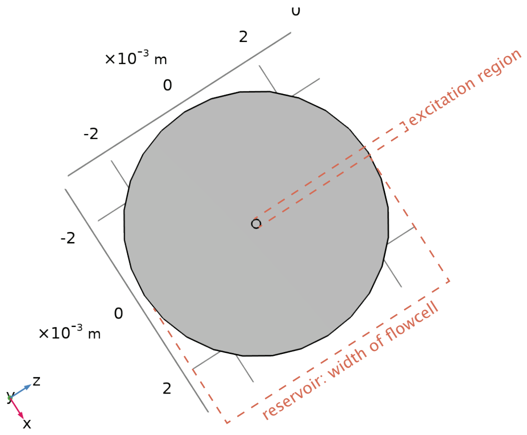

**Fig. S53. Aerial view of cylindrical geometry in COMSOL for photobleaching simulations.** We model photobleaching experiments with two concentric cylinders. The larger has a diameter of the width of the flow cell where the sample is deposited. This larger cylinder acts as a reservoir of ATP probes. The much smaller cylinder is the region where we pulse light excitations. It is inside this region where bleachign occurs.

We set the boundary conditions of the outer cylinder to have zero flux. At time zero, we fix the concentration across the geometry to a constant concentration, $c_0$.

When setting up our simulation, there are further technical specifications needed to fully describe the physical model. For those interested in precisely repeating our numerical simulations, we report the specific details of our COMSOL implementation of this problem in the following box.

*Specifics of Simulation Set-up:*

- To capture the discrete on/off nature of pulsing the excitation light, we utilize the Events Interface in COMSOL. Events trigger the solver to reassess the step size at specified times.
  - There are two event types, explicit and implicit, available in COMSOL. Here we use two explicit events, since we know the precise timings of the light stage. Implicit events should be used when the event of interest is a response to the simulation dynamics.
  - We create a discrete states variable called ONOFF, which we initially set to a value of 1. This corresponds to the light being on. We then create an explicit event, which turns the light off such that ONOFF $= 0$, which is set at time $t_{\text{excite}} = 150$ ms and repeats with an interval $t_{\text{int}} = 20\,$s. A second explicit event is set to turn the light back on at $t_{\text{int}} = 20\,$s, by setting ONOFF $= 1$, and repeats with the interval $t_{\text{int}} = 20\,$s.
- We use the Coefficient Form PDE physics to describe our model. We define two separate PDEs to simulate the bright probe concentrations in each cylinder.
  - Our first PDE is defined to act upon the smaller cylinder. In this region, probes can diffuse and also be bleached by the light pulses thus, our model is

$$\frac{\partial c}{\partial t} = D\nabla^2 c - k_{\text{bleach}} \times \text{ONOFF}. \quad [86]$$

  - The second PDE acts upon the larger cylinder excluding the inner smaller cylinder's volume. Here probes can only diffuse, thus we use the model

$$\frac{\partial c}{\partial t} = D\nabla^2 c. \quad [87]$$

- We use a physics controlled mesh with a “Fine" element size.
- To solve the equation, we use the Time Dependent Solver with output times from $0 - 4000\,$s at a step size of 0.05 s. We apply a user controlled relative tolerance of 0.01. For the absolute tolerance, we use a "Scaled" global method with the "Factor" tolerance method set to a tolerance factor of 0.1. We leave the derivative tolerance method to be "Automatic".

One goal of this simulation is to understand how the parameter values for $k_{\text{bleach}}$ and $D$ change the time dynamics of photobleaching. We run a variety of simulations sweeping the parameter values of $k_{\text{bleach}}$ and $D$ as shown in Figures S54 and S55. We find that as we increase $k_{\text{bleach}}$ and as we increase $D$, the steepness of the decay decreases and there is a higher concentration of bright probes at the end of the simulation. We fit the simulation curves to a single exponential plus a constant as seen in Figure S54. The fits are described by the equations

$$\frac{u}{u_0} = (1 - I_\infty)\exp(-t/\tau_1) + I_\infty, \quad [88]$$

in Figure S54 and a double exponential in Figure S55 which is described by

$$\frac{u}{u_0} = (1 - I_\infty)\exp(-t/\tau_1) + I_\infty \exp(-t/\tau_2), \quad [89]$$

where $u_0$ is the initial concentration of bright probes. We find a better agreement to the double exponential fit. Regardless of the fit type, we find the fit parameters for the fast decay constant, $\tau_1$, and the infinite time value of the concentration, $I_\infty$, reflect the trends in the bleaching rate and diffusion constants. The increasing steepness that results from increasing $k_{\mathrm{bleach}}$ and $D$ is reflected by a decreasing $\tau_1$. Similarly the higher ending concentration of bright probes that correlates with higher $k_{\mathrm{bleach}}$ and $D$ values is reflected by an increasing $I_\infty$ value.

Complementing our exploration of the form of the decay in probe brightness with time, we can take a more granular look into the spatial depletion profile as shown in Figures S56 and S57. Here, we draw a straight line the length of the larger cylinder diameter which passes through the center point of the geometry. Plotting the bright probe concentration as a function of position on the line, we can observe the form of depletion throughout the simulation. We find that as the diffusion constant increases, the well created by light pulsing in the inner circle becomes shallower. This makes sense because a higher diffusion constant allows bright probes to quickly replenish the darkened well. As the bleaching rate increases, the well becomes deeper. With more probes bleached per shot of light, the deficit of bright probes in the well is more significant. The simulations with zero diffusion, $D = 0$ show some noise due to numerical instability, especially at the discontinuous edges of the region of light excitation. These plots may raise concern that when treating images for photobleaching, there may need to be a spatial dependence on the corrective function. However, we find in the regions of the images we use, the spatial effects are minimal. Figure S57 zooms in on the light excitation region and also shows the width of the approximate final aster size. At early times, when the aster is first contracting and is about the size of the light excitation region, the concentration profile is mostly flat with little spatial dependence. At later times when there more curvature in the well, we highlight the percent change in bright probe concentration in the aster size region is maximally one percent. Thus, we do not believe applying a spatially-explicit correction to photobleaching is necessary for these experiments.

#### 7. Investigation of image geometry

**A. Conceivable geometric and optical distortions arising from 2D images of 3D distributions.** Monitoring the rapidly changing, and optically labile, chemical reactions driving aster formation with microscopy imposes fundamental technical constraints on our measurements. Specifically, while asters are formed by molecular motors, microtubules, and ATP arranged in rich three-dimensional structures, imaging these distributions fully in three dimensions over our conditions of biochemical interest would significantly constrain the fastest changes resolvable in these spatial fields. In addition, acquiring 3D data (via many tightly-spaced 2D images in an axial dimension) imposes a significantly larger burden of excitatory photons that can photobleach and further perturb fluorescently-labeled molecules like those we monitor. Consequently, to capture fine changes in aster structure over space and time, the bulk of our measurements are in the form of two-dimensionsal epifluorescence images of developing asters. These images are collected while focused at a central axial plane through the aster.

To interpret this slew of experimental results, in principle, we must account for the difference between our 3D objects of interest and the 2D epifluorescence microscopy images representations of these structures. Here, in the sections that follow, we assess the possible consequences/distortions following from these two-dimensional acquisitions, both conceptually in general, and empirically for our specific imaging conditions. To augur what follows, our analyses largely show that the conceptually-possible differences between two-dimensional data and three-dimensional distributions are in fact empirically negligible, largely due to the narrowness of the optical point spread functions convolving these images compared to the shape of the raw underlying structures we image.

Our discussion that follows proceeds in three parts. First, we consider the ideal case where the optical weighting function along the axial dimension is zero everywhere except the focal plane actually sampled, meaning the acquired 2D image precisely matches the equivalent 3D slice (with no distortion). Second we consider the opposing case where the weighting function is uniform along the axial dimension, resulting in a uniform projection when integrating over each slice in the axial dimension. Lastly, we consider the most realistic case falling between these two limits, where a nonzero, nonuniform, weighting function accumulates fluorescence signal from slices above and below the focal plane, giving a convolution with the signal at the focal plane. In imaging practice, we numerically verify that the narrowness of this weighting function (relative to the objects of interest) makes this case closely approximate the first case of a bona fide slice through the underlying three dimensional distribution.

**B. General form of a 2D image projection.** In general, a two dimensional image $\hat{I}(x, y)$ collected at spatial position $(x, y)$ is an integral of a true underlying source profile $I(x', y', z')$ from sources at positions $(x', y', z')$, where each source is weighted by some optical weighting function $f(x', y', z')$ throughout space,

$$\hat{I}(x, y) = \int_{-\infty}^{\infty}\int_{-\infty}^{\infty}\int_{-\infty}^{\infty} I(x', y', z')\; f(x', y', z')\,\mathrm{d}x'\mathrm{d}y'\mathrm{d}z'. \quad [90]$$

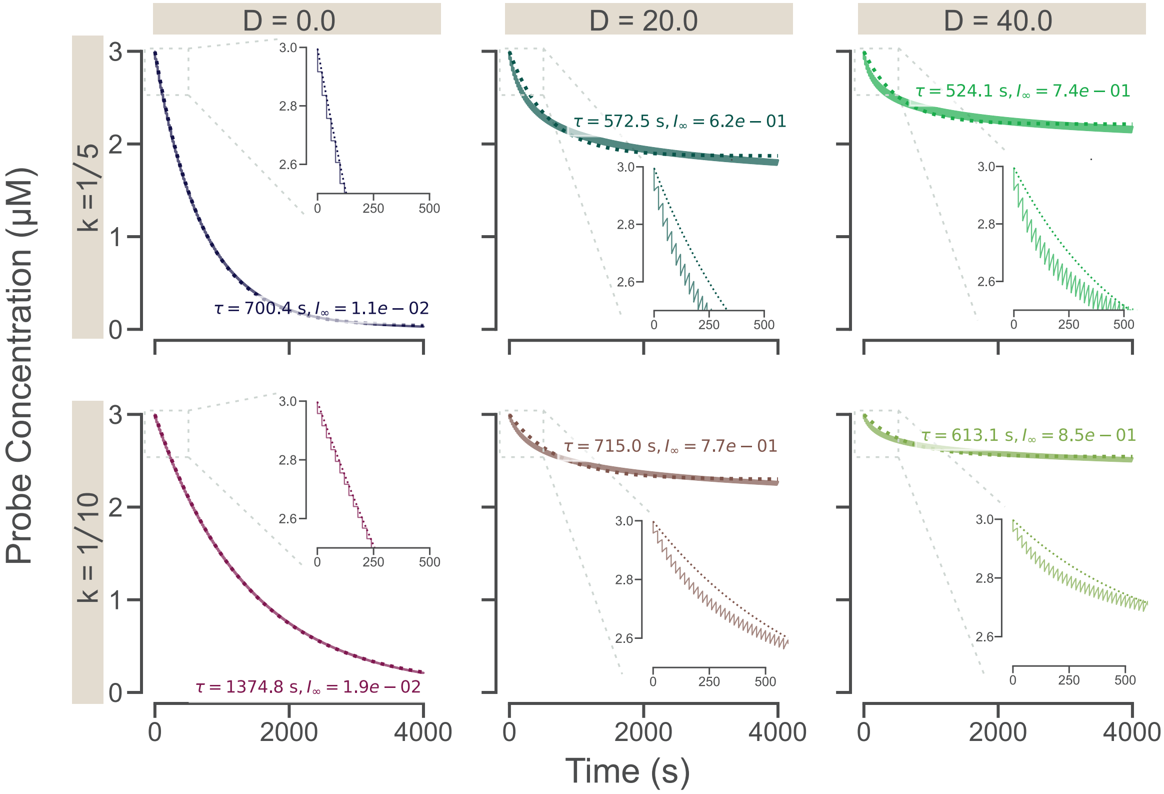

**Fig. S54. Fitting COMSOL simulations of the bright ATP probe intensity to a single exponential plus a constant.** We simulate the depletion of bright probes due to photobleaching for a sweep of bleaching rates and diffusion constants. Fitting the time traces, which report the concentration of bright probes averaged over the light excitation region, we find a single exponential plus a constant, equation 88, fits the traces moderately well. We witness trends in the fit parameters such that the decay constant $\tau$ increases with increasing $k_{\text{bleach}}$ and $D$, while $I_\infty$ increases with increasing $D$ but decreases with increasing $k_{\text{bleach}}$. The insets for each plot show a zoomed in blow up of the early times to visualize the initial depletion and recovery cycles corresponding to the pulses. The single exponential plus constant fits don't capture the initial decay well for non-zero diffusion.

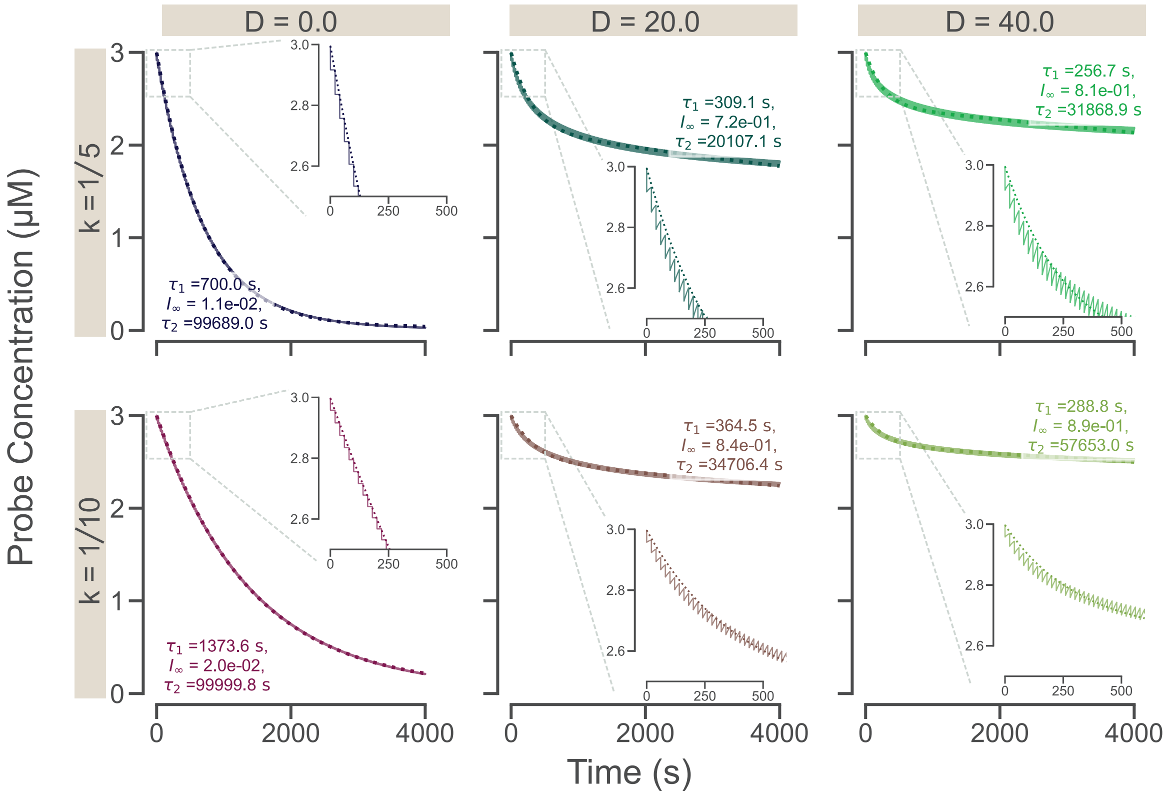

**Fig. S55. Fitting COMSOL simulations of the ATP probe intensity decay due to photobleaching to a double exponential.** We simulate the depletion of bright probes due to photobleaching for a sweep of bleaching rates and diffusion constants. Fitting the time traces, which report the concentration of bright probes averaged over the light excitation region, we find a double exponential plus a constant, equation 89, fits the traces quite well, and better than the single exponential plus a constant. We witness trends in the fit parameters such that the decay constant $\tau_1$ increases with increasing $k_{\text{bleach}}$ and $D$, while $I_\infty$ increases with increasing $D$ but decreases with increasing $k_{\text{bleach}}$. The insets for each plot show a zoomed in blow up of the early times to visualize the initial depletion and recovery cycles corresponding to the pulses. The double exponential fits well capture the initial decay.

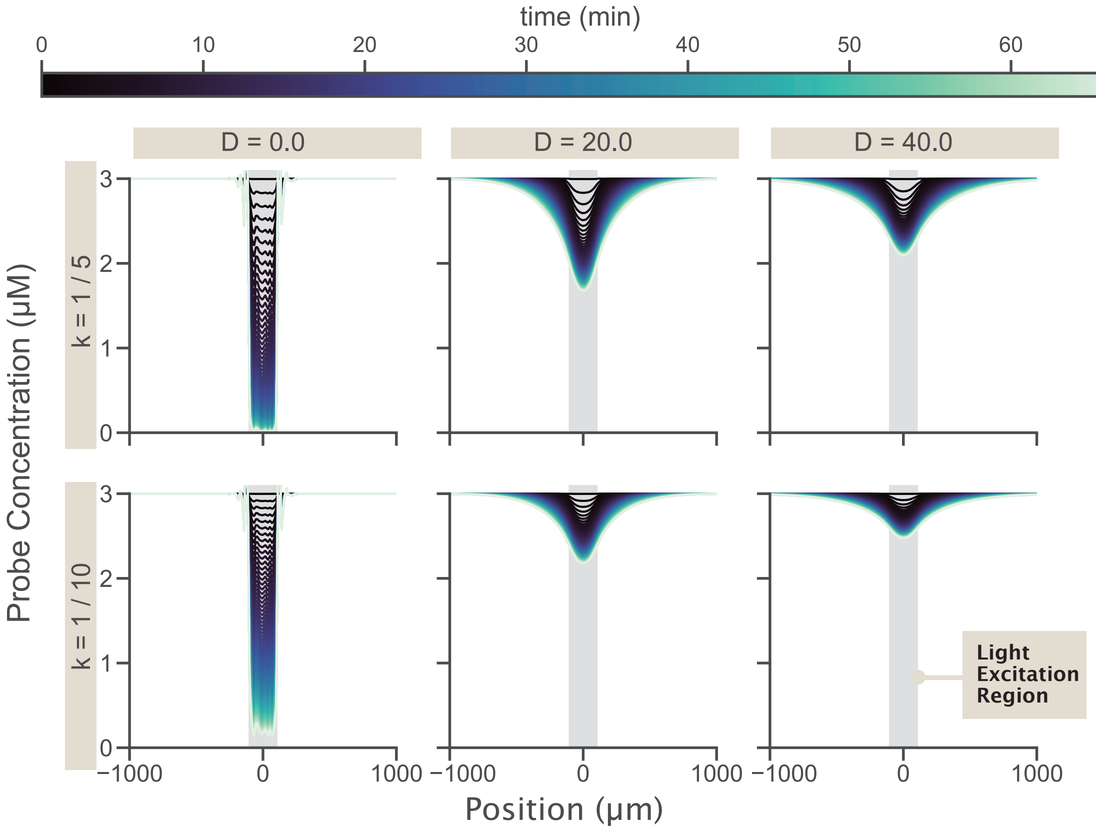

**Fig. S56. Plotting the bright probe concentration across space shows how simulation parameters impact the depletion well formed.** We draw a cut line that spans the diameter of the larger cylinder at the midpoint of the cylinder height. Each line in the figure represents the concentration of bright probes in space for a given timepoint. We find the depth of the well depleted correlates with increasing bleaching rates and decreasing diffusion constants.

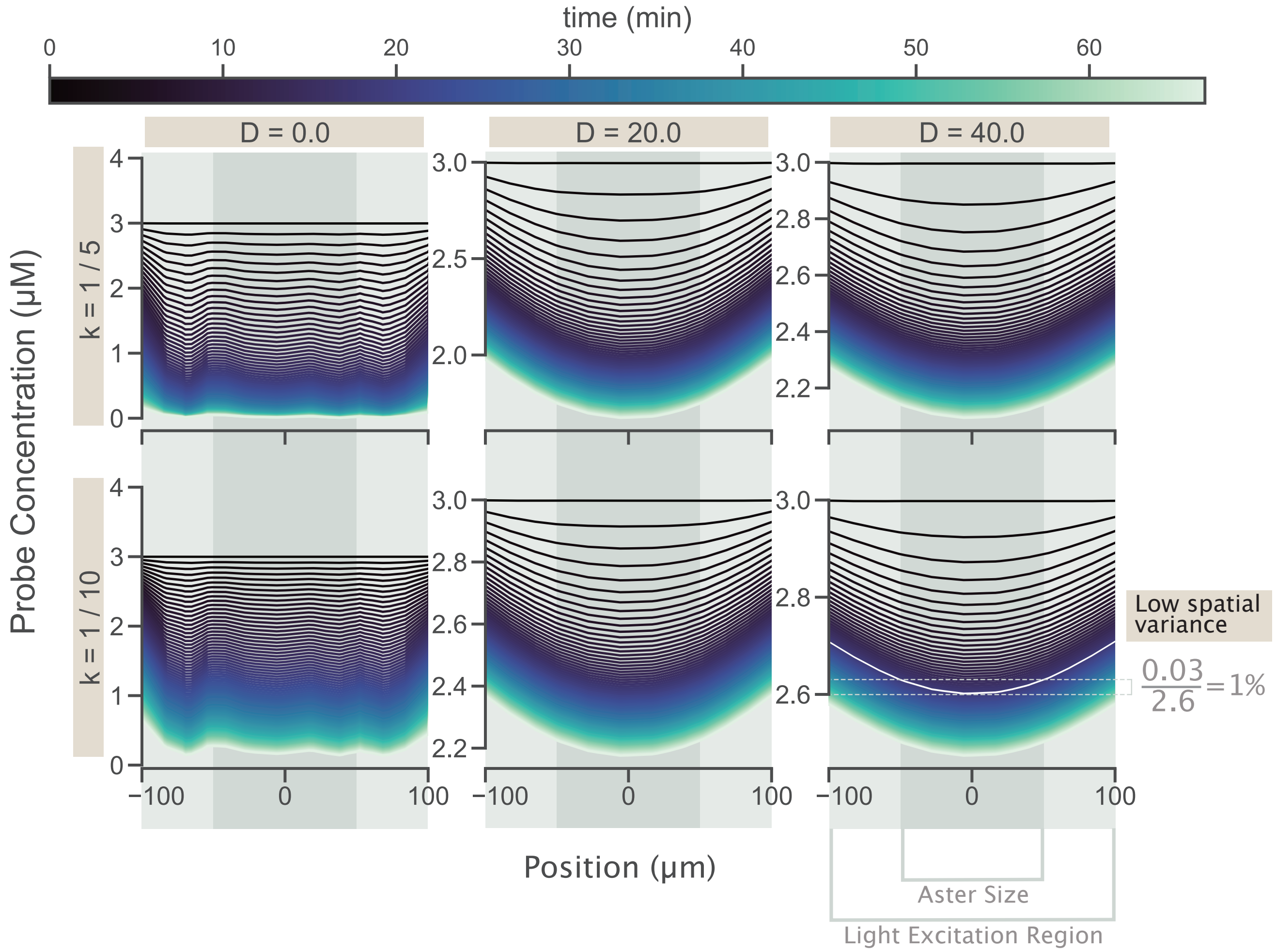

**Fig. S57. Zooming in on the spatial depletion well created by photobleaching, there is little variation within the region of interest.** If there is high levels of variation throughout space in concentration of bright probe, we would need to correct for photobleaching with a spatial dependent function. Zooming in on the light excitation region, where our analysis occurs, we find that the extent of spatial variation is small. At early times (blacker lines), the aster is approximately the size of light excitation region, but the concentration traces are mostly flat, implying little variation. For later times (bluer lines), the traces are curvier. But during these times the aster is smaller, around the width of the dark grey region. In the bottom right plot, we show for a given time around 20 minutes (highlighted by the white trace), the variation is still small, only one percent in the aster region.

***B.1. Specific imaging cases.*** In a first ideal case, when the weighting function is simply a Dirac delta function over the axial dimension centered at the focal plane at $z$, namely $f(x',y',z') = \delta(x-x')\delta(y-y')\delta(z-z')$, the image $\hat{I}^{(1)}(x,y)$ is exactly the corresponding focal slice through the three dimensional profile,

$$\hat{I}^{(1)}(x,y) = \int_{-\infty}^{\infty}\int_{-\infty}^{\infty}\int_{-\infty}^{\infty} I(x',y',z')\ \delta(x-x')\delta(y-y')\delta(z-z')\,\mathrm{d}x'\mathrm{d}y'\mathrm{d}z' \quad [91]$$

$$= I(x,y,z). \quad [92]$$

In a second opposing case, the image accumulates signal uniformly over sources along the axial dimension, namely $f(x',y',z') = \delta(x-x')\delta(y-y')\times 1$. This gives an image $I^{(2)}(x,y)$ of the form,

$$\hat{I}^{(2)}(x,y) = \int_{-\infty}^{\infty}\int_{-\infty}^{\infty}\int_{-\infty}^{\infty} I(x',y',z')\ \delta(x-x')\delta(y-y')\,\mathrm{d}x'\mathrm{d}y'\mathrm{d}z' \quad [93]$$

$$= \int_{-\infty}^{\infty} \mathrm{d}z'\ I(x,y,z). \quad [94]$$

When the underlying distributions $I(x,y,z)$ are axially- symmetric or spherically-symmetric ( $\equiv I(\rho,\theta)$ or $\equiv I(r)$), such images two dimensional images $\hat{I}$ adopt the forms of Abel transformations (23). In principle, the underlying $I(r)$ or $I(\rho,\theta)$ can be recovered by an inverse Abel transformation (23, 24) on the measured two-dimensional image $\hat{I}$, though in practice, the numerical success of these schemes is a delicate and often fraught inverse problem.

In a third, intermediate, and most realistic case, we acknowledge that the sources from above and below the focal plane at $z$ can contribute with nonzero weight to the measured signal at $z$, and also further acknowledge the possible contributions of sources at transverse positions $(x',y')$ distinct from the query position $(x,y)$. Specifically, the optical weighting function of a source at $(x',y',z')$ is a point-spread function (set by the microscope) well modeled as depending only on the displacement $(x-x', y-y', z-z')$ to the query point $(x,y)$. This gives an image $\hat{I}^{(3)}(x,y)$ of the form,

$$\hat{I}^{(3)}(x,y) = \int_{-\infty}^{\infty}\int_{-\infty}^{\infty}\int_{-\infty}^{\infty} I(x',y',z')f(x-x',y-y',z-z')\,\mathrm{d}x'\mathrm{d}y'\mathrm{d}z'. \quad [95]$$

Models of the point spread function $f(x-x',y-y',z-z')$ corresponding to circular apertures appropriate for microscope objectives are available (25–27) to varying levels of analytical and numerical detail, including the popular Born and Wolf model expressed in terms of Airy and Bessel functions. Often a very good approximation to these empirical circumstances is a Gaussian decay factorizable in transverse and axial dimensions with anisotropic widths, namely,

$$f(x-x',y-y',z-z') = \mathrm{psf}_{\mathrm{tr}}(x-x',y-y'|\sigma_{\mathrm{tr}}^2)\times \mathrm{psf}_{\mathrm{ax}}(z-z'|\sigma_{\mathrm{ax}}^2). \quad [96]$$

**C. Case Two: Abel Transformation.** When the optical weighting function is not purely localized at the focal plane, but instead is uniform throughout the axis of the imaging volume, the distortion of a three-dimensional profile is conceivably appreciable. Image sources at axial positions far away from the focal plane are convolved with sources actually residing at the focal plane, in principle posing an intractable ill-posed inverse problem: there are many different underlying 3D distributions consistent with a given 2D measured image. However, intriguingly, when the underlying distribution $I(x,y,z)$ is spherically-symmetric or axially-symmetric, a measured two-dimensional profile $\hat{I}(\rho,\theta)$ uniquely specifies the original three-dimensional profile, a relationship expressed by the so called Abel transformation and its complement, the inverse Abel transformation. Here, we explicitly describe how a radially-symmetric image (e.g. of spherically-symmetric aster) manifests a two dimensional projection forming an Abel transformation, under this pessimistic scenario when the optical weighting function does not decay away from the focal plane as rapidly as expected from epifluorescence microscopy. We also remark on qualitative features that can be analytically shown to survive in such a two dimensional uniformly-weighted projection. We provide this discussion to give maximally conservative statements about how two dimensional images still report on key features of three dimensional structures, even in the counterfactual setting where distant image sources are weighted equally to those appearing on the focal plane.

Imagining a 3D, spherical aster where we possess an omniscient knowledge of the concentration profile, we ask what will be the concentration profile reported by a microscope 2D image? To solve, we integrate the height, over z (in cylindrical coordinates), of a skyscraper cutting through the aster sphere. For a given skyscraper address with cylindrical coordinates $(\rho,\theta)$, we calculate the concentration of the image pixel value corresponding to the skyscraper,

$$\hat{I}(\rho,\theta) = \int_0^h dz\, I(\rho,\theta,z). \quad [97]$$

Since there is radial symmetry in $\theta$, we can remove the $\theta$ dependence in the integral such that,

$$\hat{I}(\rho) = \int_0^{2\pi} d\theta \int_0^h dz\, I(\rho,z) = 2\pi\int_0^h dz\, I(\rho,z). \quad [98]$$

We now change to spherical coordinates, rather than cylindrical, since we consider the aster to be a sphere. Thus, the concentration as a function of $r$, which depends on $\rho$ and $z$, can be written as,

$$\hat{I}(\rho) = 2\pi \int_0^h dz\, I(r(\rho, z)). \quad [99]$$

Since for a given skyscraper $\rho$ is constant, we can change the integration variable from $z$ to $r$ with the Equation,

$$r^2 = \rho^2 + z^2. \quad [100]$$

Differentiating with respect to $r$,

$$2r = 2z\frac{dz}{dr} \Rightarrow dz = \frac{r}{z}dr. \quad [101]$$

Using the substitution, $z = \sqrt{r^2 - \rho^2}$,

$$dz = \frac{r}{\sqrt{r^2 - \rho^2}}dr. \quad [102]$$

Thus, the concentration at a given 2D image radius, $\rho$ of a 3D aster is,

$$\hat{I}(\rho) = 2\pi \int_\rho^{\sqrt{\rho^2+h^2}} dr\, I(r)\frac{r}{\sqrt{r^2 - \rho^2}}. \quad [103]$$

As a sanity check, we can ensure that a uniform concentration in 3D, $I(r, \theta, \phi) = I_0$ is still uniform in 2D. Plugging this constant into Equation 103,

$$\hat{I}(\rho) = 2\pi I_0 \int_\rho^{\sqrt{\rho^2+h^2}} dr\, \frac{r}{\sqrt{r^2 - \rho^2}}, \quad [104]$$

and performing a u-substitution where $u = \sqrt{r^2 - \rho^2}$ and $du = r/\sqrt{r^2 - \rho^2}$,

$$\hat{I}(\rho) = 2\pi I_0 \int_0^h du = 2\pi I_0 h. \quad [105]$$

We can ensure that if we simply integrated over a constant sample from the perspective of the microscope, we would obtain the same constant,

$$\hat{I}(\rho, \theta) = \int_0^h dz\, I_0 = I_0 h, \quad [106]$$

and assuming spherical symmetry,

$$\hat{I}(\rho) = \int_0^{2\pi} d\theta\, I_0 h = 2\pi I_0 h. \quad [107]$$

The transform defined in Equation 103, is called an Abel Transform, and has an analytical inverse function, under the condition that $I(r) \to 0$ faster than $\frac{1}{r}$ (24).

**D. Empirical deconvolution on an aster.** Here we acknowledge the contributions of intensity signals from nearby pixels on a queried pixel as directed by a finite width point spread function. The majority of data in this study analyzes 2D images that pass through the aster center. Figure S58 depicts an image of this type. In analyzing the intensity "smearing" that occurs from neighboring pixels, it is important to not only evaluate the contribution of other pixels in the same plane but also pixels in planes above and below the pixel of interest. In this section, we evaluate how the results of deconvolution on 3D image stacks of an aster modify the intensity value of the resultant image. We acquire images of an aster motor distribution every $0.5\,\mu$m in the axial direction. We then perform a deconvolution procedure using an ImageJ plugin, Deconvolution Lab2 (25), employing the Richardson-Lucy deconvolution method. As inputs, we provide an image or image stack and also a point spread function. For circular apertures, the point spread function has been shown to be an Airy function (26). However, for ease of use, practitioners often approximate these functions with a Gaussian. We demonstrate the results of deconvolving with both of these methods.

We generate an Airy PSF using an ImageJ PSF Generator plugin (28). The plugin requests as inputs the refractive index, which we set to 1 for an air objective, the wavelength of light, which we input as 480 nm, the numerical aperture of the objective, which we set to 0.45, and the nanometer to pixel conversions in each dimension, which for the $x$ and $y$ dimensions, we take to be 578 nm/px and for the z dimension we use 500 nm/px. In Figure S59 we show an example of the Airy PSF at the center plane.

Using Deconvolution Lab2, we perform 50 iterations of Richardson-Lucy deconvolution, at first only in 2D, on the sample image shown in Figure S58 and deconvolve with the Airy PSF shown in Figure S59. In Figure S60(A) we show the result of this deconvolution when implemented this way. In Figure S60(B) we plot the trace of a cut line through the center of the aster for the initial raw image in black and the Fiji deconvolved Airy PSF in blue. We find that this deconvolution returns a value practically identical to the raw image trace.

Complementary to using the Airy function, we explore the results of using a Gaussian as already described earlier in the context of synthetic data. In Figure S61 we fit the length parameters for the Gaussian to the Airy PSF generated in ImageJ and find that the transverse direction has a length scale of $\sigma_{\mathrm{tr}} = 0.33\,\mu$m and the axial dimension has a length scale of $\sigma_{\mathrm{ax}} = 2.32\ \mu$m. These characteristic widths, and the conceit of Gaussian approximations, to the point spread function are highly consistent with reported values and procedures in related literature; e.g. references (29, 30) report similar widths of $0.3 \sim \mu$m to the transverse character of experimental point spread functions under related conditions.

Upon deconvolving with the Gaussian point spread function, as shown in Figure S62, the output again appears nearly identical to the raw image as shown in Figure S58.

These tests of 2D deconvolutions give us a sense that any effect of convolution by the microscope is minimal. However, it is important to check modifications created by convolutions in the axial dimension, especially since this is the widest dimension of the point spread function, as can be seen in Figure S61. To this end, we perform 100 iterations of Richardson-Lucy deconvolution for both the generated Airy PSF and the fit Gaussian PSF. Again, we plot the cut lines through the center of the aster in Figure S64 and find little deviation from the raw image. The Gaussian deconvolution again looks identical to the raw image, but for the Airy deconvolution, there is a slight difference at the center of the aster. Here, the Airy deconvolved output produces higher intensities in the aster core and becomes less smooth.

Rather than taking a cut line, we also plot the polar averages of the images centered about the aster with respect to the aster radius shown in Figure S65. Once again, we largely see agreement with the raw images. The 3D airy deconvolution is again the only deconvolved output that show any difference with an increased intensity at the aster center.

Having learned that the quantitative impact of deconvolution in two or three dimensions in real aster images of interest is extremely numerically modest, we now explicitly verify that the ambient background level of fluorescence signal does not change the (small) impact of deconvolution. Figure S66 compares how performing background subtraction before or after deconvolution affects the recovered fluorescence profiles. The raw, deconvolved, and background-subtracted then deconvolved traces of Figure S66(A) agree up to the background offset, and very small numerical noise. This agreement, arguing that deconvolution and background subtraction largely commute numerically, shows that the presence of an ambient signal background in real aster images leaves the result of deconvolution essentially unchanged (up to exactly the background offset). We further visualize this agreement in Figure S66(B) by plotting the difference between the deconvolved image $D[I]$ and the background-subtracted then deconvolved image $D[I - \langle I_{\mathrm{bg}} \rangle]$, relative to the background level $\langle I_{\mathrm{bg}} \rangle$. Over the central line profile of the aster, this difference is very close to unity throughout the image, namely $\frac{D[I]-D[I-\langle I_{\mathrm{bg}}\rangle]}{\langle I_{\mathrm{bg}}\rangle} \approx 1$.

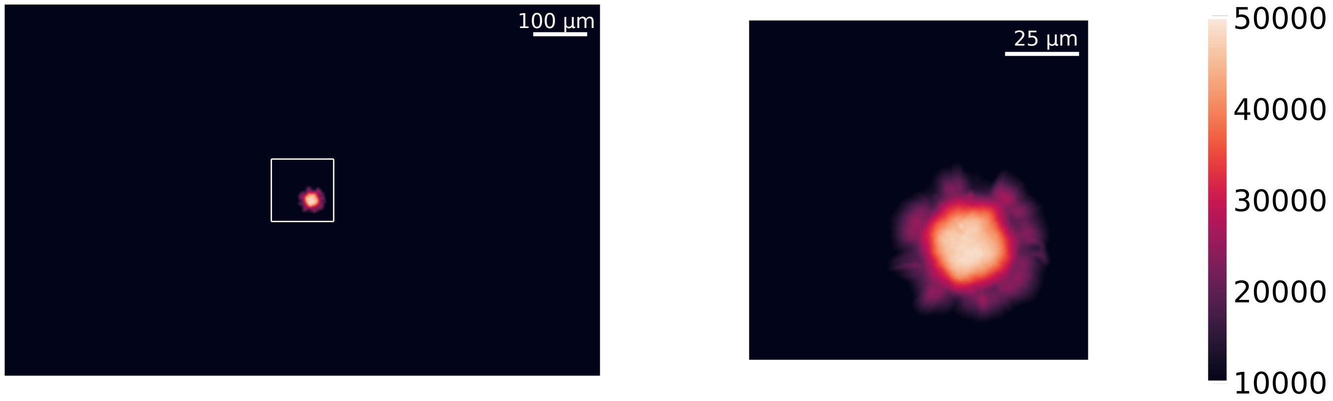

**Fig. S58. Image of the center of an aster for deconvolution.** As an example image for deconvolution, the center plane of a z-stack of a fully formed aster is shown. On the left, the full figure dimensions are shown, while on the right, a cropped view of the center of the image is shown, corresponding to the white square on the left. We note the aster is not perfectly centered within the image which is typical for asters as they dynamically form. Our deconvolution method needs to work regardles of the aster centering.

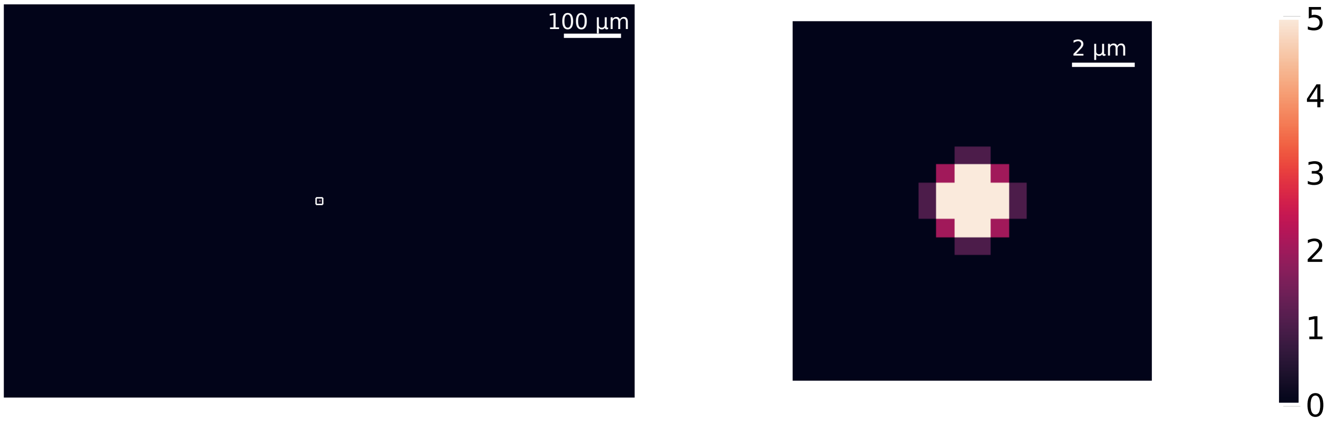

**Fig. S59. Generated Airy point spread function.** Using the PSF Generator in ImageJ with the Born and Wolf 3D model, we have created a point spread function based on our microscope's optical parameters. The images here are the center slice of the 3D PSF where the left shows the full size of the image and the right is cropped to the center to show the bright center of the PSF.

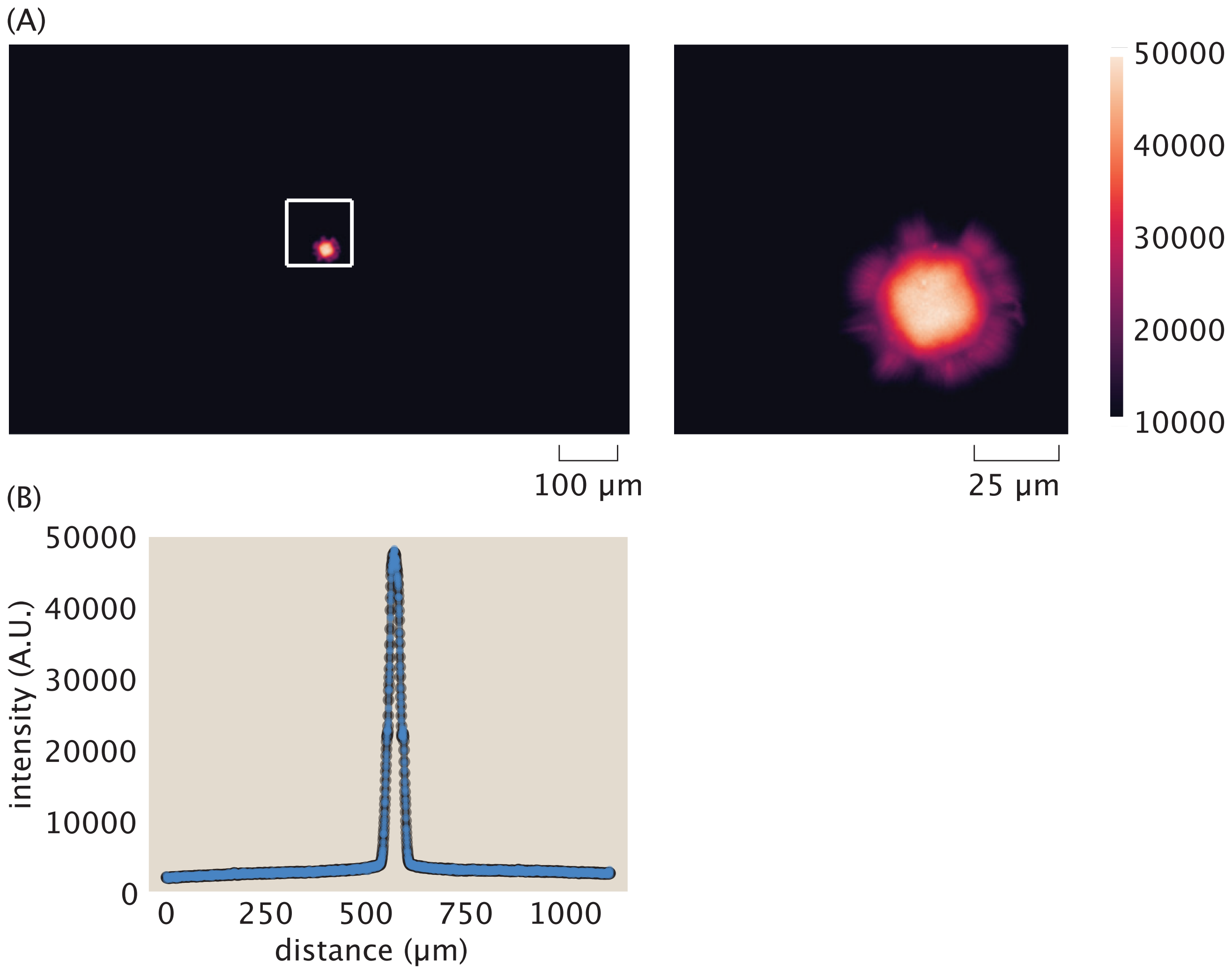

**Fig. S60. Deconvolving using an Airy PSF in ImageJ reveals a similar output to the raw image input.** (A) The full deconvolution output and cropped deconvolution output are depicted. (B) Taking a cut line through the center of the aster for both the raw image (in black) and the deconvolved output (in blue), we find near perfect agreement in intensity signal. This indicates that the convolution done by the microscope had very little effect on the image.

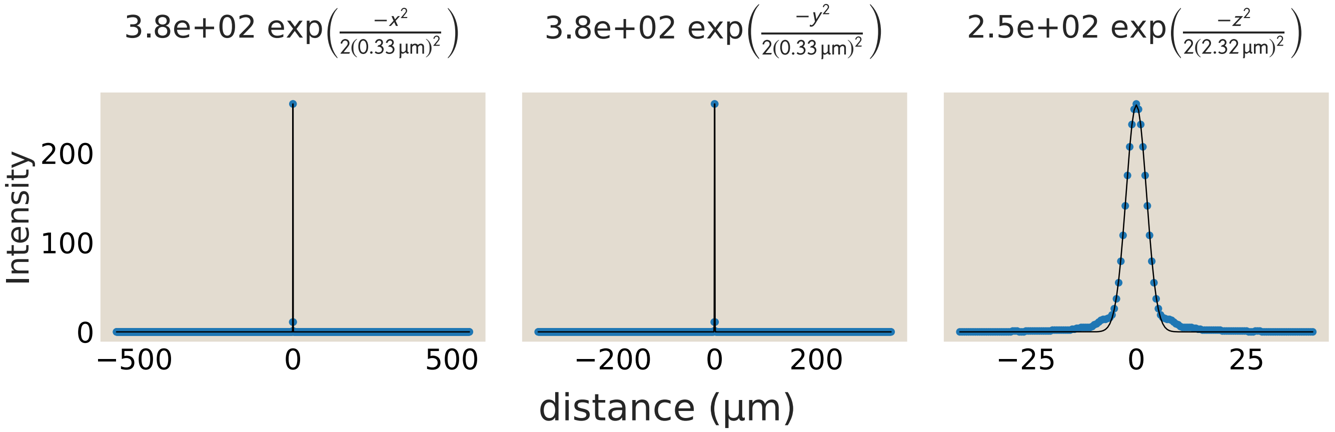

**Fig. S61. Gaussian length scales are determined by fitting the generated Airy point spread function.** Gaussian functions provide a very good fit to the decay of the point spread function in space. The transverse plane has a small length scale, only $0.33\ \mu m$, while the axial direction has a longer, but still small length scale of $2.32\ \mu m$.

 

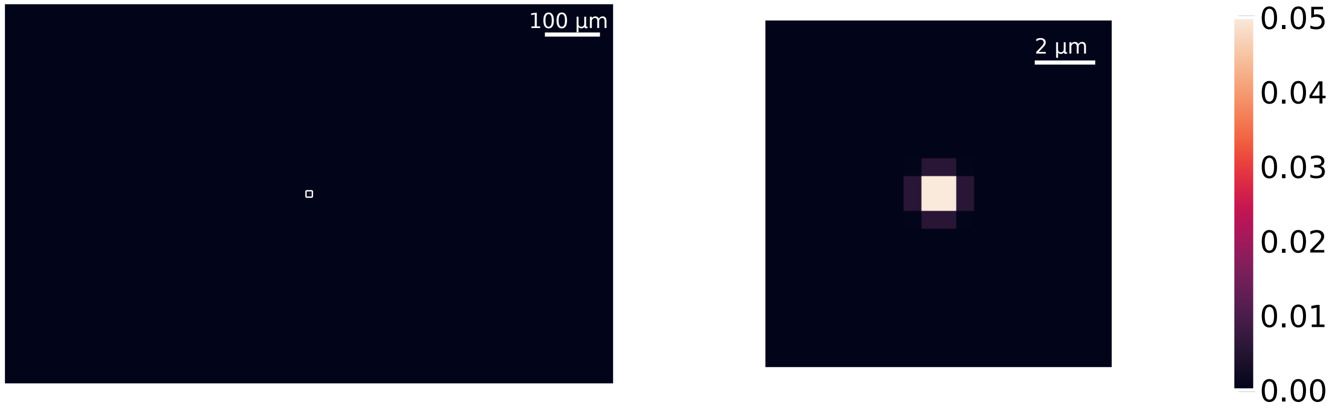

**Fig. S62. GaussPSF with length scales as determined by the Airy function fits.** .

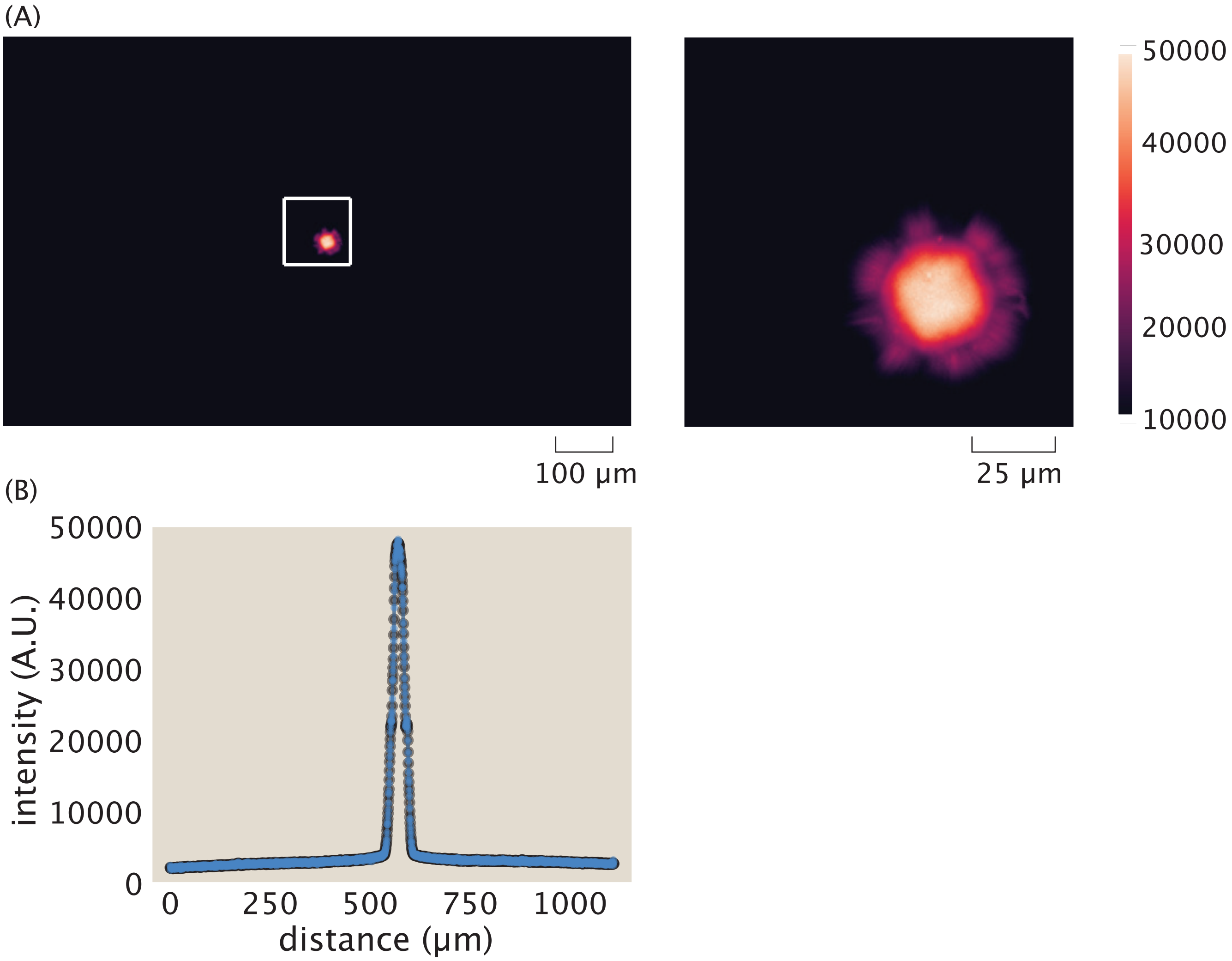

**Fig. S63. Deconvolving using a Gaussian PSF reveals a similar looking output to the raw image input.** (A) The full deconvolution output and cropped deconvolution output are depicted. (B) Taking a cut line through the center of the aster for both the raw image (in black) and the deconvolved output (in blue), we find near perfect agreement in intensity signal. This indicates that the convolution done by the microscope had very little effect on the image.

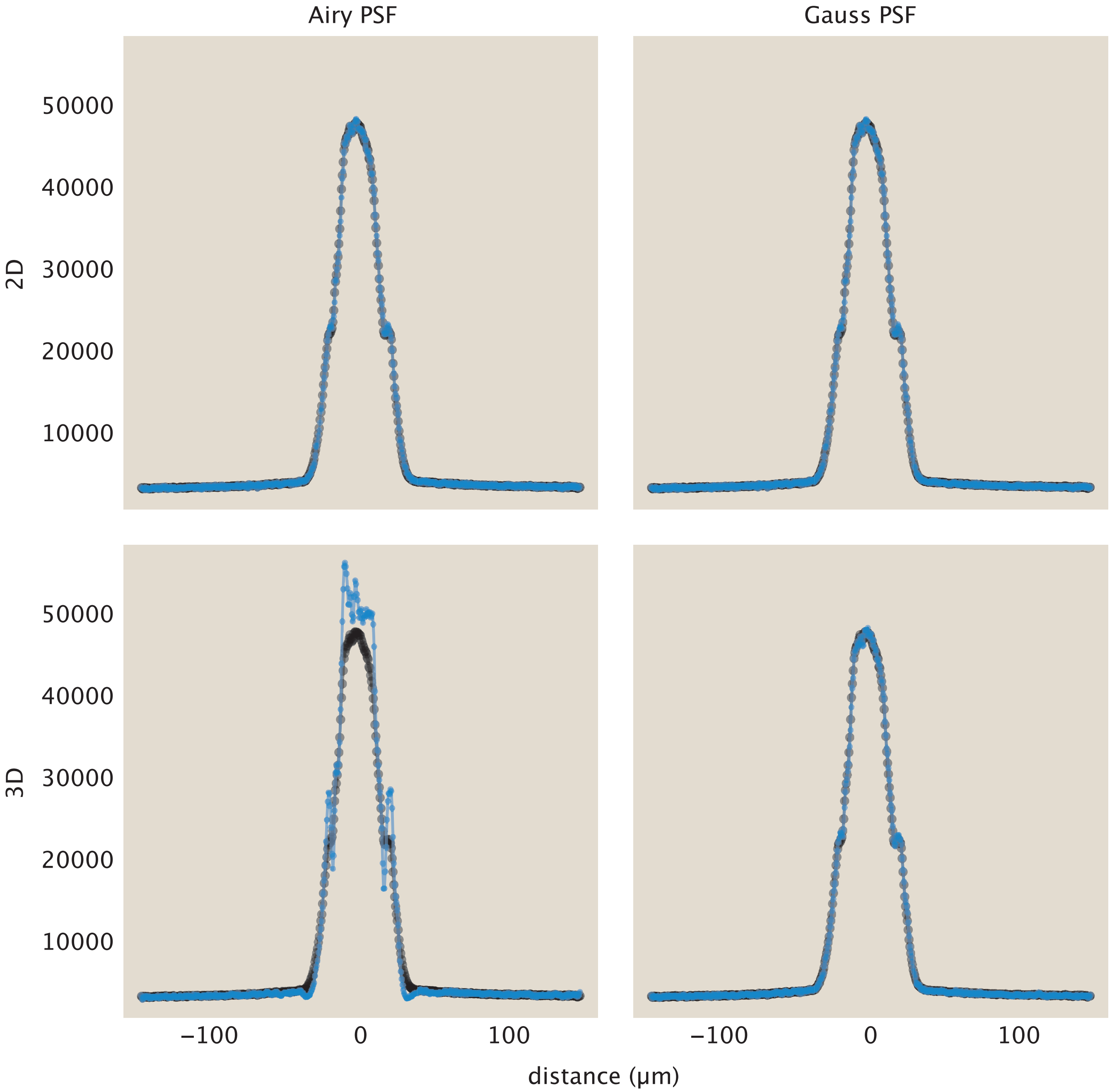

**Fig. S64. 3D deconvolution in ImageJ for Airy or Gaussian PSFs show little deviation from the original image.** For all plots, we include the raw image intensity cutline in black and the deconvolved cutline in blue. The top row highlights the ImageJ 2D deconvolutions while the bottom row highlights their 3D counterparts.

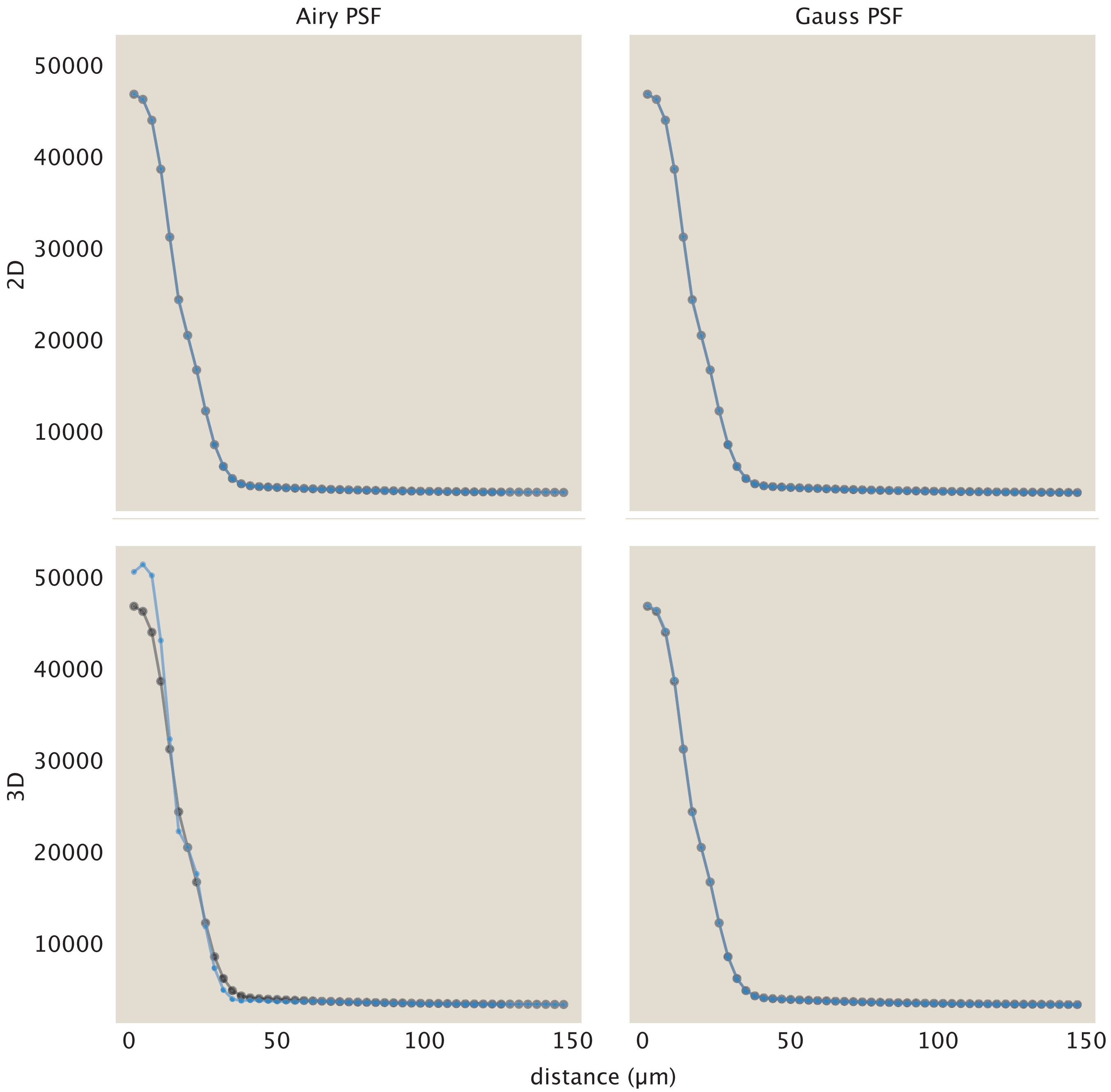

**Fig. S65. Polar averaging deconvolved outputs also largely agree with the raw polar averages.** We take the averaged intensity of volume normalized radial shells and plot how the intensity changes with respect to the distance from the aster center. In black, we show the traces of the raw image and in blue the traces of the deconvolved output. Largely, the deconvolved images in both 2D, on the top row, and in 3D, on the bottom row, match the raw image.

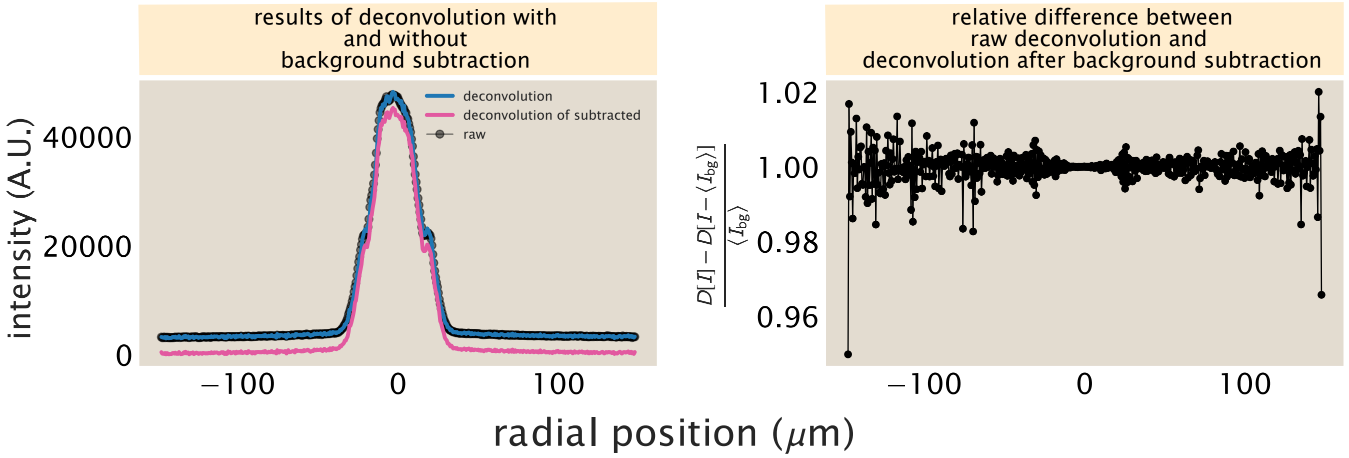

**Fig. S66. Subtraction of ambient background fluorescence leaves deconvolution results unchanged**. (A) Profiles (slices) of fluorescence along a a line through an aster's center before deconvolution (in black, denoted $I$); after deconvolution (in blue, denoted $\mathcal{D}[I]$ where $\mathcal{D}$ is the deconvolution operation); and when deconvolution is performed after the ambient $\langle I_{\text{bg}} \rangle$ average level of fluorescence in the regions of space outside the aster is subtracted, $\mathcal{D}[I - \langle I_{\text{bg}} \rangle]$,shown in pink. These traces affirm that the subtraction of ambient fluorescence before or after deconvolution does not affect the shape of the intensity fluorescence profile. These deconvolutions were performed in two dimensions using an Airy-function based point-spread function in DeconvolutionLab2 (via Imagej), as described above. (B) The difference $\frac{D[I]-D[I-\langle I_{\text{bg}} \rangle]}{\langle I_{\text{bg}} \rangle}$ of the blue and black traces in panel (A) relative to the ambient $\langle I_{\text{bg}} \rangle$ level of fluorescence. The fact that this value is consistently very close to unity affirms that $D[I] - D[I - \langle I_{\text{bg}} \rangle] \approx \langle I_{\text{bg}} \rangle$, namely that the background subtraction operation is essentially commutative with respect to deconvolution.

This finding that a background does not nontrivially modify the result of deconvolution (for relevant aster data of interest) complements generic numerical guarantees applying to Richardson-Lucy deconvolution. Studying the deconvolution operation in the context of astronomical images, Prato and coworkers (31) establish that performing Richardson-Lucy deconvolution conserves intensity flux both locally and globally in input images, when the background of the image is zero. Our numerical analysis suggests that the latter proviso that background is zero may not be required for these algorithms on our class of data.

In sum, we find deconvolving has little to no impact on the image. This is especially clear in Figure S65 where only the 3D Airy deconvolution shows any discrepancy between the raw and deconvolved images. And even here, there is a maximum of approximately a 10% difference between the intensity values within 15 $\mu$m of the aster center. Based on these results, we do not find it imperative to include a deconvolution procedure when analyzing aster data.

**E. Effects of convolution on image formation.** We were curious to better understand why the deconvolution algorithms led to very modest changes in our measurements. To investigate these questions more deeply, in this section, we examine synthetic data that we subject to the deconvolution procedure to dissect the physics of the deconvolution process.

As noted in the previous section, the formation of an image in a microscope includes convolution with the microscope's point spread function. The point spread function provides a mathematical description of how light from a point source is affected by diffraction and other optical limitations as it passes through the optical path of the microscope. This convolution causes the resultant image to appear smoother, or blurred, compared to the object itself. As this work measures the number of ATPs at a given location in time via fluorescence, it is important to determine how the PSF spreads and attenuates fluorescence signal, affecting the apparent localization and intensity of ATPs. To get a feeling for the impact of convolution, we mimic the image acquisition procedure on synthetic data. In particular, we specify a symmetric sphere of fluorescent motor proteins whose concentration decays radially. We model the motor concentration field using an exponential decay corresponding to the object

$$\text{obj} = \exp\left(\frac{-\sqrt{x^2+y^2+z^2}}{\lambda_o}\right), \tag{108}$$

where $\lambda_o$ is the decay length, which we take to be $\approx 30\,\mu$m for a fully formed aster. For convenience, we do not normalize the object, ensuring that at the origin the intensity value is one.

Our first objective is to explore how convolution modifies the “measured" intensities. To perform a convolution mimicking the imaging process in the microscope, we define a point spread function as a 3D Gaussian. We assume that the point spread function has different length scales in the transverse ($x$-$y$) plane compared to the axial ($z$) direction. The form of our synthetic point spread function is

$$\text{psf}(x,y,z) = \text{psf}_{\text{tr}}(x,y) \times \text{psf}_{\text{ax}}(z) = \frac{1}{2\pi\sigma_{\text{tr}}^2}\exp\left(\frac{-(x^2+y^2)}{2\sigma_{\text{tr}}^2}\right) \times \frac{1}{\sqrt{2\pi\sigma_{\text{ax}}^2}}\exp\left(\frac{-z^2}{2\sigma_{\text{ax}}^2}\right) \tag{109}$$

We estimate the length scale for the transverse point spread function based on the minimum resolvable distance. For a self-luminous light source of incoherent light, which encompasses fluorescent signals, the minimum distance is

$$\sigma_{\text{tr}} \approx \frac{0.61\lambda}{\text{N.A.}}, \tag{110}$$

where $\lambda$ is the wavelength of light and N.A. is the numerical aperture (26). For our setup, we use a wavelength of 513 nm and our objective has an N.A. of 0.45. These parameters yield the transverse length scale,

$$\sigma_{\text{tr}} = \frac{0.61 \times 513 \text{ nm}}{0.45} = 695 \text{ nm} \approx 0.7\,\mu\text{m}. \tag{111}$$

For the axial length scale, we compute the depth of focus as

$$\sigma_{\text{ax}} \approx \frac{\lambda\; n}{\text{N.A.}^2}, \tag{112}$$

as stated in (32), where $n$ is the index of refraction, which we take as $n = 1$ for an air objective. Plugging in values, we estimate

$$\sigma_{\text{ax}} \approx \frac{0.513\ \mu m \times 1}{(0.45^2)} \approx 2.5\ \mu m. \tag{113}$$

Now that we have the parameters describing the point spread function in hand, we now turn to examining its effects. The convolution of the object with the point spread function is characterized by the integral

$$\text{im}(x,y,z) = \int_{-\infty}^{\infty}\int_{-\infty}^{\infty}\int_{-\infty}^{\infty} \text{obj}(x',y',z')\text{psf}(x-x',y-y',z-z')\,\mathrm{d}x'\mathrm{d}y'\mathrm{d}z' \tag{114}$$

which amounts to weighting each point in the image with all points in the object, each appropriately multiplied by the point spread function itself.

We use the scipy package, *scipy.signal.convolve* to perform this convolution. We set the "mode" keyword argument to "same" which ensures the returned convolved matrix has the same size as the inputted object. The microscope would acquire an image that reflects the center slice of the convolution. We compare the center slice of the synthetic specified object and the convolved object in figure S67(A). By eye it is difficult to distinguish the two slices from one another. To be more precise, we draw a cutline through the center of the object as depicted in figure S67(B). The difference after convolution is still quite minimal, though we do see a difference near the center such that the convolved image becomes smoothed and has a lower intensity value. We demonstrate the relative change in figure S67(C) as the difference of the convolved object from the raw object divided by the raw object,

$$\Delta I = \frac{\text{obj} - \text{obj} \star \text{psf}}{\text{obj}}. \qquad [115]$$

There is about a 10% change in the signal at the center of the object and very little change beyond $10\,\mu$m outside the center. The spikes at the ends of the plot are due to edge effects occurring from convolution.

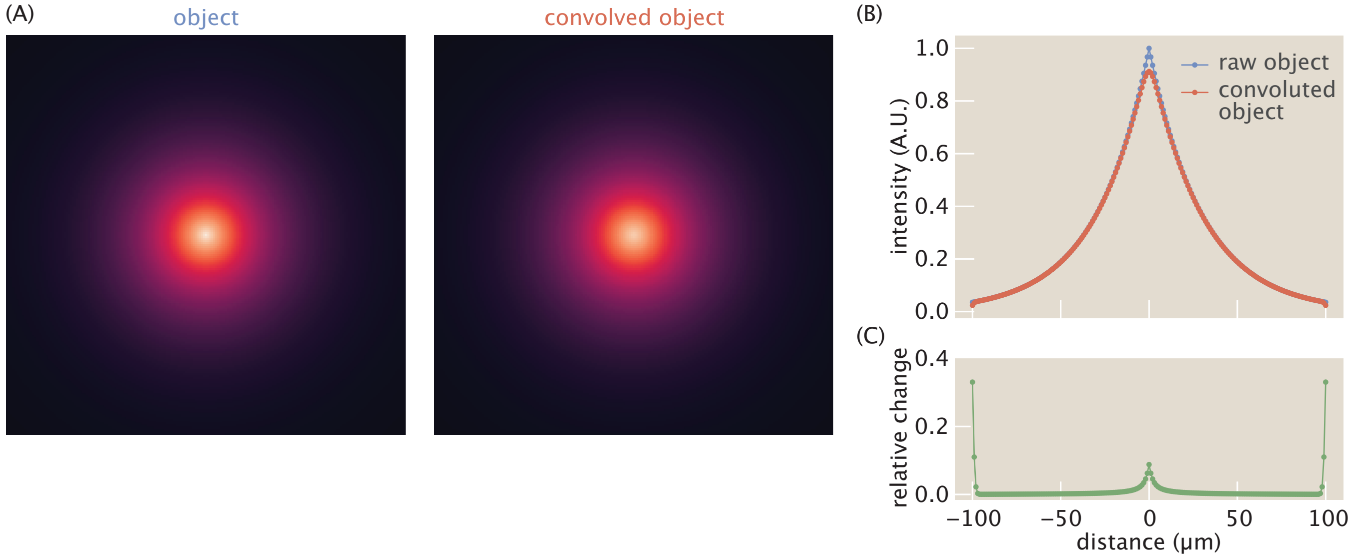

**Fig. S67. Exploring the effects of convolution with the point spread function for synthetically generated data.** (A) Images of the center plane for both the raw object, a radially decaying exponential sphere of motors, and the convolved image with a gaussian point spread function. (B) More quantitatively, we plot the intensity of both the object and convolved object along a cut line passing through the object origin. (C) We find most of the relative change, as defined as $\frac{\text{obj}-\mathrm{obj}\star\mathrm{psf}}{\text{obj}}$, occurs in the center of the object. The peaks at the ends are due to edge effects in the convolution.

The variation in the image intensity as a result of the convolution with the point spread function appears to be minimal, and is only noticeable at the very center. This begs the question: how necessary is incorporating the corrections due to the point spread function in the analysis of our images, and specifically, how will this procedure alter our measured concentrations? To that end, we perform a sweep of sigma values to determine at what Gaussian length scales convolution significantly alters the intensity reported in the image as contrasted with the intensity of the object.

From the data reported in figure S68, we find that when convolving with small $\sigma$ values ($\sigma \leq 3\,\mu$m), the resultant traces are very similar to the original object. The bottom plot gives the relative change of the intensity values after convolving as described in Equation 115. Except for the edges of the plot, where we believe boundary effects create increases, convolutions with small $\sigma$ values reveal discrepancies of under ten percent.

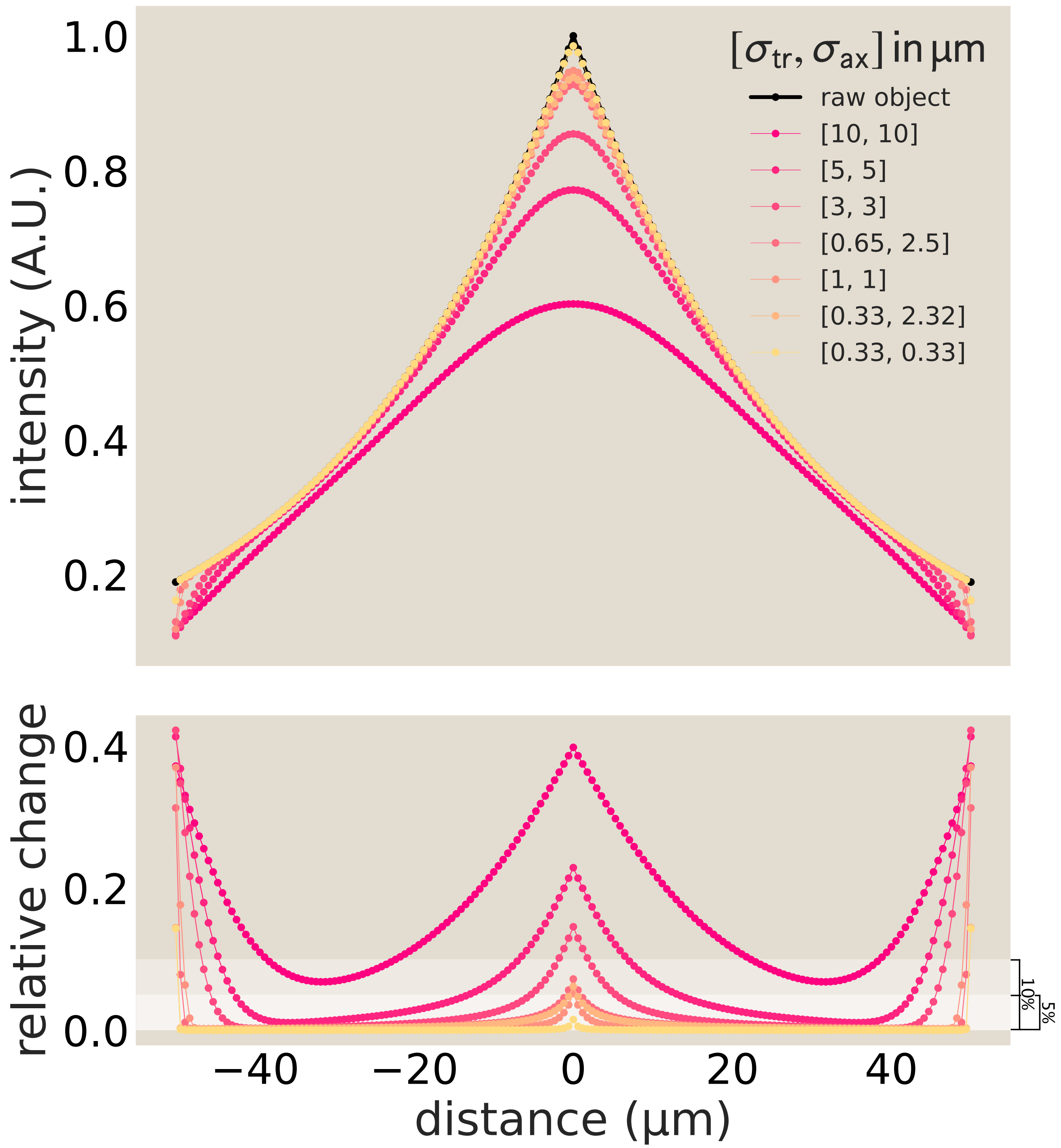

**Fig. S68. Comparison of object and object convoluted with Gaussian point spread function.** (A) The profile of the synthetic object is plotted in black with convolved "images" with varying PSF length scales overlayed. For small length scales, $\sigma < 3\,\mu\mathrm{m}$, the "image" does a good job of approximating the object. The discrepancies are highlighted in (B) where we plot the relative change between the image and object. Most of the variation occurs in the center. Increases at the edges are thought to be boundary effects.

**F. Dimensionality of reaction diffusion systems.** The data presented in this manuscript is presented as two-dimensional images as the optical micrographs are two dimensional as well. For both quantitative and qualitative analysis, it is crucial to consider a *full three-dimensional treatment*, as the reaction–diffusion equation behaves qualitatively differently in 2D versus 3D. To illustrate this, consider Fick's first law for a diffusing species with diffusion coefficient $D$, concentration $\rho$, and particle flux $\vec{j}$

$$\vec{j} = -D\vec{\nabla}\rho \qquad , \tag{116}$$

which, together with the continuity equation that expresses particle conservation

$$\frac{\partial \rho}{\partial t} + \vec{\nabla} \cdot \vec{j} = 0 \qquad , \tag{117}$$

yields Fick's second law:

$$\frac{\partial \rho}{\partial t} = D\vec{\nabla}^2 \rho \qquad . \tag{118}$$

For a perfectly absorbing sink at the origin with absorption rate $c$ in a round domain with radial, *i.e.* spherical or circular, symmetry, the particle flux $\vec{j}$ depends only on the radial distance $r$ (figure S69). The particle flow $I$ through a closed gaussian surface is

$$I = \oint \vec{j} \cdot \mathrm{d}\vec{S} \qquad . \tag{119}$$

Due to symmetry, the flux $\vec{j}$ is in the $\hat{r}$ direction, therefore $I = -jA$ if $j$ is inwards, where $A$ is the surface area of the gaussian surface. Particle conservation requires that $I$ is not a function of $r$ between the outer edge of the domain and the absorber. Hence $I(a \leq r \leq b) = c$, a constant, and equal to the absorber's rate. Therefore, the steady-state fluxes are

$$\vec{j}_{3D} = -\frac{c}{4\pi r^2}\hat{r}, \qquad \vec{j}_{2D} = -\frac{c}{2\pi r}\hat{r} \qquad , \tag{120}$$

where $c$ is the total number of particles absorbed per unit time. Corresponding steady-state concentrations are

$$\rho_{3D} = \frac{c}{4\pi r D} \qquad , \qquad \rho_{2D} = \log(r)\frac{c}{2\pi D} \qquad , \tag{121}$$

or equivalently, the Green's function of the Laplacian is $1/r$ in 3D and $\log(r)$ in 2D. The solutions behave qualitatively very differently.

The particle current $I$ through a small spherical or circular object of radius $a$ around the origin remains finite in both cases as $a \to 0$

$$\lim_{a \to 0} I_{3D} = \lim_{a \to 0} I_{2D} = c \qquad . \tag{122}$$

However, the number of particles near the sink $N$ behaves differently:

$$N_{3D} = \int \rho \, \mathrm{d}V = \int_0^a \frac{c}{4\pi r D} 4\pi r^2 \mathrm{d}r = ca^2/2D \to 0 \quad \text{as } a \to 0 \qquad , \tag{123}$$

while

$$N_{2D} = \int_0^a \log(r)\frac{c}{2\pi D}\, 2\pi r \, \mathrm{d}r \sim ca^2 \log(a)/2D \to \infty \quad \text{as } a \to 0. \tag{124}$$

Thus, in 2D, the divergence requires regularization through a microscopic cutoff radius $a$.

This has a profound effect on the timescales obtained for depletion of species with an initial uniform density by an absorbing sink. Through separation of variables, $\rho(r,t) = R(r)T(t)$, Fick's second law reduces to

$$\frac{1}{D}\frac{T'(t)}{T(t)} = \frac{\nabla^2 R}{R} = -k^2 \quad \Rightarrow \quad T(t) = e^{-Dk^2 t}, \quad \lambda = Dk^2, \tag{125}$$

where $k^2$ is the eigenvalue of the radial Laplacian subject to boundary conditions.

For a spherical domain of radius $b$ with a finite sink of radius $a$, the radial solution is

$$R(r) = \frac{A\sin(kr) + B\cos(kr)}{r} \qquad . \tag{126}$$

Applying the inner boundary condition $R(a) = 0$ gives $B = -A\tan(ka)$, so that

$$R(r) = \frac{A}{\cos(ka)}\frac{\sin(kr - ka)}{r} \qquad , \tag{127}$$

and the no-flux condition at $r = b$ leads to the transcendental equation

$$\tan(kb - ka) = kb \qquad . \tag{128}$$

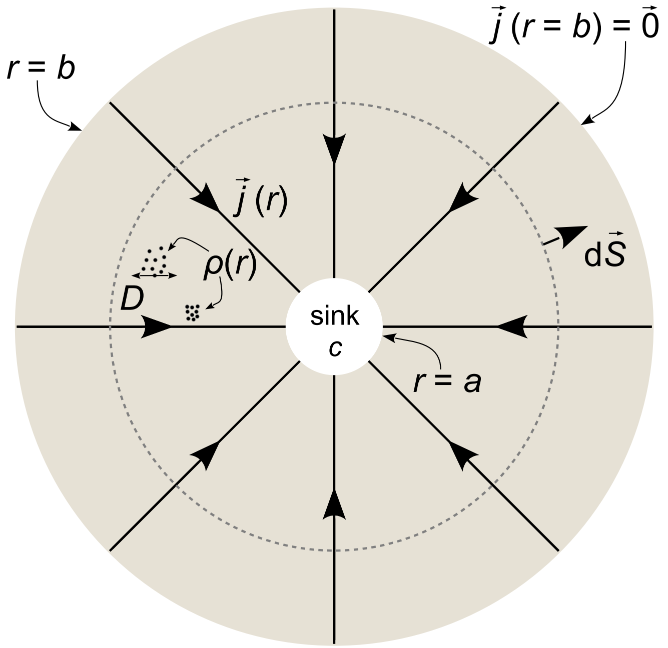

**Fig. S69. Concentration of a reactive species in a spherically symmetric domain around a perfect absorber.** An absorber with rate $c$ is located in a region $r \leq a$ centrally located in a round region with boundary $r = b$. A diffusing species with diffusion coefficient $D$ has a spatial concentration $\rho(r)$ (black dots). The absorption causes an inward particle flux $\vec{j} = -D\vec{\nabla}\rho$ (inward arrows). The particle flux through a gaussian surface (dashed circle) is $I = \oint \vec{j} \cdot \mathrm{d}\vec{S}$.

In the limit of a small sink $a \ll b$, the lowest eigenvalue is

$$k \approx k_0 \left(1 + \frac{a}{b}\right), \quad k_0 b \approx 4.493 \tag{129}$$

so that the slowest decay rate becomes

$$\lambda_{3D} \approx D k_0^2 \left(1 + 4\frac{a}{b}\right) = D\frac{(4.493)^2}{b^2}\left(1 + 4\frac{a}{b}\right) \quad . \tag{130}$$

Notice that the decay rate is only weakly dependent on the sink radius $a$.

For a circular domain of radius $b$, the radial solution involves Bessel functions:

$$R(r) = AJ_0(kr) + BY_0(kr) \quad , \tag{131}$$

where $J_0$ and $Y_0$ are Bessel functions of order zero. Applying the inner boundary $R(a) = 0$ and the no-flux condition at $r = b$ gives the transcendental equation

$$\frac{J_0(ka)}{Y_0(ka)} = \frac{J_1(kb)}{Y_1(kb)} \quad . \tag{132}$$

In the limit of a small sink $a \ll b$, this is solved approximately by

$$k^2 \approx \frac{2}{b^2 \log(2b/a)} \quad \Rightarrow \quad \lambda_{2D} \approx \frac{2D}{b^2 \log(2b/a)} \quad . \tag{133}$$

Here, the logarithmic dependence on the sink size $a$ leads to a much slower relaxation compared with 3D. More importantly, this implies the relaxation rate depends on the grid size in finite element situations, the logarithmic cutoff in analytic treatments, or the finite size of the reactive species in experiments. This is in stark contrast to the 3D result.

#### 8. Effects of Competitive Inhibition by ADP and Phosphate on ATP Hydrolysis Rates

**A. States and Weights Modeling of ATP Hydrolysis.** To quantitatively dissect the relationship between motor proteins and ATP, we develop a model describing the concentration of ATP throughout space and time. ATP can diffuse and undergo hydrolysis due to motor proteins throughout our system; however, ATP cannot regenerate. The diffusive term is straight forward and takes the form of Fick's Law, $D\nabla^2 A$. The reaction term needs to take into account the rate at which ATP can bind to a motor, along with the probability of binding. As a first pass, we model this reaction term just taking into account the statistical mechanics of ATP binding to a motor. However, this fails to acknowledge the dynamics of ATP being converted to ADP and phosphate. We elaborate on the ATP only model to include competitive inhibition resulting from ADP, as well as phosphate.

***A.1. ATP Only Model - Assume No Products Present.*** We use a states and weights approach, as outlined in reference (33), to determine the probability of an ATP molecule binding to a motor protein for $T$ ATP molecules in a lattice with $\Omega$ sites, one of which is the motor protein binding site. In this case, there are two states for the motor protein, as depicted in Figure S70. It can be unbound with energy $E = 0$ and multiplicity $\frac{\Omega!}{T!(\Omega-T)!} \approx \frac{\Omega^T}{T!}$ for $\Omega >> T$. Alternatively the motor protein can be bound, with an energy of $E = \epsilon_T$ and multiplicity $\approx \frac{\Omega^{(T-1)}}{(T-1)!}$. Putting this together, we find the probability of binding is

$$p_{\text{bound}} = \frac{\frac{\Omega^{(T-1)}}{(T-1)!}e^{-\beta\epsilon_T}}{\frac{\Omega^T}{T!} + \frac{\Omega^{(T-1)}}{(T-1)!}e^{-\beta\epsilon_T}}. \tag{134}$$

We simplify the probability by multiplying the numerator and denominator by $\frac{T!}{\Omega^T}$ to get

$$p_{\text{bound}} = \frac{\frac{T}{\Omega}e^{-\beta\epsilon_T}}{1 + \frac{T}{\Omega}e^{-\beta\epsilon_T}}. \tag{135}$$

To convert $\Omega$, the number of "lattice sites", to a volume, we multiply the numerator and denominator by $\frac{\Delta V}{\Delta V}$, the small volume each lattice site represents. By specifying $c_0 = \frac{1}{\Delta V}$ and $[\text{ATP}] = \frac{T}{\Omega \Delta V}$, our probability now depends on concentrations and becomes

$$p_{\text{bound}} = \frac{\frac{[\text{ATP}]}{c_0}e^{-\beta\epsilon_T}}{1 + \frac{[\text{ATP}]}{c_0}e^{-\beta\epsilon_T}}. \tag{136}$$

Lastly, we define a constant $K_{\text{T}} = c_0 e^{\beta\epsilon_{\text{T}}}$ which gives the result

$$p_{\text{bound}} = \frac{\frac{[\text{ATP}]}{K_{\text{T}}}}{1 + \frac{[\text{ATP}]}{K_{\text{T}}}}, \tag{137}$$

in the form of a Michaelis-Menten (Langmuir) binding curve. Thus the constant $K_{\text{T}}$ is interpreted as the concentration at which the chance of an ATP binding to a motor is fifty percent.

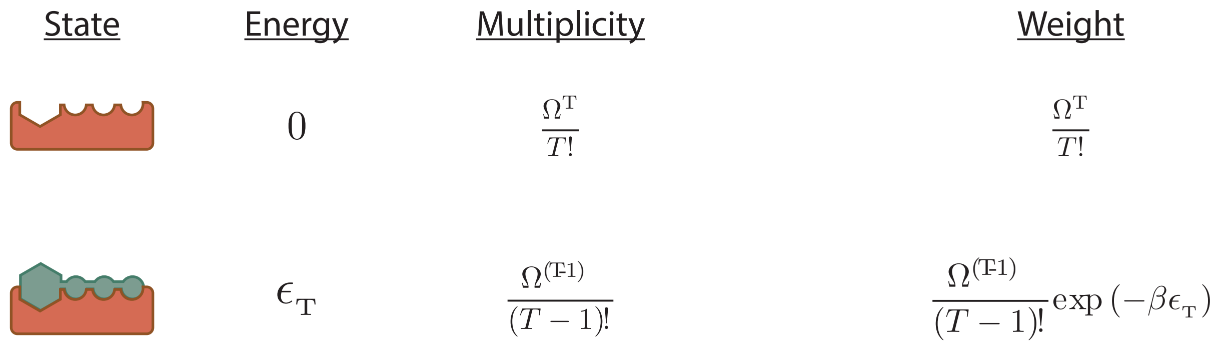

**Fig. S70. States and Weights of ATP binding to Motors.** Energy, multiplicity and weights for the binding states of a single motor in a lattice with $\Omega$ sites and $T$ ATP molecules.

***A.2. ADP Competitive Inhibition - Assume a Single Product.*** We now add in the effect of accumulating ADP into our model. If ADP binds to the motor, ATP cannot bind which creates competitive inhibition. Similar to our treatment of the ATP only system, we write down probability in terms of the weights of each state. This system has three states: the motor protein is unbound, ATP is bound, or ADP is bound, as shown in Figure S71. Once again, we denote the lattice sites as $\Omega$, the number of ATP molecules as T, and the number of ADP molecules as D. We write the probability of the ATP bound state as

$$p_{\mathrm{bound}} = \frac{\frac{\Omega^{T-1}\Omega^{D}}{(T-1)!D!}\exp\left(-\beta\epsilon_{\mathrm{T}}\right)}{\frac{\Omega^{T}\Omega^{D}}{T!D!} + \frac{\Omega^{T-1}\Omega^{D}}{(T-1)!D!}\exp\left(-\beta\epsilon_{\mathrm{T}}\right) + \frac{\Omega^{T}\Omega^{(D-1)}}{T!(D-1)!}\exp\left(-\beta\epsilon_{\mathrm{D}}\right)}. \quad [138]$$

Multiplying the numerator and denominator by $\frac{T!D!}{\Omega^{T}\Omega^{D}}$ simplifies the expression to

$$p_{\mathrm{bound}} = \frac{\frac{T}{\Omega}\exp\left(-\beta\epsilon_{\mathrm{T}}\right)}{1 + \frac{T}{\Omega}\exp\left(-\beta\epsilon_{\mathrm{T}}\right) + \frac{D}{\Omega}\exp\left(-\beta\epsilon_{\mathrm{D}}\right)}. \quad [139]$$

We can convert this into a concentration friendly equation through the substitutions $[\mathrm{ATP}] = \frac{T}{\Omega\Delta V}$, $[\mathrm{ADP}] = \frac{D}{\Omega\Delta V}$, and $K_{T,D} = \frac{1}{\Delta V}e^{\beta\epsilon_{T,D}}$, which results in

$$p_{\mathrm{bound}} = \frac{\frac{[\mathrm{ATP}]}{K_T}}{1 + \frac{[\mathrm{ATP}]}{K_T} + \frac{[\mathrm{ADP}]}{K_D}}. \quad [140]$$

Equation 140 is of a similar form to equation 137, but has an extra term in the denominator corresponding to the likelihood of binding ADP. Thus as more ATP turns to ADP, the probability of ATP binding to the motor decreases.

Note that if we consider a model of ATP and phosphate with no ADP present, the form should be the same as in equation 140 where [ADP] is replaced with $[\mathrm{P}_i]$ and $K_{\mathrm{D}}$ is replaced with $K_{\mathrm{P}}$.

***A.3. ADP and Phosphate Competitive Inhibition - Assume Both Products are Present.*** We can go a step further with this model by considering both hydrolysis products, ADP and phosphate, to be present in solution. This creates two new inhibition states: phosphate is bound, or phosphate and ADP are bound. We follow a similar procedure as in the ADP only model, though now we must take into account three species, ATP ($T$), ADP ($D$), and phosphate ($P$). As depicted in Figure S72, let us define the energies of the bound phosphate, and the ADP and phosphate bound states, as $\epsilon_{\mathrm{P}}$ and $\epsilon_{\mathrm{D \cdot P}} = \epsilon_{\mathrm{D}} + \epsilon_{\mathrm{P}} + \sigma'$ respectively. Here we guess the energy of both species binding is the sum of the individual binding energies, plus or minus some interaction energy, $\sigma'$. For, again, $\Omega$ binding sites, the probability of an ATP being bound to the motor is:

| State | Energy | Multiplicity | Weight |
|---|---|---|---|
| 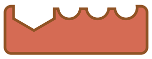 | $0$ | $\frac{\Omega^{\mathrm{T}}}{T!}\frac{\Omega^{\mathrm{D}}}{D!}$ | $\frac{\Omega^{\mathrm{T}}}{T!}\frac{\Omega^{\mathrm{D}}}{D!}$ |
| 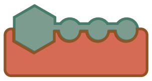 | $\epsilon_{\mathrm{T}}$ | $\frac{\Omega^{(\mathrm{T}\text{-}1)}}{(T-1)!}\frac{\Omega^{\mathrm{D}}}{D!}$ | $\frac{\Omega^{(\mathrm{T}\text{-}1)}}{(T-1)!}\frac{\Omega^{\mathrm{D}}}{D!}\exp\left(-\beta\epsilon_{\mathrm{T}}\right)$ |
| 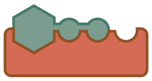 | $\epsilon_{\mathrm{D}}$ | $\frac{\Omega^{\mathrm{T}}}{T!}\frac{\Omega^{(\mathrm{D}\text{-}1)}}{(D-1)!}$ | $\frac{\Omega^{\mathrm{T}}}{T!}\frac{\Omega^{(\mathrm{D}\text{-}1)}}{(D-1)!}\exp\left(-\beta\epsilon_{\mathrm{D}}\right)$ |

**Fig. S71.** States and Weights for Competitive Inhibition of ADP on a Motor. Energy, multiplicity and weights for three binding states: unbound motor, bound ATP, and bound ADP. Assume a lattice with $\Omega$ sites, one of which is the motor protein, $T$ ATP molecules and D ADP molecules.

$$p_{\text{bound}} = \underbrace{\frac{\Omega^{T-1}\Omega^{D}\Omega^{P}}{(T-1)!D!P!}\exp\left(-\beta\epsilon_{\mathrm{T}}\right)}_{\text{ATP Bound}} \cdot \left[\underbrace{\frac{\Omega^{T}\Omega^{D}\Omega^{P}}{T!D!P!}}_{\text{Unbound}} + \underbrace{\frac{\Omega^{T-1}\Omega^{D}\Omega^{P}}{(T-1)!D!P!}\exp\left(-\beta\epsilon_{\mathrm{T}}\right)}_{\text{ATP Bound}} + \underbrace{\frac{\Omega^{T}\Omega^{(D-1)}\Omega^{P}}{T!(D-1)!P!}\exp\left(-\beta\epsilon_{\mathrm{D}}\right)}_{\text{ADP Bound}} + \underbrace{\frac{\Omega^{T}\Omega^{D}\Omega^{(P-1)}}{T!D!(P-1)!}\exp\left(-\beta\epsilon_{\mathrm{P}}\right)}_{\text{P Bound}} + \underbrace{\frac{\Omega^{T}\Omega^{(D-1)}\Omega^{(P-1)}}{T!(D-1)!(P-1)!}\exp\left(-\beta(\epsilon_{\mathrm{D}}+\epsilon_{\mathrm{P}}+\sigma')\right)}_{\text{ADP and P Bound}}\right]^{-1}. \quad [141]$$

We can simplify this expression by multiplying the numerator and denominator by $\frac{T!D!P!}{\Omega^{T}\Omega^{D}\Omega^{P}}$, which gives us

$$p_{\text{bound}} = \frac{\frac{T}{\Omega}\exp\left(-\beta\epsilon_{\mathrm{T}}\right)}{1+\frac{T}{\Omega}\exp\left(-\beta\epsilon_{\mathrm{T}}\right)+\frac{D}{\Omega}\exp\left(-\beta\epsilon_{\mathrm{D}}\right)+\frac{P}{\Omega}\exp\left(-\beta\epsilon_{\mathrm{P}}\right)+\frac{DP}{\Omega\Omega}\exp\left(-\beta(\epsilon_{\mathrm{D}}+\epsilon_{\mathrm{P}}+\sigma')\right)}. \quad [142]$$

Once more converting to units of concentration, we take $[\mathrm{ATP}] = \frac{T}{\Omega\Delta V}$, $[\mathrm{ADP}] = \frac{D}{\Omega\Delta V}$, $[\mathrm{P}] = \frac{P}{\Omega\Delta V}$, $K_{T,D,P} = \frac{1}{\Delta V}e^{\beta\epsilon_{T,D,P}}$, and define $\sigma = e^{\beta\sigma'}$ which results in

$$p_{\text{bound}} = \frac{\frac{[\mathrm{ATP}]}{K_T}}{1+\frac{[\mathrm{ATP}]}{K_T}+\frac{[\mathrm{ADP}]}{K_D}+\frac{[\mathrm{P}]}{K_P}+\frac{[\mathrm{ADP}][\mathrm{P}]}{\sigma K_D K_P}}. \quad [143]$$

This model demonstrates how products of ATP hydrolysis compete with the motor binding sites. As more possible states emerge, the probability of ATP will binding to the motor protein reduces. We can directly see an impact on ATP hydrolysis by defining the hydrolysis rate as the probability of binding ATP multiplied by the maximum hydrolysis rate. This is written as

$$\Gamma\left([\mathrm{ATP}],[\mathrm{ADP}],[\mathrm{P}_i]\right) = \gamma\cdot\frac{\frac{[\mathrm{ATP}]}{K_{\mathrm{T}}}}{1+\frac{[\mathrm{ATP}]}{K_{\mathrm{T}}}+\frac{[\mathrm{ADP}]}{K_{\mathrm{D}}}+\frac{[\mathrm{P}]}{K_{\mathrm{P}}}+\frac{[\mathrm{ADP}][\mathrm{P}]}{\sigma K_{\mathrm{D}}K_{\mathrm{P}}}}, \quad [144]$$

where $\gamma$ is the ATP hydrolysis per motor per second for saturating ATP conditions with zero product concentrations.

| State | Energy | Multiplicity | Weight |
|---|---|---|---|
| 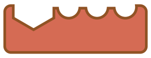 | $0$ | $\frac{\Omega^{T}}{T!}\frac{\Omega^{D}}{D!}\frac{\Omega^{P}}{P!}$ | $\frac{\Omega^{T}}{T!}\frac{\Omega^{D}}{D!}\frac{\Omega^{P}}{P!}$ |
| 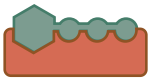 | $\epsilon_{T}$ | $\frac{\Omega^{(T-1)}}{(T-1)!}\frac{\Omega^{D}}{D!}\frac{\Omega^{P}}{P!}$ | $\frac{\Omega^{(T-1)}}{(T-1)!}\frac{\Omega^{D}}{D!}\frac{\Omega^{P}}{P!}\exp\left(-\beta\epsilon_{T}\right)$ |
| 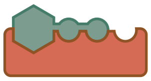 | $\epsilon_{D}$ | $\frac{\Omega^{T}}{T!}\frac{\Omega^{(D-1)}}{(D-1)!}\frac{\Omega^{P}}{P!}$ | $\frac{\Omega^{T}}{T!}\frac{\Omega^{(D-1)}}{(D-1)!}\frac{\Omega^{P}}{P!}\exp\left(-\beta\epsilon_{D}\right)$ |
| 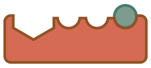 | $\epsilon_{P}$ | $\frac{\Omega^{T}}{T!}\frac{\Omega^{D}}{D!}\frac{\Omega^{(P-1)}}{(P-1)!}$ | $\frac{\Omega^{T}}{T!}\frac{\Omega^{D}}{D!}\frac{\Omega^{(P-1)}}{(P-1)!}\exp\left(-\beta\epsilon_{P}\right)$ |
| 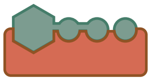 | $\epsilon_{D\cdot P}$ | $\frac{\Omega^{T}}{T!}\frac{\Omega^{(D-1)}}{(D-1)!}\frac{\Omega^{(P-1)}}{(P-1)!}$ | $\frac{\Omega^{T}}{T!}\frac{\Omega^{(D-1)}}{(D-1)!}\frac{\Omega^{(P-1)}}{(P-1)!}\exp\left(-\beta\epsilon_{DP}\right)$ |

**Fig. S72.** States and Weights for Competitive Inhibition of ADP and Phosphate on a Motor. Energy, multiplicity and weights for five binding states: unbound motor, bound ATP, bound ADP, bound phosphate, and bound ADP and phosphate. Once more, assume a lattice with $\Omega$ sites, one of which is the motor protein, $T$ ATP molecules, D ADP molecules, and P phosphate molecules.

#### B. Testing Our Model Against Published Data .

***B.1. Fitting Motor Speeds Versus ATP Concentrations.*** The paper, *Inhibition of kinesin motility by ADP and phosphate supports a hand-over-hand mechanism*, by Schief et al. (10), examines how the speed of kinesin motors vary with ADP and phosphate concentrations. This measurement is very useful to us because speed is proportionally related to ATP hydrolysis rates through the motor step size. The authors define the speed of motors as

$$S = d \cdot k_{\mathrm{cat}} \frac{[\mathrm{ATP}] - \frac{[\mathrm{ADP}][\mathrm{P_i}]}{K_{\mathrm{eq}}}}{K_{\mathrm{M}} + [\mathrm{ATP}]}, \quad [145]$$

where $d$ is the motor step size, $k_{\mathrm{cat}}$ is the per second hydrolysis rate, $K_{\mathrm{M}}$ is the Menten constant, $K_{\mathrm{eq}}$ is the equilibrium constant for hydrolysis, and all terms in brackets are concentrations. Hydrolysis of ATP highly favors the forward direction with $K_{\mathrm{eq}} = 4.9 \cdot 10^{11}$ µM (10), thus the second term in the numerator is negligible, as no concentrations considered exceed $10^{6}$ µM. We now write the speed as

$$S = d \cdot k_{\mathrm{cat}} \frac{[\mathrm{ATP}]}{K_{\mathrm{M}} + [\mathrm{ATP}]}. \quad [146]$$

We scan the measured data from Schief et al. and test if our model fits to it. We first examine Figure 2 from Schief et al., which depicts motor speeds versus ATP concentrations for different levels of hydrolysis products. We fit the data to equation 146, where $k_{\mathrm{cat}}$ and $K_{\mathrm{M}}$ are the fitting parameters. We determine $k_{\mathrm{cat}}$ for the no product data (Figure S73 black curve) and then keep it fixed for all other data sets, thus all other data only has one fitting parameter, $K_{\mathrm{M}}$.

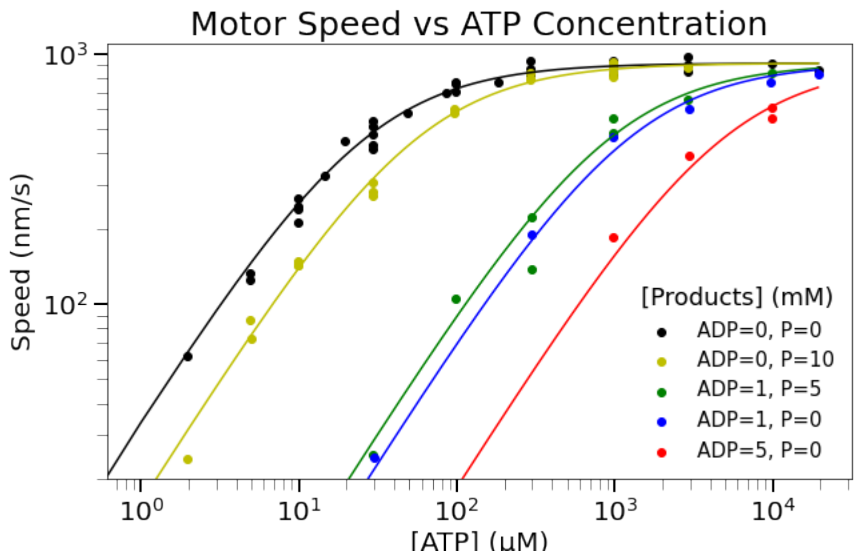

**Fig. S73. Motor speeds for a given ATP concentration depends on the concentrations of ADP and phosphate.** Here, we replicate Figure 2 from Schief et al. The data points were determined by scanning the original figure with WebPlotDigitizer. The lines are fit to a Michaelis-Menten function. For the black curve (where the product concentrations are zero), we fit both $k_{\text{cat}}$ and $K_{\text{T}}$ parameters. For all other curves where products are present, we fit a single parameter, $K_{\text{M}}$, and use $k_{\text{cat}}$, from the no product condition, as an input for this function.

We can express our hydrolysis expression from equation 144 in the form of equation 146 with the following substitutions:

$$\begin{aligned} \Gamma &= \frac{S}{d}, \\ k_{\text{cat}} &= \gamma, \\ K_{\text{M}} &= K_{\text{T}}\left[1 + \frac{[\text{ADP}]}{K_{\text{D}}} + \frac{[\text{P}]}{K_{\text{P}}} + \frac{[\text{ADP}][\text{P}]}{\sigma K_{\text{D}} K_{\text{P}}}\right] \end{aligned} \qquad [147]$$

With these substitutions, we can determine the binding constants for each species. We find $K_{\text{T}}$ from the no products experiment (Figure S73 black curve), which is simple,

$$K_{\text{M}} = K_{\text{T}}. \qquad [148]$$

To determine $K_{\text{D,P}}$, we use the equation

$$K_{\text{D,P}} = [\text{ADP, P}]\left(\frac{K_{\text{M}}}{K_{\text{T}}} - 1\right)^{-1}, \qquad [149]$$

(Figure S73 blue, red, and yellow curves). Finally to determine sigma, we write

$$\sigma = \frac{[\text{ADP}][\text{P}]}{K_{\text{D}} K_{\text{P}}}\left(\frac{K_{\text{M}}}{K_{\text{T}}} - 1 - \frac{[\text{ADP}]}{K_{\text{D}}} - \frac{[\text{P}]}{K_{\text{P}}}\right)^{-1}, \qquad [150]$$

(Figure S73 green curve). This means the equation to find $K_{\text{D}\cdot\text{P}}$ is

$$K_{\text{D}\cdot\text{P}} = [\text{ADP}][\text{P}]\left(\frac{K_{\text{M}}}{K_{\text{T}}} - 1 - \frac{[\text{ADP}]}{K_{\text{D}}} - \frac{[\text{P}]}{K_{\text{P}}}\right)^{-1}. \qquad [151]$$

The tabulated parameters are listed in Table S1

Through this analysis, we find that the values of $K_{\text{T}}$ and $K_{\text{D}}$ are about equivalent, perhaps this is due to similar chemical structure. It can thus be expected that the presence of ADP in our assay will cause significant slowing in the motor hydrolysis rate. On the other hand, $K_{\text{P}}$ is two orders of magnitude larger than $K_{\text{T}}$, implying a small chance of phosphate binding to the motor protein. While we can expect some inhibition due to phosphate, it appears that competitive inhibition due to ADP dominates.

Strangely, $K_{\text{D}\cdot\text{P}}$, and thus $\sigma$, is negative. It is also very large, four orders of magnitude larger than $K_{\text{T}}$. A negative binding constant would mean that the presence of both ADP and phosphate reduces competitive inhibition for the motor protein, while only having one of the species present creates increases the inhibition. This does not seem physical. At best we would expect that the ADP and phosphate always bind together, so the total inhibition is equivalent to inhibition of ADP alone, as this is the more dominant inhibitor. Mathematically, this value for $K_{\text{D}\cdot\text{P}}$ does not jibe with our model because it requires that

| Figure S73 Parameter Fits | | |
|---|---|---|
| Parameter | Value | Description |
| $\gamma$ | $114.5$ s$^{-1}$ | Fitted Parameter |
| $K_{\text{M (Black)}}$ | $26.2$ µM | Fit for 0 mM ADP, 0 mM P |
| $K_{\text{M (Yellow)}}$ | $55.8$ µM | Fit for 0 mM ADP, 10 mM P |
| $K_{\text{M (Blue)}}$ | $1.2$ mM | Fit for 1 mM ADP, 0 mM P |
| $K_{\text{M (Red)}}$ | $4.9$ mM | Fit for 5 mM ADP, 0 mM P |
| $K_{\text{M (Green)}}$ | $930$ µM | Fit for 1 mM ADP, 5 mM P |
| $K_{\text{T}}$ | $26.2$ µM | Equation 148 |
| $K_{\text{D}}$ | $24.5$ µM | Equation 149 |
| $K_{\text{P}}$ | $8.9$mM | Equation 149 |
| $K_{\text{D}\cdot\text{P}}$ | $-714$ mM | Equation 151 |
| $\sigma$ | $-3.3$ µM | Equation 150 |

**Table S1. All the fitted parameters from Figure S73 are reported, along with the resulting Menten constants for each species.**

$\sigma$ is a negative value. We defined $\sigma = e^{-\beta\sigma'}$, where $\sigma'$ is an interaction energy for the duel binding of ADP and phosphate. A negative $\sigma$ implies an imaginary $\sigma'$.

Regardless of the sign, the magnitude of $K_{\mathrm{D\cdot P}}$ indicates that the inhibition due to ADP and P being bound is much smaller than the effect of either product alone and could possibly be negligible. Schief et al. only present one data set (green) with both products present in solution. These experiments seem worth replicating to understand if there is some behavior that is incorrectly incorporated into our model.

Figure S74 examines how the fit changes for different values of $K_{\mathrm{D\cdot P}}$. Setting $K_{\mathrm{D\cdot P}} = \infty$, thereby neglecting the interaction term, is the next best fit to the data after setting $K_{\mathrm{D\cdot P}} = \sigma \cdot K_{\mathrm{D}} \cdot K_{\mathrm{P}}$. Both taking the absolute value of $\sigma$ and setting $\sigma = 1$ produce worse fits.

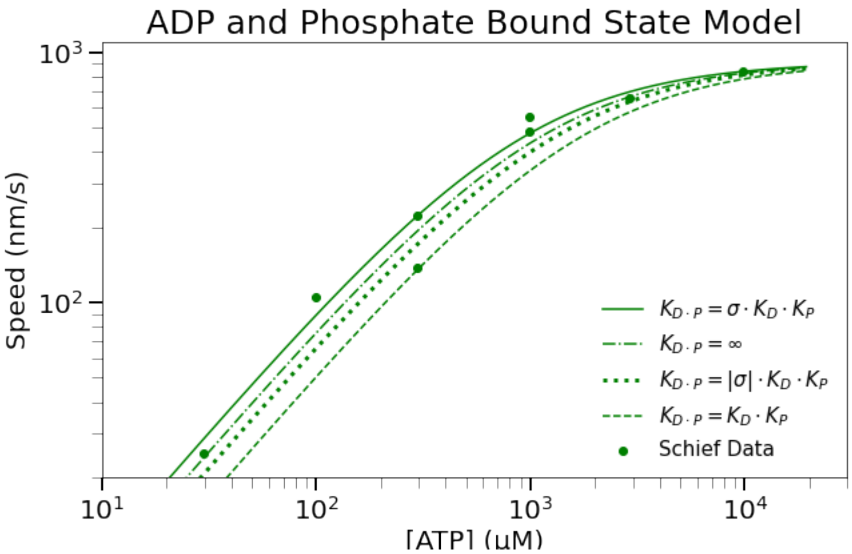

**Fig. S74. Exploration of $K_{\mathrm{D\cdot P}}$ Values for the ADP and Phosphate Bound State**. We examine the fits from varying the $\sigma$ value in $K_{\mathrm{D\cdot P}}$. Using $\sigma = -713$ mM as found from our model, we get the best fit to the data. However, because a negative $\sigma$ does not seem physical, we explore how setting $\sigma$ to 713 mM, 1, and $\infty$ change the fits. We find that entirely neglecting the ADP/Phosphate bound state, setting $\sigma = \infty$ is the next best fit to the data. This is followed by taking $|\sigma| = 713$ mM, as the magnitude reduces the amount of inhibition experienced. Lastly, ignoring an interaction energy, setting $\sigma = 1$ produces the worst fit, as it maximizes the inhibition due to each product.

Additionally from Table S1, we learn about the hydrolysis rate of the motors in question. Our model quotes a rate of 114.5 ATP hydrolyzed per second per motor, for saturating ATP conditions with no products in solution. Previous work also using full length *Drosophila* conventional kinesin measures consistent hydrolysis rates with our fitted rate (34).

***B.2. Fitting Motor Speeds Versus Product Concentrations.*** The authors took additional data investigating motor speeds versus various levels of products at various ATP levels. Using our model, we fit the data to

$$S = d \cdot \gamma \frac{\frac{[\mathrm{ATP}]}{K_T}}{1 + \frac{[\mathrm{ATP}]}{K_{\mathrm{T}}} + \frac{[\mathrm{ADP,\ P}]}{K_{\mathrm{D,\ P}}}}, \qquad [152]$$

where we fit one parameter, $K_{\mathrm{D,\ P}}$. The first set of data, depicted in Figure S75, illustrates the variation in motor speed for different phosphate concentrations. Motor speeds drop with increasing phosphate, especially as ATP concentrations drop, which implies inhibition is occurring.

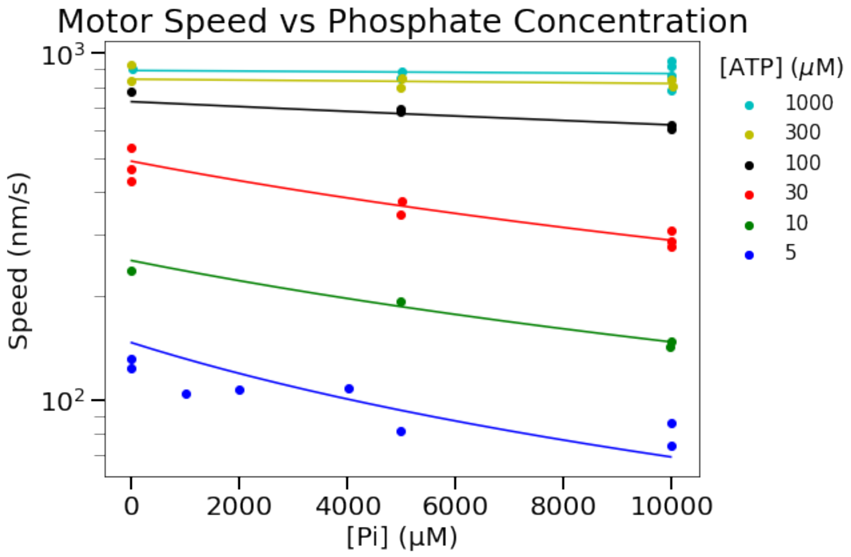

**Fig. S75.** Motor speeds are reduced with increasing phosphate concentrations. Here, we replicate Figure 4A from Schief et al. The data points were determined by scanning the original figure with WebPlotDigitizer. The lines are fit to a Michaelis-Menten function. For each curve we fit a single parameter, $K_{\rm P}$ , and use $k_{\rm cat}$ and $K_{\rm T}$, from Table S1, as inputs.

Our fits result in the $K_{\rm P}$ values quoted in Table S2. This data set produces an average $K_{\rm P}$ of 11.3 mM, comparable to the 8.9 mM value found from the previous data set in Table S1. There does not appear to be a trend in the $K_{\rm P}$ value with ATP concentration.

The authors also examined the variation in speed with ADP concentration, as depicted in Figure S76. Once again, we have plotted the digitized data from Schief et al. and fit the points to our model in equation 152. Motor speeds drop significantly with higher levels of ADP present. Again, this is more drastic for lower ATP concentrations.

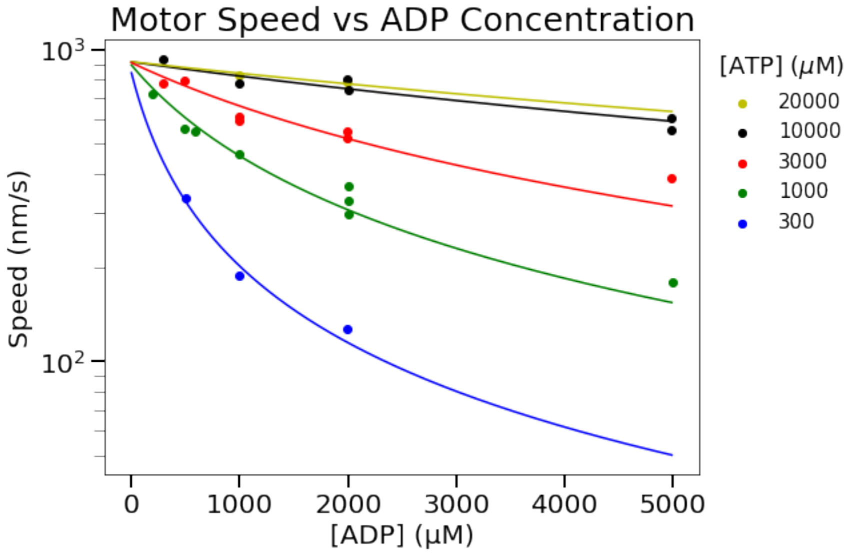

**Fig. S76.** Motor speeds are significantly reduced with increasing ADP concentrations. Here, we replicate Figure 4B from Schief et al. The data points were determined by scanning the original figure with WebPlotDigitizer. The lines are fit to a Michaelis-Menten function. For each curve we fit a single parameter, $K_{\rm D}$, and use $k_{\rm cat}$ and $K_{\rm T}$, from Table S1, as inputs.

Table S3 reports the fitted $K_{\rm D}$ values. On average, $K_{\rm D} = 23.6$ matching the $K_{\rm D} = 24.5$ from table S1. Again, $K_{\rm D}$ values do not appear correlated with ATP concentration.

***B.3. Comparison of Our Model to the Schief Model.*** We now compare our model with the model used in Schief et al. The authors defined $k_{\rm cat}$ and $K_{\rm M}$ as

| $K_P$ Fitted Values for Various ATP Concentrations | |
|---|---|
| ATP Concentration ($\mu$M) | $K_P$ (mM) |
| 1000 | 12.2 |
| 300 | 27.9 |
| 100 | 12.5 |
| 30 | 6.8 |
| 10 | 10.1 |
| 5 | 7.4 |
| Average | 11.3 |

**Table S2. Here we report upon the fits of $K_P$ from the data in Figure S75. We do not find a correlation with ATP concentrations, as expected from our model.**

| $K_D$ Fitted Values for Various ATP Concentrations | |
|---|---|
| ATP Concentration (mM) | $K_D$ ($\mu$M) |
| 20 | 14.7 |
| 10 | 23.8 |
| 3 | 23.0 |
| 1 | 26.7 |
| 0.3 | 25.4 |
| Average | 23.6 |

**Table S3. Here we report upon the fits of $K_D$ from the data in Figure S76. We do not find a correlation with ATP concentrations, as expected from our model.**

$$k_{\mathrm{cat}} = \frac{k_{\mathrm{cat}}^{00}}{1 + \frac{[\mathrm{ADP}]}{K_{\mathrm{ii}}^{\mathrm{ADP}}} + \frac{[\mathrm{P_i}]}{K_{\mathrm{ii}}^{\mathrm{P}}} + \frac{[\mathrm{ADP}][\mathrm{P_i}]}{K_{\mathrm{ii}}^{\mathrm{ADP\cdot P}}}}, \qquad [153]$$

and

$$\frac{K_{\mathrm{M}}}{k_{\mathrm{cat}}} = \frac{K_{\mathrm{M}}^{00}}{k_{\mathrm{cat}}^{00}} \left[1 + \frac{[\mathrm{ADP}]}{K_{\mathrm{i}}^{\mathrm{ADP}}} + \frac{[\mathrm{P_i}]}{K_{\mathrm{i}}^{\mathrm{P}}} + \frac{[\mathrm{ADP}][\mathrm{P_i}]}{K_{\mathrm{i}}^{\mathrm{ADP\cdot P}}}\right] \qquad [154]$$

respectively. The subscript 00 denotes the constants when the products are at zero concentration. The subscript $i$ describes competitive inhibition by the products on the motor protein impacting the Menten term. The subscript $ii$ is described as non-competitive inhibition which modifies the overall stepping rate and the Menten constant. Inputting equations 153 and 154 into equation 146, we find

$$S = d \cdot \frac{k_{\mathrm{cat}}^{00}}{1 + \frac{[\mathrm{ADP}]}{K_{\mathrm{ii}}^{\mathrm{ADP}}} + \frac{[\mathrm{P_i}]}{K_{\mathrm{ii}}^{\mathrm{P}}} + \frac{[\mathrm{ADP}][\mathrm{P_i}]}{K_{\mathrm{ii}}^{\mathrm{ADP\cdot P}}}} \cdot \frac{[\mathrm{ATP}]}{\frac{K_{\mathrm{M}}^{00} k_{\mathrm{cat}}^{00}}{k_{\mathrm{cat}}^{00}} \frac{1 + \frac{[\mathrm{ADP}]}{K_{\mathrm{i}}^{\mathrm{ADP}}} + \frac{[\mathrm{P_i}]}{K_{\mathrm{i}}^{\mathrm{P}}} + \frac{[\mathrm{ADP}][\mathrm{P_i}]}{K_{\mathrm{i}}^{\mathrm{ADP\cdot P}}}}{1 + \frac{[\mathrm{ADP}]}{K_{\mathrm{ii}}^{\mathrm{ADP}}} + \frac{[\mathrm{P_i}]}{K_{\mathrm{ii}}^{\mathrm{P}}} + \frac{[\mathrm{ADP}][\mathrm{P_i}]}{K_{\mathrm{ii}}^{\mathrm{ADP\cdot P}}}} + [\mathrm{ATP}]}. \qquad [155]$$

Simplifying the denominator and dividing the numerator and denominator by $K_{\mathrm{M}}$ yields

$$S = \frac{d \cdot k_{\mathrm{cat}}^{00} \cdot \frac{[\mathrm{ATP}]}{K_{\mathrm{M}}^{00}}}{1 + \frac{[\mathrm{ATP}]}{K_{\mathrm{M}}^{00}} + \frac{[\mathrm{ADP}]}{K_{\mathrm{i}}^{\mathrm{ADP}}} + \frac{[\mathrm{P_i}]}{K_{\mathrm{i}}^{\mathrm{P}}} + \frac{[\mathrm{ADP}][\mathrm{P_i}]}{K_{\mathrm{i}}^{\mathrm{ADP\cdot P}}} + \frac{[\mathrm{ATP}]}{K_{\mathrm{M}}^{00}}\left(\frac{[\mathrm{ADP}]}{K_{\mathrm{ii}}^{\mathrm{ADP}}} + \frac{[\mathrm{P_i}]}{K_{\mathrm{ii}}^{\mathrm{P}}} + \frac{[\mathrm{ADP}][\mathrm{P_i}]}{K_{\mathrm{ii}}^{\mathrm{ADP\cdot P}}}\right)}. \qquad [156]$$

Comparing the Schief et al. model (equation 156) with our model of binding (equation 143), we note many parallels. Dividing $d \cdot k_{\mathrm{cat}}^{00}$, from the Schief model gives the probability of ATP binding to the motor. Note that the hydrolysis rate $k_{\mathrm{cat}}^{00}$ is what we denote $\gamma$. The numerators are of the same form with $K_T = K_{\mathrm{M}}^{00}$. The Schief model contains three additional terms in the denominator, as compared to our model. These three terms imply three additional binding states, ATP and ADP bound, ATP and phosphate bound, or all three species bound.

We report the fitted parameters found by the Schief model, as well as the comparable values we determined in table S1, in table S4.

The values found by both models are very similar with the exception of the $K_{\mathrm{D\cdot P}}$ value. The Schief model yields a positive value about an order of magnitude lower than our model. The low Schief value results in a larger overall $K_{\mathrm{M}}$, which in our model pushes the fit farther away from the measured data. This makes the Schief $K_{\mathrm{D\cdot P}}$ the worst of all the fits for both species present according to figure S74. However, with this exception, we have excellent agreement with all of our other parameters. Finally, the Schief model has three additional parameters, the $K_{\mathrm{ii}}$. These are all very large, in the tens to hundreds of millimolar, and are multiplied by $K_{\mathrm{T}} = 23$ µM, as seen in equation 156. This renders the cross terms effectively negligible to the model and causes the Schief model to be equivalent to our model.

#### 9. Investigation of gradient smoothing due to ATP to probe binding

It is important to take into account the on/off times of ATP to the ATP reporter to understand how reported ATP gradients may be affected by the probe. We can do an estimate to determine how far a probe can travel while bound to an ATP. We estimate the diffusion constant of the ATP probe to be $D = 45\,\mu\mathrm{m}^2/\mathrm{s}$ using equation 33. We then need to take into account the binding rates for the ATP reporter. We use the reporter Queen-7µ A81D which has a reported dissociation constant of $K_{\mathrm{d}} = 7.02 \times 10^{-2}$ mM (35). The on and off rates for this mutant are not listed, however, the authors do list the rates for other mutants. Queen-2m has a reported on rate of $k_{\mathrm{on}} = 2.7 \times 10^{-2}\,\mathrm{mM}^{-1}\mathrm{s}^{-1}$ and an off rate of $k_{\mathrm{off}} = 9.4 \times 10^{-2}\,\mathrm{s}^{-1}$ (4). Queen-37C has a reported on rate of $k_{\mathrm{on}} = 3.5 \times 10^{-2}\,\mathrm{mM}^{-1}\mathrm{s}^{-1}$ and an off rate of $k_{\mathrm{off}} = 1.7 \times 10^{-1}\,\mathrm{s}^{-1}$ (3). Based on these reported off rates, the bound time of an ATP to a probe is roughly,

$$t_{\mathrm{b}} = \frac{1}{k_{\mathrm{off}}} = \frac{1}{0.1\,\mathrm{s}^{-1}} \approx 10\ \mathrm{s}. \qquad [157]$$

Given that we image the aster every 20 s, the bound time is long. The distance the probe can travel during the bound time of ATP is on the order of

$$l_{\mathrm{b}} = \sqrt{D t_{\mathrm{b}}} = \sqrt{45\,\mu\mathrm{m}^2/\mathrm{s} \times 10\ \mathrm{s}} \approx 20\,\mu\mathrm{m}. \qquad [158]$$

This length scale is also long since the gradients we measure are also on the order of few $\times\, 10\,\mu$m. Thus, diffusing probes bound to ATP could contribute to a significant smoothing of gradients as bound probes diffuse.

To better understand this, we can write a model for the ATP bound probes over space and time. In this model, we take into account the diffusion of bound probes, the dissociation of the ATP to the probe, and the binding of an ATP to the probe,

$$\frac{\partial Q_{\mathrm{b}}(r,t)}{\partial t} = D_{\mathrm{Q}} \nabla^2 Q_{\mathrm{b}}(r,t) - k_{\mathrm{off}} Q_{\mathrm{b}}(r,t) + k_{\mathrm{on}} Q_{\mathrm{f}}(r,t) A_{\mathrm{f}}(r,t), \qquad [159]$$

where $Q_{\mathrm{b}}(r,t)$ is the concentration of the ATP bound probe, $D_{\mathrm{Q}}$ is the diffusion coefficient of the probe, $Q_{\mathrm{f}}(r,t)$ is the concentration of the free probe, and $A_{\mathrm{f}}$ is the free concentration of ATP. We turn to COMSOL to solve this partial differential equation.

| Model Comparisons | | |
| --- | --- | --- |
| Parameter | Schief Model | Our Model |
| $k_{\text{cat}}^{00}$ | $113.2$ s$^{-1}$ | $114.5$ s$^{-1}$ |
| $K_{\text{M}}^{00}$ | $28.1$ µM | $26.2$ µM |
| $K_{\text{i}}^{\text{P}}$ | $9$ mM | $8.9$ mM |
| $K_{\text{i}}^{\text{ADP}}$ | $34.6$ µM | $24.5$ µM |
| $K_{\text{i}}^{\text{ADP}\cdot\text{P}}$ | $95$ mM | $-714$ mM |
| $K_{\text{ii}}^{\text{P}}$ | $200$ mM | - |
| $K_{\text{ii}}^{\text{ADP}}$ | $23$ mM | - |
| $K_{\text{ii}}^{\text{ADP}\cdot\text{P}}$ | $30$ mM | - |

**Table S4. We compare the fitted parameters from our model versus the fitted parameters found by the Schief model. Overall the fits are in agreement with the exception of the Menten constant for the ADP and phosphate bound state.**

To gain intuition for the problem, we first make a few approximations. First, we approximate the total concentration of probe, $Q$ to be homogeneous across the experiment and conserved throughout time. Due to the ratiometric nature of this probe, any spatial features due to elevated concentrations of the probe are eliminated. We substitute $Q_{\rm f}(r,t) = Q - Q_{\rm b}(r,t)$ in Equation 159. Additionally, we take the concentration of ATP to be much larger than the concentration of the probe, our experiment uses about 500 $\mu$M ATP and 2.8 $\mu$M probe, such that if all probes were bound to an ATP, the impact on the free ATP distribution would be negligible, $A(r,t) \approx A_{\rm f}(r,t)$. Lastly, we take the ATP distribution to be in a steady state gradient to get a sense for how the timescale of blurring created by the probes.

$$\begin{aligned}\frac{\partial Q_{\rm b}(r,t)}{\partial t} &= D_{\rm Q}\nabla^2 Q_{\rm b}(r,t) - k_{\rm off}Q_{\rm b}(r,t) + k_{\rm on}\left(Q - Q_{\rm b}(r,t)\right)A(r),\\ &= D_{\rm Q}\nabla^2 Q_{\rm b}(r,t) - (k_{\rm off} + k_{\rm on}A(r))Q_{\rm b}(r,t) + k_{\rm on}QA(r)\end{aligned} \qquad [160]$$

We turn to COMSOL to numerically solve this partial differential equation. For our parameters, we take $D_{\rm Q} = 45\,\mu{\rm m}^2/{\rm s}$, as estimated using equation 33, $K_{\rm D} = 70\,\mu$M (3), $k_{\rm off} = 0.1$ s$^{-1}$(3, 4), $k_{\rm on} = \frac{k_{\rm off}}{K_{\rm D}} = 10^{-3}\,\mu{\rm M}^{-1}{\rm s}^{-1}$, and $Q = 2.5\,\mu$M. We set the ATP profile to be a Gaussian distributed following the form

$$A(r) = A_0\left(1 - w\exp\left(\frac{-r^2}{\lambda^2}\right)\right), \qquad [161]$$

where $A_0$ is initial and maximal ATP concentration in the experiment, $w$ is the fractional reduction of the Gaussian amplitude from $A_0$, and $\lambda$ is the decay length of the Gaussian profile. For our initial condition, we set all of the probes to be unbound with $Q_b(r,0) = 0$ and we impose a no flux boundary condition.

As a start, we run a simulation exploring the binding in the first 20 seconds, the interval between image acquisitions. In Figure S77, we show both (A) the inputted ATP distribution and (B) the bound probe response over time. It appears that it takes under 20 seconds for the bound probe concentration to reach an initial steady state. To translate the probe distribution into a representative measurement of ATP concentrations, we need a calibration curve. Given Langmuir binding, we expect

$$Q_{\rm b}(r,t) = Q\frac{\frac{A(r,t)}{K_{\rm D}}}{1 + \frac{A(r,t)}{K_{\rm D}}}. \qquad [162]$$

As a sanity check, we confirm that when taking a homogeneous constant value of ATP, $A(r,t) = A_0$, and simulating Equation 160 for several $A_0$ values, the steady state probe concentration at each $A_0$ falls on the curve defined by Equation 162.

For each $A_0$ value, we plot the resultant bound concentration of probe averaged across the specified geometry and indeed find agreement to Equation 162. We plot the calibration curve in Figure S78. We now turn resulting bound probe plots into "measured" ATP plots by inverting Equation 162 yielding,

$$A(r,t) = \frac{Q_{\rm b}(r,t)K_{\rm D}}{Q - Q_{\rm b}(r,t)}. \qquad [163]$$

Applying this transformation on the data in Figure S77(B), we report the "measured" ATP in this simulation in Figure S77(C).

We find that the "measured" ATP is greater in the center of the depletion region than the prescribed ATP distribution, implying that probes that diffuse with ATP bound effectively blur the curve we observe. We next explore the effects of modulating the well depth of the prescribed ATP gradient. In Figure S79 we show the "measured" ATP when making the initial ATP well deeper.

We now explore how different the blurring effect of bound probe diffusion is as compared with dynamically changing ATP resulting from a dynamic motor profile. Using a sigmoid based motor profile (Figure S80(A)), we simulate the resulting ATP curves based on our reaction-diffusion equation. Then using Equation 159, we determine the bound probe concentrations and convert to a "measured" ATP profile via Equation 163(Figure S80(B). We find that while blurring does occur at intermediate times in aster formation, it is very minimal compared to the simulated "true" ATP profiles. We see that the blurring always makes gradients smoother than they actually are, and proceed in this study without further correction for this effect.

**A. Binding Model of Diffusion.** ATP in our system can either be bound to the fluorescent ATP probe, at a concentration $A_b$, or free, with a concentration $A_f$. Let us begin this section by determining the fraction of bound ATP, $p_b$. The binding of ATP to the probe QUEEN-7μ is described by the kinetic equation

$$[\text{ATP}] + [\text{QUEEN-7μ}] \underset{k_{\rm off}}{\overset{k_{\rm on}}{\rightleftharpoons}} [\text{ATP}\cdot\text{QUEEN-7μ}]. \qquad [164]$$

The kinetics of binding is described by

$$\frac{d[\text{ATP}\cdot\text{QUEEN-7μ}]}{dt} = k_{\rm on}[\text{ATP}][\text{QUEEN-7μ}] - k_{\rm off}[\text{ATP}\cdot\text{QUEEN-7μ}]. \qquad [165]$$

In steady state, we find

$$\frac{[\text{ATP}][\text{QUEEN-7μ}]}{[\text{ATP}\cdot\text{QUEEN-7μ}]} = \frac{k_{\rm off}}{k_{\rm on}} = K_{\rm d}. \qquad [166]$$

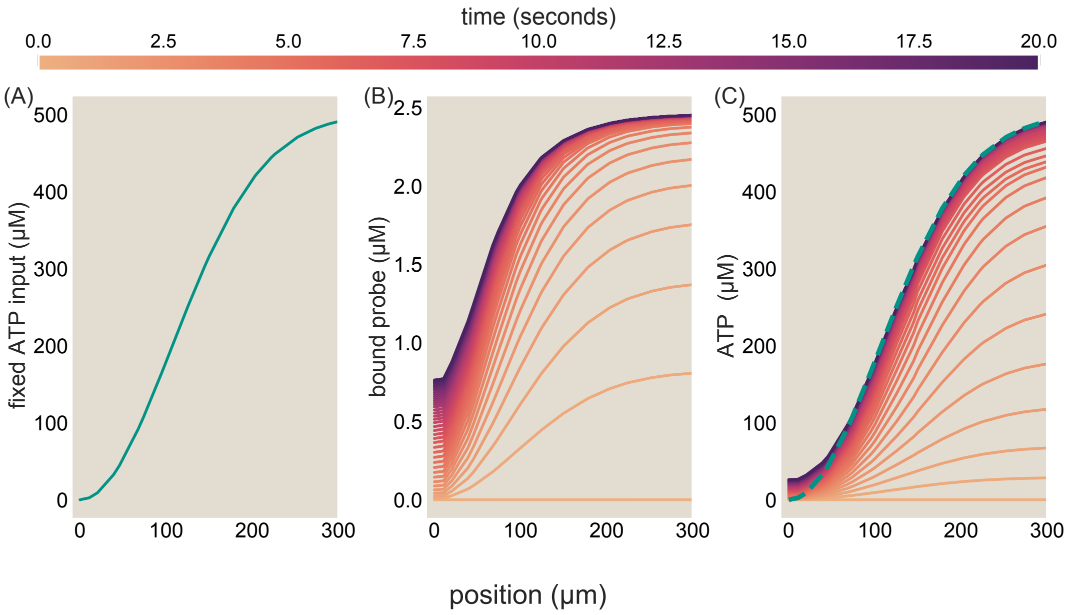

**Fig. S77. Binding curve evolution to steady state for a fixed ATP profile.** (A) Defined fixed ATP profile as described by Equation 161, where $A_0 = 500\,\mu\mathrm{M}$, $w = 1$, and $\lambda = 150\,\mu\mathrm{m}$. (B) Simulated results for the bound concentration of ATP probes, $Q_{\mathtt{b}}(r,t)$, with an initial condition of $Q_{\mathtt{b}}(r,0) = 0$. (C) "Measured" ATP concentration of ATP by the simulated probe distribution. These profiles are found by applying Equation 162 to the $Q_{\mathtt{b}}(r,t)$ profiles simulated in (B) and solving for $A(r,t)$. The dashed teal line is the defined fixed ATP profile, same as the line shown in (A).

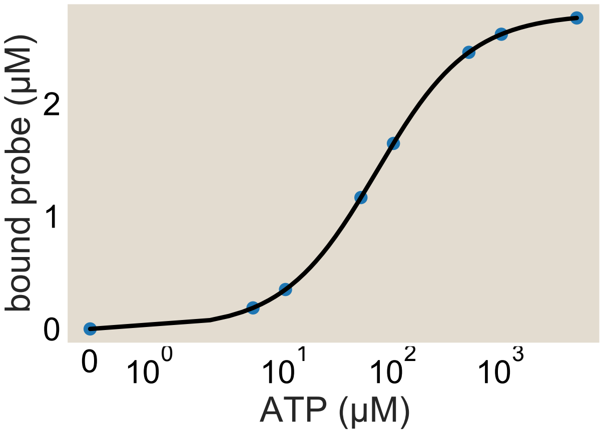

**Fig. S78. Calibration of bound probe concentration to ATP concentration.** For homogeneously defined ATP profiles at varying concentrations, we find the steady state concentration of bound probes in our simulation. We plot these values as a function of ATP concnetration as the blue dots. The black curve is the Michaelis-Menten curve for the probe as defined in Equation 162, where we take the probe's Menten constant to be $K_{\mathrm{d}} = 70\,\mu\mathrm{M}$ and the total probe concentration to be $Q = 2.8\,\mu\mathrm{M}$.

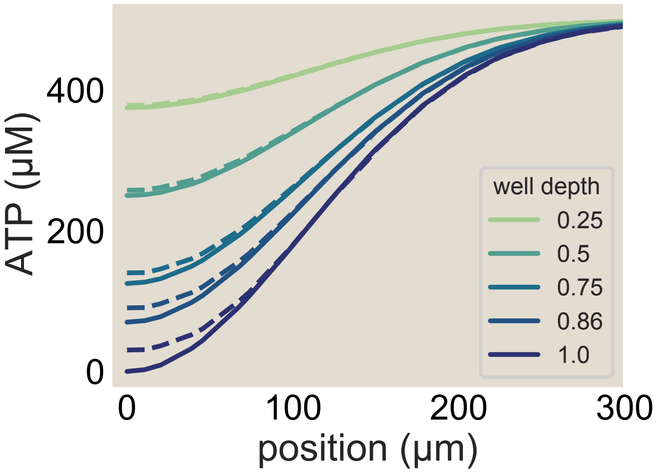

**Fig. S79. More blurring due to bound probe diffusion occurs for larger well depths.** We modulate the well depth of the inputted fixed ATP profiles (solid lines), while keeping the gaussian radius constant. We sweep over a variety of depths notably plotting a depth such that the lowest value of the ATP is equivalent to the probe Michaelis-Menten constant, $K_\mathrm{d}/A_0 = 0.86$ for our probe. The dashed lines are the "measured" ATP values determined by using Equation 163 on the simulation for $Q_\mathrm{b}(r,t)$. Larger well depth experience higher "measured" ATP concentrations than the defined ATP in the center of the well. The "measured" curves are taken from simulations that have run for $t = 4000$ seconds, well past the time to reach steady state.

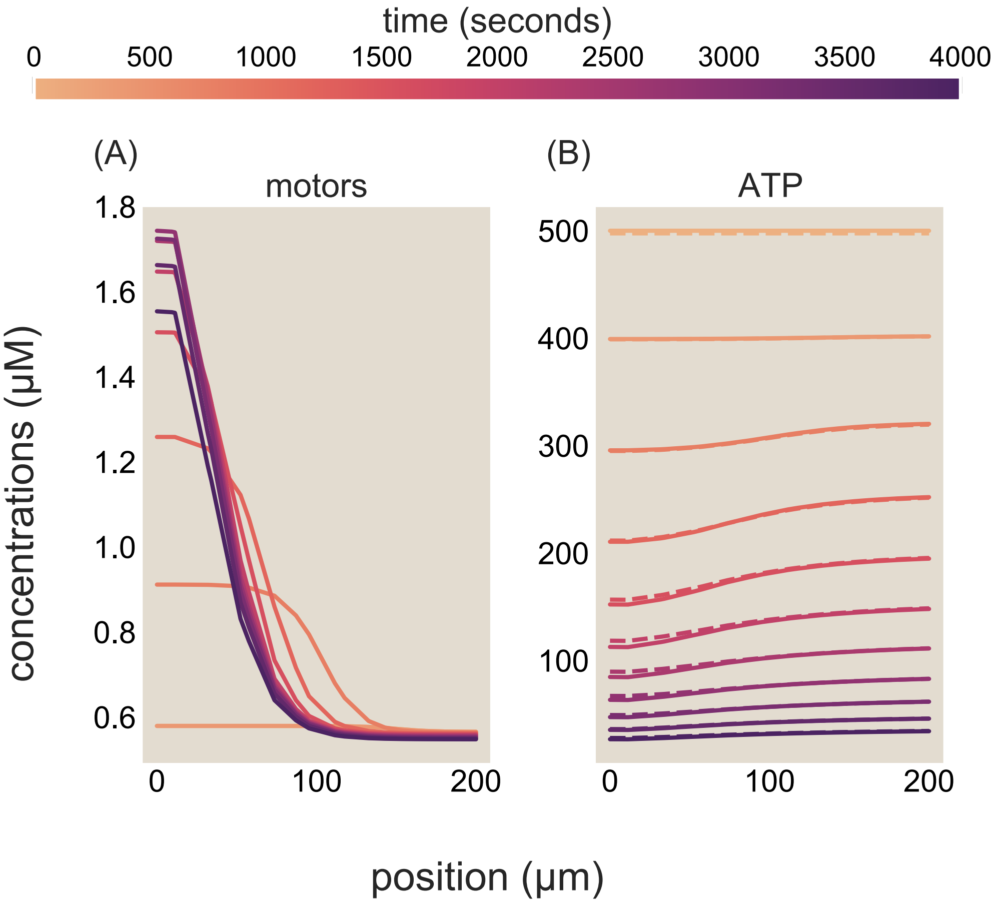

**Fig. S80. The result of blurring on dynamic ATP profiles is small.** (A) The prescribed motor profile based on a sigmoidal trial function. (B) The resultant ATP profiles given the motor distributions are plotted as solid lines. The bound probe distributions are converted to ATP units using the calibration curve specified by Equation 163 and then plotted as dashed lines.

As reported by Yaginuma et al. (4), the dissociation constant of ATP to the QUEEN-7µ ATP probe is $K_\mathrm{d} = 70\,\mu$M.

This equation is of the form of a simple binding model of a ligand to a receptor. For brevity, we write $[\mathrm{ATP}] = [L]_\mathrm{free}$, $[\text{QUEEN-7µ}] = [R]_\mathrm{free}$, and $[\mathrm{ATP}\cdot\text{QUEEN-7µ}] = [C]$, where $[L]_\mathrm{free}$ is the concentration of free ATP, $[R]_\mathrm{free}$ is the concentration of free QUEEN-7µ, and $[C]$ is the concentration of bound ATP-QUEEN-7µ complex. We denote the total ATP concentration by $[L]_\mathrm{tot}$ and the total QUEEN-7µ concentration by $[R]_\mathrm{tot}$. Thus,

$$[R]_\mathrm{tot} = [R]_\mathrm{free} + [C], \qquad [167]$$

and

$$[L]_\mathrm{tot} = [L]_\mathrm{free} + [C]. \qquad [168]$$

Assuming that ATP binding to QUEEN-7µ equilibrates quickly as ATP is depleted, we can compute the fraction of ATP bound to QUEEN-7µ,

$$p_b = \frac{[C]}{[L]_\mathrm{tot}}. \qquad [169]$$

The binding equilibrium is

$$K_\mathrm{d} = \frac{[L]_\mathrm{free}[R]_\mathrm{free}}{[C]} = \frac{([L]_\mathrm{tot} - [C])\,([R]_\mathrm{tot} - [C])}{[C]}. \qquad [170]$$

Rearranging gives the quadratic equation

$$[C]^2 - ([L]_\mathrm{tot} + [R]_\mathrm{tot} + K_\mathrm{d})\,[C] + [L]_\mathrm{tot}[R]_\mathrm{tot} = 0. \qquad [171]$$

Solving for the concentration of bound ATP-QUEEN-7µ complex gives

$$[C] = \frac{[L]_\mathrm{tot} + [R]_\mathrm{tot} + K_\mathrm{d} - \sqrt{([L]_\mathrm{tot} + [R]_\mathrm{tot} + K_\mathrm{d})^2 - 4[L]_\mathrm{tot}[R]_\mathrm{tot}}}{2}. \qquad [172]$$

The minus sign is the physical root because the plus sign would give a bound-complex concentration larger than the available reservoir. The fraction of ATP bound to QUEEN-7µ is therefore

$$p_b = \frac{[C]}{[L]_\mathrm{tot}} = \frac{[L]_\mathrm{tot} + [R]_\mathrm{tot} + K_\mathrm{d} - \sqrt{([L]_\mathrm{tot} + [R]_\mathrm{tot} + K_\mathrm{d})^2 - 4[L]_\mathrm{tot}[R]_\mathrm{tot}}}{2[L]_\mathrm{tot}}. \qquad [173]$$

We now evaluate the range of $p_b$ values that occur during the course of our experiment. At the beginning of the experiment, we pipette 500 $\mu$M of ATP into the system. Throughout the experiment the concentration of the ATP reporter remains constant at 1.5 $\mu$M. At the start of the experiment, we find

$$p_b(t=0) = \frac{571.5\,\mu\mathrm{M} - \sqrt{(571.5\,\mu\mathrm{M})^2 - 4(500)(1.5\,\mu\mathrm{M})}}{2(500\,\mu\mathrm{M})} \approx 0.003. \qquad [174]$$

where we have used the $K_\mathrm{d}$ value from (4), after ATP has been hydrolyzed, the limit $[L]_\mathrm{tot} \to 0$ gives

$$p_b \to \frac{1.5\,\mu\mathrm{M}}{1.5\,\mu\mathrm{M} + 70\,\mu\mathrm{M}} \approx 2 \times 10^{-2}. \qquad [175]$$

We plot the full trajectory of the bound fraction and the bound concentration of ATP in Figure S81. We observe that over the course of the experiment, the fraction of ATP bound to QUEEN-7µ is always small and varies little. We therefore treat $p_b$ as a small constant in space and time for the rest of this section. Let us now explore if the diffusion of ATP is altered by binding to the probe.

We need two coupled partial differential equations to model the evolution of ATP through space and time. In a given time step, the concentration of bound ATP at a given position can change if the ATP and probe diffuse, if bound ATP unbinds from the probe, or free ATP binds to the probe. We formalize this as

$$\frac{\partial A_b(r,t)}{\partial t} = D_b \nabla^2 A_b(r,t) + k_\mathrm{on} A_f(r,t) - k_\mathrm{off} A_b(r,t), \qquad [176]$$

where $D_b$ is the diffusion constant of ATP bound to the probe and $k_\mathrm{on}/k_\mathrm{off}$ are the on/off rates of the ATP probe respectively. Similarly, we write a partial differential equation for free ATP. In a given time step and position, the concentration of free ATP can change if ATP diffuses, bound ATP unbinds from the probe, free ATP binds to the probe, or a motor protein hydrolyzes ATP. We formalize this as

$$\frac{\partial A_f(r,t)}{\partial t} = D_f \nabla^2 A_f(r,t) - k_\mathrm{on} A_f(r,t) + k_\mathrm{off} A_b(r,t) - \Gamma\left(r,t,A_f,m\right) A_f(r,t), \qquad [177]$$

where $D_f$ is the diffusion constant of ATP and $\Gamma\left(r,t,A_f,m\right)$ is a function of the hydrolysis rate of ATP, which is dependent on $m(r,t)$, the motor concentration through space and time. The presence of the motor function introduces a third partial

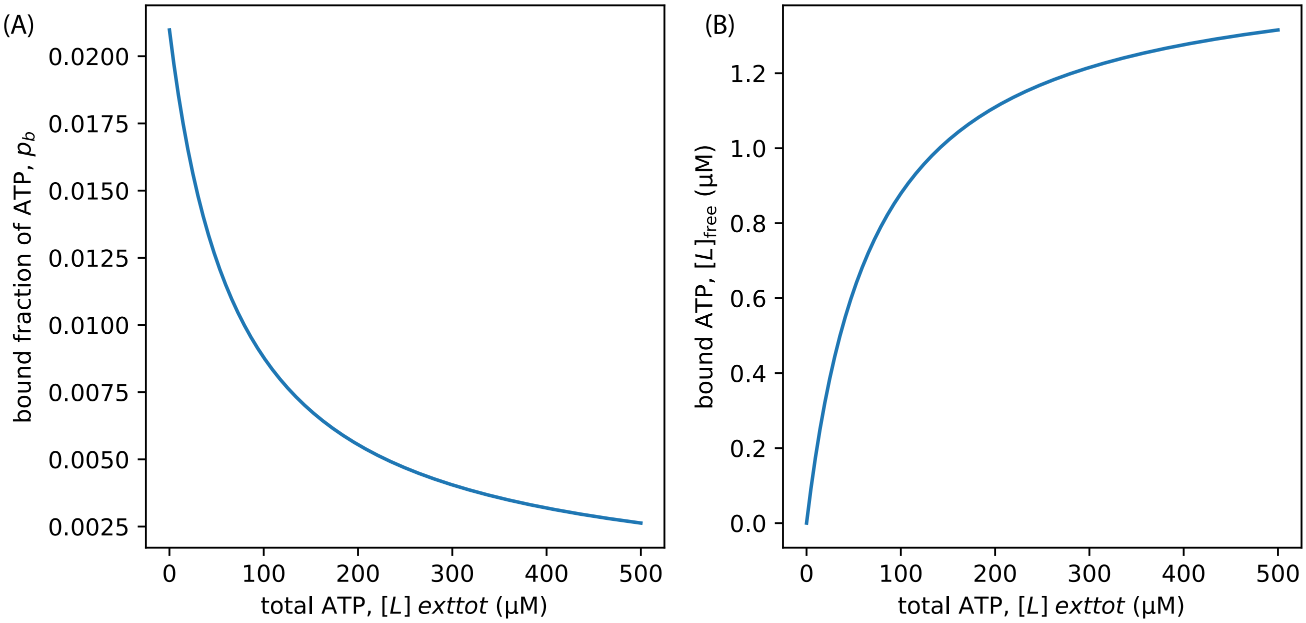

**Fig. S81. The amount of ATP bound to the probe is small throughout the course of the experiment.** (A) The bound fraction of ATP , as computed with equation 173, and (B) the bound concentration of ATP are plotted as a function of total ATP concentration.

differential equation to consider, the evolution of motors. The concentration of motors at a given time and position can change if motors diffuse or walk along microtubules (advection), which we will write as

$$\frac{\partial m(r,t)}{\partial t} = D_m \nabla^2 m(r,t) + \nabla \cdot v(r) m(r,t), \quad [178]$$

where $D_m$ is the diffusion constant and $v(r)$ is the velocity of the motor proteins.

In this section, we aim to understand the role of binding on diffusion, so we next write an equation for the total ATP concentration, $A$, which is the sum of the free and bound species, $A(r,t) = A_f(r,t) + A_b(r,t)$. By linearity of differentiation, the partial derivative of the total ATP with time is the sum of the partial derivative of each species with time, $\frac{\partial A(r,t)}{\partial t} = \frac{\partial A_f(r,t)}{\partial t} + \frac{\partial A_b(r,t)}{\partial t}$. Thus, we write an equation for the total ATP by adding equations 176 and 177 which yields

$$\frac{\partial A(r,t)}{\partial t} = D_f \nabla^2 A_f(r,t) + D_b \nabla^2 A_b(r,t) - \Gamma(r,t,A_f,m) A_f(r,t) \quad [179]$$

Here, we still have terms containing $A_f$ and $A_b$. Let us now write equation 179 only in terms of the total ATP and the fraction of bound ATP, $p_b$, which we solved for in equation 173. We now express $A_b(r,t) = p_b A(r,t)$ and $A_f(r,t) = \big(1-p_b\big)A(r,t)$. Thus, equation 179 can be written as

$$\frac{\partial A(r,t)}{\partial t} = D_f\big(1-p_b\big)\nabla^2 A(r,t) + D_b p_b \nabla^2 A(r,t) - \Gamma(r,t,p_b,m) A(r,t)\big(1-p_b\big). \quad [180]$$

Again by linearity of differentiation, we can rearrange the equation to

$$\frac{\partial A(r,t)}{\partial t} = \underbrace{\big(D_f - p_b(D_f - D_b)\big)\nabla^2 A(r,t)}_{\text{Diffusion Term}} - \underbrace{\Gamma(r,t,p_b,m) A(r,t)\big(1-p_b\big)}_{\text{Reaction Term}}. \quad [181]$$

From this form, we can infer that the value of the diffusion constant is a sliding scale based on the fraction of ATP that is bound. If no ATP is bound, $p_b = 0$, then the diffusion term simplifies to $D_f\nabla^2 A$, whereas if all the ATP is bound, $p_b = 1$, then the diffusion term becomes $D_b\nabla^2 A$. In equations 174 and 175, we estimated the values of $p_b$ in the range of experimental conditions and found that $p_b$ is very small. So, it is appropriate to treat the diffusion constant of the total ATP as the diffusion constant of free ATP, $D_f - p_b(D_f - D_b) \approx D_f$. Thus, we conclude that the binding kinetics of ATP to the probe are not relevant to the diffusion of the total ATP field, only the ATP diffusion constant contributes.

#### 10. Comparison to other biological gradients.

**A. Simple tools for comparing gradients.** By what measures should gradients be regarded as steep or shallow, functionally meaningful or inconsequential? To contextualize the concentration gradients uncovered in this work found in contracting asters, we discuss three simple operational notions of how large a gradient is. These numerical summaries capture distinct and complementary aspects of how a gradient might matter to organisms.

- **Absolute concentration gradient,** $\left|\frac{\partial c}{\partial r}\right|$: How does the absolute number of molecules at a position (per unit volume) change per linear increment in spatial position? This primordial number, absent further normalization by any reference concentrations or spatial scales, most literally expresses the size of a gradient. This is reasonably approximated by the representative drop in concentration $\Delta c \equiv |c(r+\Delta r) - c(r+\Delta r)|$ over a characteristic lengthscale $\Delta r$ spanning a biological region of interest where the gradient functionally operates.

- **Fractional concentration gradient,** $\left|\frac{\partial \log c}{\partial r}\right| = \left|\frac{1}{c}\frac{\partial c}{\partial r}\right|$: By what fraction does the number of molecules (per unit volume) change per linear increment in spatial position? This notion measures concentration gradients in units of the typical ambient concentration $c$. The reciprocal of this quantity reflects the typical distance over which the gradient elicits a concentration change of order the ambient concentration.

- **Spatial log-derivative of concentration gradient,** $\left|\frac{\partial \log c}{\partial \log r}\right| = \left|\frac{\partial c/\partial r}{c/r}\right|$: What is the fractional change in the abundance of molecules per fractional change in spatial position? This logarithmic derivative (namely the slope of a concentration gradient when plotted on a log-log plot) is a unitless handle on the scale of gradients.

Note that the comparative concentration scales $c$ and lengthscales $r$ used to inform these normalizations may be most informative when taken not as literal local values, but instead more characteristic scales that more fully express each context's physics. For instance, in the context of ATP consumed by motors, one can take the Michaelis constant $K_M$ setting the concentration of ATP required for motors to step at meaningful fractions of their maximal rates. (This is a comparison mentioned in the main text for asters: the manifested changes in ATP concentration are several multiples of characteristic $K_M$ values).

Another natural normalization scale emerges in the example of positional information in body plans: the informational value of a concentration profile $g(x)$ with respect to space $x$ can be related to the average gradient $dg/dx$ measured in units of the local variability $\sigma_g(x)$ in the concentration over positions, namely depending on $\int\ dx\ p(x)\ \frac{1}{\sigma_g(x)}\left|\frac{dg}{dx}\right|$, where $p(x)$ is the probability distribution over space (36). This connection argues for assessing the functionality of a gradient by its size relative to a separate concentration scale, $\sigma_g$, expressing the variability of profiles.

Similarly, the relevant lengthscale used to normalize gradients may not be a literal position $r$ but instead another scale provided by separate physics. For instance, in the example of eukaryotic or bacterial chemotaxis, a cell's own lengthscale $L$ conspires with the magnitude of an environmental gradient to set whether the change in concentration over its own length can be substantial enough to sense gradients instantaneously over their length (37). Or else, the relevant $L$ can be the required run length in run-and-tumble motion that permits a minimum accumulation of a concentration gradient for sensing. The takeaway urged by these examples is that the ultimate consequences of a gradient are refracted by intrinsic normalizations.

Armed by these conceptual frames and provisos, we now detail a small set of evocative biological gradients for comparison to the scales registered in ATP in the present work's asters. Our aim is far from encyclopedic. Rather, we explore these few numbers to build a rough feeling for scales of gradients that manifest in biology.

We summarize numerical estimates from the following analysis in Table S5. By most measures, the ATP gradients explored in this present work are comparable to or significantly larger than other biological gradients.

| | **this work's contracting asters (§E)** | chemotactic gradients (§B) | Pom1 in yeast (§C) | Bicoid in *Drosophila* (§D) |
|---|---|---|---|---|
| absolute gradient, $\left\lvert\frac{\partial c}{\partial r}\right\rvert \approx \frac{\Delta c}{\Delta r}$ | $\approx$ (few $\times 10^2$) - $10^3$ $\frac{\text{molecules}/\mu\text{m}^3}{\mu\text{m}}$ | $\approx$ few $\times\, 10 \frac{\text{molecules}/\mu\text{m}^3}{\mu\text{m}}$ | $\approx$ few $\frac{\text{molecules}/\mu\text{m}^3}{\mu m}$ | $\approx$ few $\times\, 10^{-1}$ $\frac{\text{molecules}/\mu\text{m}^3}{\mu m}$ |
| fractional gradient, $\left\lvert\frac{1}{c}\frac{\partial c}{\partial r}\right\rvert \approx \frac{\Delta c}{c}$ | $\approx$ few $\times\, 10^{-1}$ $\mu\text{m}^{-1}$ | $\approx$ few $\times\, 10^{-3}$ $\mu\text{m}^{-1}$ | $\approx$ few $\times\, 10^{-1}$ $\mu\text{m}^{-1}$ | $\approx$ few $\times\, 10^{-3}$ $\mu\text{m}^{-1}$ |
| spatial log derivative, $\left\lvert\frac{\partial c/\partial r}{c/r}\right\rvert \approx \frac{\Delta c}{c}$ | $\approx$ few $\times\, 10^{-1}$ | $\approx$ few $\times\, 10^{-2}$ | $\approx 1$ | $\approx 10^{-1}$ |

**Table S5. Summary of order of magnitude estimates on the sizes of different biological gradients, evaluated by complementary measures of gradient size.**

**B. Scale of measured gradients navigated by bacterial chemotaxis.** One of the most famous gradient examples in all of biology is that of bacterial chemotaxis. The phenomenon of interest is that a bacterial cell finds itself in a spatially inhomogeneous field of chemoattractant, such as chemical proxies for carbon sources. Specifically, these cells detect such gradients and use them to "decide" which way to swim by adjusting the frequency of tumble events which reorder their swim direction.

In foundational experiments like those of Sourjik and Berg (38, 39), a capillary tube filled with chemoattractant at a concentration of $c_0 \approx 1$ mM is positioned amidst bacteria. This pipette tip has radius of order $R = 10$ $\mu$m. The bacteria they observe are at a distance of approximately $r \approx 0.6$ mm away from the capillary tube. We may idealize the concentration profile established by the pipette as a source in a free homogenous medium as that corresponding to the steady-state of a spherical source in an empty bath. Specifically, when $c(r,t)$ evolves according to the diffusion equation $\frac{\partial c}{\partial t} = D\nabla^2 c$ expressed in spherical coordinates, with the boundary conditions $c(r = R) = c_0$ and $c(r \to \infty) = 0$, the classical power law solution profile $c_{\text{ss}}(r) = -\frac{A}{r} + B$ following at steady-state (for some constants $A$ and $B$) adopts the form,

$$c_{\text{ss}}(r) = c_0 \frac{R}{r}. \quad [182]$$

This expresses an absolute gradient of,

$$\frac{dc}{dr} = -c_0 \frac{R}{r^2}, \quad [183]$$

It is also interesting to express the absolute gradient expressed in Eq. 183 in normalized senses. The fractional concentration gradient is locally,

$$\frac{d \log c}{dr} = \frac{1}{c(r)} \frac{dc}{dr} = \frac{-c_0 R/r^2}{R c_0/r} = 1/r, \quad [184]$$

giving a fractional change in concentration per fractional change in position of,

$$\frac{d \log c}{d \log r} = \frac{\partial c/\partial r}{c/r} = 1. \quad [185]$$

The above expressions give local descriptions of the gradients (which can be evaluated to describe the gradient at a particular spatial position $r$). However, it is useful to summarize the aggregate scale of such gradients in an accumulated sense. To do so, we compute the average gradient $\approx \frac{\Delta c}{\Delta r}$ manifested over a distance $\Delta r \approx 30$ $\mu$m; this is a revealing lengthscale as it corresponds to the typical displacement of an *E. coli* bacterium over one of its one-to-two second ballistic runs (40), while traveling at ten to a few tens of micron per second. (Indeed, one might speculate that these run lengths could be tuned to be sufficiently long to satisfactorily sample concentration gradients typical of environments.) At the start of such a run, bacteria located at $r \approx 0.6$ mm $= 600$ $\mu$m experience a concentration of,

$$c(r = 600\ \mu\text{m}) = c_0 \frac{R}{r} = (10^3\ \mu\text{M}) \frac{10\ \mu\text{m}}{600\ \mu\text{m}} \approx 16.7\ \mu\text{M}, \quad [186]$$

and at the end of the run ($r + \Delta r = 630$ $\mu$m) they experience,

$$c(r = 630\ \mu\text{m}) = c_0 \frac{R}{r + \Delta r} = 15.9\ \mu M. \quad [187]$$

This corresponds to an average absolute gradient of

$$\frac{\Delta c}{\Delta r} \approx \frac{0.8\ \mu\text{M}}{30\ \mu\text{m}} = 0.027\ \frac{\mu\text{M}}{\mu\text{m}} \tag{188}$$

$$\approx \text{few} \times 10^{-2}\ \frac{\mu\text{M}}{\mu\text{m}} \tag{189}$$

$$\approx \text{few} \times 10 \frac{\text{molecules}/\mu\text{m}^3}{\mu\text{m}}. \tag{190}$$

A rough estimate of the fractional concentration gradient evaluated on this lengthscale is then,

$$\frac{1}{c(r)}\frac{\Delta c}{\Delta r} \approx \frac{1}{16.7\ \mu\text{M}}\left(0.027\ \frac{\mu\text{M}}{\mu\text{m}}\right) \tag{191}$$

$$\approx 1.6 \times 10^{-3}\ \mu\text{m}^{-1}. \tag{192}$$

Last, a rough estimate of the gradient estivated as a log-derivative over this lengthscale is,

$$\frac{\Delta r}{c(r)}\frac{\Delta c}{\Delta r} \approx (30\ \mu\text{m})\left(1.6 \times 10^{-3}\ \mu\text{m}^{-1}\right) \tag{193}$$

$$\approx 4.8 \times 10^{-2} \tag{194}$$

$$\approx \text{few} \times 10^{-2}. \tag{195}$$

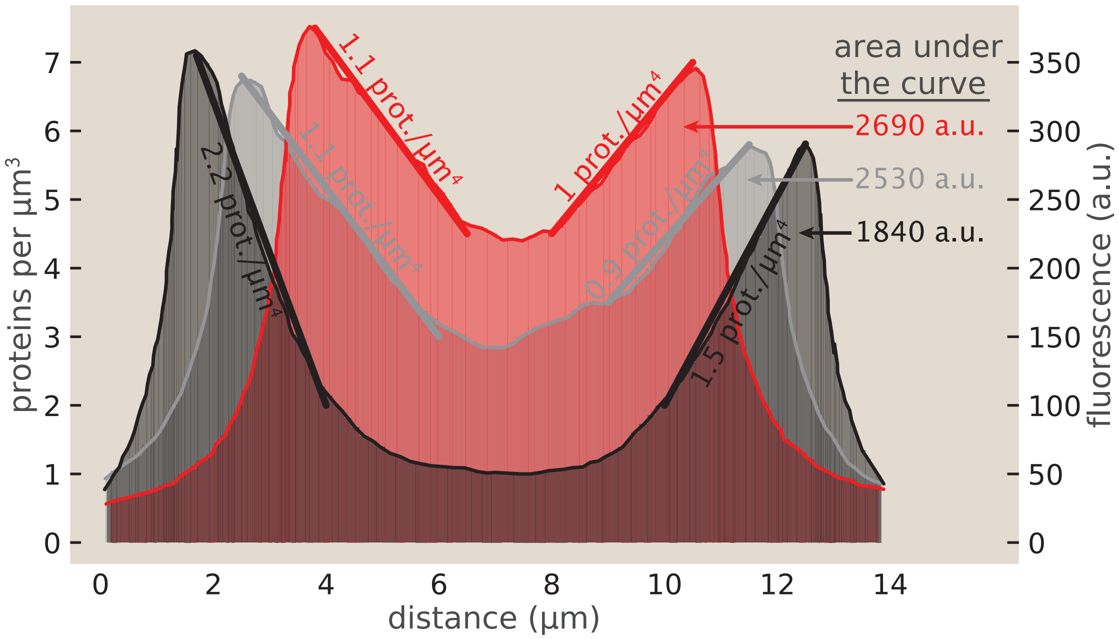

**Fig. S82.** The gradient of Pom1 in fission yeast. Three representative fluorescence profiles are shown corresponding to different stages during the cell cycle. Given a previous measurement of cytoskeletal and motor proteins from fission yeast (41), we converted arbitrary fluorescence units into protein counts. Plot adapted from J. Moseley et al., Nature 857:860, 2009, ( 18).

**C. Scale of the gradient in protein kinase, Pom1, in fission yeast.** As discussed in the main text, one point of context for the scales of gradients we considered is the gradient of the protein kinase Pom1 in fission yeast. As part of the signaling pathway that controls entry to mitosis, Pom1 localizes in the poles of the yeast cell (42). As shown in the main text, we were interested in characterizing the magnitude of different biological gradients in units of $M/\mu m$. Our goal was to quantify gradients across scales ranging from intracellular up to entire organisms. As a single-cell example, the fission yeast cell cycle offered an interesting opportunity. It has been shown that fission yeast develops a gradient of the protein kinase Pom1 as part of cell division signaling (42). We aim to compare the scale of this biological gradient with the ATP gradients we measure. Figure S82 is adapted from (42) where the authors show Pom1 localizes at the cell poles. Their results are plotted in arbitrary units which we attempt to convert to concentration units using a census of protein counts conducted by Wu and Pollard (41). Protein kinases in Wu and Pollard's study have approximately 5000 copies per cell with a cell volume of $92\,\mu m^3$. The microscopy images associated with Figure S82 suggest approximating the cell to be a cylinder with a diameter of approximately

3.5 $\mu$m. Thus the volume of the cell for the three different measurements we extracted from Figure S82 are 73 $\mu$m$^3$, 94 $\mu$m$^3$, and 111 $\mu$m$^3$ associated with the reported cell lengths 7.6 $\mu$m, 9.8 $\mu$m, and 11.5 $\mu$m respectively. Using PlotDigitizer (43), we extracted the data points in Figure 4b of (42) and approximate the integral of fluorescence across the cell with thin rectangles. What is missing is a conversion factor between fluorescence in “arbitrary units (au)” and number of proteins. To that end, we convert the arbitrary units of fluorescence to concentration by summing up all the fluorescence as

$$\mathcal{N} \int_0^L I \, dl = \frac{p}{V}, \quad [196]$$

where $\mathcal{N}$ is the unknown calibration factor, $L$ is the length of the cell $I$ is the fluorescent intensity in arbitrary units, $p$ is the number of proteins per cell, and $V$ is the volume of the cell. Given these definitions, we approximate the calibration factor as

$$\mathcal{N} = \frac{p}{V \int_0^L I \, dl} = \frac{5000 \, \text{proteins}}{92 \, \mu\text{m}^3 \times 2500 \, \text{a.u.}} \approx 0.02 \, \frac{\text{proteins}}{\mu\text{m}^3 \cdot \text{a.u.}}, \quad [197]$$

where we used the integrated fluorescence of the 9.8 $\mu$m long cell. Applying the calibration factor allows us to now replot the original figure, but now with the concentration of Pom1 across the cell in proteins per $mu$m$^3$ units in Figure S82. Using this crude calibration, we estimate the maximum gradient by drawing a line from the peak to trough of the curve and finding the corresponding slope which was then used in the main text as part of our series of estimates of biological gradients.

Consulting the steepest spatial gradient in Fig. S82 (the black curve), we see that $c(r = 2\ \mu\text{m}) \approx 7.1$ molecules/$\mu$m$^3$ drops to $c(r = 4\ \mu\text{m}) \approx 1.7$ molecules/$\mu$m$^3$, namely a change $\Delta c \approx 5.7$ molecules/$\mu$m$^3$ over $\Delta r \approx 2\ \mu$m. Using these scales, we compute the following rough estimates of the gradient. The absolute gradient is approximately,

$$\begin{aligned} \frac{\Delta c}{\Delta r} &\approx \frac{5.7 \text{ molecules}/\mu\text{m}^3}{2\ \mu\text{m}} && [198] \\ &\approx 2.85 \frac{\text{molecules}/\mu\text{m}^3}{\mu m} && [199] \\ &\approx \text{few } \frac{\text{molecules}/\mu\text{m}^3}{\mu m}. && [200] \end{aligned}$$

The fractional gradient is approximately,

$$\begin{aligned} \frac{1}{c(r)} \frac{\Delta c}{\Delta r} &\approx \left( \frac{1}{7.1 \text{ molecules}/\mu\text{m}^3} \right) \left( 2.85 \frac{\text{molecules}/\mu\text{m}^3}{\mu m} \right) && [201] \\ &\approx 0.4\ \mu\text{m}^{-1} && [202] \\ &\approx \text{few} \times 10^{-1}\ \mu\text{m}^{-1}. && [203] \end{aligned}$$

The spatial log derivative over this lengthscale can be estimated as,

$$\begin{aligned} \frac{\Delta r}{c(r)} \frac{\Delta c}{\Delta r} &\approx (2\ \mu\text{m})(0.4\ \mu\text{m}^{-1}) && [204] \\ &\approx 0.8 && [205] \\ &\approx 1. && [206] \end{aligned}$$

**D. Scale of the Bicoid gradient in the fly.** As another example, take the transcription factor *Bicoid* that famously contributes to delivering the positional information to organize the fruit fly *Drosophila*’s body plan as shown in the main text’s Figure 4(D). This molecular gradient develops with high precision across embryos, and robustly adopts a characteristic profile of approximately,

$$[\text{Bicoid}](x) \approx (55\ nM) \exp \left[ -\frac{x}{100\ \mu m} \right],$$

where $x$ is the distance along the embryo’s anterior-posterior axis (44). This means that at half the length of the embryo, $\ell \approx 250\ \mu$m, concentration has dropped from $c(r = 0) \approx 55 \times 10^{-3} \mu$M to about $c(r = 250\ \mu\text{m}) \approx c(r = 0) \exp\left[-(250\ \mu\text{m})/(100\ \mu\text{m})\right] \approx 4.5 \times 10^{-3} \mu$M. These give an estimated absolute gradient of,

$$\begin{aligned} \frac{\Delta c}{\ell} &\approx \frac{50\ \times 10^{-3}\ \mu\text{M}}{250\ \mu\text{m}} && [207] \\ &\approx 2 \times 10^{-4}\ \mu\text{M}/\mu\text{m} && [208] \\ &\approx 0.2\ \frac{\text{molecules}/\mu\text{m}^3}{\mu\text{m}}. && [209] \end{aligned}$$

The fractional gradient is then estimated as,

$$\frac{1}{c(r=0)}\frac{\Delta c}{\ell} \approx \frac{1}{55\ \text{molecules}/\mu\text{m}^3}\left(0.2\ \frac{\text{molecules}/\mu\text{m}^3}{\mu\text{m}}\right) \tag{210}$$

$$\approx 3.6 \times 10^{-3}\ \mu\text{m}^{-1}, \tag{211}$$

and the spatial log-derivative over this lengthscale is roughly,

$$\frac{\ell}{c(r=0)}\frac{\Delta c}{\ell} \approx \frac{4.5\ \text{molecules}/\mu\text{m}^3}{55\ \text{molecules}/\mu\text{m}^3} \tag{212}$$

$$\approx 8 \times 10^{-2} \tag{213}$$

$$\approx 10^{-1}. \tag{214}$$

**E. Scale of this work's average ATP gradients.** In the cytoskeletal network asters explored in this work, the average gradient evaluated between $r = 200\ \mu$m (near each aster's edge), and $r = 100\ \mu$m, reflects gradient scales of order

$$\frac{\Delta c}{\Delta r} \approx \frac{\text{ATP}(r = 100\mu\text{m}, t) - \text{ATP}(r = 200\mu\text{m}, t)}{100\mu\text{m}} \tag{215}$$

$$\approx (0.5 - 1)\ \mu\text{M}/\mu\text{m} \tag{216}$$

$$\approx (500 - 1000)\ \frac{\text{molecules}/\mu m^3}{\mu m}. \tag{217}$$

as for example shown across replicate asters in §3,Fig. S31. Taking typical ambient concentrations of order $ATP \approx 150\ \mu$M when these gradients have developed, this suggests a fractional concentration gradient of,

$$\frac{1}{c(r=200\mu\text{m})}\frac{\Delta c}{\Delta r} \approx \frac{1}{150\ \mu M}(0.5-1)\ \mu\text{M}/\mu\text{m} \tag{218}$$

$$\approx (3.3 \times 10^{-3}) - (6.7 \times 10^{-3})\ \mu\text{m}^{-1}, \tag{219}$$

and a spatial log derivative of order,

$$\frac{\Delta c}{c} \approx \frac{50\ \mu\text{M}}{150\ \mu\text{M}} \tag{220}$$

$$\approx 1/3 \tag{221}$$

$$\approx \text{few} \times 10^{-1}. \tag{222}$$

#### 11. Comparison to other (bulk) measurements of biological dissipation

Here, we assemble a small group of relevant scales of biological dissipation published elsewhere that enjoy interesting comparison with our measurements of dissipation in the contractile active matter system in this work. The spirit of this modest comparison is two-fold. First, naturalistically, it is interesting to ask how different corners of the biological world are powered by different scales of dissipation. Second, methodologically, it is edifying to ask how extremely different means of measurement—such as bulk calorimetry, oxygen turnover, chemostat measurements of sugar depletion, etc.—cohere (or do not) in the impressions they capture of such numbers. (Precisely assessing such coherence is a valuable activity for our community in a broader sense: understanding which numbers disagree often catalyzes wider discovery, as for instance in Perrin's comparative measurements of Avogadro's constant (45).)

In general, these comparative numbers suggest that the power expenditures disclosed by our ATP consumption rate measurement scheme in asters fall in the vicinity of those measured in other cellular contexts by completely different means, but with rich, revealing, system-specific differences.

Below we discuss assumptions to convert these numbers to common units for this comparison. Throughout this brief discussion, we recall the following characteristic scales. Under standard biochemical conditions hydrolyzing ATP yields a free energy $\Delta G \approx 50$ kJ/mol (or $\approx 20\ k_BT$/ATP) (46, 47), making hydrolysis of one ATP equivalent to $\approx (8 \times 10^{-20})$ J $\approx 10^{-19}$ J at $T = 300$ K, affirming the natural scale that 1 ATP/s corresponds to $10^{-19}\ W$ (namely a tenth of an attowatt). In addition we recall that 1 $\mu$M $\approx 602$ molecules/$\mu$m$^3$ underlying the operational rule of thumb that 1 $\mu$M $\approx 10^3$ molecules/$\mu$m$^3$.

**A. This work's aster contractile measurements.** As shown in Fig. S32, this work resolved total aster powers varying between $(3-8) \times 10^8$ ATP/s/aster. Assuming, as our imaging geometry suggests, an aster's dissipation stems from a cylindrical measurement volume of height $h = 70\ \mu$m and the aster has an initial radius of $r_0 \approx 170\ \mu$m when sufficiently formed to allow tracking of its boundary and dissipation, then the initial aster volume is

$$V_0 \approx h\pi r_0^2 \approx (70\ \mu\text{m})(\pi)(170\ \mu\text{m})^2 \tag{223}$$

$$\approx 6 \times 10^6\ \mu\text{m}^3. \tag{224}$$

Then, since most asters dissipate most quickly at such early times with initial powers of order $P_0 \sim$ few $\times\, 10^8$ ATP/s $\approx$ few $\times\, 10^{-11}$ W, asters are manifesting a volumetric power density $\rho_0$ of roughly,

$$\rho_0 = \frac{P_0}{V_0} \approx \frac{\text{few} \times 10^8 \ \text{ATP/s}}{6 \times 10^6 \ \mu\text{m}^3} \tag{225}$$

$$\approx 10^2 \ \text{ATP/s}/\mu\text{m}^3 = (\text{1-few}) \times 10^{-17} \ \text{W}/\mu\text{m}^3. \tag{226}$$

So, for instance, asters as a whole dissipate at total rates analogous to a few tens of exponentially growing *E. coli* cells, but are large enough in volume so that their power densities are about a tenth of those of starving microbial cells on a per volume basis.

**B. Comparison to calorimetric measurements on extensile active matter systems.** Foster and coworkers carefully measured the dissipation in an extensile active matter *in vitro* motor-microtubule system (48), in interesting contrast to our contractile active matter system here. Their paper reports a total dissipation rate $P_{\text{diss}} = 94 \pm 20$ nW in a $V = 0.5$ $\mu$L droplet. So that system exhibits a power density,

$$\rho = \frac{P}{V} \approx \frac{10^{-7} \ \text{W}}{5 \times 10^8 \ \mu m^3} \tag{227}$$

$$\approx \text{few} \times 10^{-16} \ W/\mu m^3 \approx \text{few} \times 10^3 \ \text{ATP/s}/\mu\text{m}^3. \tag{228}$$

Thus, this extensile active matter system studied by Foster and coworkers shows a power density about a few times ten fold larger than the contractile asters characterized in our present work, on a per-volume basis.

**C. Comparison to bacterial metabolisms.** Bacteria can show very plastic metabolic rates. To capture some characteristic scales, we refer to three empirical traditions.

First, as one extreme, exponentially growing model organisms such as *E. coli* consume of order $P \approx (\text{few} \times 10^6) - 10^7$ ATP/cell/s, as inferred both by explicit chemostat turnover measurements and modeling (49–51), in their approximately cubic micron volumes. (Much of this expenditure can be adequately explained by the cost of just duplicating the proteome turnover (52).)

In the metabolic slow lane, microbes can consume energy via ATP at radically lower rates. One tradition of measurement involves extrapolating the rate at which sugar consumption slows down with slower growth rate down to zero growth rate in chemostats; such measurements report highly typical power densities of $\langle \rho \rangle \approx 9 \times 10^5$ ATP/s/$\mu$m$^3$, (52–54), a maintenance energy expenditure scale in common with a number of eukaryotic cells as well.

More direct measurements on basal microbial metabolism add resolution to these smaller expenditures. For instance, Schink *et al.* use rates of dying to infer that starving *E. coli* cells consume energy at a rate $\beta \approx 0.6$ fmol glucose/cell/day (55). Since the complete oxidation of 1 molecule of glucose generates the energetic equivalent of 26 ATP molecules (BNID 114969; (56), this gives,

$$\beta \approx 0.6 \ \frac{\text{fmol glu}}{\text{CFU day}} \times \frac{10^{-15} \ \text{mol}}{\text{fmol}} \frac{26 \ \text{mol ATP}}{1 \ \text{mol glu}} \frac{6 \times 10^{23} \ \text{ATP}}{\text{mol}} \frac{1 \ \text{day}}{86400 \ s} \tag{229}$$

$$\approx 10^5 \ \text{ATP/s/cell}. \tag{230}$$

In other environmental settings, starving bacteria can operate at even lower metabolic rates; for instance, new direct measurements from the laboratory of Dianne Newman and colleagues (57) indicate that anoxic *P. aueriginosa* cells can operate metabolically at rates of order $10^3$ ATP/s/cell (in their order cubic micron volumes).

**D. Comparison to mouse oocytes.** Yang, Needleman, and coworkers (58) infer NADH flux in mouse oocytes using fluorescence lifetime imaging and a metabolic model. There, an oocyte shows an inferred/predicted NADH flux of $J_{ox}$ ranging from 55 $\mu$M/s to 80 $\mu$M/s (see their Figure 9). Recalling $1\mu\text{M} \approx 10^3$ molecules/$\mu$m$^3$, this suggests taking a value about $J_{ox} \approx (\text{few} \times 10^4 \text{NADH}/\mu\text{m}^3/s)$. Assuming 2.5 mol ATP/1 mol NADH turned over (see e.g. reference (59)), this implies a total ATP consumption rate per volume of order

$$\rho = J_{ox} \times \frac{\text{few ATP}}{\text{NADH}} \approx 10^5 \ \text{ATP}/\mu\text{m}^3/s. \tag{231}$$

So, these mouse oocytes dissipate about a thousand times faster per volume than the contracting asters explored by our measurements.

Towards a sense of the absolute oocyte dissipation budget, this flux occurs in an oocyte of apparent radius $r \approx 40\mu m$ (see their scale bar in their Figure 9), which we might assume has a spherical volume $V = \pi r^3 = 2 \times 10^5 \mu m^3$. (The calculation above does not require this value as input as the consumption rate is already registered in concentration depleted per time units.) However the total metabolic power of the whole oocyte would be implicated to be,

$$P = \rho V \approx (10^5 \text{ATP}/\mu\text{m}^3/s) \times (2 \times 10^5 \ \mu\text{m}^3) = 2 \times 10^{10} \ \text{ATP/s/oocyte}. \tag{232}$$

**E. Comparison to zebrafish embryos.** Rodenfels and colleagues (60) report beautiful calorimetric measurements of zebrafish embryos. Intriguingly, these measurements resolve both a total average net heat flow (putatively from total average metabolism), and also finer metabolic oscillations in net heat flow atop this average trend, presumably corresponding to the expenditures specific to the cell cycle.

The authors report that the *average* heat outflow per embryo changes from $P_{\text{avg}} = 60$ nJ/s at a two-cell-stage (which they map to $\rho_{\text{avg}} \approx 25$ $\mu$M ATP/s $\approx 1.5 \times 10^4$ ATP/$\mu$m$^3$/s, or 1 W/kg, and which we can equivalently express as $10^3$ W/$\mu$m$^3$) to $P_{\text{avg}}$ =84 nJ/s (translatable to $\rho_{\text{avg}}$ ≈1.4 W/kg = ($1.4 \times 10^3$ W/$\mu$m$^3$) or 35 $\mu$M ATP/s $\approx 2.1 \times 10^4$ ATP/$\mu$m$^3$/s) when at cleavage stage. So, the total average dissipation in such embryos is a few hundred times that of our contracting aster measurements on a per-volume basis.

The cell cycle dissipation oscillations registered by these embryos are about a hundred times smaller than the average expenditure, specifically $\rho_{\text{osc.}}$ falls between the scales of $\approx 0.25\ \mu M\ ATP/s$ to $\approx 0.75\ \mu M\ ATP/s$, namely a few tenths of micromolar ATP per second. This corresponds to a few hundred ATP per second per cubic micron, or within a factor few the power densities measured in our contracting asters.

The authors acknowledge an embryo volume of $V = 6 \times 10^7$ $\mu$m$^3$. This means,

$$P_{\text{avg}} \approx 10^{12}\ \text{ATP/embryo}, \qquad [233]$$

$$P_{\text{osc}} \approx 10^{10}\ \text{ATP/embryo}. \qquad [234]$$

#### 12. Jensen's inequality shows that hydrolysis rates can be underestimated by spatially-averaged values

What new theoretical insights can be unlocked by the hard-won measurement of spatiotemporal dissipation rates? Here, we highlight a simple, generic consequence of spatially-nonuniform concentration fields on the net dissipation sustaining nonequilibrium in a system. Due just to their nonlinear dependence on underlying chemical fields, true dissipation rates averaged or accumulated over space can substantially depart from their values predicted from spatially-averaged information—for instance, as predicted by more experimentally-straightforward, bulk biochemical measurements.

To gain quick intuition for how spatially-averaged quantities such as concentrations can systematically distort estimates of dissipation, we begin with a first toy kinetic model that illustrates the essential point (before graduating to nonlinearities apt for motors and ATP). Consider a system where an energy consumption rate $P$ depends on a reactant concentration $c$ according to a simple power law, namely $P = kc^\alpha$, for some effective kinetic exponent $\alpha$ and some constant of proportionality $\gamma_0$. (This relation may seem contrived, but in fact e.g. reasonably describes an energy consumption reaction under mass action kinetics, or indeed locally approximates more feedbacked reaction dynamics.) Then the true energy dissipation rate aggregated over the volume is proportional to $\langle P \rangle = \gamma_0 \langle c^\alpha \rangle$. However, if separate measurements furnish only the knowledge of the spatially-averaged concentration $\langle c \rangle$, then these yield instead an estimate $P_{\text{unif}} = \gamma_0 \langle c \rangle^\alpha$. The generic discrepancy $\langle c^\alpha \rangle \neq \langle c \rangle^\alpha$ when $\alpha \neq 1$ starkly exposes that averaged information will give systematic distortions to true aggregate behavior. In particular, whether the uniform mean-field $P_{\text{unif}}$ systematically exceeds or underestimates the true $\langle P \rangle$ depends on the convexity of the nonlinearity, as encoded by the particular numerical value of the exponent $\alpha$. The sign of the discrepancy is a mathematical object that can be commented usefully by Jensen's inequality, which we now discuss and employ for realistic motor-ATP kinetics.

In particular, using Jensen's inequality, along with negative correlations between energy-consuming agents and their fuel, we establish that whenever the dissipation rate is a concave function of some spatially-varying quantities—a functional dependence commonly observed or assumed in Michaelis-Menten kinetics of molecular motors, *inter alia*—the dissipation rate estimated by mean-field quantities can be an overestimate of the true dissipation rate.

Specifically, consider the dissipation rate $P \equiv \frac{d\text{ATP}}{dt}$ at which ATP locally changes in time. For concreteness, imagine motors at local concentration $m(\mathbf{r}, t)$ consume ATP at a maximum rate $\gamma_0$ modulated by a Michaelis-Menten dependence on ATP, subject to ATP's diffusion, namely,

$$\frac{\partial[\text{ATP}]}{\partial t} = -\gamma_0\ m(\mathbf{r}, t)\ \frac{[\text{ATP}](\mathbf{r}, t)}{[\text{ATP}](\mathbf{r}, t) + K} + D\nabla^2[\text{ATP}](\mathbf{r}, t), \qquad [235]$$

where ATP diffuses at diffusion coefficient $D$ and motors operate with a Michaelis constant $K$.

(For clarity of illustration, this is a significantly simplified reaction-diffusion system from the dynamics explored elsewhere in this text. More complex functional dependencies, including invoking the inhibitory effects of accumulating ADP and phosphate, can also be adopted without changing our conclusions, as long as these functions retain a concave dependence on the local value of ATP, and provided (as physically expected) additional covariance terms do not overwhelm an overall Jensen-gap contribution.) Integrated over some whole system volume $V$, the true total dissipation rate $P$—for instance, as measured by calorimetry and experienced as an energy need in a cell—is,

$$P^{\text{tot}} \equiv \int dV\ \frac{d[\text{ATP}]}{dt} = \int dV\ \left[ -\gamma_0\ m(\mathbf{r}, t)\ \frac{[\text{ATP}](\mathbf{r}, t)}{[\text{ATP}](\mathbf{r}, t) + K} + D\nabla^2[\text{ATP}](\mathbf{r}, t) \right], \qquad [236]$$

$$V \left\langle \frac{d[\text{ATP}](\mathbf{r}, t)}{dt} \right\rangle = -V\ \gamma_0\ \left\langle m(\mathbf{r}, t)\ \frac{[\text{ATP}](\mathbf{r}, t)}{[\text{ATP}](\mathbf{r}, t) + K} \right\rangle + \int_{\partial V} d(\partial V)\ \ (D\nabla\, [\text{ATP}]) \cdot \mathbf{n}, \qquad [237]$$

where we have used angular brackets to denote averages of quantities over spatial variation in the system, $\langle x\rangle \equiv \frac{1}{V}\int x\, dV$. We also applied the divergence theorem on the diffusion term on the right, to attribute the net diffusive change in ATP in the whole volume as the flux $\propto D\nabla\,[\mathrm{ATP}]$ integrated over the surface area of the system volume. For no-flux boundary conditions the latter term vanishes, and we see

$$\frac{P^{\mathrm{tot}}}{V} = \left\langle \frac{d[\mathrm{ATP}](\mathbf{r},t)}{dt} \right\rangle = -\gamma_0 \left\langle m(\mathbf{r},t)\, \frac{[\mathrm{ATP}](\mathbf{r},t)}{[\mathrm{ATP}](\mathbf{r},t)+K} \right\rangle. \qquad [238]$$

Often, more accessible experiments report only coarser, spatially-averaged information such as the system's total amount of ATP or motors, $\mathrm{ATP}^{\mathrm{tot}} = V\langle[\mathrm{ATP}]\rangle$ and $m^{\mathrm{tot}} = V\langle m\rangle$. Our motivating question is, **how well or badly does spatially-averaged information report on the true total dissipation rate?** In other words—given confidence in the form of the dissipation's dependence on underlying fields—how different are $\langle P([\mathrm{ATP}],m)\rangle \equiv \left\langle \frac{d[\mathrm{ATP}](\mathbf{r},t)}{dt}\right\rangle$ and $P\left(\langle[\mathrm{ATP}]\rangle,\langle m\rangle\right) \equiv \left.\frac{d[\mathrm{ATP}](\mathbf{r},t)}{dt}\right|_{\langle[\mathrm{ATP}]\rangle,\langle m\rangle}$?

To approach this question, we treat the underlying chemical fields $[\mathrm{ATP}](\mathbf{r},t)$ and $m(\mathbf{r},t)$ as random variables drawn from joint probability distributions over space in the system volume. Denoting a covariance between two random variables $X$ and $Y$ as $\sigma_{X,Y} \equiv \langle XY\rangle - \langle X\rangle\langle Y\rangle$, Eq. 238 can be expressed as,

$$\left\langle \frac{d[\mathrm{ATP}](\mathbf{r},t)}{dt} \right\rangle = -\gamma_0 \left( \langle m(\mathbf{r},t)\rangle \left\langle \frac{[\mathrm{ATP}](\mathbf{r},t)}{[\mathrm{ATP}](\mathbf{r},t)+K} \right\rangle + \sigma_{m,ATP/(ATP+K)} \right). \qquad [239]$$

Importantly, note that when variations in the underlying chemical fields have been self-organized by the action of motor consumption, we should generically expect that on average where there are many motors, there is a deficit of ATP, and vice versa. Such anticorrelations can develop in many closed self-organizing systems after an intermediate time when agents have exercised their activity to consume fuel. Thus, we expect that the motor distribution and the ATP distribution will be negatively correlated, and since $ATP/(K+ATP)$ is monotonic in ATP, we anticipate that $\boxed{\sigma_{m,ATP/(ATP+K)} \leq 0}$. *

Next, we turn to the second part of the first term, $\left\langle \frac{[\mathrm{ATP}](\mathbf{r},t)}{[\mathrm{ATP}](\mathbf{r},t)+K} \right\rangle$, namely the average of the dissipation rate's dependence on ATP. This is a concave function in the ATP concentration. This invites us to apply *Jensen's inequality*, which declares that if a function $f(X)$ is concave in $X$, then when $X$ experiences some variation, the average of the function is smaller than the function evaluated at the average, $f(\langle X\rangle) \geq \langle f(x)\rangle$. In the case of the Michaelis-Menten functional dependence, this means that

$$\left\langle \frac{[\mathrm{ATP}]}{[\mathrm{ATP}]+K} \right\rangle \leq \frac{\langle[\mathrm{ATP}]\rangle}{\langle[\mathrm{ATP}]\rangle+K}, \qquad [240]$$

which says that the so-called *Jensen gap*, the difference between the function of the average and the average of the function, is positive, $\boxed{\frac{\langle[\mathrm{ATP}]\rangle}{\langle[\mathrm{ATP}]\rangle+K} - \left\langle \frac{[\mathrm{ATP}]}{[\mathrm{ATP}]+K} \right\rangle \geq 0}$. Together, the two simple boxed facts—that the underlying fields are negatively correlated, and the concavity of the dissipation rate's dependence on the underlying field—mean that

$$\left(\langle P([\mathrm{ATP}],m)\rangle - P\left(\langle[\mathrm{ATP}]\rangle,\langle m\rangle\right)\right) = \left\langle \frac{d[\mathrm{ATP}](\mathbf{r},t)}{dt} \right\rangle - \left.\frac{d[\mathrm{ATP}](\mathbf{r},t)}{dt}\right|_{\langle[\mathrm{ATP}]\rangle,\langle m\rangle} \qquad [241]$$

$$= -\gamma_0 \left( \langle m(\mathbf{r},t)\rangle \left\langle \frac{[\mathrm{ATP}](\mathbf{r},t)}{[\mathrm{ATP}](\mathbf{r},t)+K} \right\rangle + \sigma_{m,ATP/(ATP+K)} \right) - \left( -\gamma_0\, \langle m(\mathbf{r},t)\rangle\, \frac{\langle[\mathrm{ATP}]\rangle}{\langle[\mathrm{ATP}]\rangle+K} \right) \qquad [242]$$

$$= \gamma_0 \left( \langle m(\mathbf{r},t)\rangle \underbrace{\left( \frac{\langle[\mathrm{ATP}]\rangle}{\langle[\mathrm{ATP}]\rangle+K} - \left\langle \frac{[\mathrm{ATP}]}{[\mathrm{ATP}]+K} \right\rangle \right)}_{\geq 0,\ \text{Jensen gap}} \underbrace{- \underbrace{\sigma_{m,ATP/(ATP+K)}}_{\leq 0,\ \text{correlation}}}_{\geq 0} \right). \qquad [243]$$

Note that the fact that this difference of two negative quantities—ATP dissipation rates in time, which are negative—is *positive* means that the magnitude of the first is smaller than the second. In other words, thanks to spatial variation, the magnitude of the true dissipation rate $\left|\left\langle \frac{d[\mathrm{ATP}](\mathbf{r},t)}{dt}\right\rangle\right|$ will be *smaller* than that predicted by taking spatially-averaged values for underlying fields, $\left| \left.\frac{d[\mathrm{ATP}](\mathbf{r},t)}{dt}\right|_{\langle[\mathrm{ATP}]\rangle,\langle m\rangle} \right|$.

For moderately-small variation in the underlying field variables, the size of the Jensen gap can be estimated by a Taylor expansion about the mean values; namely,

$$\langle f(x)\rangle \approx f(\langle x\rangle) + \frac{1}{2} f''(\langle x\rangle)\sigma_x^2. \qquad [244]$$

*Put more precisely, a sufficient condition for the required $\sigma_{m,ATP/(ATP+K)} \leq 0$ negative correlation to develop between the Michaelis Menten kinetic factor $ATP/(ATP+K)$ and motors $m$ is that $\langle m|A > a\rangle \leq \langle m\rangle$ for all thresholds $a$, e.g. the average value of a motor concentration given an that ATP concentration is at least some value $a$ should be at most the unconditional average motor concentration.

So the larger the variability $\sigma_x^2$ in the underlying variable $x$, the larger the discrepancy between the mean-field and true dissipation rates.

For the Michaelis-Menten hydrolysis function $f(x) = \frac{x}{x+K}$ in our model of dissipation in Eq. 235, we calculate that the second derivative evaluated at the average ATP concentration, $f''(\langle x \rangle)$, is

$$f''(\langle x \rangle) = \frac{\partial}{\partial x}\left(\frac{\partial f}{\partial x}\right)\bigg|_{\langle x \rangle} \qquad [245]$$

$$= \frac{\partial}{\partial x}\left(\frac{K}{(x+K)^2}\right)\bigg|_{\langle x \rangle} \qquad [246]$$

$$= -2\frac{K}{(\langle x \rangle + K)^3}. \qquad [247]$$

So a first order estimate of this hydrolysis function's Jensen gap is

$$\text{Jensen gap} \equiv |\langle f(x) \rangle - f(\langle x \rangle)| \approx \frac{1}{2}\left|f''(\langle x \rangle)\right| \sigma_x^2 \qquad [248]$$

$$\approx \frac{\sigma_{ATP}^2}{2}\left|-2\frac{K}{(\langle[\mathrm{ATP}]\rangle + K)^3}\right| \qquad [249]$$

$$\approx \sigma_{ATP}^2 \frac{K}{(\langle[\mathrm{ATP}]\rangle + K)^3}. \qquad [250]$$

#### 13. Spatial gradients in ATP are upper bounded by those in underlying reaction rates (accumulated over time)

Our finite element simulations discussed earlier show that when the hydrolysis rate of ATP $h(r, t, \mathrm{ATP}(r,t))$ is controlled only by a motor concentration $m(r,t)$,
namely,

$$h(r, t, \mathrm{ATP}) \equiv \gamma\; m(r,t)\; p_{\text{mot-ATP}}(\mathrm{ATP}(r,t)), \qquad [251]$$

where,

$$p_{\text{mot-ATP}}(\mathrm{ATP}(r,t)) \equiv \frac{\frac{\mathrm{ATP}}{K_T}}{1 + \frac{\mathrm{ATP}}{K_T} + \frac{\mathrm{ADP}}{K_D} + \frac{\mathrm{P}_i}{K_D}} \qquad [252]$$

expresses the binding probability of ATP to a motor), gradients do *not* appreciably develop in finite element modeling with physically-realistic parameters. However, when the hydrolysis rate also acknowledges dependence on the microtubule concentration (because only a motor bound both to ATP and to microtubules should be expected to step and appreciably hydrolyze ATP), meaningful gradients can and *do* develop. (That is, when

$$h(r, \mathrm{ATP}) \equiv \gamma\; m(r,t)\; p_{\text{mot-ATP}}(\mathrm{ATP}(r,t))\; p_{\text{mot-tub}}\left(\mathrm{tub}(r,t), m(r,t)\right), \qquad [253]$$

where $p_{\text{mot-tub}}$ is the probability that a motor is bound to a microtubule, gradients in ATP easily develop in numerical simulations.) The reason gradients become newly feasible is that including the itself spatially-varying $p_{\text{mot-tub}}\left(\mathrm{tub}(r,t), m(r,t)\right)$ further sharpens the spatial localization of the hydrolysis rates.

In fact, this link between concentration gradients and reaction rate gradients expresses a more general principle. Here we set up and justify the claim in the main text (found in the finite element modeling Results section) that a spatial gradient in a concentration field (such as ATP) at any time is upper bounded by maximal gradients in underlying consumption rates, accumulated over time. Specifically, we will establish that if $h\left(r, \mathrm{ATP}(r,t)\right)$ is the local consumption rate at radial position $r$ and time $t$, then

$$\boxed{\max_{r\in[0,R]} \left|\frac{\partial \mathrm{ATP}}{\partial r}(r,t)\right| \leq \int_0^t dt' \max_{r'\in[0,R]} \left|\frac{\partial h}{\partial r}\bigg|_{\mathrm{ATP}}(r',t')\right|}, \qquad [254]$$

This inequality, while very simple, is illuminating. It makes both general and precise the understanding that extant gradients in observed continuum fields emerge only thanks to sufficiently large spatial variation in the underlying primordial reaction rates. Accordingly, observing a gradient in a concentration of a given magnitude implicates that gradient in governing reaction rates accumulate to values at least this large.

Our approach will be as follows. To learn about $\max_{r\in[0,R]} \frac{\partial \mathrm{ATP}}{\partial r}\Big|_t$, the maximum concentration gradient over all points in space at a given time $t$, we write an expression for the time evolution of the gradient at any position $r$ at any instant in time $t' < t$, then find the maximum size of this rate of change in the gradient over all positions at this instant (by connecting this to the rate of change of currently steepest gradient). Then, focusing on this central location of the steepest gradient, we discard diffusive terms (which can only smoothen gradients there), yielding an upper bound for this maximal rate of change over all positions and time-integrate it, leading to the bound on the right hand side of Eq. 254.

To proceed, specifically, let a concentration field $\mathrm{ATP}(r,t)$ change due to a generic consumption reaction $h \geq 0$ smoothed by diffusion, namely,

$$\frac{\partial \mathrm{ATP}}{\partial t} = D\nabla^2 \mathrm{ATP}(r,t) - h\left(r, \mathrm{ATP}(r,t)\right) \tag{255}$$

$$= D\left(\frac{\partial^2 \mathrm{ATP}}{\partial r^2} + \frac{2}{r}\frac{\partial \mathrm{ATP}}{\partial r}\right) - h\left(r, \mathrm{ATP}(r,t)\right), \tag{256}$$

where we adopt the case of spherical symmetry in the second line. Differentiating this equation with respect to position $r$ then gives how the spatial gradient in ATP (which for convenience we may call $g(r,t) \equiv \frac{\partial \mathrm{ATP}}{\partial r}$) itself changes in time, namely,

$$\frac{\partial}{\partial r}\frac{\partial \mathrm{ATP}}{\partial t} = \frac{\partial}{\partial t}\frac{\partial \mathrm{ATP}}{\partial r} = \frac{\partial g}{\partial t} \tag{257}$$

$$\rightarrow \frac{\partial g}{\partial t} = \frac{\partial}{\partial r}\left[D\left(\frac{\partial^2 \mathrm{ATP}}{\partial r^2} + \frac{2}{r}\frac{\partial \mathrm{ATP}}{\partial r}\right) - h\left(r, \mathrm{ATP}(r,t)\right)\right] \tag{258}$$

$$= D\left(\frac{\partial^3 \mathrm{ATP}}{\partial r^3} - \frac{2}{r^2}\frac{\partial \mathrm{ATP}}{\partial r} + \frac{2}{r}\frac{\partial}{\partial r}\frac{\partial \mathrm{ATP}}{\partial r}\right) - \left(\left.\frac{\partial h}{\partial r}\right|_{\mathrm{ATP}} + \frac{\partial h}{\partial \mathrm{ATP}}\frac{\partial \mathrm{ATP}}{\partial r}\right) \tag{259}$$

$$= D\left(\frac{\partial^2 g}{\partial r^2} - \frac{2}{r^2}\, g + \frac{2}{r}\frac{\partial g}{\partial r}\right) - \left(\left.\frac{\partial h}{\partial r}\right|_{\mathrm{ATP}} + \frac{\partial h}{\partial \mathrm{ATP}}\, g\right). \tag{260}$$

Having written an equation for the time evolution of the gradient as Eq. 260, we now pursue information about its maximum value over all positions in some finite domain $r \in [0, R]$ of physical interest. Specifically, consider the case of physical interest where the ATP begins at some ambient value uniform in space, $\mathrm{ATP}(r, t=0) = \mathrm{ATP}_0$, and the volume is closed giving no flux boundary conditions, $\frac{\partial \mathrm{ATP}}{\partial r}(r=R,t) = 0$, along with the regularity condition $\lim_{r\to 0}\frac{\partial \mathrm{ATP}}{\partial r}(r,t) = 0$. These boundary conditions in the concentration field imply the corresponding boundary conditions in the spatial gradient field of $g(r, t=0) = 0$ and $g(r=R,t) = g(r\to 0,t) = 0$. This means that,

$$g(r,t) = g(r, t=0) + \int_0^t dt'\ \frac{\partial g}{\partial t} \tag{261}$$

$$= \int_0^t dt'\ \frac{\partial g}{\partial t}. \tag{262}$$

We are interested in the steepest gradient over all positions $r \in [0, R]$, and to learn about this at a final time $t$ it is useful to understand how it changes in time at earlier times $t' < t$. First, we bound the positive maximum of the gradient $g$. (Any negative parts are confronted by applying the same argument to the negated gradient $-g$.) Let $r_*(t')$ be the location of the steepest gradient (namely, satisfying $g(r_*(t'), t') = \max_r g(r,t')$).Assuming for simplicity (as should apply to scenarios of most physical interest) that the positive maximum is unique in space, varies smoothly in time, and found in the interior of the spatial volume (as implicated by the vanishing boundary conditions), then the steepest gradient's rate of change can be expressed with the chain rule as,

$$\frac{\partial}{\partial t'}\max_r g(r,t') = \frac{\partial}{\partial t'} g\left(r_*(t'), t'\right) \tag{263}$$

$$= \frac{\partial g\left(r_*(t'), t'\right)}{\partial r_*}\frac{\partial r_*(t')}{\partial t'} + \left.\frac{\partial g}{\partial t}\right|_{(r_*(t'),t')} \tag{264}$$

$$= \left.\frac{\partial g}{\partial t}\right|_{(r_*(t'),t')}. \tag{265}$$

The last line follows since $r_*(t')$ maximizes the gradient (and the vanishing boundary conditions ensure this steepest location $r_*(t') \in (0, R)$ occurs in the interior (not the boundaries) of our spatial domain of interest), we know $\frac{\partial g\left(r_*(t'),t'\right)}{\partial r_*} = 0$ by virtue of the maximum. (In words, this expression says that the rate at which the maximal gradient changes with time is the same as the gradient's rate of change evaluated at the current maximum; this is extremely natural, apart from the possibility that the location $r_*(t')$ of the gradient can itself change, but the flatness at the maximum suppresses this effect from changing the rate of change of the maximum. Interestingly, this reasoning occurs in other guises as the "envelope theorem" in economics and optimization.) Integrating in time, these statements give,

$$\max_r g(r,t) = \int_0^t dt'\ \frac{\partial}{\partial t'}\max_r g(r,t') \tag{266}$$

$$= \int_0^t dt'\ \left.\frac{\partial g}{\partial t}\right|_{(r_*(t'),t')}, \tag{267}$$

and now we may substitute the expression for the time evolution $\frac{\partial g}{\partial t'}$ expressed in Eq. 260, with the important detail that we will evaluate all the terms in Eq. 260 at the point $r_*(t')$ of the steepest gradient:

$$\max_r g(r,t) = \int_0^t dt' \left[ D \left( \frac{\partial^2 g}{\partial r^2} - \frac{2}{r^2}\, g + \frac{2}{r}\frac{\partial g}{\partial r} \right)\Bigg|_{r_*(t')} - \left( \frac{\partial h}{\partial r}\bigg|_{\mathrm{ATP}} + \frac{\partial h}{\partial \mathrm{ATP}}\, g \right)\Bigg|_{r_*(t')} \right]. \quad [268]$$

Now, we claim that the first diffusive-smoothing terms, $D\left(\frac{\partial^2 g}{\partial r^2} - \frac{2}{r^2}\, g + \frac{2}{r}\frac{\partial g}{\partial r}\right)\Big|_{r_*(t')}$, is not positive. We validate this term-by-term. The curvature term $\frac{\partial^2 g}{\partial r^2}\Big|_{r_*(t')}$ is less than or equal to zero by the fact that $r_*(t')$ is an interior maximum (namely, via the second derivative test). The term $-\frac{2}{r^2}\, g$ is clearly negative (or zero) at the positive maximum under consideration ($g(r_*(t'),t') \geq 0$). Last, $\frac{2}{r}\frac{\partial g}{\partial r}\Big|_{r_*(t')}$ vanishes, because the gradient's spatial derivative vanishes when evaluated its maximizing position. We conclude that indeed, as a group, these diffusive terms are definitely nonpositive. (This is only true at the gradient-maximizing position $r_*(t')$, as at other spatial positions e.g. valleys, diffusion can fill in and boost other local gradients; but the important point is that we care about the behavior specifically at the point of the steepest gradient in order to harness Eq. 267.)

Accordingly, neglecting the diffusive smoothing terms is guaranteed to overestimate the rate of change of the maximum gradient: this gives an upper bound as,

$$\frac{\partial}{\partial t'} \max_r g(r,t') \leq -\left( \frac{\partial h}{\partial r}\bigg|_{\mathrm{ATP}} + \frac{\partial h}{\partial \mathrm{ATP}}\, g \right)\Bigg|_{r_*(t')}. \quad [269]$$

The time integral gives,

$$\max_r g(r,t) \leq -\int_0^t dt' \left( \frac{\partial h}{\partial r}\bigg|_{\mathrm{ATP}} + \frac{\partial h}{\partial \mathrm{ATP}}\, g \right)\Bigg|_{r_*(t')}. \quad [270]$$

The two terms in the integrand reflect, respectively, intrinsic spatial variation in hydrolysis rate $\frac{\partial h}{\partial r}\Big|_{\mathrm{ATP}}$, and variation inherited from any underlying extant ATP gradient, $\frac{\partial h}{\partial \mathrm{ATP}}\, g$. While this expression registers that both sources of variation can accumulate to accentuate an ATP gradient, it is interesting to isolate the first intrinsic effect. Note that when the hydrolysis rate $h$ depends on the underlying ATP it consumes in a monotonically increasing way, as is very natural in nearly all kinetic circumstances such as Michaelis Menten or mass action dynamics, then its variation with ATP is nonnegative, $\frac{\partial h}{\partial \mathrm{ATP}} \geq 0$. This means neglecting that term also gives a further upper bound expressed purely in terms of the intrinsic spatial variation of the hydrolysis rate:

$$\max_r g(r,t) \leq -\int_0^t dt' \; \frac{\partial h}{\partial r}\bigg|_{\mathrm{ATP},r_*(t')} \leq -\int_0^t dt' \max_{r' \in [0,R]} \frac{\partial h}{\partial r}\bigg|_{r'}. \quad [271]$$

This gives our claimed original inequality in Eq. 254. Our argument can be repeated, *mutatis mutandis*, for negative gradients, to give an inequality where unsigned steepness magnitude is bounded:

$$\boxed{\max_r \left| g(r,t') \right| \leq \int_0^t dt' \max_{r' \in [0,R]} \left| \frac{\partial h}{\partial r}\bigg|_{r'} \right|}. \quad [272]$$

#### 14. Power Estimates

For decades, the question of how energy is invested in cellular processes has been of great interest. In the 1970s, several independent threads converged on these same questions with one set of efforts focused on the cost of cytoplasm (49, 54), others focused on the apparently futile cycles of GTP hydrolysis in protein translation (61) and yet others investigating the role of GTP hydrolysis in the context of microtubules (62, 63). Our aim in this section is to build on these early efforts (and many others) to carefully characterize the power cost of a variety of processes that attend the rearrangements of the motors and microtubules that occur during aster formation in our experiments. Broadly speaking, the philosophy of the approach is to make a list of various processes that we know take place during aster formation and to examine quantitative estimates of the power associated with these processes. Conceptually, the structure of all of the estimates will be the same, with the generic functional form

$$\text{power of process} = J_{process} \times \Delta\mu_{process}. \quad [273]$$

Here we have defined the quantity $\Delta\mu_{process}$ as the free energy cost of a unit process such as a single step of a motor or the movement of a single molecule up a gradient. $J_{process}$ refers to the flux associated with the process of interest, meaning how many unit processes occur per unit time. For example, in the context of constantly pumping ions up a concentration gradient, $\Delta\mu_{process}$ refers to the free energy cost of taking a single ion from one side of the membrane to the other. Similarly, the flux in that case would be given by a phenomenological linear transport law relating the concentration jump across the membrane to the flux itself. Using this quantitative structure, we carry out a series of estimates for each of the processes we think is implicated in the structural rearrangements of our motor-microtubule systems.

**A. Experimentally Measured Power.** During aster formation, we measure $\approx f \times 10^8$ ATPs are consumed per second, as shown in the Main Text Figure 5. As noted above, we aim to put this measured value into context through a series of estimates geared towards understanding the power of various processes that attend the formation of an aster. In particular, we are intrigued by the relative costs of processes such as the entropy of orientational ordering, the reduction in volume (i.e. pV work), the maintenance of gradients and so on. Each of the sections below examines one such process in detail.

**B. Estimate of the Power of Stepping Motors.** At the mechanistic level, we know what is happening to the ATP. It is being consumed by motors. Hence, the simplest statement about the ATP consumption is that every time a motor takes a step it hydrolyzes an ATP. As a proof of principle check, we ask if the power expenditure based on the measured motor hydrolysis rate matches the experimentally measured power. We measure a motor hydrolysis rate of $\gamma \approx 0.5 \frac{\mathrm{ATP}}{\mathrm{s \cdot motor}}$. If all motor proteins consume ATP at this rate, we can multiply the hydrolysis rate by the number of motors in the system to find the expected power. Our experiments contain 1 µM motors, in an initial cylindrical volume of

$$V_i = \pi R^2 d = \pi \times (125\,\mu\mathrm{m})^2 \times (70\,\mu\mathrm{m}) \approx 3 \times 10^6\,\mu\mathrm{m}^3, \tag{274}$$

which translates to

$$N_{\mathrm{mot}} = C_i V_i = 10^{-6}\,\frac{\mathrm{mol}}{\mathrm{L}} \times \left(10^3\,\frac{\mathrm{L}}{\mathrm{m}^3}\right) \times \left(3 \times 10^{-12}\,\mathrm{m}^3\right) \times \left(6 \times 10^{23}\,\frac{\mathrm{MT}}{\mathrm{mol}}\right) \approx 2 \times 10^9\,\mathrm{motors}. \tag{275}$$

Thus the power expected of the system is

$$P = N_{\mathrm{mot}}\gamma = 0.5 \frac{\mathrm{ATP}}{\mathrm{s \cdot motor}} \times \left(2 \times 10^9\ \mathrm{motors}\right) \approx 10^9\,\frac{\mathrm{ATP}}{\mathrm{s}}. \tag{276}$$

Compared to the $f \times 10^8\,\frac{\mathrm{ATP}}{\mathrm{s}}$ measured in our experiment, this estimate is within a factor of a few of the measured ATP consumption rate. In this sense, the experimental measurements are coherent with what we know about the agents that are consuming ATP. But our question is really a deeper one. We ask what is it that the motors are doing with their capacity to do work through the application of forces?

**C. Estimate of the Power of Aster Contraction via pV Work.** During the process of aster formation, the volume occupied by the microtubules that make up that aster decreases. Initially, microtubules of fixed length are uniformly distributed in a cylinder with a height, $h$, and radius, $R$. This cylinder corresponds to the region where we project light to induce motor dimerization, which drives aster formation. Some time after turning on the light, $\Delta\mathcal{T}$, the microtubules are organized in a sphere with radius, $r$, where $R > r$. This process is illustrated in Figure S83. Here, we perform a simple estimate to determine the pressure-volume work to contract an ideal gas of microtubules into a smaller aster volume. We are cognizant that a collection of microtubules is not a gas, but we suspect this the entropic cost will exceed the value we estimate here. We express the pressure of the microtubule gas as a function of volume based on the ideal gas law as

$$PV = Nk_{\mathrm{B}}T \Rightarrow P = \frac{Nk_{\mathrm{B}}T}{V}, \tag{277}$$

and input it into the equation for pressure-volume work

$$W = -\int_{V_i}^{V_f} P dV = -\int_{V_i}^{V_f} \frac{Nk_{\mathrm{B}}T}{V} dV = -Nk_{\mathrm{B}}T \ln\left(\frac{V_f}{V_i}\right), \tag{278}$$

where $V_f$ is the aster sphere volume and $V_i$ is the cylindrical microtubule gas volume. Note that $V_f$ is smaller than $V_i$ which implies that $V_f/V_i < 1$, and hence the work done is positive since the system is compressed. We now use numbers characteristic of our experiments to calculate the amount of work performed in the process of contracting the network into its final shape. First, we take the initial cylindrical volume from equation 274 and the final spherical aster volume to be

$$V_f = \frac{4}{3}\pi r^3 = \frac{4}{3}\pi \times (30\,\mu\mathrm{m})^3 \approx 10^5\,\mu\mathrm{m}^3. \tag{279}$$

Given the initial concentration of microtubules $C_i = 1$ nM, we can find the number of microtubules as

$$N_{\mathrm{MT}} = C_i V_i = 10^{-9}\frac{\mathrm{mol}}{\mathrm{L}} \times \left(10^3\,\frac{\mathrm{L}}{\mathrm{m}^3}\right) \times \left(3 \times 10^{-12}\,\mathrm{m}^3\right) \times \left(6 \times 10^{23}\,\frac{\mathrm{MT}}{\mathrm{mol}}\right) \approx 2 \times 10^6\,\mathrm{MT}. \tag{280}$$

We can now combine all of these results to find the pressure-volume work as

$$W = -2 \times 10^6 \times \ln\left(\frac{1}{30}\right)\,k_{\mathrm{B}}T \approx 7 \times 10^6\,k_{\mathrm{B}}T. \tag{281}$$

Under physiological conditions the energy derived from an ATP hydrolysis event is $\approx 20\,k_{\mathrm{B}}T$ (51). Thus, in ATP units the work is equivalent to

$$W \approx 7 \times 10^6\,k_{\mathrm{B}}T \times \frac{\mathrm{ATP}}{20\,k_{\mathrm{B}}T} = 4 \times 10^5\ \mathrm{ATP}. \tag{282}$$

We can also compute the power of volume contraction by dividing the work by the time it takes to form the aster. In our experiments, asters form in approximately 10 minutes, thus

$$P = \frac{W}{\Delta \mathcal{T}} = \frac{4 \times 10^5 \text{ATP}}{600 \text{ s}} \approx 10^3 \frac{\text{ATP}}{\text{s}}. \qquad [283]$$

From this estimate, we find that the pressure-volume contribution to the power of aster contraction is negligible since the pV work effect is five orders of magnitude smaller than measured powers. We will approach this same question differently using ideas of nematohydrodynamics in a later section.

**D. The Entropy Cost of Bundling.** Here we investigate the power to reduce the entropy of the system by dimerizing motor proteins. For this simple thought experiment suggested to us in conversations with Erwin Frey, we compare the number of microstates of a pair of independent microtubules, not bridged by motors, in comparison to linked microtubules by a dimerized motor pair. Say that two independent microtubules of length $l$ are placed on separate one dimensional tracks with lattice sites at a spacing $a$, the step size of the motor. If each track is twice the length of a microtubule, $2l$, then there are $l/a$ lattice sites that the microtubule center can occupy, see Figure S84. For two independently moving microtubules, this amounts to $(l/a)^2$ microstates. Thus the entropy of the independent microtubule case is

$$S_{\text{indep}} = 2k_{\text{B}} \ln \left( \frac{l}{a} \right). \qquad [284]$$

If we now imagine a motor protein aligning the two microtubules, we now consider both microtubules to be on the same track. The microtubules now must move together. For the sake of this estimate, lets say that the microtubules are perfectly aligned with their ends at the same positions. The number of microstates of the coupled system is now only $l/a$, giving an entropy of

$$S_{\text{coup}} = k_{\text{B}} \ln \left( \frac{l}{a} \right). \qquad [285]$$

We can compute the free energy of the microtubule coupling as

$$\begin{aligned} \Delta G = -T\Delta S = -T(S_{\text{coup}} - S_{\text{indep}}) \\ \Delta G = k_{\text{B}} T \ln \left( \frac{l}{a} \right) \end{aligned} \qquad [286]$$

The length of stabilized microtubules in our experiment are $1\,\mu$m and the size of a motor step is 8 nm. Plugging in numbers, we find the free energy to couple two microtubules is

$$\Delta G = k_{\text{B}} T \ln \left( \frac{1000 \text{ nm}}{8 \text{ nm}} \right) \approx 5\, k_{\text{B}} T \approx 0.25 \text{ ATP}, \qquad [287]$$

where ATP $= 20\, k_{\text{B}} T$ (51). Say we wanted all of the microtubules in our aster system to fo rm one aligned bundle. Calculating the energy to couple all $2 \times 10^6$ microtubules together, the total energy is

$$\Delta G_{\text{tot}} = \Delta G \times N_{\text{MT}} = 5 \times 10^5 \text{ ATP}. \qquad [288]$$

If the bundle is formed on the same timescale as the aster, about 10 minutes, the power of bundling is

$$P = \frac{\Delta G}{t} = \frac{5 \times 10^5 \text{ ATP}}{600 \text{ s}} \approx 10^3 \frac{\text{ATP}}{\text{s}}. \qquad [289]$$

This estimate is five orders of magnitude smaller that the measured power, indicating that the entropic cost of microtubule coupling through motor dimerization is negligible.

**E. Estimate of Power Associated with Dragging a Microtubule.** One of the ways in which motors act is by producing motion. Like with most all real world motions, these processes dissipate energy. That is, some of the energy is transferred from the center of mass of the motor-cargo complex to the microscopic motions of the molecules making up the surrounding medium. This is revealed as dissipation. Here we ask what fraction of the ATP being consumed by each motor is converted into heat energy. In particular, we compute the power of a motor protein dragging a microtubule into an aster. This power can be expressed as

$$P = F_{\text{D}} v, \qquad [290]$$

where $F_{\text{D}}$ is the drag force and $v$ is the velocity of the motor.

To estimate the magnitude of this dissipation, we use the Stokes law to compute the drag force. Before making the calculation itself, we first examine the Reynolds number which gives us a measure of the dissipative forces and tells us whether we are in the regime of validity of the Stokes law. Since Stokes' law requires the system to be at low Reynolds number, we first make the following upper bound calculation for the Reynolds number in our experiment, namely,

$$Re = \frac{\rho v l}{\mu} = \frac{\left(10^3 \frac{\text{kg}}{\text{m}^3}\right) \times \left(10^{-7} \frac{\text{m}}{\text{s}}\right) \times \left(3 \times 10^{-5} \text{ m}\right)}{10^{-3} \frac{\text{N} \cdot \text{s}}{\text{m}^2}} \approx 10^{-6}, \qquad [291]$$

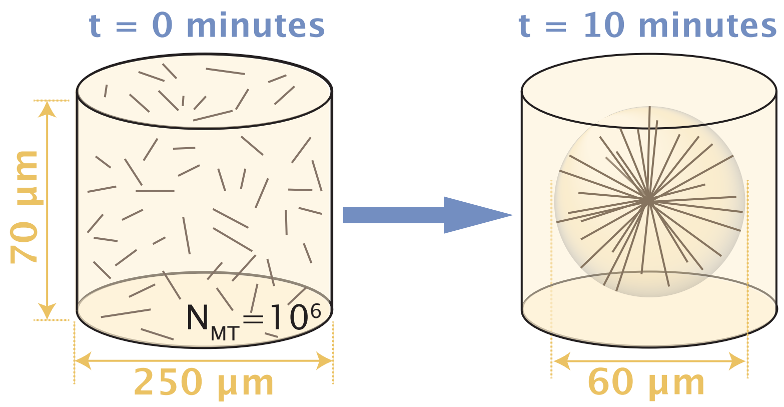

**Fig. S83. The work of aster compression.** At the beginning of the experiment, microtubules are uniformly mixed throughout a cylinder of projected light. After some time, $\mathcal{T}$, microtubules organize into a spherical aster with a volume smaller than the initial cylinder volume.

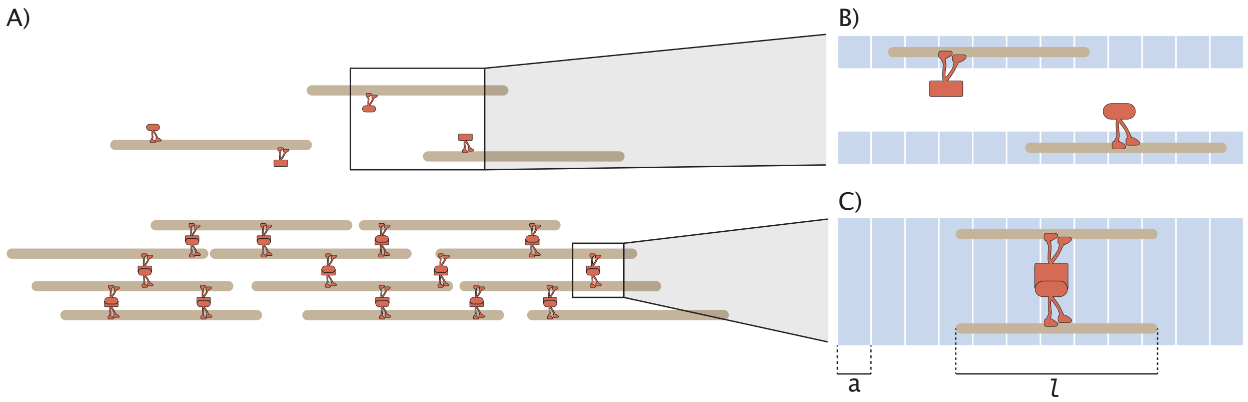

**Fig. S84. Microtubule Coupling.** A) Cross-linked microtubules form a bundle. B) Two microtubules on separate 1D tracks moving independently. C) Two microtubules aligned on the same track moving in unison.

where $\rho = 1000\,\mathrm{kg/m^3}$ is the fluid density, $v = 10^{-7}\,\mathrm{m/s}$ is the measured motor speed of our NCD motors, $l = 3 \times 10^{-5}$ m is the measured aster radius, and $\mu = 0.003\,\mathrm{N \cdot s/m^2}$ is the viscosity (11). We see that the Reynolds number satisfies $Re << 1$. Since our experiment meets the low Reynolds number criterion, we can use the Stokes law ($F_D = 6\pi\mu r v$) for the drag force in equation 290, resulting in a power of the form

$$P_{\text{per MT}} = 6\pi\mu r v^2, \qquad [292]$$

where $r$ is an effective radius as we will describe below.

We begin with the most extreme model of the drag force which is to imagine a microtubule as a sphere. This approach gives us a first order-of-magnitude look at the scale of the mechanical power associated with fluid drag. This calculation provides an upper bound as it assumes the moving object to be a sphere, which we will take to a have a radius of half the length of a microtubule, as depicted in Figure S85. Using our measurements for microtubule length, $1\,\mu$m, and motor speed, 100 nm/s, we find the power for a single motor to drag a microtubule is

$$P_{\text{per MT}} = 6\pi \times \left(3 \times 10^{-3}\,\frac{\mathrm{N \cdot s}}{\mathrm{m^2}}\right) \times \left(5 \times 10^{-7}\,\mathrm{m}\right) \times \left(10^{-7}\,\frac{\mathrm{m}}{\mathrm{s}}\right)^2 \approx 0.3\,\frac{\mathrm{pN \cdot nm}}{\mathrm{s}}. \qquad [293]$$

Taking $k_B T = 4\,\mathrm{pN \cdot nm}$ and the energy of ATP hydrolysis $\Delta\mu_{\text{ATP}} = 20\,k_B T$, we can convert to units of ATP/s, resulting in

$$P_{\text{per MT}} = 4 \times 10^{-3}\,\frac{\mathrm{ATP}}{\mathrm{MT \cdot s}}. \qquad [294]$$

The concentration of motor proteins to microtubules is 1000-fold, so if every single microtubule was being dragged into the aster at the same time, the maximum power from dragging microtubules would be the power per microtubule, as found in equation293, multiplied by the number of microtubules, as estimated in equation 280, resulting in

$$P = P_{\text{per MT}} \times N_{\text{MT}} = \left(4 \times 10^{-3}\,\frac{\mathrm{ATP}}{\mathrm{s \cdot MT}}\right) \times \left(2 \times 10^{6}\,\mathrm{MT}\right) \approx 10^4\,\frac{\mathrm{ATP}}{\mathrm{s}}. \qquad [295]$$

Again, we find that this estimate is far lower than our measured power by four orders of magnitude. Note that we can go farther and more precisely treat the fluid drag on a cylinder, taking care to distinguish motion parallel or perpendicular to the long axis of the microtubule. But these estimates lead to an even smaller power and we leave them as an exercise for the reader.

Though it will result in an even smaller estimate, we now refine our previous estimates. We refine our sphere estimate, to more accurately represent the surface area of a cylinder. Our previous estimate significantly overestimates the size of the microtubule by taking the sphere diameter to be the length of the microtubule. Now we will compute the spherical drag for a sphere with the same surface area as our microtubule. For a cylindrical microtubule of length $l = 1\,\mu$m as measured, and a radius $r_{\text{cyl}} = \frac{25\text{ nm}}{2} \approx 10$ nm (51), the surface area is

$$S_{\text{cyl}} = 2\pi r_{\text{cyl}} l + 2\pi r_{\text{cyl}}^2 \approx 2\pi \times 10^{-2}\,\mu\mathrm{m}^2. \qquad [296]$$

We now find the spherical radius for a sphere with the same surface area as a cylinder,

$$\begin{aligned} S_{\text{sph}} &= 4\pi r_{\text{sph}}^2 = 2\pi(r_{\text{cyl}} l + r_{\text{cyl}}^2) = S_{\text{cyl}} \\ \Rightarrow r_{\text{sph}} &= \left(\frac{1}{2}(r_{\text{cyl}} l + r_{\text{cyl}}^2)\right)^{-2} = \left(\frac{1}{4\pi} S_{\text{cyl}}\right)^{-2}. \end{aligned} \qquad [297]$$

Thus in our case,

$$r_{\text{sph}} = \left(5 \times 10^{-3}\right)^{-2}\,\mu\mathrm{m} \approx 7 \times 10^{-2}\,\mu\mathrm{m}. \qquad [298]$$

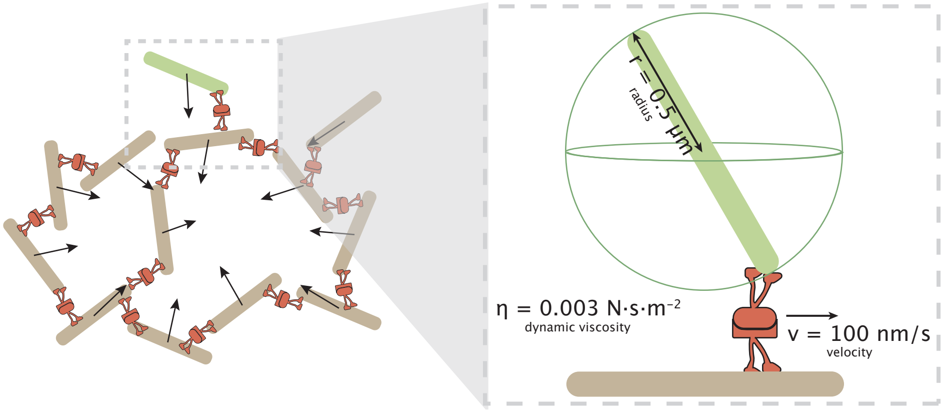

**Fig. S85. Drag on a Microtubule.** This figure illustrates the set up to calculate the drag on a motor protein carrying a microtubule. On the left, many microtubules are dragged into the center of mass to create an aster. The inset shows the force of drag due to pulling a single microtubule. As an upper bound, the microtubule is depicted as a sphere with a diameter of the microtubule length, allowing the application of Stokes' law.

Applying Stokes' Law for the new radius, we find

$$P_{\text{per MT}} = -6\pi \left(3 \times 10^{-3}\,\frac{\text{N}\cdot\text{s}}{\text{m}^2}\right)\left(7\times 10^{-8}\,\text{m}\right)\left(10^{-7}\,\frac{\text{m}}{\text{s}}\right)^2 \approx f\times 10^{-2}\,\text{pN}\cdot\text{nm} \approx f \times 10^{-4}\,\frac{\text{ATP}}{\text{s}\cdot\text{MT}}, \quad [299]$$

for each microtubule and

$$P = P_{\text{per MT}} \times N_{\text{MT}} = \left(f \times 10^{-4}\,\frac{\text{ATP}}{\text{s}\cdot\text{MT}}\right)\left(2\times 10^6\ \text{MT}\right) \approx 10^3\,\frac{\text{ATP}}{\text{s}}, \quad [300]$$

for the process of dragging all microtubules. This is one order of magnitude lower than our previous estimate.

***E.1. Cylindrical Drag.*** According to Finn (1953) (64), the cylindrical drag coefficient when the long axis is perpendicular to the fluid flow can be calculated as

$$C_{\text{cyl}} = \frac{8\pi}{\frac{1}{2} - \gamma - \ln(N_{Re}/8)}, \quad [301]$$

where $\gamma \approx 0.577$ is the Euler-Mascheroni constant and $N_{Re}$ is the Reynolds number at the length scale of the cylinder diameter. Finding the Reynolds number for a diameter of $l = 25$ nm,

$$N_{Re} = \frac{\rho v l}{\mu} = \left(10^3\,\frac{\text{kg}}{\text{m}^3}\right)\left(10^{-7}\,\frac{\text{m}}{\text{s}}\right)\left(2.5\times 10^{-8}\ \text{m}\right)\left(\frac{\text{m}^2}{0.003\,\text{N}\cdot\text{s}}\right) \approx 10^{-9}. \quad [302]$$

Thus, the coefficient of drag for the microtubule is

$$C_{\text{cyl}} = \frac{8\pi}{\frac{1}{2} - 0.577 - \ln(10^{-9}/8)} \approx \frac{24}{20} \approx 1, \quad [303]$$

as predicted in Chapter 12 of Physical Biology of the Cell (33). We can then find the force of drag with the formula (33),

$$F_{\text{cyl}} = C_{\text{cyl}}\mu L v, \quad [304]$$

where $L$ is the length of the microtubule. Thus, we find the drag on a cylinder perpendicular to the flow as

$$P_{\text{MT}} = F_{\text{cyl}} v = C_{\text{cyl}}\mu L v = 1\left(3\times 10^{-3}\,\frac{\text{N}\cdot\text{s}}{\text{m}^2}\right)\left(10^{-6}\,\text{m}\right)\left(10^{-7}\,\frac{\text{m}}{\text{s}}\right)^2 \approx 0.03\,\text{pN}\cdot\text{nm} \quad [305]$$

Taking $k_{\text{B}}T = 4\,\text{pN}\cdot\text{nm}$ and $\text{ATP} = 20\,k_{\text{B}}T$,

$$P_{\text{per MT}} = 4\times 10^{-4}\,\frac{\text{ATP}}{\text{s}\cdot\text{MT}}. \quad [306]$$

Finally, calculating the power of dragging all microtubules, we find

$$P = P_{\text{per MT}} \times N_{\text{MT}} = \left(4\times 10^{-4}\,\frac{\text{ATP}}{\text{s}\cdot\text{MT}}\right)\left(2\times 10^6\ \text{MT}\right) \approx 10^3\,\frac{\text{ATP}}{\text{s}}. \quad [307]$$

This value is an order of magnitude smaller than the sphere with a microtubule length diameter calculation, and the same order of magnitude as the sphere with matching surface area calculation. However, despite these calculations, the estimated power of microtubule drag is still many orders of magnitude smaller than the measured power.

**F. Nematic Order Parameter.** Here, we attempt to find the free energy of organizing a two dimensional aster using the order parameter, **Q**. We denote the orientation of each microtubule as

$$\mathbf{u} = (\cos\theta,\, \sin\theta) \quad [308]$$

and the order tensor as

$$Q_{ij} = \langle u_i u_j\rangle - \frac{1}{2}\delta_{ij} = \left\langle \begin{pmatrix} \cos^2\theta - \frac{1}{2} & \cos\theta\sin\theta \\ \cos\theta\sin\theta & \sin^2\theta - \frac{1}{2}\end{pmatrix}\right\rangle. \quad [309]$$

We assert that the aster is perfectly ordered, as locally all the microtubules point at the same $\theta$. This eliminates the average surrounding the order parameter resulting in,

$$Q_{ij} = \begin{pmatrix} \cos^2\theta - \frac{1}{2} & \cos\theta\sin\theta \\ \cos\theta\sin\theta & \sin^2\theta - \frac{1}{2}\end{pmatrix}. \quad [310]$$

In the form written, $Q_{ij}$ is in Cartesian coordinates, however the matrix elements are written in terms of $\theta$, a polar coordinate. We write the elements in terms of Cartesian elements, where $\cos\theta = \frac{x}{\sqrt{x^2+y^2}}$ and $\sin\theta = \frac{y}{\sqrt{x^2+y^2}}$,

$$Q_{ij} = \begin{pmatrix} \frac{x^2}{x^2+y^2} - \frac{1}{2} & \frac{xy}{x^2+y^2} \\ \frac{xy}{x^2+y^2} & \frac{y^2}{x^2+y^2} - \frac{1}{2}\end{pmatrix}. \quad [311]$$

Note that this tensor is both traceless,

$$Q_{11} + Q_{22} = \frac{x^2}{x^2+y^2} - \frac{1}{2} + \frac{y^2}{x^2+y^2} - \frac{1}{2} = 0$$
$$\Rightarrow Q_{11} = -Q_{22}, \qquad [312]$$

and symmetric,

$$Q_{ij} = \begin{pmatrix} Q_{11} & Q_{12} \\ Q_{21} & Q_{22} \end{pmatrix} = \begin{pmatrix} Q_{11} & Q_{21} \\ Q_{12} & Q_{22} \end{pmatrix} = Q_{ji} \qquad [313]$$

given that,

$$Q_{12} = \frac{xy}{x^2+y^2} = Q_{21}. \qquad [314]$$

***F.1. Free Energy Density.*** With the nematic order parameter in toe, we can now compute the free energy density gradient using the Landau-Ginsberg theory. According to Julia Yeomans' lecture notes (65), there are two components to the free energy density: the bulk free energy density, which follows the Landau-de Gennes expression,

$$F_{\text{bulk}} = \frac{A}{2}(Q_{ij}Q_{ji}) + \frac{B}{3}(Q_{ij}Q_{jk}Q_{ki}) + \frac{C}{4}(Q_{ij}Q_{ji})^2, \qquad [315]$$

and the elastic free energy density, which in 2D is,

$$F_{\text{el}} = \underbrace{\frac{L_1}{2}(\partial_k Q_{ij})^2}_{\text{splay}} + \underbrace{\frac{L_3}{2}Q_{ij}(\partial_i Q_{kl})(\partial_j Q_{kl})}_{\text{bend}}. \qquad [316]$$

All together, the free energy density in terms of $Q_{ij}$ is,

$$F = F_{\text{bulk}} + F_{\text{el}} = \frac{A}{2}(Q_{ij}Q_{ji}) + \frac{B}{3}(Q_{ij}Q_{jk}Q_{ki}) + \frac{C}{4}(Q_{ij}Q_{ji})^2 + \cdots$$
$$+ \frac{L_1}{2}(\partial_k Q_{ij})^2 + \frac{L_3}{2}Q_{ij}(\partial_i Q_{kl})(\partial_j Q_{kl}) + \cdots. \qquad [317]$$

Here, we will evaluate each listed term. We begin with the power series and examine up to fourth order. First expanding the summation notation of the quadratic term, we find

$$\begin{aligned} Q_{ij}Q_{ji} &= \sum_j \sum_i Q_{ij}Q_{ji} \\ &= \sum_j (Q_{1j}Q_{j1} + Q_{2j}Q_{j2}) \\ &= Q_{11}Q_{11} + Q_{21}Q_{12} + Q_{12}Q_{21} + Q_{22}Q_{22}. \end{aligned} \qquad [318]$$

Due to the tracelessness of $Q_{ij}$, $Q_{11}^2 = Q_{22}^2$, and due to the matrix being symmetric, $Q_{12} = Q_{21}$, thus, we can simplify the result to

$$Q_{ij}Q_{ji} = 2Q_{11}^2 + 2Q_{12}^2. \qquad [319]$$

Plugging in the matrix elements of equation 311, the quadratic term evaluates to

$$\begin{aligned} Q_{ij}Q_{ji} &= 2\left(\frac{x^2}{x^2+y^2} - \frac{1}{2}\right)^2 + 2\left(\frac{xy}{x^2+y^2}\right)^2 \\ &= 2\left(\frac{x^4}{(x^2+y^2)^2} - \frac{x^2}{x^2+y^2} + \frac{1}{4} + \frac{x^2y^2}{(x^2+y^2)^2}\right) \\ &= 2\left(\frac{x^4}{(x^2+y^2)^2} - \frac{x^4+x^2y^2}{(x^2+y^2)^2} + \frac{1}{4} + \frac{x^2y^2}{(x^2+y^2)^2}\right) \\ &= \frac{1}{2} \end{aligned} \qquad [320]$$

Let us now compute the cubic term,

$$\begin{aligned} Q_{ij}Q_{jk}Q_{ki} &= \sum_k \sum_j \sum_i Q_{ij}Q_{jk}Q_{ki} \\ &= \sum_k \sum_j (Q_{1j}Q_{jk}Q_{k1} + Q_{2j}Q_{jk}Q_{k2}) \\ &= \sum_k (Q_{11}Q_{1k}Q_{k1} + Q_{21}Q_{1k}Q_{k2} + Q_{12}Q_{2k}Q_{k1} + Q_{22}Q_{2k}Q_{k2}) \\ &= Q_{11}Q_{11}Q_{11} + Q_{21}Q_{11}Q_{12} + Q_{12}Q_{21}Q_{11} + Q_{22}Q_{21}Q_{12} \\ &\quad + Q_{11}Q_{12}Q_{21} + Q_{21}Q_{12}Q_{22} + Q_{12}Q_{22}Q_{21}Q_{22}Q_{22}Q_{22} \\ &= Q_{11}^3 + 3Q_{11}Q_{12}^2 + 3Q_{22}Q_{12}^2 + Q_{22}^3. \end{aligned} \quad [321]$$

By the symmetry of $Q_{ij}$,

$$Q_{ij}Q_{jk}Q_{ki} = Q_{11}^3 + 3Q_{11}Q_{12}^2 + 3Q_{22}Q_{12}^2 + Q_{22}^3, \quad [322]$$

and given that the matrix is traceless,

$$Q_{ij}Q_{jk}Q_{ki} = Q_{11}^3 + 3Q_{11}Q_{12}^2 - 3Q_{11}Q_{12}^2 + (-Q_{11})^3 = 0, \quad [323]$$

so the cubic term vanishes. We now compute the quartic term using the result from equation 319

$$\begin{aligned} (Q_{ij}Q_{ji})^2 &= \left( \sum_j \sum_i Q_{ij}Qji \right)^2 \\ &= \left( 2Q_{11}^2 + 2Q_{12}^2 \right)^2. \end{aligned} \quad [324]$$

Thus, the quartic term is simply the square of the quadratic term, which evaluates to

$$(Q_{ij}Q_{ji})^2 = \left(\frac{1}{2}\right)^2 = \frac{1}{4}. \quad [325]$$

We now compute the derivative terms. We write the partial first derivatives of $Q_{ij}$ as follows.

**First Derivatives:**

$$\frac{\partial}{\partial x}Q_{ij} = \begin{pmatrix} \partial_1 Q_{11} & \partial_1 Q_{12} \\ \partial_1 Q_{21} & \partial_1 Q_{22} \end{pmatrix} = \begin{pmatrix} \frac{2xy^2}{(x^2+y^2)^2} & \frac{y(-x^2+y^2)}{(x^2+y^2)^2} \\ \frac{y(-x^2+y^2)}{(x^2+y^2)^2} & -\frac{2xy^2}{(x^2+y^2)^2} \end{pmatrix} \quad [326]$$

$$\frac{\partial}{\partial y}Q_{ij} = \begin{pmatrix} \partial_2 Q_{11} & \partial_2 Q_{12} \\ \partial_2 Q_{21} & \partial_2 Q_{22} \end{pmatrix} = \begin{pmatrix} -\frac{2x^2y}{(x^2+y^2)^2} & \frac{x(x^2-y^2)}{(x^2+y^2)^2} \\ \frac{x(x^2-y^2)}{(x^2+y^2)^2} & \frac{2x^2y}{(x^2+y^2)^2} \end{pmatrix} \quad [327]$$

Note that the matrices representing the first derivatives are also traceless and symmetric, meaning $\partial_k Q_{11} = -\partial_k Q_{22}$ and $\partial_k Q_{12} = \partial_k Q_{21}$.

Computing the first term in the elastic free energy density, we find

$$\begin{aligned} (\partial_k Q_{ij})^2 &= \sum_k \sum_j \sum_i (\partial_k Q_{ij})^2 \\ &= \sum_k \sum_j (\partial_k Q_{1j})^2 + (\partial_k Q_{2j})^2 \\ &= \sum_k (\partial_k Q_{11})^2 + (\partial_k Q_{12})^2 + (\partial_k Q_{21})^2 + (\partial_k Q_{22})^2 \\ &= (\partial_1 Q_{11})^2 + (\partial_2 Q_{11})^2 + (\partial_1 Q_{12})^2 + (\partial_2 Q_{12})^2 + (\partial_1 Q_{21})^2 + (\partial_2 Q_{21})^2 + (\partial_1 Q_{22})^2 + (\partial_2 Q_{22})^2 \end{aligned} \quad [328]$$

By the properties $Q_{11} = -Q_{22}$, and $Q_{12} = Q_{21}$,

$$(\partial_k Q_{ij})^2 = 2(\partial_1 Q_{11})^2 + 2(\partial_2 Q_{11})^2 + 2(\partial_1 Q_{12})^2 + 2(\partial_2 Q_{12})^2. \quad [329]$$

Plugging in the matrix elements,

$$\begin{aligned} (\partial_k Q_{ij})^2 &= \frac{8x^2y^4}{(x^2+y^2)^4} + \frac{8x^4y^2}{(x^2+y^2)^4} + \frac{2y^2(-x^2+y^2)^2}{(x^2+y^2)^4} + \frac{2x^2(x^2-y)^2}{(x^2+y^2)^4} \\ &= \frac{2}{x^2+y^2} \end{aligned} \quad [330]$$

We now compute the second term in the elastic free energy density and use the simplifications that $Q_{ij}$ is symmetric and traceless,

$$\begin{aligned}
Q_{ij}(\partial_i Q_{kl})(\partial_j Q_{kl}) =& \sum_l \sum_k \sum_j \sum_i Q_{ij}(\partial_i Q_{kl})(\partial_j Q_{kl}) \\
=& \sum_l \sum_k Q_{11}(\partial_1 Q_{kl})^2 + 2Q_{12}(\partial_1 Q_{kl})(\partial_2 Q_{kl}) + Q_{22}(\partial_2 Q_{kl})^2 \\
=& Q_{11}(\partial_1 Q_{11})^2 + 2Q_{11}(\partial_1 Q_{12})^2 + Q_{11}(\partial_1 Q_{22})^2 \\
&+ 2Q_{12}(\partial_1 Q_{11})(\partial_2 Q_{11}) + 4Q_{12}(\partial_1 Q_{12})(\partial_2 Q_{12}) + 2Q_{12}(\partial_1 Q_{22})(\partial_2 Q_{22}) \\
&+ Q_{22}(\partial_2 Q_{11})^2 + 2Q_{22}(\partial_2 Q_{12})^2 + Q_{22}(\partial_2 Q_{22})^2 \\
=& 2Q_{11}(\partial_1 Q_{11})^2 + 2Q_{11}(\partial_1 Q_{12})^2 \\
&+ 4Q_{12}(\partial_1 Q_{11})(\partial_2 Q_{11}) + 4Q_{12}(\partial_1 Q_{12})(\partial_2 Q_{12}) \\
&- 2Q_{11}(\partial_2 Q_{11})^2 - 2Q_{11}(\partial_2 Q_{12})^2 \\
=& 2Q_{11}\Big[(\partial_1 Q_{11})^2 + (\partial_1 Q_{12})^2 - (\partial_2 Q_{11})^2 - (\partial_2 Q_{12})^2\Big] \\
&+ 4Q_{12}\Big[(\partial_1 Q_{11})(\partial_2 Q_{11}) + (\partial_1 Q_{12})(\partial_2 Q_{12})\Big]
\end{aligned}$$

[331]

Plugging in matrix values and simplifying with Mathematica, this term reduces to

$$Q_{ij}(\partial_i Q_{kl})(\partial_j Q_{kl}) = -\frac{1}{x^2+y^2}$$ [332]

Putting everything together, we reveal an equation for the free energy density throughout an ordered aster

$$F = \frac{A}{2}\left(\frac{1}{2}\right) + \frac{C}{4}\left(\frac{1}{4}\right) + \cdots + \frac{L_1}{2}\frac{2}{x^2+y^2} - \frac{L_3}{2}\frac{1}{x^2+y^2} + \cdots .$$ [333]

We can translate this back into polar coordinates using the transforms $r = \sqrt{x^2+y^2}$, $\cos\theta = \frac{x}{\sqrt{x^2+y^2}}$ and $\sin\theta = \frac{y}{\sqrt{x^2+y^2}}$

$$F = \frac{4A+C}{16} + \frac{2L_1 - L_3}{2}\frac{1}{r^2} + \cdots .$$ [334]

Now it is left to find the energy of the aster state by integrating the free energy density over the area of the aster. We integrate over all angles from $0 \leq \theta \leq 2\pi$ and radius values from a core cutoff, $R_c$ to the aster radius $R$,

$$\begin{aligned}
E &= \int_0^{2\pi}\int_{R_c}^{R} F\, r\, \mathrm{d}r\, \mathrm{d}\theta \\
&= \int_0^{2\pi}\int_{R_c}^{R} \left(\frac{4A+C}{16}r + \frac{2L_1 - L_3}{2r}\right) \mathrm{d}r\, \mathrm{d}\theta \\
&= \frac{4A+C}{8}\pi(R^2 - R_c^2) + (2L_1 - L_3)\pi \ln\left(\frac{R}{R_c}\right)
\end{aligned}$$ [335]

Now that we have solved the equation for the energy of the ordered state, we can input parameters to find the energy scale of perfect order.

***F.2. Order of Magnitude Estimate.*** To get a sense of scale of the energy in equation 335, we explore literature values for the constants in front of each term. Zhang et al. use modeling, simulations, and experiments to find how the elasticity of a liquid crystal (LC) depends on filament length, density, and rigidity. Their system, a thin film nematic of actin and microtubules, follows the same free energy density form as equation 334. In their study, they measure elastic constants, $K$, using elastic beam theory. They find that $L_1 = \frac{K}{2q_0^2}$, where $L_1$ is the coefficient of the $(\partial_k Q_{ij})^2$, same as us, and $q_0$ is the scalar order parameter. Thus, if we assume the $L_1$ term is at least of the same order as the other terms, we can get an order of magnitude estimate for the ordered energy of the aster. Since our aster is assumed to be perfectly ordered, we take $q_0 = 1$. Zhang et al. find $K \sim f \times 10^{-1}$ pN (66). Thus, $L_1 \approx 0.1$ pN. Note that for the constant $L_1$ to have units of piconewtons, the free energy density must be integrated over a volume rather than an area. Since their system was a very thin film of $\delta_z = 300$ nm, their system is still quasi 2D in comparison to their large film area $4\,\mathrm{mm}^2$. While the thickness of the aster is non-trivial, we will simply proceed with this estimate to find the energy scale of ordering. Thus, we will integrate over z, the thickness of the aster, in our estimate,

$$E = \frac{4A+C}{8}\pi z(R^2 - R_c^2) + (2L_1 - L_3)\pi z \ln\left(\frac{R}{R_c}\right).$$ [336]

Taking the aster radius to be $R \approx f \times 10\,\mu$m, the cutoff radius to be $R_c \approx 1\,\mu$m, and the thickness to be $z \approx 100\,\mu$m, we find the value of the $L_1$ term of the energy,

$$E \sim 2\pi L_1 z \ln\left(\frac{R}{R_c}\right) = 2\pi \times 10^{-1}\ \text{pN} \times 10^5\ \text{nm} \times \ln(f \times 10) \approx f \times 10^5\,\text{pN}\cdot\text{nm} \qquad [337]$$

With the conversion $\text{ATP} = 20\,k_\text{B}T\left(\frac{4\,\text{pN}\cdot\text{nm}}{k_\text{B}T}\right) = 80\,\text{pN}\cdot\text{nm}$, the free energy of the ordered aster is $f \times 10^3$ ATP. If an aster takes around 10 minutes to form, then the power of organization is

$$P = \frac{f \times 10^3\ \text{ATP}}{600\ \text{s}} = f\,\frac{\text{ATP}}{\text{s}}. \qquad [338]$$

This is VERY low in comparison with our measured power of $10^8\,\frac{\text{ATP}}{\text{s}}$. While other studies have found $K$ constant values as large as 10 pN (67), this would only increase our power by a factor of 100, still six orders of magnitude smaller than the measured value. Thus, the power of ordering microtubules appears to be negligible.

**G. Power to Maintain a Motor Gradient.** As noted in the main body of the paper, the largest power estimate out of the suite of processes we considered is associated with the formation of the gradient of motors. Because of the gradient in motor density across the aster, it is of interest to estimate the free energy required to maintain that gradient. We begin by examining the free energy cost of a one-dimensional gradient to set notation and to explain the concept and follow that discussion by the case of interest involving spherical symmetry. Here we follow nearly verbatim our earlier treatment of the one-dimensional problem in "The Restless Cell" (68). The concept of the estimate is to compute the free energy change when we take one particle from a region with one concentration and put that particle in a nearby region with a slightly different concentration.

The left panel of Figure S86 illustrates the free energy change we need to evaluate. In particular, we compute the free energy of the system before and after allowing a single molecule to cross from one side of the partition to the other. The free energy is defined as

$$F = U - TS, \qquad [339]$$

where $U$ is either the energy or the enthalpy and $S$ is the entropy. We ignore the subtleties associated with Helmholtz vs Gibbs free energies. We interest ourselves in the change in free energy

$$\Delta F = F_{final} - F_{initial} = (U_{final} - TS_{final}) - (U_{initial} - TS_{initial}). \qquad [340]$$

Further, we simplify by ignoring any contribution due to energy (or enthalpy) changes (i.e. $\Delta U = 0$).

To compute the entropy change, we need to compute the entropy of the solutions on both sides of the partition, both before and after we have plucked a molecule from the left side and installed it on the right side. The total entropy is given by

$$S_{tot} = S_1 + S_2 \qquad [341]$$

where the subscripts refer to the two compartments. This expression implies that the free energy in this state is given by

$$F = -TS_{tot}. \qquad [342]$$

We can compute the entropy by counting the number of microscopic states both before and after the particle transfer. In particular, we have

$$S_{tot}^{(final)} = k_B\ \ln W_1^{(final)} + k_B\ \ln W_2^{(final)}, \qquad [343]$$

with a similar expression for the entropy in the initial state. This can be simplified to the form

$$S_{tot}^{(final)} = k_B\ \ln\,(W_1^{(final)} W_2^{(final)}), \qquad [344]$$

which makes sense given that the total number of microscopic states is equal to the product of the number of states in the first box and the number of states in the second box.

In a lattice model, we make the abstraction that space is subdivided into tiny lattice sites with a characteristic dimension of 1 nm$^3$ (i.e. molecular sizes). To compute the number of microstates $W$, we count the number of ways of arranging our $L_i$ motors among the $\Omega$ lattice sites as

$$W_i(L_i) = \frac{\Omega!}{L_i!(\Omega - L_i)!},. \qquad [345]$$

When $\Omega \gg L_i$ (i.e. the dilute limit), we can make the much simpler approximation

$$W_i(L) = \frac{\Omega^{L_i}}{L_i!}, \qquad [346]$$

which amounts to the idea that every motor can sit on any of the $\Omega$ lattice sites.

We can now write the change in free energy which is strictly entropic as

$$\Delta F = -k_B T\left(\ \ln\,\frac{\Omega^{L_1+1}}{(L_1+1)!}\frac{\Omega^{L_2-1}}{L_2-1)!} -\ \ln\,\frac{\Omega^{L_1}}{L_1!}\frac{\Omega^{L_2}}{L_2!}\right). \qquad [347]$$

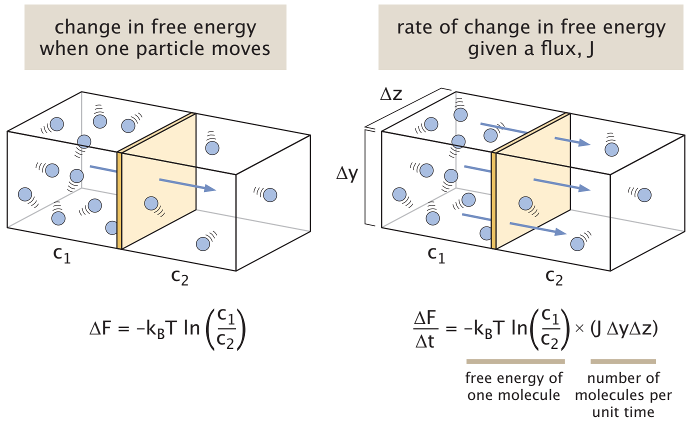

**Fig. S86. The change in free energy when a particle is transported in a gradient.** The left panel shows the free energy change upon moving a particle from the left to the right.

This can be simplified to

$$\Delta F = -k_B T \ \ln \ \frac{L_1!}{(L_1+1)!}\frac{L_2!}{(L_2-1)!} \approx -k_B T \ \ln \ \frac{L_2}{L_1}. \quad [348]$$

We can rewrite this in a more familiar form using the language of concentrations. If we multiply numerator and denominator within the logarithm by $\Omega v$, where $v$ is the volume of a single lattice site in our lattice model, then $\Omega v = V_{tot}$ and hence $c_1 = L_1/V_{tot}$ and $c_2 = L_2/V_{tot}$, permitting us to write

$$\Delta F = k_B T \ \ln \ \frac{c_2}{c_1}. \quad [349]$$

Now, as seen in the right panel of Figure S86 if we want to find the total rate of free energy change, we need to multiply the free energy per particle by the total number of particles transported between the two adjacent reservoirs per unit time using the flux resulting in

$$\left(\frac{\Delta F}{\Delta t}\right) = -\Delta\mu(J\Delta y\Delta z). \quad [350]$$

The factor $J\Delta y\Delta z$ counts the number of molecules carried down the gradient per unit time and the minus sign guarantees that if the left reservoir has more molecules than the left, then $\Delta F/\Delta t < 0$. We now interest ourselves in the case where the concentration is slowly varying, permitting us to write

$$\left(\frac{\Delta F}{\Delta t}\right) = k_B T \ln \frac{c(x+\Delta x)}{c(x)} \times (J\Delta y\Delta z), \quad [351]$$

where we have introduced the notation $c_1 = c(x)$ and $c_2 = c(x+\Delta x)$. Note now we switched the sign because we inverted the ratio in the logarithm with the concentration on the right now appearing in the numerator. Note that this expression is valid regardless of whether $c_1 > c_2$ or $c_2 > c_1$ since in those two cases the flux is in opposite directions and our expression reflects that. By invoking Fick's law we now have

$$\left(\frac{\Delta F}{\Delta t}\right) = k_B T \ln \frac{c(x+\Delta x)}{c(x)} \times \left(-D\frac{dc}{dx}\right)\Delta y\Delta z. \quad [352]$$

By carrying out the Taylor expansion $c(x+\Delta x) = c(x) + (\partial c/\partial x)\Delta x$, we can rewrite the logarithmic term as

$$\begin{aligned} \ln \frac{c(x+\Delta x)}{c(x)} &= \ln \left[\frac{c(x) + \frac{\partial c}{\partial x}\Delta x}{c(x)}\right] \\ &= \ln \left[1 + \frac{1}{c}\frac{\partial c}{\partial x}\Delta x\right]. \end{aligned} \quad [353]$$

Finally, we invoke a second Taylor series in the form $\ln \ (1+\epsilon) \approx \epsilon$, resulting in the simple and powerful expression

$$\left(\frac{\Delta F}{\Delta t}\right) = -Dk_B T \frac{1}{c(x)}\left(\frac{\partial c(x)}{\partial x}\right)^2 \Delta x\Delta y\Delta z. \quad [354]$$

This result tells us the free energy loss for two adjacent planes if the gradient is allowed to dissipate. We can now put this all together to explore the free energy dissipated over a continuous concentration field. In this case, we take the expression seen in eqn. 354 and add up the contribution from each set of planes in our discrete representation of the concentration field. We can work out the minimum power to maintain such a gradient where we imagine every time a molecule goes down its gradient, energy is consumed to push it back where it came from.

Building on this one-dimensional analysis, we now turn our attention to a three-dimensional structure with a spherically symmetric concentration gradient of motors of the form $c(r)$, where $r$ is the radial distance from the aster center. Figure S87(A) shows the amendment that needs to be made to our one-dimensional analysis, where now in the three dimensional case, the "boxes" are spherical shells. Interestingly, the expression we derived earlier goes through essentially unchanged except that now we consider the radial concentration, and instead of integrating over planes along the x-direction, we now integrate over spherical shells in the $r$ direction. Given these adjustments, the power to sustain a gradient is now given as

$$P = k_{\mathrm{B}}TD \int_0^{2\pi}\int_0^{\pi}\int_0^{\infty} \frac{1}{c(r)}\left(\frac{\partial c(r)}{\partial r}\right)^2 r^2 \sin\theta \, dr \, d\theta \, d\phi, \quad [355]$$

where $D$ is the diffusion constant and $c(r)$ is the concentration profile.

To get a qualitative feeling for the power scales, we begin by considering a fixed, radial distribution of motors described as a decaying exponential of the form

$$c(r) = c_0 e^{-r/\lambda}, \quad [356]$$

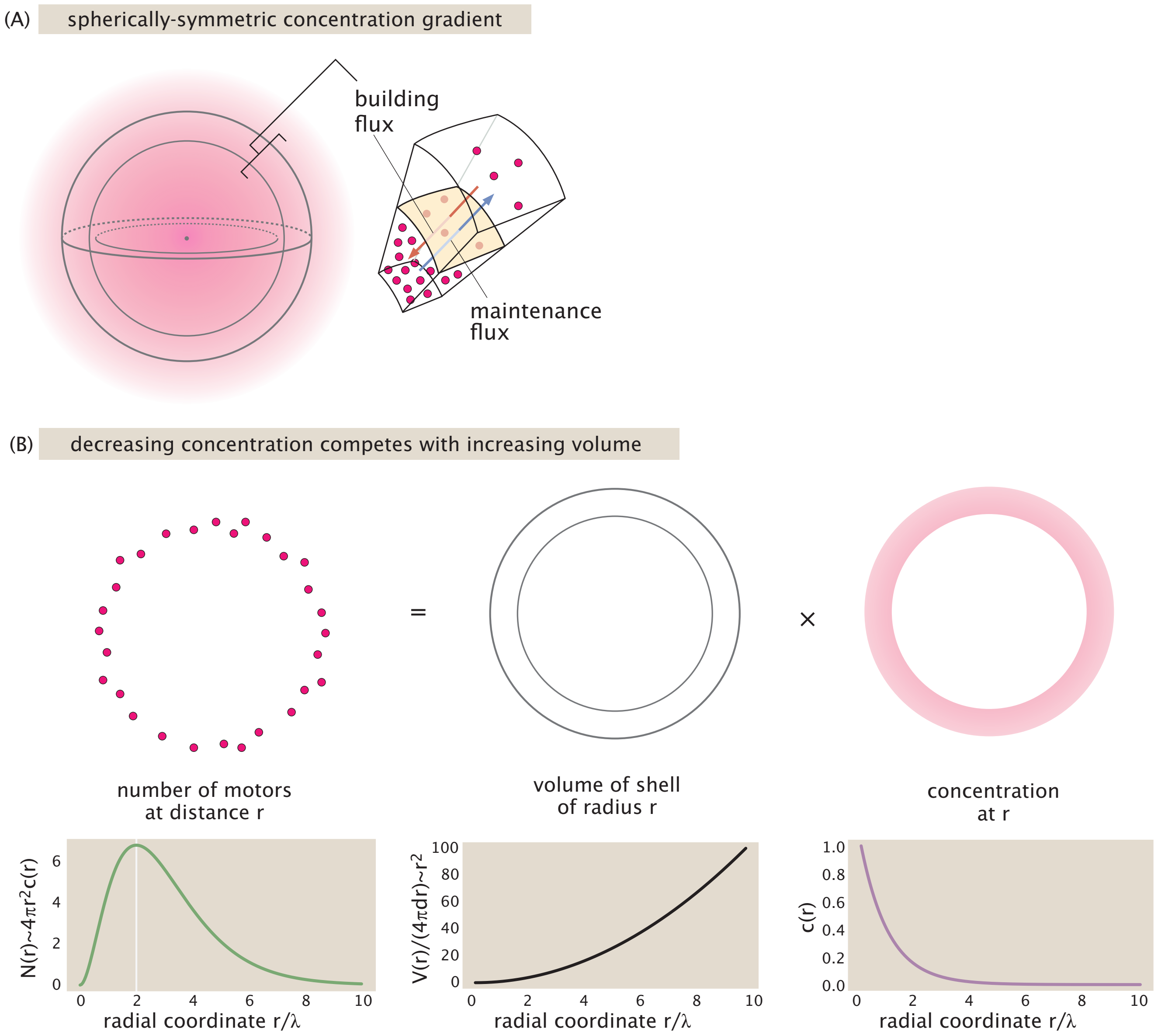

**Fig. S87.** Dissipation in a spherically symmetric concentration gradient. (A) Maintaining a spherically symmetric concentration gradient results from a competition between the outward diffusive flux and the inward active flux provided by motors moving on microtubules. (B) There is a peak in the distribution of motors as a function of radial distance resulting from the competition between the monotonically decreasing motor concentration and the geometric effect that the spherical shells get larger with increasing $r$.

where $c_0$ is the motor concentration at $r = 0$ and $\lambda$ is the decay length. Plugging this profile into eqn. 355, we can immediately compute the power that must be expended to prevent the diffusive relaxation of this gradient as

$$\begin{aligned} P &= 4\pi k_B T D \int_0^\infty \frac{e^{r/\lambda}}{c_0} \left(-\frac{c_0}{\lambda} e^{-r/\lambda}\right)^2 r^2\, dr \\ &= \frac{4\pi k_B T D c_0}{\lambda^2} \int_0^\infty r^2\, e^{-r/\lambda} dr. \end{aligned} \quad [357]$$

Integrating by parts, we find that the integral evaluates to $\lambda^2 \int_0^\infty r^2\, e^{-r/\lambda} dr = 2\lambda^3$ yielding,

$$P = 8\pi k_B T D c_0 \lambda. \quad [358]$$

With this equation in hand, we estimate the magnitude of power required to prevent diffusive smoothing of gradients. We take an aster radius of $60\,\mu$m as the value for $\lambda$, and the maximum motor concentration $c_0 = 3\,\mu$M as the $r = 0$ concentration. We use the size of a motor protein to estimate the diffusion constant. Our motors are approximately 100 kDa in weight. The molecular weight of a protein scales approximately linearly with its volume (51). We make the simplifying assumption that our motor proteins have a "radius" of 3 nm. Given the motor protein radius, we can compute the diffusion constant for motors using the Stokes-Einstein equation, where we approximate our reaction buffer to be a 30% glycerol solution with a viscosity of $2.4 \times 10^{-3}\,\mathrm{N \cdot s/m^2}$ (11),

$$D = \frac{k_B T}{6\pi\mu r} = \frac{4\,\mathrm{pN \cdot nm}}{6\pi \times (2.4 \times 10^{-3}\,\mathrm{pN \cdot s/\mu m^2}) \times (3\ \mathrm{nm})} \approx 30 \frac{\mu\mathrm{m}^2}{\mathrm{s}}. \quad [359]$$

Plugging in all these constants into equation 358, we estimate the power to prevent diffusive spreading as,

$$P = 8\pi k_B T \left(30\ \frac{\mu\mathrm{m}^2}{\mathrm{s}}\right)(3\,\mu\mathrm{M})(60\,\mu\mathrm{m}) \approx 10^8\ \frac{k_B T}{\mathrm{s}} \approx 5 \times 10^6\ \frac{\mathrm{ATP}}{\mathrm{s}}. \quad [360]$$

This is the greatest power we have estimated. As seen from examining the powers across asters in Figure S32, at late times, the power is $\approx f \times 10^7\ \frac{\mathrm{ATP}}{\mathrm{s}}$ about an order of magnitude larger than our estimate. Thus, we expect the power to maintain a motor gradient could be a significant fraction of the energetic cost on the motor-microtubule system.

#### References

1. SJ DeCamp, Dogic lab - acrylamide coating protocol (Web Page) (2016) Accessed April 15, 2026.
2. S Hirokawa, et al., Motor-driven microtubule diffusion in a photobleached dynamical coordinate system. *Proc. Natl. Acad. Sci.* **122**, e2417020122 (2025).
3. H Yaginuma, Y Okada, Live cell imaging of metabolic heterogeneity by quantitative fluorescent ATP indicator protein, QUEEN-37c. *bioRxiv* (2021).
4. H Yaginuma, et al., Diversity in atp concentrations in a single bacterial cell population revealed by quantitative single-cell imaging. *Sci. reports* **4**, 6522 (2014).
5. B Najma, WS Wei, A Baskaran, PJ Foster, G Duclos, Microscopic interactions control a structural transition in active mixtures of microtubules and molecular motors. *Proc. Natl. Acad. Sci.* **121**, e2300174121 (2024).
6. T Photometrics, Camera Test Protocol (https://www.photometrics.com/learn/camera-test-protocol) (2019) Accessed: (2024).
7. N Dey, Uneven illumination correction of digital images: A survey of the state-of-the-art. *Optik* **183**, 483–495 (2019).
8. HG Garcia, R Phillips, Quantitative dissection of the simple repression input–output function. *Proc. Natl. Acad. Sci.* **108**, 12173–12178 (2011).
9. F Van Breugel, JN Kutz, BW Brunton, Numerical differentiation of noisy data: A unifying multi-objective optimization framework. *IEEE Access* **8**, 196865–196877 (2020).
10. WR Schief, RH Clark, AH Crevenna, J Howard, Inhibition of kinesin motility by adp and phosphate supports a hand-over-hand mechanism. *Proc. Natl. Acad. Sci.* **101**, 1183–1188 (2004).
11. C Westbrook, Calculate density and viscosity of glycerol/water mixtures (2018) Accessed: 2024.
12. J Howard, *Mechanics of motor proteins and the cytoskeleton.* (Sunderland, Mass. : Sinauer Associates, Publishers), (2001).
13. RA Walker, ED Salmon, SA Endow, The drosophila claret segregation protein is a minus-end directed motor molecule. *Nature* **347**, 780–782 (1990).
14. K Furuta, YY Toyoshima, Minus-end-directed motor ncd exhibits processive movement that is enhanced by microtubule bundling in vitro. *Curr. Biol.* **18**, 152–157 (2008).
15. K Furuta, et al., Measuring collective transport by defined numbers of processive and nonprocessive kinesin motors. *Proc. Natl. Acad. Sci., USA110* **110**, 501–506 (2012).
16. RA Banks, et al., Motor processivity and speed determine structure and dynamics of microtubule-motor assemblies. *Elife* **12**, e79402 (2023).
17. MJ deCastro, CH Ho, RJ Stewart, Motility of dimeric ncd on a metal-chelating surfactant: evidence that ncd is not processive. *Biochemistry* **38**, 5076–5081 (1999).
18. HB McDonald, RJ Stewart, LS Goldstein, The kinesin-like ncd protein of drosophila is a minus end-directed microtubule motor. *Cell* **63**, 1159–1165 (1990).

19. KA Foster, JJ Correia, SP Gilbert, Equilibrium binding studies of non-claret disjunctional protein (ncd) reveal cooperative interactions between the motor domains. *J. Biol. Chem.* **273**, 35307–35318 (1998).
20. M Lab, Tubulin basics (Web Page) (Accessed 2026) https://mitchison.hms.harvard.edu/resources.
21. DL Jones, RC Brewster, R Phillips, Promoter architecture dictates cell-to-cell variability in gene expression. *Science* **346**, 1533–6 (2014).
22. RW Pan, T Röschinger, K Faizi, HG Garcia, R Phillips, Deciphering regulatory architectures of bacterial promoters from synthetic expression patterns. *PLOS Comput. Biol.* **20**, e1012697 (2024).
23. LM Smith, DR Keefer, S Sudharsanan, Abel inversion using transform techniques. *J. Quant. Spectrosc. Radiat. Transf.* **39**, 367–373 (1988).
24. S Gibson, et al., Pyabel/pyabel: v0.9.0 (2022).
25. D Sage, et al., Deconvolutionlab2: An open-source software for deconvolution microscopy. *Methods* **115**, 28–41 (2017).
26. M Born, E Wolf, *Principles of optics: electromagnetic theory of propagation, interference and diffraction of light.* (Elsevier), (2013).
27. E Hecht, *Optics.* (Addison Wesley, San Francisco), 4th edition, (2003).
28. H Kirshner, F Aguet, D Sage, M Unser, 3-d psf fitting for fluorescence microscopy: implementation and localization application. *J. microscopy* **249**, 13–25 (2013).
29. S Stallinga, B Rieger, Accuracy of the gaussian point spread function model in 2d localization microscopy. *Opt. express* **18**, 24461–24476 (2010).
30. B Zhang, J Zerubia, JC Olivo-Marin, Gaussian approximations of fluorescence microscope point-spread function models. *Appl. optics* **46**, 1819–1829 (2007).
31. M Prato, R Cavicchioli, L Zanni, P Boccacci, M Bertero, Efficient deconvolution methods for astronomical imaging: algorithms and idl-gpu codes. *Astron. & Astrophys.* **539**, A133 (2012).
32. F Aguet, D Van De Ville, M Unser, Model-based 2.5-d deconvolution for extended depth of field in brightfield microscopy. *IEEE Transactions on Image Process.* **17**, 1144–1153 (2008).
33. R Phillips, J Kondev, J Theriot, HG Garcia, N Orme, *Physical biology of the cell.* (Garland Science), (2012).
34. DL Coy, M Wagenbach, J Howard, Kinesin takes one 8-nm step for each ATP that it hydrolyzes. *J. Biol. Chem.* **274**, 3667–3671 (1999).
35. H Imamura, et al., Visualization of ATP levels inside single living cells with fluorescence resonance energy transfer-based genetically encoded indicators. *Proc. Natl. Acad. Sci.* **106**, 15651–15656 (2009).
36. G Tkačik, JO Dubuis, MD Petkova, T Gregor, Positional information, positional error, and readout precision in morphogenesis: a mathematical framework. *Genetics* **199**, 39–59 (2015).
37. RG Endres, NS Wingreen, Accuracy of direct gradient sensing by single cells. *Proc. Natl. Acad. Sci.* **105**, 15749–15754 (2008).
38. V Sourjik, HC Berg, Binding of the escherichia coli response regulator chey to its target measured in vivo by fluorescence resonance energy transfer. *Proc. Natl. Acad. Sci.* **99**, 12669–12674 (2002).
39. V Sourjik, HC Berg, Receptor sensitivity in bacterial chemotaxis. *Proc. Natl. Acad. Sci.* **99**, 123–127 (2002).
40. AJ Waite, NW Frankel, T Emonet, Behavioral variability and phenotypic diversity in bacterial chemotaxis. *Annu. review biophysics* **47**, 595–616 (2018).
41. JQ Wu, TD Pollard, Counting cytokinesis proteins globally and locally in fission yeast. *Science* **310**, 310–314 (2005).
42. JB Moseley, A Mayeux, A Paoletti, P Nurse, A spatial gradient coordinates cell size and mitotic entry in fission yeast. *Nature* **459**, 857–860 (2009).
43. PlotDigitizer: Version 3.1.6 (2026) https://plotdigitizer.com.
44. T Gregor, DW Tank, EF Wieschaus, W Bialek, Probing the limits to positional information. *Cell* **130**, 153–164 (2007).
45. JB Perrin, Discontinuous structure of matter (year?).
46. H Wackerhage, et al., Recovery of free adp, pi, and free energy of atp hydrolysis in human skeletal muscle. *J. applied physiology* **85**, 2140–2145 (1998).
47. QH Tran, G Unden, Changes in the proton potential and the cellular energetics of escherichia coli during growth by aerobic and anaerobic respiration or by fermentation. *Eur. journal biochemistry* **251**, 538–543 (1998).
48. PJ Foster, et al., Dissipation and energy propagation across scales in an active cytoskeletal material. *Proc. Natl. Acad. Sci.* **120**, e2207662120 (2023).
49. AH Stouthamer, Theoretical Study on Amount of Atp Required for Synthesis of Microbial Cell Material. *Antonie Van Leeuwenhoek J. Microbiol.* **39**, 545–565 (1973).
50. M Mori, C Cheng, BR Taylor, H Okano, T Hwa, Functional decomposition of metabolism allows a system-level quantification of fluxes and protein allocation towards specific metabolic functions. *Nat. Commun.* **14**, 4161 (2023).
51. R Milo, R Phillips, *Cell biology by the numbers.* (Garland Science), (2015).
52. G Salmon, PhD thesis (California Institute of Technology, Pasadena, CA) (2025).
53. J Kondev, M Kirschner, HG Garcia, GL Salmon, R Phillips, Biological processes as exploratory dynamics. *Biophys. J.* **125** (2025).
54. M Lynch, GK Marinov, The bioenergetic costs of a gene. *Proc. Natl. Acad. Sci. United States Am.* **112**, 15690–15695 (2015).
55. SJ Schink, E Biselli, C Ammar, U Gerland, Death rate of e. coli during starvation is set by maintenance cost and biomass

recycling. *Cell systems* **9**, 64–73 (2019).

56. R Milo, P Jorgensen, U Moran, G Weber, M Springer, Bionumbers—the database of key numbers in molecular and cell biology. *Nucleic acids research* **38**, D750–D753 (2010).
57. JA Ciemniecki, CL Ho, RD Horak, A Okamoto, DK Newman, Mechanistic study of a low-power bacterial maintenance state using high-throughput electrochemistry. *Cell* **187**, 6882–6895 (2024).
58. X Yang, G Ha, DJ Needleman, A coarse-grained nadh redox model enables inference of subcellular metabolic fluxes from fluorescence lifetime imaging. *Elife* **10**, e73808 (2021).
59. PC Hinkle, P/o ratios of mitochondrial oxidative phosphorylation. *Biochimica et Biophys. Acta (BBA)-Bioenergetics* **1706**, 1–11 (2005).
60. J Rodenfels, KM Neugebauer, J Howard, Heat oscillations driven by the embryonic cell cycle reveal the energetic costs of signaling. *Dev. cell* **48**, 646–658 (2019).
61. JJ Hopfield, Kinetic proofreading: a new mechanism for reducing errors in biosynthetic processes requiring high specificity. *Proc Natl Acad Sci U S A* **71**, 4135–9 (1974).
62. TL Hill, MW Kirschner, Bioenergetics and kinetics of microtubule and actin filament assembly-disassembly. *Int Rev Cytol* **78**, 1–125 (1982).
63. TJ Mitchison, MW Kirschner, Properties of the Kinetochore In vitro. II. Microtubule Capture and ATP-Dependent Translocation. *J. Cell Biol.* **101**, 766–777 (1985).
64. R Finn, Determination of the drag on a cylinder at low reynolds numbers. *J. Appl. Phys.* **24**, 771–773 (1953).
65. JM Yeomans, The hydrodynamics of active systems. *La Rivista del Nuovo Cimento* **40**, 1–31 (2017).
66. R Zhang, N Kumar, JL Ross, ML Gardel, JJ De Pablo, Interplay of structure, elasticity, and dynamics in actin-based nematic materials. *Proc. Natl. Acad. Sci.* **115**, E124–E133 (2018).
67. R Zhang, Y Zhou, M Rahimi, JJ De Pablo, Dynamic structure of active nematic shells. *Nat. communications* **7**, 13483 (2016).
68. C Hueschen, R Phillips, *The Restless Cell: Continuum Theories of Living Matter.* (Princeton University Press), (2024).
